# No Ramp Needed: Spandrels, Statistics, and a Slippery Slope

**DOI:** 10.1101/2022.06.27.497802

**Authors:** Richard Sejour, Janet Leatherwood, Alisa Yurovsky, Bruce Futcher

## Abstract

Previously, Tuller et al. found that the first 30 to 50 codons of the genes of yeast and other eukaryotes are slightly enriched for rare codons, so are presumably translated somewhat slowly. They argued, based on informatics, that this initial slow translation “ramp” was adaptive; and increased efficiency of translation by queuing ribosomes to prevent collisions. Today, the translational speeds of different codons are known, and indeed rare codons are translated slowly. We re-examined the slow translation ramp. We confirm the finding that 5’ regions are enriched for rare codons. However, we also find that the 5’ ends of yeast genes are poorly conserved in evolution, suggesting that they are unstable and turn over relatively rapidly. When a new 5’ end forms *de novo*, it is likely to include codons that would otherwise be rare. Because evolution has had a relatively short time to select against these codons, 5’ ends are typically slightly enriched for rare, slow codons. Opposite to the expectation of Tuller et al., we show by direct experiment that genes with slowly translated codons at the 5’ end are expressed relatively poorly, and substituting faster codons improves expression. Further informatic studies suggest that for natural genes, slow 5’ ends are correlated with poor gene expression, opposite to the expectation of Tuller et al. Thus we conclude that slow 5’ translation is a “spandrel”; it is a non-adaptive consequence of something else, in this case the turnover of 5’ ends in evolution, and it does not improve translation.

**Highlight:** The 5’ ends of yeast genes are unstable over evolutionary time, enriching for rare codons, slowing translation; slow initial translation does not enhance expression.

## Introduction

Tuller et al. ^1^ were interested in the idea that a slow translational ramp at the beginning of a gene might queue ribosomes in an orderly way, thereby preventing ribosome traffic jams and collisions. However, at that time, translation speeds for the 61 sense codons were not known from direct measurement. Therefore, as a proxy for codon translation speed, Tuller et al. devised a proxy speed measurement based on the tRNA-adaptation index (tAI), a measure of the abundance of each tRNA. The assumption is that codons recognized by more abundant tRNAs would be translated faster. Using this proxy, Tuller et al. found that in yeast and other eukaryotes, the first 30 to 100 codons are enriched for codons for which tRNAs are rare (generally, rare codons), and are presumably translated slowly. The size of the effect is small (about a 3% difference, Tuller et al. Figure 2C), but is statistically significant.

At the time of the work of Tuller et al., ribosome profiling had recently been developed^2^, and early ribosome profiling results showed a very high density of ribosomes near the 5’ end of the mRNA, consistent with very slow translation in this region, and the rare codons found by Tuller et al. could have contributed to this. However, later work showed that this 5’ high density of ribosomes was an artefact of the way cycloheximide was used to arrest translation in the original protocol^3^. With newer protocols for ribosome profiling, which use cycloheximide only at later steps, the region of 5’ high ribosome density largely, but not entirely, disappears^3^ (see Discussion).

Since then, many workers have used ribosome profiling to directly measure the speed of translation of individual codons (cited below). With such data in hand, we re-visited the issues addressed by Tuller et al. ^1^. On the one hand, our analyses confirm that the 5’ regions of genes are typically slightly enriched for rare codons, and these encodings likely slow translation. On the other hand, various aspects of the data led us to an alternative hypothesis, namely that the 5’ ends were turning over relatively rapidly in evolution; that these 5’ ends were therefore relatively young; and that selection had not yet succeeded in removing all the rare, slow codons initially present in the *de novo* 5’ ends. We did a direct experimental test of the effects of slow or fast initial translation. Exactly opposite to Tuller et al., we found that encoding slow initial translation resulted in lower protein production than fast initial translation. This continued to be true even when we placed ribosome collision sites inside the reporter gene. Thus a slow initial translation ramp, though present, does not improve gene expression.

It is natural to assume that the enrichment of slow codons near 5’ ends is a product of selection. However, as elegantly argued by Gould and Lewontin^4^ in their classic paper “The Spandrels of San Marco and the Panglossian Paradigm: A Critique of the Adaptionist Programme”, not all biological phenomena are adaptive, or even a direct product of selection. They argued from the example of a “spandrel”: in architecture, a triangular space created when an arch supports a lintel. There is no architectural role for spandrels as such; they are the indirect and inevitable result of the juxtaposition of two other functional architectural elements. We argue that the slightly slow initial translation of eukaryotic genes may likewise be a spandrel, a non-adaptive consequence of something else, the instability of 5’ ends in evolution.

## Results

### Calculations of encoded translation speed imply slow initial translation

Tuller et al. ^1^ used codon-specific tRNA abundance as a proxy to estimate speed of translation of codons. Since then, analysis of ribosome profiling data has yielded direct measurements of the translation speed of individual codons^3, 5–11^. Accordingly, we have repeated some of the work of Tuller et al., but using the Ribosome Residence Time (RRT, Methods and Materials, Table S1) ^6^ of each of the 61 sense codons as a measure of translation speed. We refer to “encoded translation speed” to specify that we are focusing purely on the effects of different codons on translation speed, and not on other factors that might differentially affect translation speed at different regions of the mRNA, such as secondary structure.

Using the RRT, encoded translation speeds were calculated in sliding windows across all coding ORFs from *S. cerevisiae*. (The start codon was omitted because it is constant across genes and is an unusually “fast” codon.) Consistent with Tuller et al., we find that the first 30-100 codons had lower calculated translation speeds than the rest of the gene (Figure 1). We focused our analyses on the first 40 codons for comparability to Tuller et al.; we call this the “Slow Initial Translation” region, or SIT. Although the tendency towards slow translation near the beginning of the gene is very highly statistically significant, the size of the effect is small. When we compare the first 40 codons of a gene to the rest of the same gene, we find that the average difference in encoded translation speed is about 1.2% (similar to ^1^), with a p-value < 0.001. For comparison, codons can vary in translation speed by about two-fold (Table S1), or perhaps as much as 6-fold (Weinberg et al., their Table S2, also shown in our Table S1).

**Figure 1.**
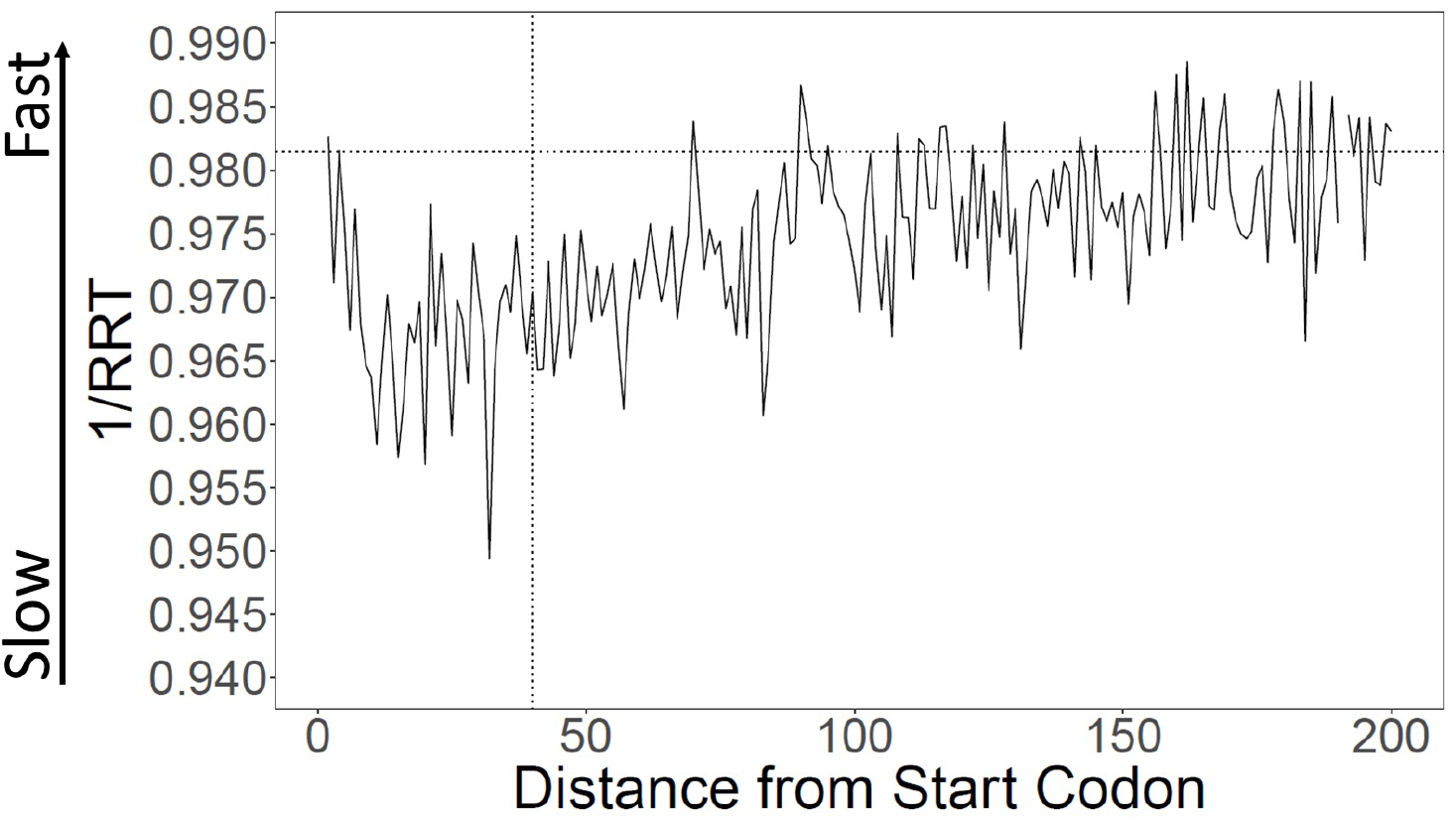
Calculation of translation speed confirms slow initial translation (SIT). Translation speeds were calculated using Ribosome Residence Time (RRT) (Gardin et al. 2014) (Table S1 for RRT values) as a measure of codon-specific translation speed over 5694 *S. cerevisiae* ORFs. The horizontal line indicates average inverse RRT across all ORFs. The average speed in the first 40 amino acids is about 1.1% slower than in the rest of the gene (p < 0.001).

We observed a qualitatively similar, but smaller, encoded slow translation at the 3’ ends of genes, of borderline statistical significance (Figure S1).

Although on average genes are translated slowly near their 5’ ends, there is variability. Figure S2 shows the distribution of initial encoded translation speed for all yeast ORFs. About 57% of genes have a Slow Initial Translation (SIT) region, while the remainder have Fast Initial Translation (FIT).

### Rare (slow) codons are enriched within the first 40 codons

Rare codons tend to be slow codons, and *vice versa*^6^ (Figure S3). But the correlation is not perfect—other things being equal, A/T-rich codons tend to be faster than G/C-rich codons, and codons with a wobble base at the third position tend to be faster than codons with the cognate base^6^. To better understand why gene beginnings are more slowly translated, we examined the relative usage of each of the 61 sense codons in the first 40 codons (Figure 2).

**Figure 2.**
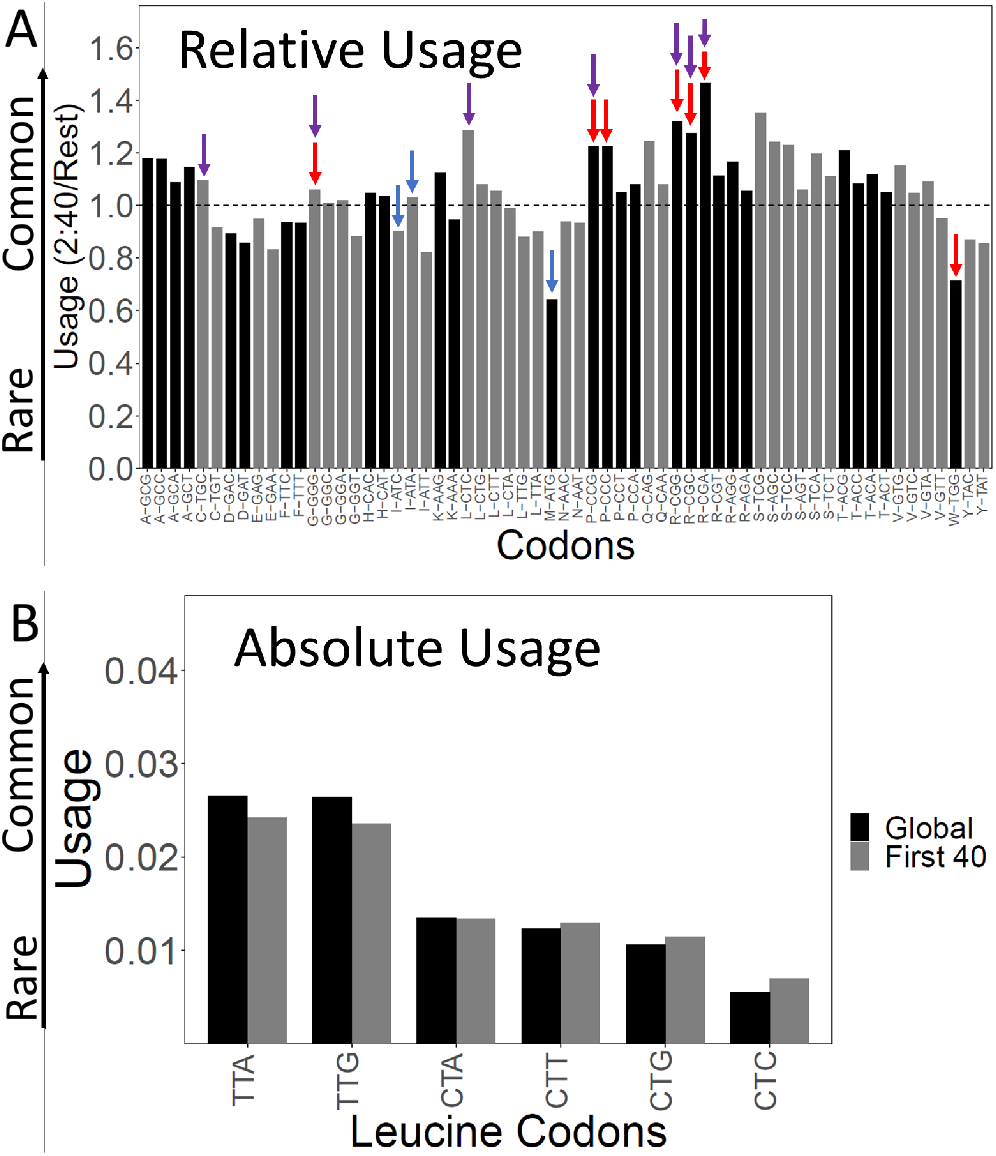
Codon usage in the Slow Initial Translation (SIT) region. A. Relative usage of each codon in the SIT versus the rest of the gene. The Y-axis shows the usage of each codon in the first 40 amino acids (omitting ATG) divided by its usage in the rest of gene. Each of the 61 sense codons is displayed along the X-axis, grouped by amino acid. Within each group, codons are ordered, left to right, from least frequent to most frequent over the whole transcriptome (i.e., the leftmost codon in each group is the rarest). Red arrows show the seven slowest codons by RRT, purple arrows show the seven rarest codons by total usage over the transcriptome, and blue shows Start and possible Start codons (ATG, ATA, ATC). Ratios above 1 indicate enrichment in the first 40 amino acids. Typically, the rarest codons for each amino acid are enriched. B. Absolute usage of each leucine codon in the SIT. The absolute usage frequency of each leucine codon is shown globally, and for the first 40 amino acids. Rare codons are still rare in the SIT, just not as rare as elsewhere. The same pattern holds for the other amino acids.

In the first 40 codons, we saw notable depletion of AUG (by nearly 50%), ATT, and ATC (Figure 2A). These are all fast codons, and their depletion would lead to slower translation. But AUG is the usual Start codon, and ATT and ATC can also be used as Start codons. Possibly these codons are depleted to reduce the possibility of translation initiating at the wrong place. We considered whether the SIT might be due to depletion of Start codons. We recalculated translation speeds after assigning AUG a neutral speed. On the one hand, this did reduce the difference in translation speed between the SIT and the rest of the gene by about 13% of the difference, but even so, the slow initial translation was highly significant. Thus, bias against Start codons contributes only modestly to slow translation.

We also saw enrichments of all six arginine codons (likely because of their use in N-terminal signal sequences, data not shown), especially the slow, rare codons CGA, CGC, and CGG. Slow, rare codons for proline (CCC, CCG), leucine (CTC), glycine (GGG) and cysteine (TGC) were also enriched (Figure 2A).

### Rare codons are rare in the first 40 codons, just not as rare as elsewhere

Although the proportion of rare, slow codons in the SITs is relatively higher than in the body of genes, in absolute terms rare codons are still rare compared to more common synonymous codons (e.g., Figure 2B, Leu codons). That is, rare, slow codons are still strongly disfavored in the SIT, though they are less strongly disfavored than elsewhere. This was true for all rare codons.

### Why are there relatively more rare, slow codons at 5’ ends?

Our analysis is consistent with that of Tuller et al. to the extent that we find a slight relative enrichment of rare, slow codons near the 5’ ends of coding regions. Tuller et al. interpret the slow translation ramp as an adaptation—they believe there is selection for slow codons near the 5’ end to enhance efficiency of translation. But there are other possibilities.

### The Young Spandrel Hypothesis

We noticed (see below) that the N-termini of yeast genes are often poorly conserved, and otherwise highly homologous genes often vary at the N-termini between different closely-related species. This suggests a different idea: N-termini are unstable and variable in evolution. They form *de novo* from new DNA sequence, and so all codons may initially occur at similar frequencies. Since these N-termini are, on average, younger than the remainder of the gene, selection has worked on them for a shorter time. Therefore selection against rare, slow codons may be less complete, and N-termini, due to their relative youth, may still retain some extra rare codons.

In this idea, in contrast to Tuller et al., the slight excess of rare codons near N-termini is not adaptive; it is not at all a product of selection. Instead, in the words of Gould and Lewontin ^4^, it is a spandrel. It is a non-adaptive by-product of something else, in this case the evolutionary instability of N-termini. This idea is explored below.

### Poor 5’ conservation is a feature of many yeast genes

To illustrate the idea, we picked example genes. We used protein-protein BLAST at NCBI to blast several query genes against species of the subphylum *Saccharomycotina* (but having subtracted out all of *Saccharomyces*). Figure 3 shows that for these chosen example genes, the middle portions of the proteins are highly conserved, but the N-termini are not. We suggest that these non-conserved N-terminal regions are young in evolution, and therefore likely contain an excess of rare codons.

**Figure 3.**
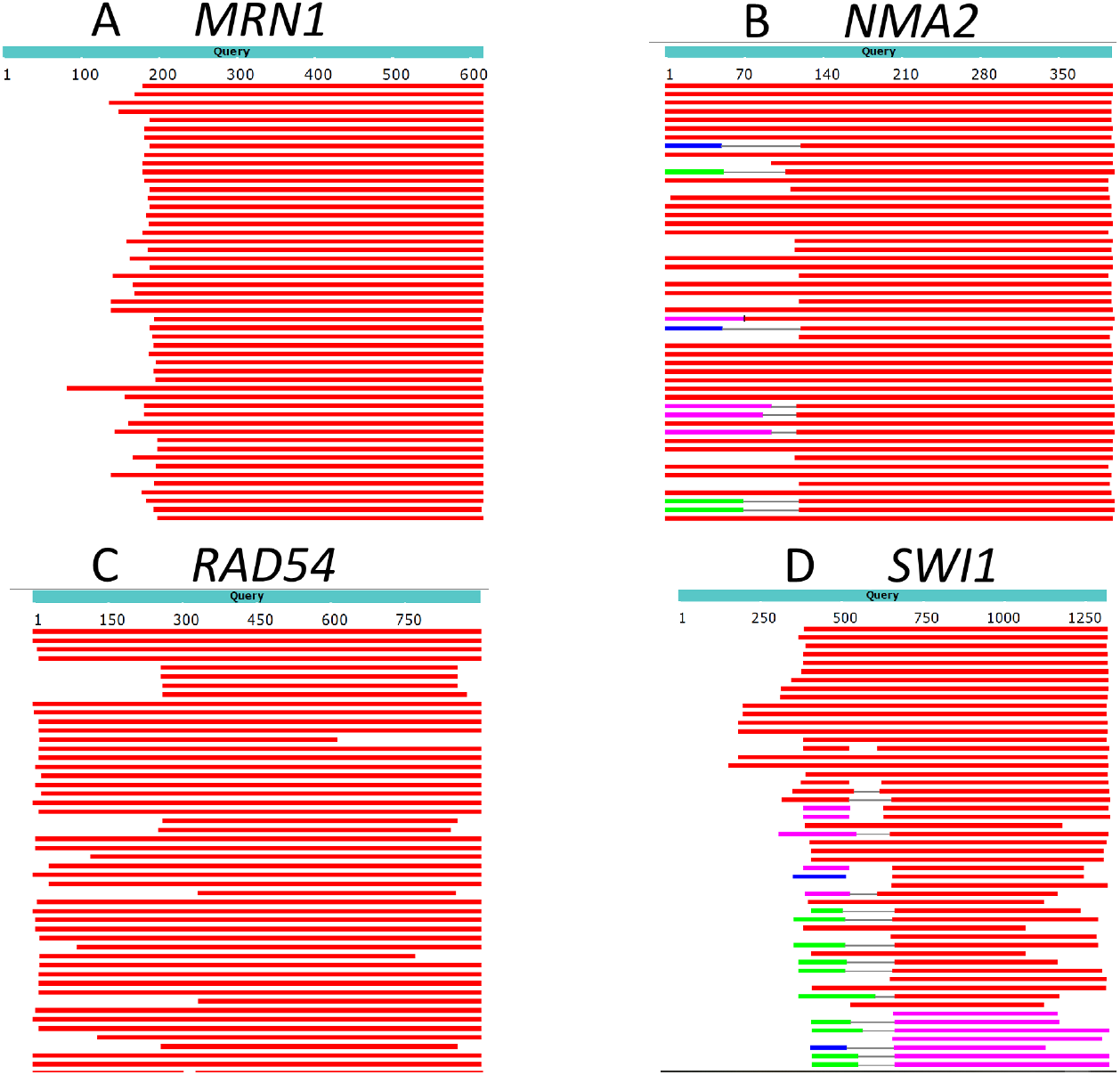
The N-termini of proteins can vary in evolution. BLAST of four example *S. cerevisiae* proteins against proteins in the subphylum “*Saccharomycotina*” (taxid: 147537) (excluding *Saccharomyces,* taxid 4930) was performed. Top hits are shown. Red regions indicate homology with an alignment score > 200, while white indicates no detected homology (BLAST default parameters). Even though all hits have high to moderate homology towards the center of the protein, many have little or no homology at the N-terminus.

In the next section, we ask whether the findings of Figure 3 can be generalized to proteins of *S. cerevisiae*.

### Method of scoring N-terminal conservation, and rationale for using Saccharomycotina

We continued our investigations into N-termini, protein conservation, and translation speed using a more systematic approach. We ran local protein-protein BLAST for all *S. cerevisiae* genes against a database consisting of the *Saccharomycotina* subphylum, omitting *Saccharomyces cerevisiae*. *Saccharomycotina* was chosen because almost every gene from *S. cerevisiae* has a recognizable, conserved homolog in almost every species in *Saccharomycotina*, and yet the evolutionary distances are long enough that there is considerable sequence variability. We excluded species of *Saccharomyces* as they are too closely related to *S. cerevisiae*, and they are also very numerous in the database, and would overwhelm results from the other members of *Saccharomycotina*. We believe, however, that this particular choice of subphylum does not greatly affect the final result.

Conservation at the N-terminus was calculated as the weighted proportion of yeast species with sequence matches (a match by default BLAST parameters) beginning in the first 40 amino acids. The lowest conservation score is 0 (no hits in the first 40 amino acids), whereas the highest conservation score is 40 indicating that every species had a match (default BLAST parameters) starting at the first amino acid (Table S2, S3). The length of genes is negatively correlated with the N-termini conservation score (rho= −.47; p<0.001)—that is, short genes tended to have more conserved N-termini.

### N-termini are variable and poorly conserved

We measured protein conservation across more than 3,000 *S. cerevisiae* proteins with orthologues among 822 closely related yeasts from *Saccharomycotina*. For each protein, we developed a conservation score for the first 40 amino acids to represent the N-termini, an equivalent conservation score for the middle 40 amino acids, and an equivalent conservation score for the C-terminal 40 amino acids (Methods and Materials; Tables S2, S3).

Strikingly, the N-termini of *S. cerevisiae* orthologs had conservation scores that were much lower, and very differently distributed than the middle of the same orthologs (Figure 4). The first 40 amino acids had a flat distribution of protein conservation scores, indicating high levels of variability amongst these orthologs. That is, many of the orthologs had no detectable homology with the first 40 amino acids of the *cerevisiae* protein.

**Figure 4.**
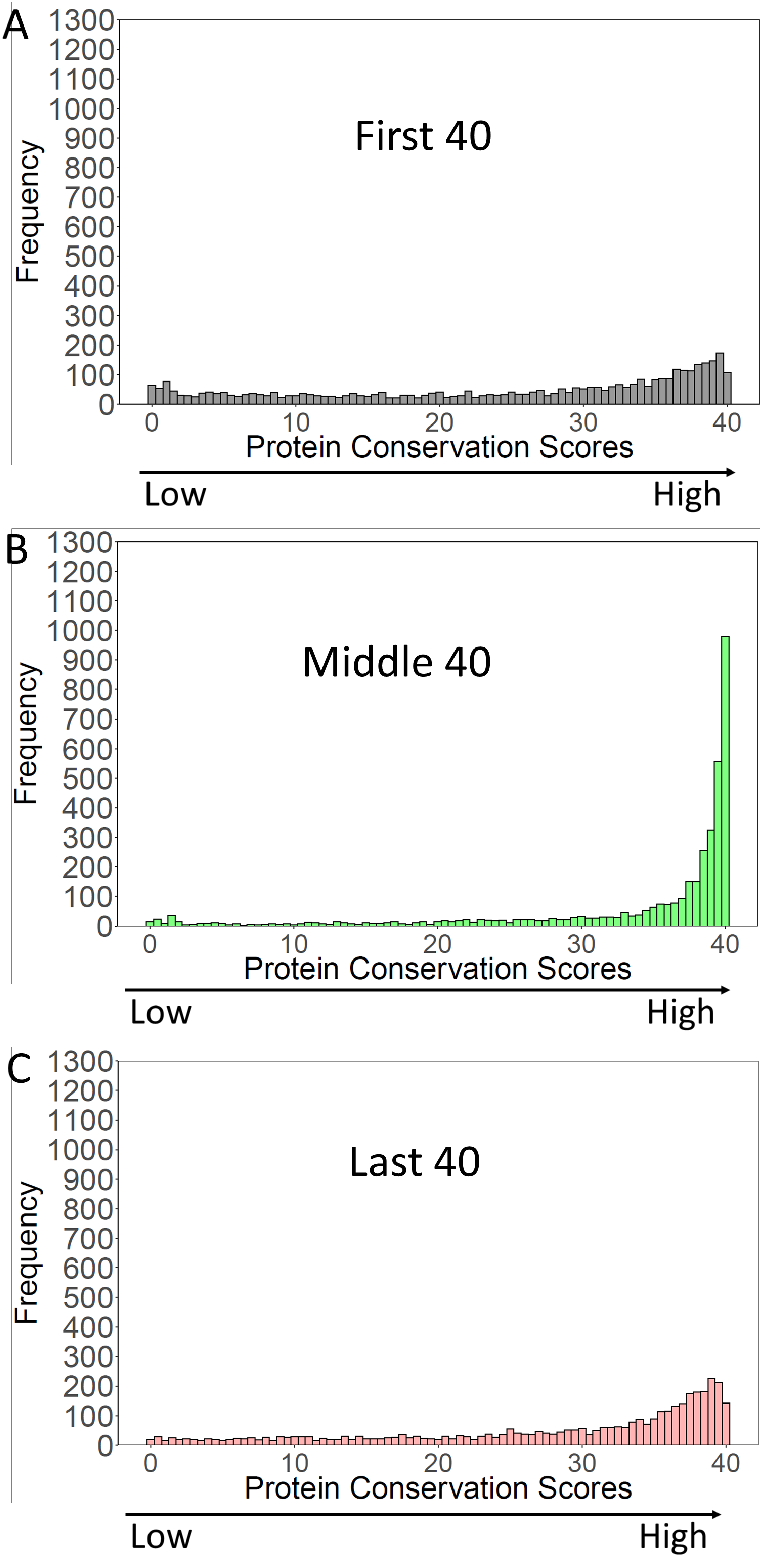
Conservation of *S. cerevisiae* proteins over the N-terminal, Middle, and C-terminal 40 amino acids. *S. cerevisiae* proteins were blasted against proteins of *Saccharomycotina* (excluding *cerevisiae*). “Conservation Scores” (Methods and Materials) were calculated for the N-terminal, Middle, and C-terminal 40 amino acids of the *S. cerevisiae* proteins. Scores range from 0 (no conservation) to 40 (perfect conservation). The frequency of each conservation score (3964 *S. cerevisiae* proteins) was plotted.

In contrast, the middle 40 amino acids were highly conserved, with conservations scores peaking at 40, the highest possible score. It Is evident that for the middle 40 amino acids, a large fraction of orthologs had a region of high homology to the *S. cerevisiae* protein, whereas this was not true for the N-termini. Finally, the last 40 amino acids had conservation scores similar to those of the first 40 amino acids, though a bit higher (more conserved) (see Figure S4 for a comparison). These results suggest both ends of the gene “breathe”, gaining and losing new sequences during evolution, whilst the middles of the same genes stay constant. Thus the ends of the genes are younger than the middles. At their first formation, they would likely have contained some rare codons, which selection may not yet have had time to remove.

### 5’ translation speeds positively correlate with 5’ conservation scores

If the “Young Spandrel” hypothesis is true, and slow 5’ translation is caused by evolutionary instability of 5’ ends, then there should be a correlation between encoded slow translation, and poor N-terminal conservation. Our model predicts the newest N-termini to have the lowest conservation and the rarest codons, and, vice versa, the termini with the rarest codons should have the lowest conservation scores. To test this, we ranked all genes by the conservation scores of their first 40 amino acids. We then divided this ranked list into thirds. For each of the thirds (i.e., the bottom, middle, and top conservation scores) we plotted the average relative initial translation speeds.

As shown in Figure 5A, the genes with the most poorly conserved N-termini also had the slowest initial translation, while the genes with the most conserved N-termini had the fasted initial translation, supporting the Spandrel hypothesis, and opposite to the Ramp hypothesis.

**Figure 5.**
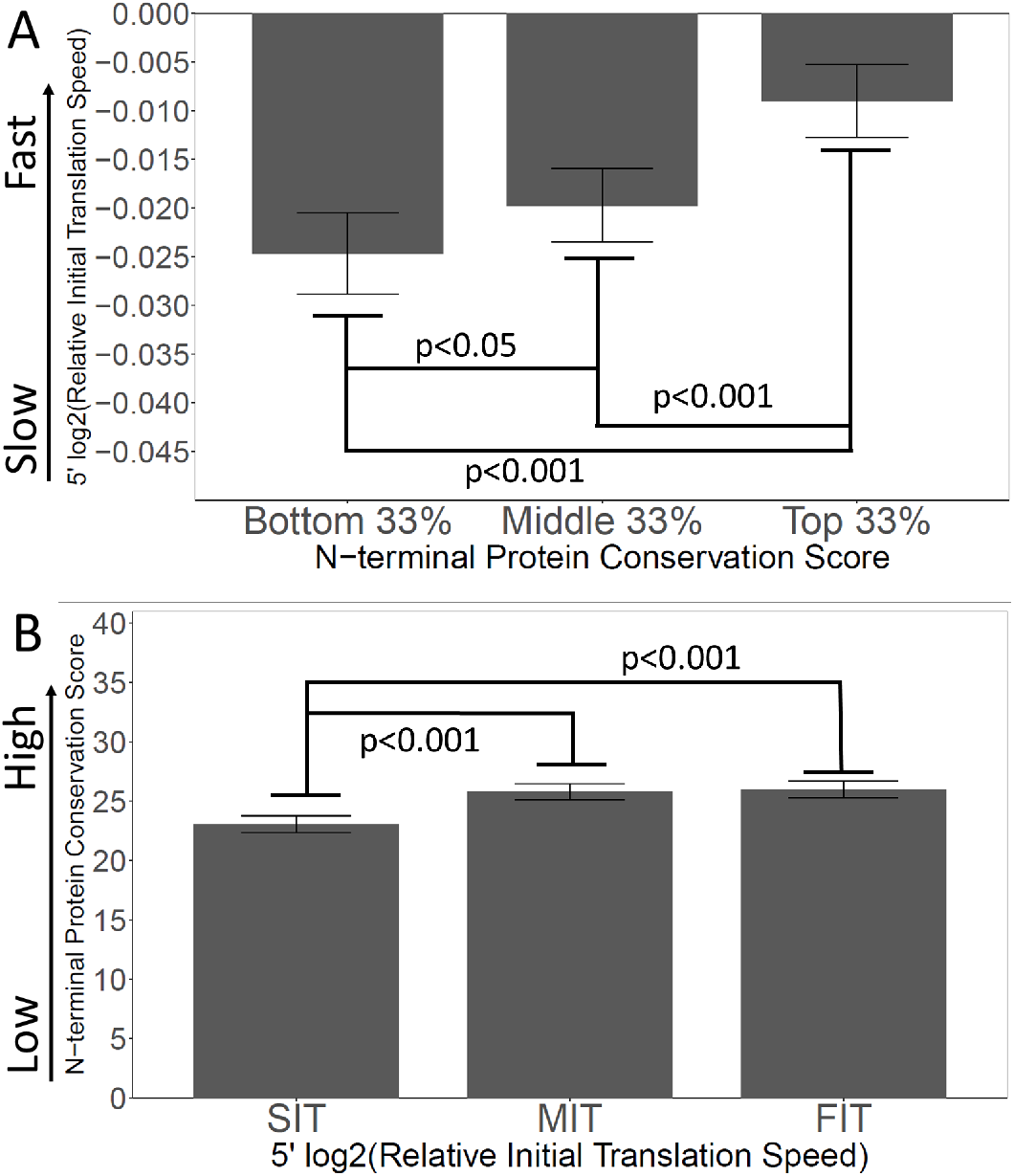
Slow Initial Translation is correlated with poor N-terminal conservation. A. Proteins were grouped by their N-terminal conservation scores (top, middle, and bottom thirds), and then the relative initial translation rate was plotted for each group. More conserved N-termini have faster initial translation. B. Proteins were grouped by their initial translation rate (top, middle, and bottom thirds), and then the N-terminal conservation scores were plotted for each group. Genes with faster initial translation have more conserved N-termini. Relative Initial Translation Speed is the log2 of (average RRT of the first 40 amino acids divided by the average RRT of the rest of the same gene) (Methods and Materials).

We also looked at the correlation in the other direction (Figure 5B). We ranked genes by their relative initial translation speed, and divided the ranked list into thirds, then plotted N-terminal conservation scores. Again the effects are correlated: genes with the slowest initial translation have the lowest N-terminal conservation scores. Thus, overall, there is a strong correlation between N-terminal instability (i.e., newness in evolution, low conservation scores) and slow initial translation (i.e., the presence of slow/rare codons).

### The Ramp hypothesis is inconsistent with observations of ribosome density and gene expression

In the Tuller “Ramp” hypothesis, in which the purpose of the slow translational ramp is to queue ribosomes and prevent collisions, genes with the highest ribosome occupancy would be in the most danger of ribosome collisions, and would therefore presumably have pronounced SITs. SITs might not be necessary on genes with low ribosome density, since there would not be much danger of collision in any case. To test this, we used the Arava et al. 2003 dataset which measured the density of ribosomes on all *S. cerevisiae* mRNAs^12^ (Figure 6). We ranked genes by ribosome density, then grouped them in thirds. Opposite to the expectation from the Tuller et al. “Ramp” theory, the genes with the highest ribosome densities had the fastest initial translation, whereas the genes with the lowest ribosome densities had the slowest initial translation (Figure 6C). While these findings are opposite to the expectation of the “Ramp” theory, they are consistent with the spandrel theory, because genes with high ribosome density would be subject to more intense selection against slow codons, thus leading to faster 5’ ends. Furthermore, analysis of Conservation Scores on the same genes showed that the genes with the lowest ribosome densities also had the lowest Conservation Scores (Figure 6D), as predicted by the Spandrel hypothesis.

**Figure 6.**
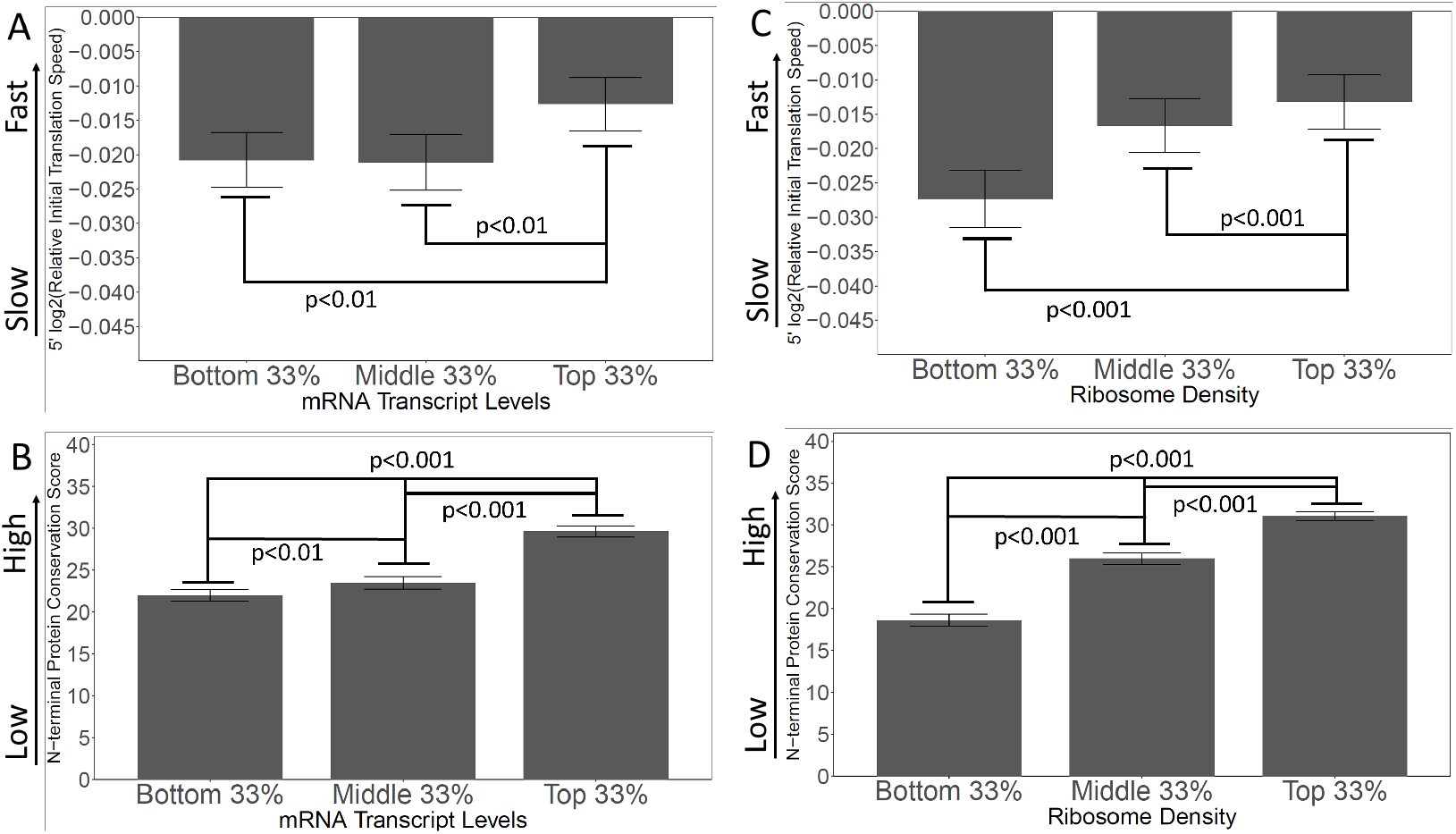
Genes with high levels of expression, and high ribosome densities, generally have rapidly-translated N-termini, and high N-terminal conservation scores. **A and B**. Genes were grouped by expression level (bottom, middle, and top)(except that genes with fewer than 10 read-counts were omitted to reduce noise) (Lipson et al. 2009). In A, the initial translation rate is shown; in B, the conservation scores are shown. The correlation between speed and transcript abundance fails for the bottom third of genes; possibly these are genes expressed at high levels under other conditions (e.g., meiosis and sporulation). **C and D**. Genes were grouped by ribosome density (Arava et al. 2003) as a measure of intensity of translation. In C, the initial translation rate is shown; in D, the conservation scores are shown. High ribosome density correlates with high initial translation speed and high conservation score.

Similarly, in the “Ramp” hypothesis, ribosome collisions on highly-expressed genes would presumably have more serious consequences for the cell than collisions on poorly-expressed genes, and so highly expressed genes ought to have the most pronounced SITs. To test this, we used the Lipson et al. 2009 transcriptome dataset which measured the number of mRNA transcripts for *S. cerevisiae* genes^13^. Again, we ranked genes by expression, then grouped them by thirds. Exactly contrary to the “Ramp” hypothesis, we found that genes with the highest expression had the fastest initial translation (Figure 6A). In this analysis, the middle and bottom genes are not significantly different from each other; possibly some of the poorly expressed genes are inducible genes that would be highly expressed under some other condition (e.g., the sporulation genes, the *GAL* genes). In addition, the most highly expressed genes had the highest conservation scores, consistent with the Spandrel hypothesis (Figure 6B).

### Experimentally, encoded slow initial translation does not increase gene expression; the opposite is true

Tuller et al. hypothesized that slow initial translation was adaptive, and improved efficiency of translation and gene expression by minimizing ribosome collisions. However, in the Spandrel hypothesis, slow initial translation is not generated by selection and is not adaptive. It might not have any effect on gene expression, but, if anything, slow translation might reduce gene expression. While informatics is wonderful, it is always nice to do an experiment, and in this section, we present direct experimental results regarding the effect of encoded slow, medium, and fast initial translation on gene expression.

We used a gene expression reporter based on EKD1024^14^ (Methods and Materials). In this construct, GFP is the reporter, but for accuracy it is normalized against a divergently transcribed red fluorescent protein (Figure S6). Thus, GFP expression is reported as a GFP/RFP ratio. Although the reporter is GFP, the N-terminal region of this particular protein is derived from yeast *HIS3*, not GFP, and likely has little if any effect on the fluorescence of the GFP fused downstream^15–17^. We used synonymous slow, medium, or fast codons to recode some of the codons in the first 41 amino acids of this GFP reporter to generate three reporters with slow, medium, or fast translation over the first 41 amino acids. We emphasize that the amino acid sequences of the three constructs were identical. The slow, medium, and fast average RRT values over the first 41 codons were 1.20, 1.04, and 0.93, respectively. That is, this SIT is slower than most natural SITs, and this FIT is faster than most natural FITs, but the difference is moderate.

As shown in Figure 7 (left three bars), the SIT did not improve expression of GFP, contrary to Tuller et al. In fact, the GFP with the SIT was expressed at only 71% of the level of the GFP with the FIT. It was surprising to us that the difference was this large—again, recoding was limited to codons within the first 41, and the protein sequences were identical.

**Figure 7.**
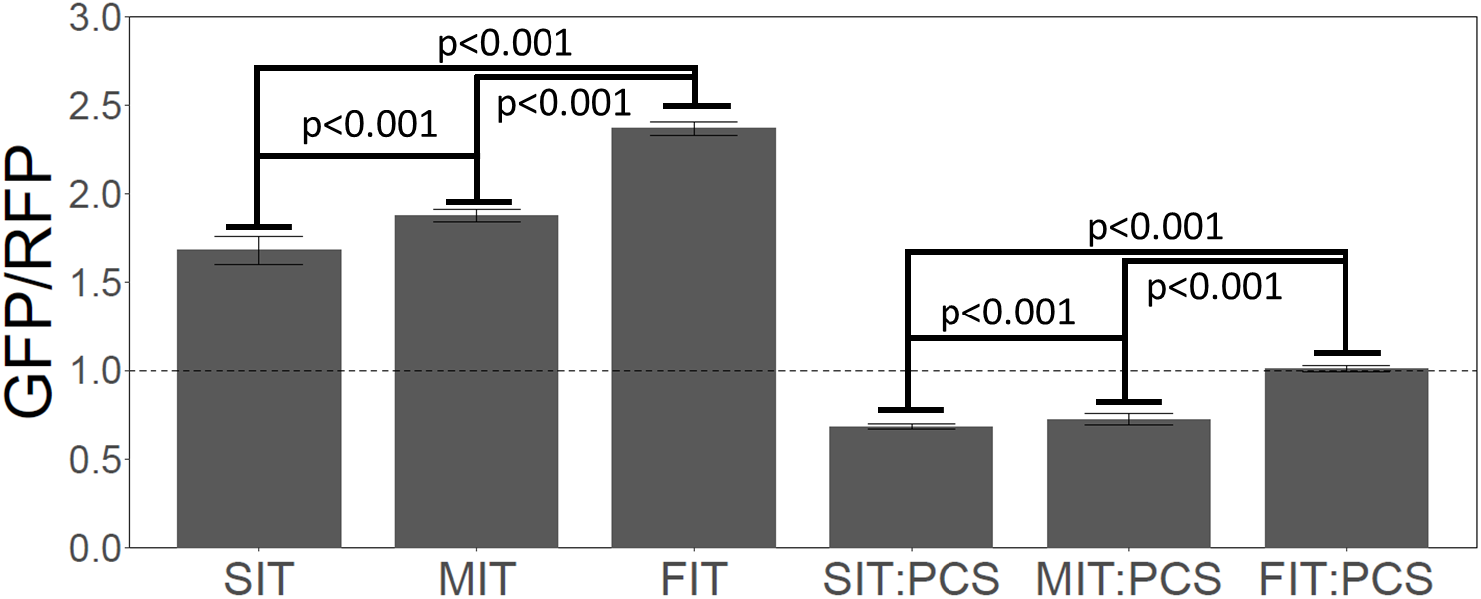
Slow Initial Translation inhibits gene expression. Left 3 bars. A synthetic GFP was constructed with a leader amino acid sequence that had little effect on GFP. The leader sequence was recoded to give slow (SIT), medium (MIT) or fast (FIT) translation speed over the first 41 amino acids, without changing the amino acid sequence—i.e., the SIT, MIT and FIT had identical amino acid sequences, but different average RRTs. Each construct (SIT, MIT, FIT) was integrated in single copy at the *ADE2* locus, and 25 independently-transformed strains were picked, and GFP fluorescence was measured for each, and the RFP-normalized mean was plotted. Numerical values were: SIT, 1.66; MIT, 1.80, FIT, 2.29. GFP was normalized to RFP expressed from the same reporter molecule, but RFP fluorescence hardly changed amongst the transformants, and non-normalized GFP would have given very similar results. Slower initial translation reduced gene expression. Right 3 bars. As above, but a Putative ribosome Collision Site (PCS) (CGA-CGG) was inserted between the leader and the GFP. Again, slower initial translation reduced gene expression. Values were: SIT:PCS, 0.69, MIT:PCS, 0.74, FIT:PCS, 0.99.

Another possibility is that a SIT can protect against ribosome collisions when there is a site downstream that induces ribosome collisions. Sites thought to induce ribosome collisions include rare Arg-Arg codon pairs^5,18^. We therefore introduced the codon pair CGA-CGG (replacing Asn-Asp, AAT-GAT) downstream of the first 41 amino acids, but still upstream of important GFP residues. Indeed, this single CGA-CGG codon pair, potentially inciting ribosome collisions, caused a large reduction--about 50%--in expression of GFP (Figure 7, right three bars). The reduction was about same in the SIT, MIT, and FIT constructs. In this case, with putative collision sites, the GFP with the SIT was expressed at only 67% of the level of the equivalent GFP with the FIT. That is, this SIT (a fairly extreme SIT) did not at all protect against the putative ribosome collisions—if anything, it made things worse. This result suggests there is no benefit to “queuing” ribosomes, if “queuing” even occurs. Instead, the fastest-translating gene once again gave the highest expression, and the highest relative expression, despite the collision site.

## Discussion

Tuller et al. found that the 5’ ends of genes are translated slowly because of the codons used at 5’ ends, and posited that this was a selective advantage because it somehow increased efficiency of translation. However, this theory predicts positive correlations between slow initial translation and high gene expression, and slow initial translation and high overall (that is, on the whole gene) ribosome density. In fact, by informatic analysis of existing data, we find the correlations are opposite to those predicted by the “Ramp” model. Most importantly, an experiment in which rates of initial translation were changed shows that faster initial translation causes higher gene expression, exactly opposite to the prediction of the “Ramp” hypothesis. We believe no ramp is needed.

Tuller et al. believed that a region of slow translation is encoded, using slowly-translated codons, and it is specifically this idea of encoded slow translation that we are addressing. However, in addition to “encoded” slow translation, there is some evidence for slow translation at 5’ ends by other, unknown mechanisms. Ribosome profiling experiments show an increased density of ribosome footprints near the 5’ end, independent of encoding^3^, and a high density of footprints could be due to slow translation. It is now known that much of the reason for increased density of footprints in early studies was the use of cycloheximide as the first step in the original ribosome footprinting protocols. Addition of cycloheximide to growing cultures allowed ribosomes to initiate at Start codons, but did not allow elongation, hence there was a pile-up of ribosomes near the 5’ end. More recently, flash-freezing, rather than cycloheximide, has been used as the first step in stopping translation, and even in these studies, there is about a 50% increase in ribosome density near 5’ ends^3^. And yet, even in these flash-freezing protocols, cycloheximide is used in a later step, when extracts are thawed, to prevent elongation, and this cycloheximide usage at a second step could again result in an artefactual increase in ribosome density at 5’ ends. Alternatively, the increased density of ribosomes at 5’ ends could mean that some proportion of ribosomes fall off the mRNA as they progress^3^. Consistent with the latter idea, ribosome profiling studies show a general trend towards lower ribosome densities at more 3’ positions in translating mRNAs (Weinberg et al., their Figure S7^3^), and studies using other experimental approaches have shown a general decrease in ribosome number or density as one progresses along a gene ^19,20^. For these reasons, it is not clear that there is any slow translation in the 5’ regions of genes.

We considered that an increased density of ribosomes at the 5’ end could be because some genes have additional ATG Start codons, sometimes upstream and sometimes downstream of the annotated Start, and translation of short open reading frames from these additional Start codons could contribute to ribosome density at the 5’ end. Using the program “Frameshift Detector”^21^ and ribosome profiling data, we quantitated the fraction of out-of-frame ribosomes both globally, and within the first 150 nucleotides of genes. We found the global proportion of out-of-frame ribosomes is about 13%, and the proportion of out-of-frame ribosomes in the first 150 nucleotides is about 14.5%. Although this increased out-of-frame ribosome presence of about 1.5% is highly significant (p ∼ 10^-28^), it is not nearly big enough to explain the observed 5’ increase (∼ 50%) in ribosome density (ref. 3, their Fig. 1C).

In any case, by direct experiment, we find that encoding a faster 5’ end using fast synonymous codons improves gene expression. In particular, even when a ribosome collision site was placed downstream, the effect of the collision site was not at all ameliorated by encoded slow translation upstream of the collision site. This seems strong evidence against the idea that slow initial translation is a defense against collisions. The basis of the idea that slow initial translation could possibly be a defense against collisions is not clear to us. Regions of slow translation would not affect the gaps between ribosomes, if measured as times, and especially not if measured at a constant finish-line, such as a putative collision site.

A potential weakness in our GFP reporter experiment is that the mRNA sequences are necessarily different, and so there are different mRNA structures. We achieved fast and slow sequences by recoding with synonymous codons, so amino acids are identical, and there are no issues of, e.g., co-translational protein folding, or interaction with the ribosome exit tunnel. But of course the RNA structures will be at least slightly different, and RNA structures at the 5’ end are known to affect translation initiation^3,22–27^. Generally, more open mRNA structures are more favorable for translation initiation, and also for translation speed. To fully resolve this would likely require the recoded GFP reporter experiment to be redone many times, starting from different underlying sequences, and even then, it might be hard to disentangle effects of open mRNA structure on initiation from effects of fast translation. But whether the increased gene expression we see for the “fast” encoding is being generated mainly by fast translation, or by efficient initiation, in neither case is there an argument that slow translation is efficient, or that it protects against collisions.

The observation that 5’ ends have low conservation, likely because of instability in evolution, provides a completely different explanation for the region of encoded slow initial translation. In this “Spandrel” hypothesis, N-termini frequently change in evolution, gathering new 5’ sequences *de novo.* These would contain all codons at similar frequencies—i.e., rare codons would not be especially rare. Although these rare codons would eventually be removed by selection, the fact that N-termini are relatively young means that this process might not be complete for all genes, and so some rare, slow codons still remain. These explain the initial region of encoded slow translation. This hypothesis is highly consistent with the observed correlations between slow initial translation and low gene expression; and slow initial translation and low ribosome density, and with the results of gene expression experiments. It is also consistent with the region of slightly slow translation we observe at the 3’ ends of genes.

We have looked at the conservation of N-termini only in *S. cerevisiae*. However, Tuller et al. found that there is a region of encoded slow initial translation in genes of a wide variety of eukaryotes. We speculate that in these other cases, too, slow initial translation is a spandrel deriving from the turnover of 5’ ends. This in turn has implications for protein structure and evolution; for the interpretation of evolutionary sequence clocks; and for the rates of selection against rare codons. We note that Bricout et al.^28^ have also recently found that N- and C-termini of proteins evolve faster than the middles, although their perspective is different from ours.

It was surprising to us that recoding just the first 41 codons of the GFP fusion protein from slow to fast increased the level of GFP expression by so much—about 30%. These 41 codons were originally derived from the yeast *HIS3* gene, and this increase in expression is roughly the proportional increase expected based on fully recoding the *HIS3* gene to preferred codons^29^. Because this recoding from slow to fast tends to replace G/C rich codons with A/T rich codons, recoding from slow to fast may decrease the stability of RNA structures near the 5’ end, and increase the accessibility of the cap. Decreased stability of mRNA structures could be responsible for the increase in gene expression, consistent with studies in both yeast^3,23^ and *E. coli*^26^.

We characterized both the 5’ and 3’ ends of genes. Qualitatively, results were similar: both 5’ and 3’ ends showed reduced conservation scores, increased proportions of rare codons, and slow translation, consistent with the Spandrel hypothesis. However, quantitatively, effects on rare codons and slow translation are much bigger at the 5’ end. We are puzzled by the asymmetry. Contributing to the asymmetry is the depletion of initiation codons from the 5’ end. These are fast codons; they are presumably depleted to prevent translation-initiation at the wrong place; and this effect is specific to the 5’ end. Also contributing are basic, arginine-rich sequences in N-terminal signal sequences; these Arg-rich sequences include many rare Arg codons, and this enrichment for N-terminal signal sequences is specific to the 5’ end. Still, even after taking these into account, quantitative effects seem bigger at the 5’ end. We speculate that although rare codons and slow translation are generated and maintained for a time as spandrels, poor translation could be a selective advantage in genes that need to be expressed at low levels. Here, we are turning the hypothesis of Tuller et al. on its head: contrary to Tuller, slow initial translation seems to reduce gene expression, and yet this may be adaptive for some genes. Similarly, although architectural spandrels were originally generated from the need to support a lintel by an arch, nevertheless, there may eventually have been “selection” for them, as church-goers found them beautiful.

We believe our study has three main limitations or uncertainties. First, in our GFP reporter experiments suggesting that a slowly-translated N-terminal region reduces gene expression, we did not address the possibility that changes in mRNA structure could have been responsible for this. We believe that experiments with a multitude of reporter sequences would be required to address this issue. Second, we found somewhat different results at the 5’ and 3’ ends of genes: both ends show poor evolutionary conservation, but the enrichment of rare codons is more pronounced at the 5’ end. Although we suggest this could be due to selection for poor translation at the 5’ end (see above), this is only a suggestion, and we feel the reasons for the difference between the 5’ and 3’ ends are unresolved. Third, ribosome profiling studies show high ribosome densities near the 5’ end of the mRNA which are not nearly fully accounted for by rare codons^3^ (and see Discussion above); although we discuss (above) several possibilities for these high 5’ ribosome densities, ultimately this issue also remains unresolved.

Finally, again, in “The Spandrels of San Marco …”, Gould and Lewontin^4^ warned that not all biological phenomena are adaptive, and it is a mistake to assume that any particular characteristic of an organism must necessarily have been generated by natural selection. We believe the encoded slow initial translation of eukaryotic genes may be an example of this.

## Methods and Materials

### Statistical calculations of relative initial rate of translation

Bioinformatics were performed on protein-coding open reading frames (ORFs) of *Saccharomyces cerevisiae* downloaded from the Saccharomyces Genome Database (SGD) website, as last modified on April 22, 2021. Protein-coding ORFs annotated as dubious or pseudogenes were not included in analyses. All statistics were performed using The R Project for Statistical Computing. Translation speed was measured using the ribosome residence time (RRT) which is a metric of the occupancy of ribosomes on each sense codon within the A-site ^6^. The RRT values we used are shown in Table S1; these are modified from the original RRT results of Gardin et al. by inclusion of the ribosome profiling data of Jan et al.^30^.

### Relative initial translation speed

Tuller et al.^1^ focused on a “ramp” of translation speed, where the first part of the gene has slow translation relative to the rest of the gene. The ramp thus refers to a rate. To quantitate this slow relative ramp for each gene, we calculated the average RRT for an initial window of the gene (e.g., 40 amino acids, see below), then divided by the average RRT of the rest of the gene. Thus, genes with a “slow ramp” have a ratio less than 1. We then took log2 of this ratio; genes with a slow ramp yield a negative number, and the more negative the number, the steeper the ramp.

For each gene, the relative initial translation speed (RIT) (explained above) was calculated across windows of the first 30, 40, 50 … and 100 codons, with all windows being statistically significant for slow translation. For these RIT calculations, the first (start) codon was omitted since all protein-coding genes in this dataset except Q0075 start with ATG, and ATG is one of the fastest codons, which would skew the RIT. Similarly, the last (stop) codon was omitted.

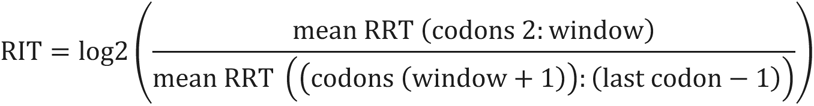

Windows of the first 30, 40, 50 … and 100 codons each had statistically significant depletion in translation speed compared to the body of genes. We chose to focus on the first 40 codons. All genes shorter than 303 nucleotides (translated into 100 amino acids) were omitted from all analyses. For all RIT analyses, 328 out of 6022 ORFs were omitted leaving a dataset of 5694 ORFs.

### Datasets

mRNA transcript readings, which we used as a proxy for gene expression, was acquired from Lipson et al. 2009^13^. Genes with a read count less than 10 were omitted due to concerns about noise. Ribosome density measurements were acquired from Arava et al. 2003^12^. These values were calculated as the number of ribosomes, detected on an mRNA, divided by the nucleotide length of the gene (including the stop codon).

### Protein BLAST setup and diagnostics

*S. cerevisiae* proteins were downloaded from the SGD (last modified on April 22, 2021). Proteins derived from ORFs annotated as dubious or pseudogene were omitted from analyses. The *Saccharomycotina* (Taxonomy ID: 147537) protein sequences were downloaded from NCBI using the links:

https://www.ncbi.nlm.nih.gov/Taxonomy/Browser/wwwtax.cgi?mode=Info&id=147537&lvl=3&lin=f&keep=1&srchmode=1&unlock

https://www.uniprot.org/uniprotkb?dir=ascend&query=(taxonomy_id:147537)&sort=organism_name

We downloaded and compiled the source databases from DDBJ, EMBL, Genbank, RefSeq, PIR, and UniProtKB. Duplicate sequences were deleted. To perform local BLAST, we downloaded the NCBI BLAST software (version 2.13.0+) and used RStudio as a wrapper to operate the software; all default BLAST parameters were selected, except that the number of alignments was changed to the maximum value of 1000000000. Local protein BLAST of every *S. cerevisiae* protein (5694 proteins, see above) was performed against all genomes of the subphylum *Saccharomycotina*, but omitting all species in the genus *Saccharomyces* (net, 822 genomes). We eliminated submissions of duplicate species by limiting our database to the highest bitscores from sequence hits derived from each unique species. We were only interested in sequences that had high homology with queried *S. cerevisiae* proteins, so all hits with bitscores lower than 50 were omitted.

We wanted to compare conservation at the beginning of proteins with conservation at the middle and end of those same proteins. For this purpose, we split each *S. cerevisiae* protein into two halves (start to middle; middle to end), then blasted each half against all genomes in the subphylum *Saccharomycotina* (omitting *Saccharomyces*). We then calculated a “conservation score” (see below) for the first 40 amino acids of the protein, and, identically, for the first 40 amino acids of the second half of the protein. (We describe the first 40 amino acids of the second half of the protein as the “middle”, but in fact the region is displaced 20 amino acids C-terminal from the exact middle.) In a parallel way, a conservation score is calculated for the last 40 amino acids of each protein. For example, the length of Swi5 is 709 amino acids, and therefore BLASTs of the first half spanned from 1:354, and BLASTs for the second half spanned from 355:709. Conservation scores were calculated for residues 1:40 (beginning) (from the BLASTs of the first half of the protein), and 355:394 (middle) and 670:709 (end) (from BLASTs of the second half of the protein). In total, BLAST of the first half of all queried *S. cerevisiae* proteins yielded a total of 477,749 high homology (minimum of bitscore of 50) sequence matches across 816 unique *Saccharomycotina* species, whereas BLAST of the second half of each protein yielded a total of 487,022 high homology matches across 812 unique *Saccharomycotina* species.

With respect to the above procedure, we note that we are relying on the BLAST algorithm to find regions of homology. Homology would be somewhat more easily found in the middle of sequences than at the ends because of seeding issues. It is for this reason that we divided proteins in half, and used a BLAST with the second half of the protein to find homologies with the first 40 amino acids of the second half. That is, in this procedure, for the middle homologies, the algorithm is being asked to find homologies at the end of a sequence, exactly as is the case for the first 40 and last 40 amino acids.

### Calculations of protein conservation scores and ratios

The general idea of the “Conservation Score” is that it represents the lengths of the regions of BLAST homology between the *S. cerevisiae* query and the *Saccharomycotina* subjects in the beginning, middle, and end 40-aminoacid windows. Each *S. cerevisiae* query sequence was separated into two equal halves, and then BLASTs were done on both halves against all of *Saccharomycotina* (omitting all submissions from the *Saccharomyces* genus). For all subject proteins with high homology over any region (i.e., a bitscore greater than 50), one finds the pair-wise regions of homology with a BLAST “Alignment Score” of 200 or more (red colored regions in the BLAST website “Graphic Summary”). The length of the high-homology region in the window of interest (but not the actual number of amino acid sequence matches within that region) contributes to the Conservation Score. That is, amino acids have to be within a region of homology found by BLAST in order to contribute to the score; even though all proteins begin with “M”, these only contribute to the conservation score if they are within a region of BLAST homology. The Conservation Score is the sum of (the length of the homology in the window, multiplied by the proportion of qualified subject proteins with that length of homology). Example Conservation Scores are shown in Table S2, and an example Conservation Score is calculated in Table S3. We only considered conservation scores from proteins that had homology with at least 40 unique species in *Saccharomycotina*. We also omitted proteins that were shorter than 100 amino acids. As shown in Table S2, conservation scores ranged from 0, meaning no BLAST homology region within the window of 40 amino acids for any qualifying homologue in *Saccharomycotina*, up to a maximum of 40, meaning that all qualifying homologues in *Saccharomycotina* had matches starting at the first amino acid. In total, protein conservation analyses used 3964 *S. cerevisiae* proteins with high homology hits for BLAST done on the first and second half of proteins (Figure 4).

#### Code

We provide the custom R code written for this project as a text file, Appendix 1.

### Design of the fluorescent reporter gene constructs

We created a reporter gene based on reporter plasmid EKD1024^14^. Briefly, a bidirectional galactose promoter simultaneously induces the expression of GFP and RFP in the presence of galactose, and we integrated the reporter into the yeast genome at the *ADE2* locus. We recoded GFP to give the first 41 codons of GFP a slow initial translation speed (SIT), medium initial translation speed (MIT), or fast initial translation speed (FIT), while maintaining the same amino acid sequence (Figure S6, sequences in Table S4). We also designed three more constructs (Figure S6, sequences in Table S4) with a SIT, MIT, or FIT upstream of one of the slowest and rarest codon pairs, CGA-CGG (replacing AAT-GAT, Asn-Asp), which is known to greatly attenuate gene expression in living yeast^16^. Other rare codon pairs (CGA-CGA and CGA-CCG) have been shown to promote ribosome stalling^18^ so in Figure 6, right, CGA-CGG operates as a putative ribosome collision site (PCS). Instead of a PCS, the constructs in Figure 6, left, had AAT-GAT (Asn Asp), a frequent codon pair with above average translation speed. The Relative Initial Translation Speed scores of the constructs were: SIT was 0.208; MIT was −.0004; FIT was −.166; SIT+PCS was 0.198; MIT+PCS was −.0109; and FIT+PCS was −0.177. (Note that the RIT scores of the constructs with the PCS change because the PCS makes the translation speed of the body of the gene slower; that is, the change is due to a change in the denominator.)

### Yeast strains

The constructs were transformed into BY4741 (*MATa*; *his3Δ1*; *leu2Δ0*; *met15Δ0*; *ura3Δ0*). The reporter expresses *MET15* allowing selection. Transformants were selected for Met+ on HULA plates (0.075g/L Histidine; 0.075g/L Uracil; 0.25g/L Leucine; 0.075g/L Adenine; 20g/L D-Glucose; 5g/L Ammonium Sulfate; 1.7g/L Yeast Nitrogen Base). The reporter gene integrates into the *ADE2* locus, and thus successful transformants are *ade2*-delete which becomes red when grown on YPD plates; this was used as a secondary biological marker to confirm successful transformants. From each strain, a total of 25 Met+, Ade-, red transformants were chosen for each recoded GFP construct.

### Flow Cytometry analysis

The strains were inoculated in liquid HULA media, with 2% galactose, until mid-log phase (around 10-14 hours) containing around 3 x 10^7 cells/mL. The strains were sonicated to separate cells and the strains were stored on ice (typically around an hour) until data collection. An LSR Fortessa Flow Cytometer was used to measure GFP levels (All Events FITC-A Mean) and RFP levels (All Events PE-Texas Red-A Mean) across 75,000 events. As a control, a strain lacking a fluorescent reporter was used. For all samples, GFP levels were normalized by RFP levels to reduce background fluorescence.

### Statistical tests

None of our analyses made assumptions regarding the normality of the data. As such, we only performed nonparametric statistics. Wilcoxon signed-rank tests were done for relevant pairwise analyses; when necessary, p-values were corrected for multiple comparisons using the Holm–Bonferroni method. Spearman correlations were used. We used the Kolmogorov–Smirnov goodness of fit test to confirm that the three distributions were significantly different (p<0.001) for Figure 4.

## Supporting information

Supplemental Table 1 RRT Values

Appendix 1: R code for analysis

## Acknowledgements

We thank Steve Ketchum and Sangeet Honey for discussions that helped form the central idea of this manuscript.

---

**Figure S1.**
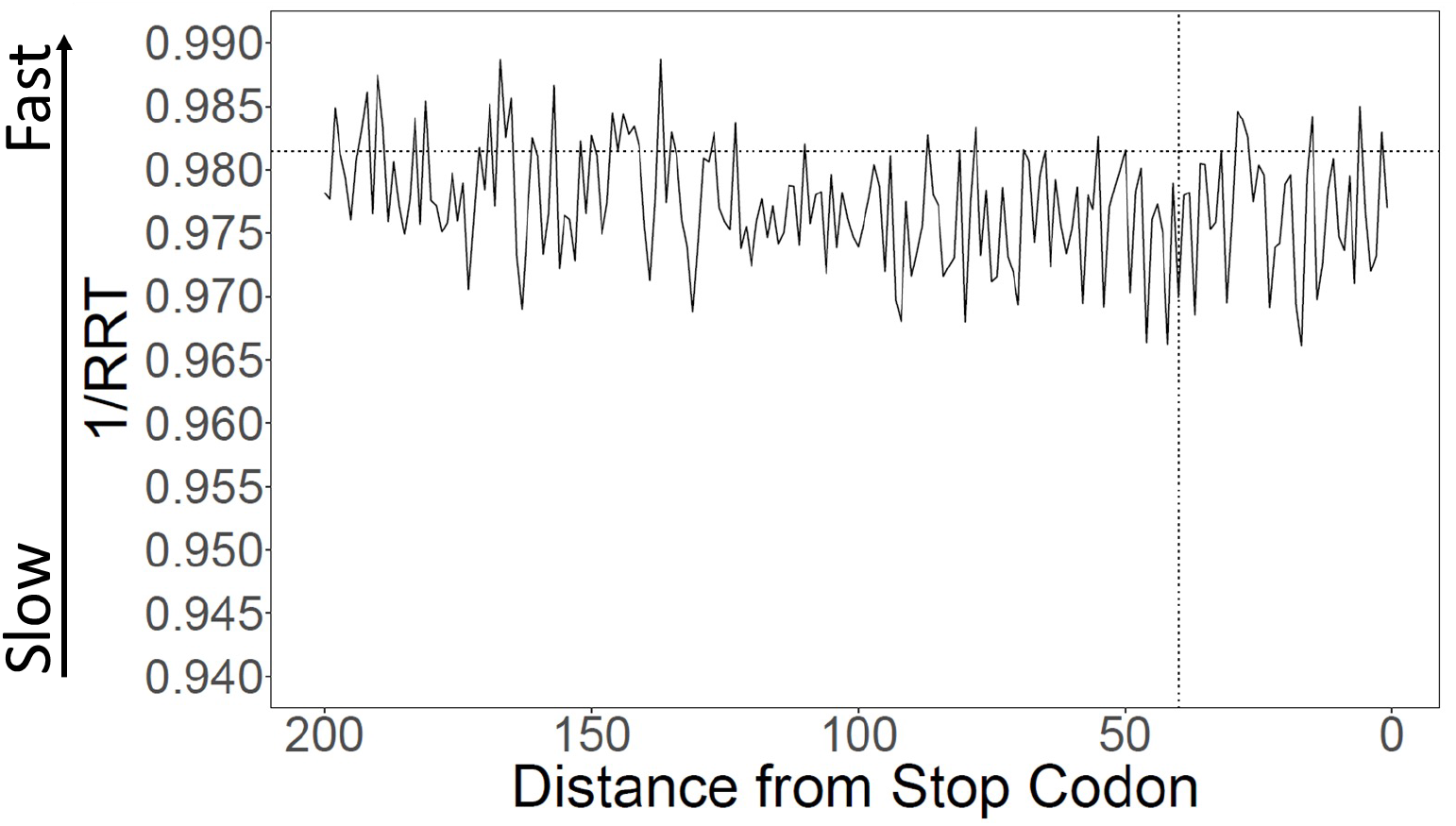
Translation speed at 3’ ends. Translation speeds at the 3’ ends of genes were calculated using Ribosome Residence Time (RRT) (Gardin et al. 2014) (Table S1 for RRT values). The average speed over the last 40 amino acids is about 0.1% slower than in the rest of the gene, not statistically significant. The average speed over the last 100 amino acids is about 0.19% slower, which is significantly different (p = 0.028).

**Figure S2.**
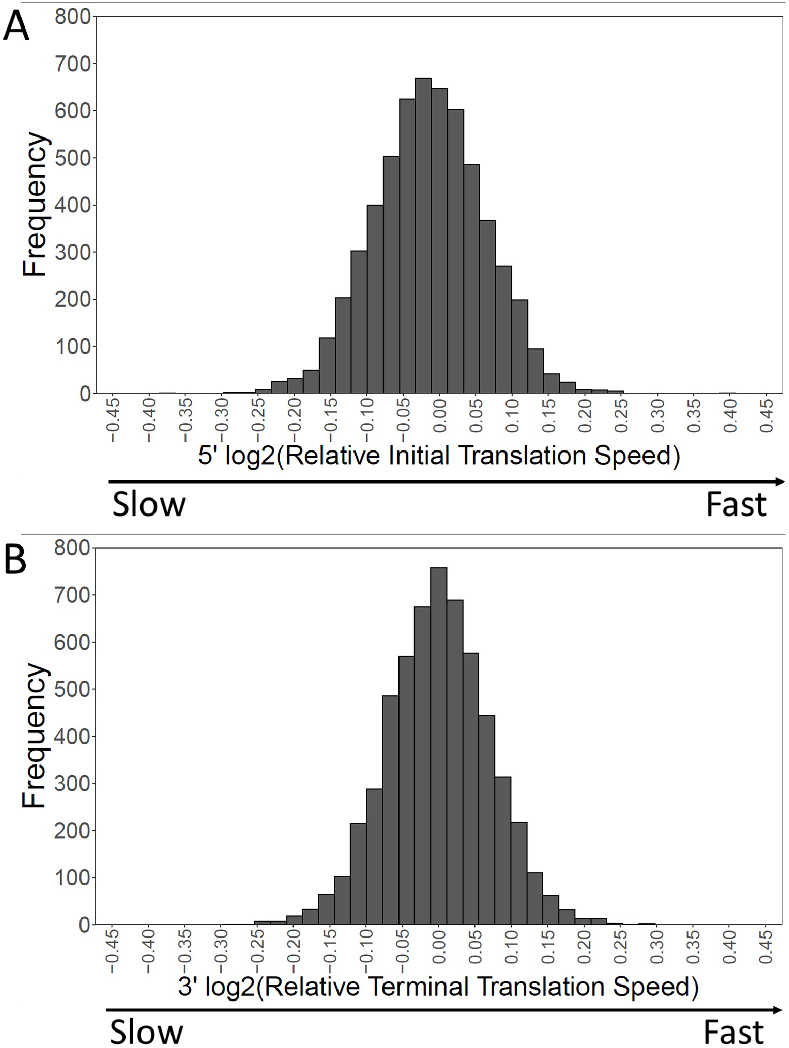
Distribution of translation speeds at 5’ and 3’ ends. The distribution of relative translation speeds over 5694 genes is shown for the first and last 40 amino acids. For the 5’ end, 57.2% of genes have relatively slow initial translation, while for the 3’ end, 50.14% of genes have slow terminal translation.

**Figure S3.**
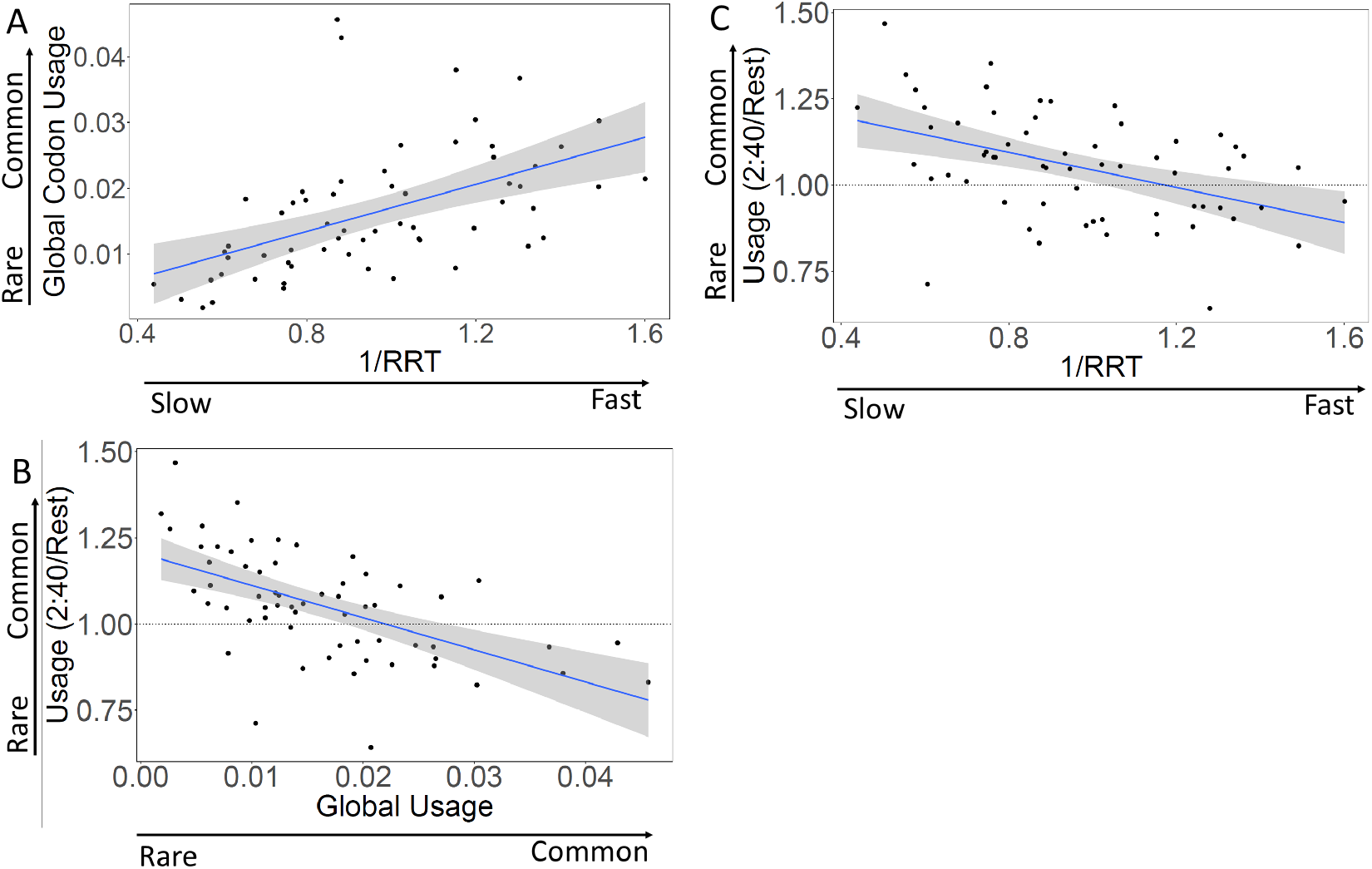
Codon speed and codon usage are correlated. **A**. Rare codons are translated slowly. Each dot represents a sense codon. The x-axis displays the translation speed of each codon (modified from Gardin et al. 2014); the y-axis displays the global frequency of usage of each codon. The correlation is 0.64, p < 0.001. **B.** The first 40 codons are enriched for rare codons. Each dot represents a sense codon. The relative usage of each type of codon in the first 40 codons of genes (i.e., in the SIT) (y-axis) is displayed against global codon usage (x-axis). The correlation is −0.61, p < 0.001. **C.** The first 40 codons are enriched for slow codons. Each dot represents a sense codon. The relative usage of each type of codon in the first 40 codons of genes (i.e., in the SIT) is displayed against codon translation speed (i.e., 1/RRT, Gardin et al. 2014). The correlation is −0.45, p < 0.001.

**Figure S4.**
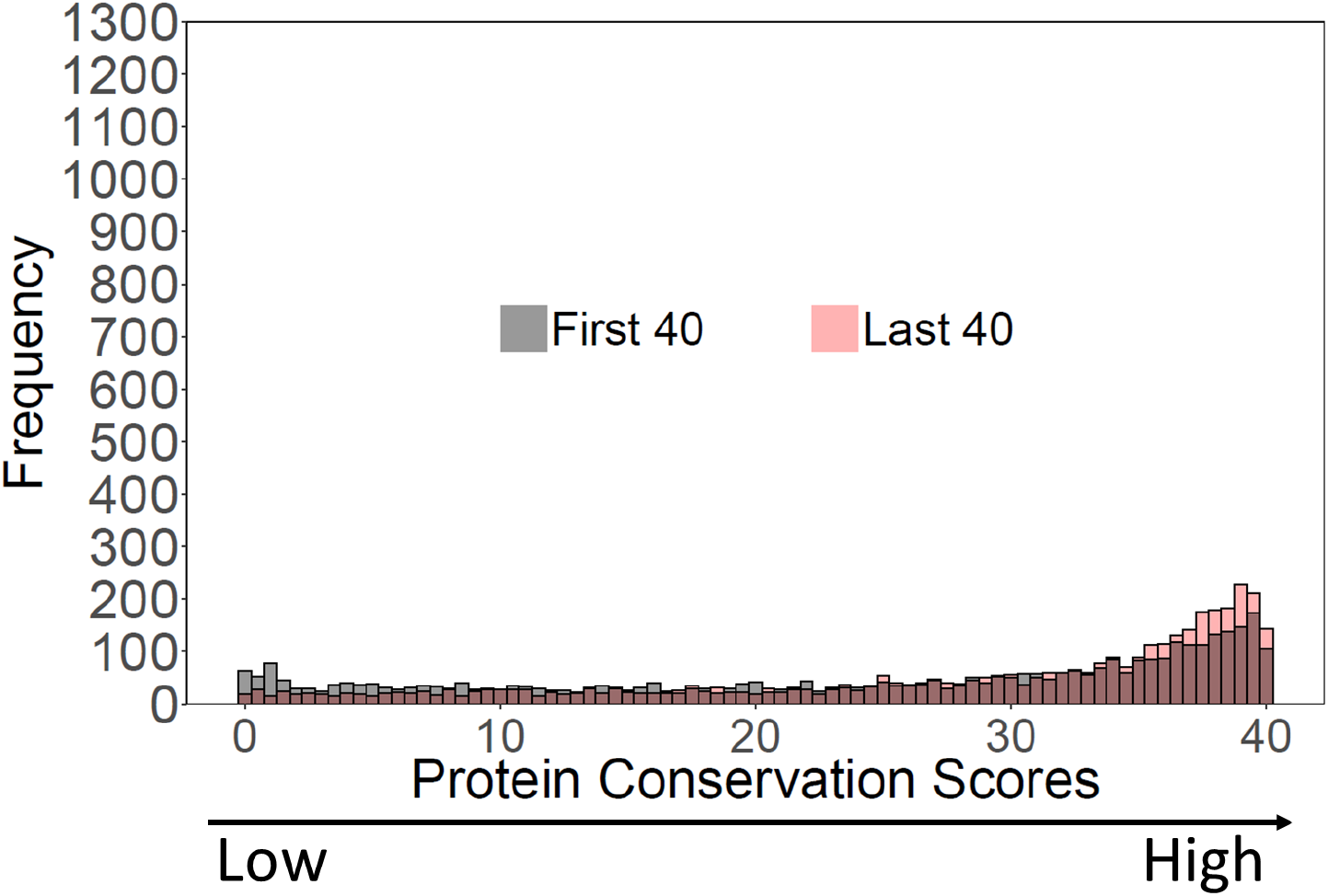
Comparison of conservation scores at the N- and C-termini. Grey, N-terminal conservation scores. Red, C-terminal conservation scores.

**Figure S5.**
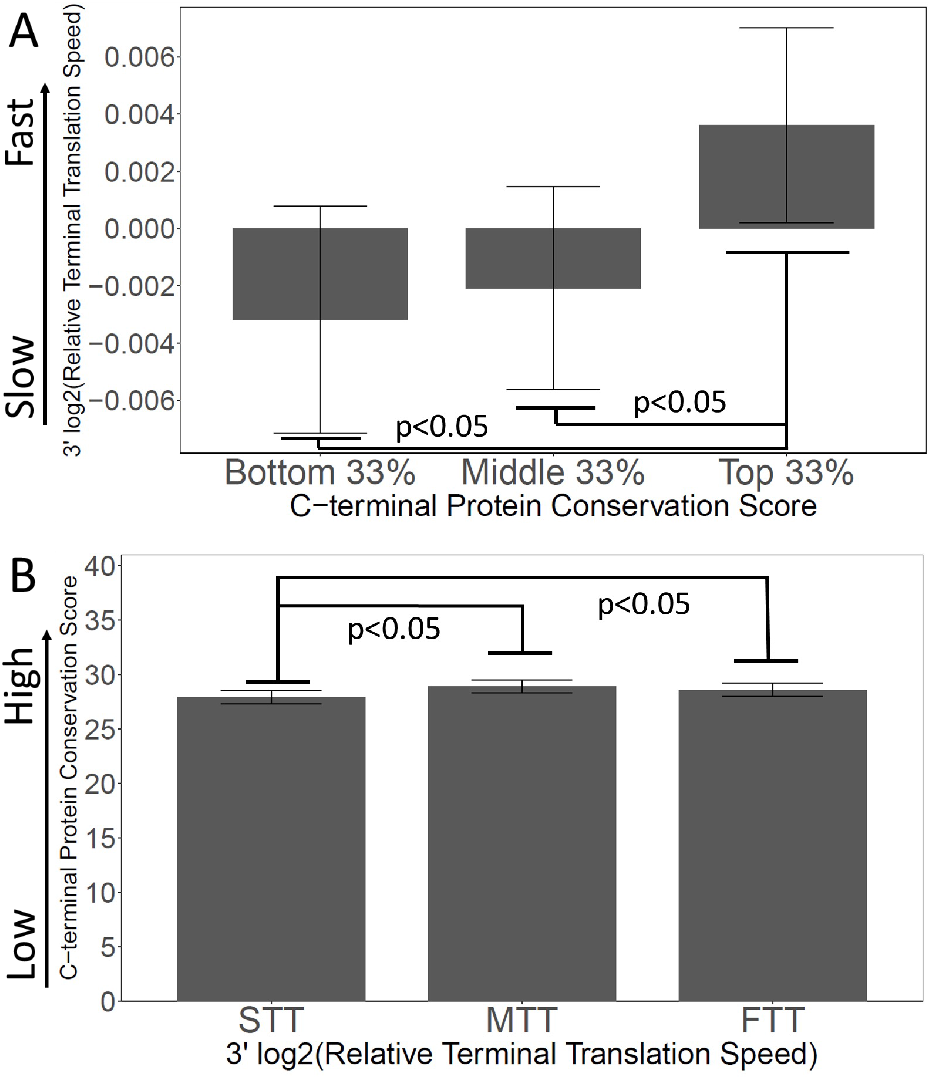
Slow 3’ Translation is correlated with poor C-terminal conservation. **A.** Proteins were grouped by their C-terminal conservation scores (top, middle, and bottom thirds), and then the terminal translation rate was plotted for each group. More conserved C-termini have faster terminal translation. **B.** Proteins were grouped by their C-terminal translation rate (top, middle, and bottom thirds), and then the C-terminal conservation scores were plotted for each group. Genes with faster terminal translation have more conserved C-termini.

**Figure S6.**
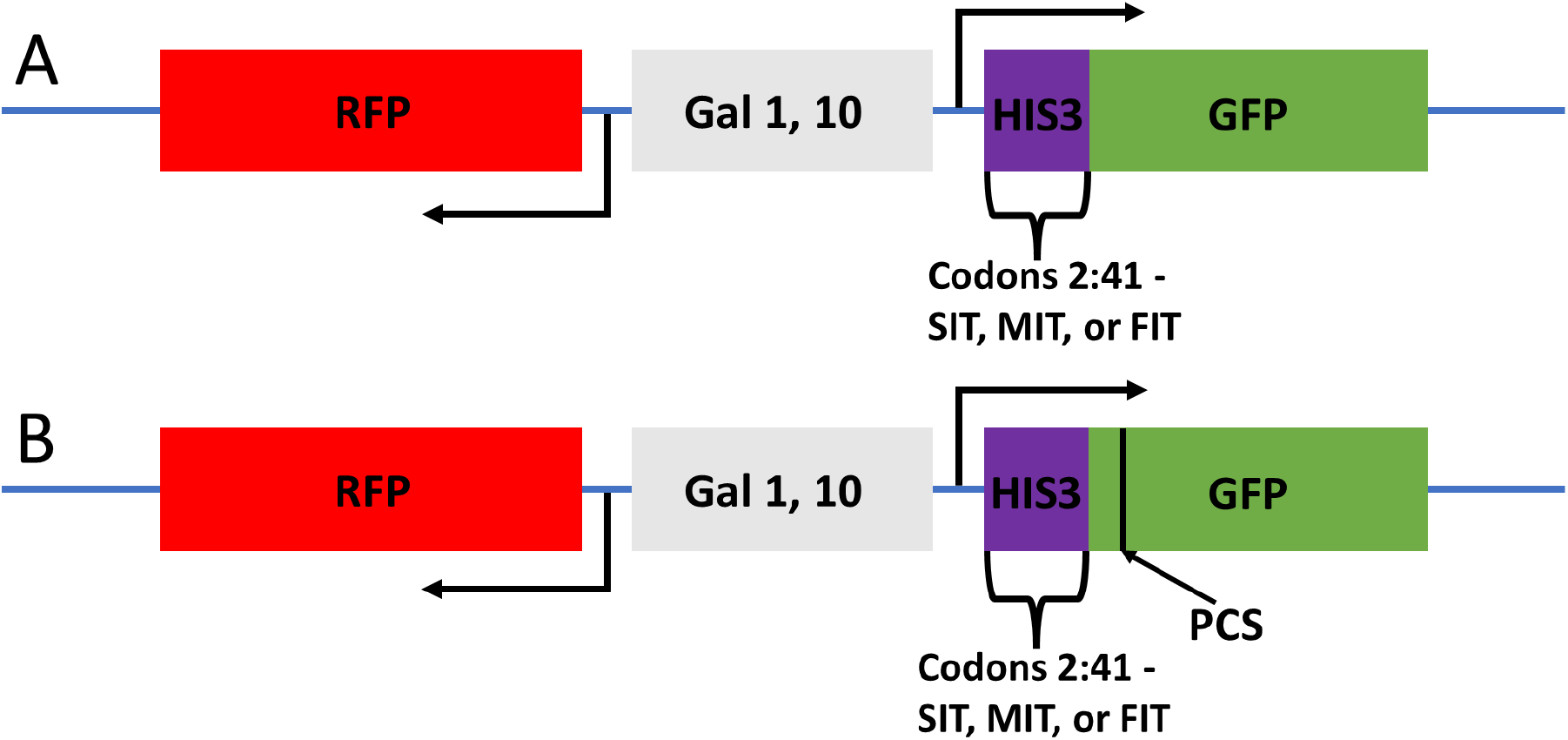
Structure of the GFP reporters. These reporters were adapted from Brule and Grayhack, 2016, and Gamble et al., 22016. A leader sequencer (purple), originally from *HIS3*, is appended upstream of GFP. **A.** For the first three constructs, recoding of some residues within the first 41 codons with synonymous codons gave either a slow (SIT), medium (MIT), or fast (FIT) initial translation rate. Protein sequences were preserved. **B.** Three analogous reporters were made with a putative ribosome collision site (PCS) at codon positions 68 and 69 (still upstream of GFP sequences). The PCS was the codon pair CGA-CGG, two rare Arg codons.

**Table S2.** Example conservation scores. Scores were calculated as described in Materials and Methods, but in this example, only for a subset of *Saccharomycotina*. “Total Hits” is the number of different proteins from the sub-phylum *Saccharomycotina* subset giving a BLAST bitscore of at least 50. “Hits in the first 40 amino acids” is the number of proteins (out of the proteins in the “Total Hits” columns) that had a BLAST alignment with an alignment score >200 matching any part of the first 40 amino acids of the query sequence (i.e., of *PCA1*, *NSR1*, etc.). “Query Start” is the range of amino acid positions in the Query protein where the BLAST alignments started. For instance, for *BUD5*, the 125 S*accharomycotina* homologs had BLAST alignments that started at positions between amino acid 211 and amino acid 420 on *S. cerevisiae BUD5*; none had an alignment starting within the first 40 amino acids. For *SNX41*, 65 of the 121 hits had an alignment beginning within the first 40 amino acids of *S. cerevisiae SNX41*. For *RPL12B*, all 121 of the *Saccharomycotina* homologs had BLAST alignments starting at amino acid 1 of *S. cerevisiae RPL12B*. The “Conservation Score” is the score calculated as described in Materials and Methods. Note that the number of hits varies in part because the genomes of the *Saccharomycotina* species were not all fully sequenced. Thus, *BNA2* likely has fifteen fewer hits than *TRP3* because the *BNA2* locus was not sequenced in some species. However, the number of hits does not affect the conservation score (see Table S2), as long as the number meets the qualifying minimum.

| Protein | Total Hits | Hits in the first 40 amino acids | Query Start | Conservation Score |
| --- | --- | --- | --- | --- |
| BUD5 | 125 | 0 | 211-420 | 0 |
| NSR1 | 122 | 0 | 61-264 | 0 |
| SNX41 | 121 | 65 | 2-199 | 10.7 |
| BNA2 | 107 | 106 | 15-86 | 20.8 |
| TRP3 | 122 | 122 | 1-23 | 30.6 |
| RPL12B | 121 | 121 | 1 | 40 |

**Table S3.** Calculation of a Conservation Score. “Query start position in BLAST alignment” is the amino acid residue of the *S. cerevisiae* query protein where a BLAST alignment (alignment score >200) begins with a protein of *Saccharomycotina*. “Proportion of hits with this Q-Start Position” is the proportion of qualifying *Saccharomycotina* hits (i.e., bitscore >50) that have their alignment begin at this position. “Weight” is multiplied by “Proportion”, and the sum is the Conservation Score.

| Query start position<br>in BLAST alignment | Weight to give | Proportion of hits<br>with this Q-Start<br>Position | Weight x Proportion |
| --- | --- | --- | --- |
| 1 | 40 | 0.22 | 8.8 |
| 2 | 39 | 0.01 | 0.39 |
| 3 | 38 | 0.01 | 0.38 |
| 4 | 37 | 0.02 | 0.74 |
| . | . | . | . |
| . | . | . | . |
| . | . | . | . |
| 38 | 3 | 0.03 | 0.09 |
| 39 | 2 | 0.03 | 0.06 |
| 40 | 1 | 0.04 | 0.04 |
|  |  |  | Sum (= Cons. Score) |

**Table S4.**
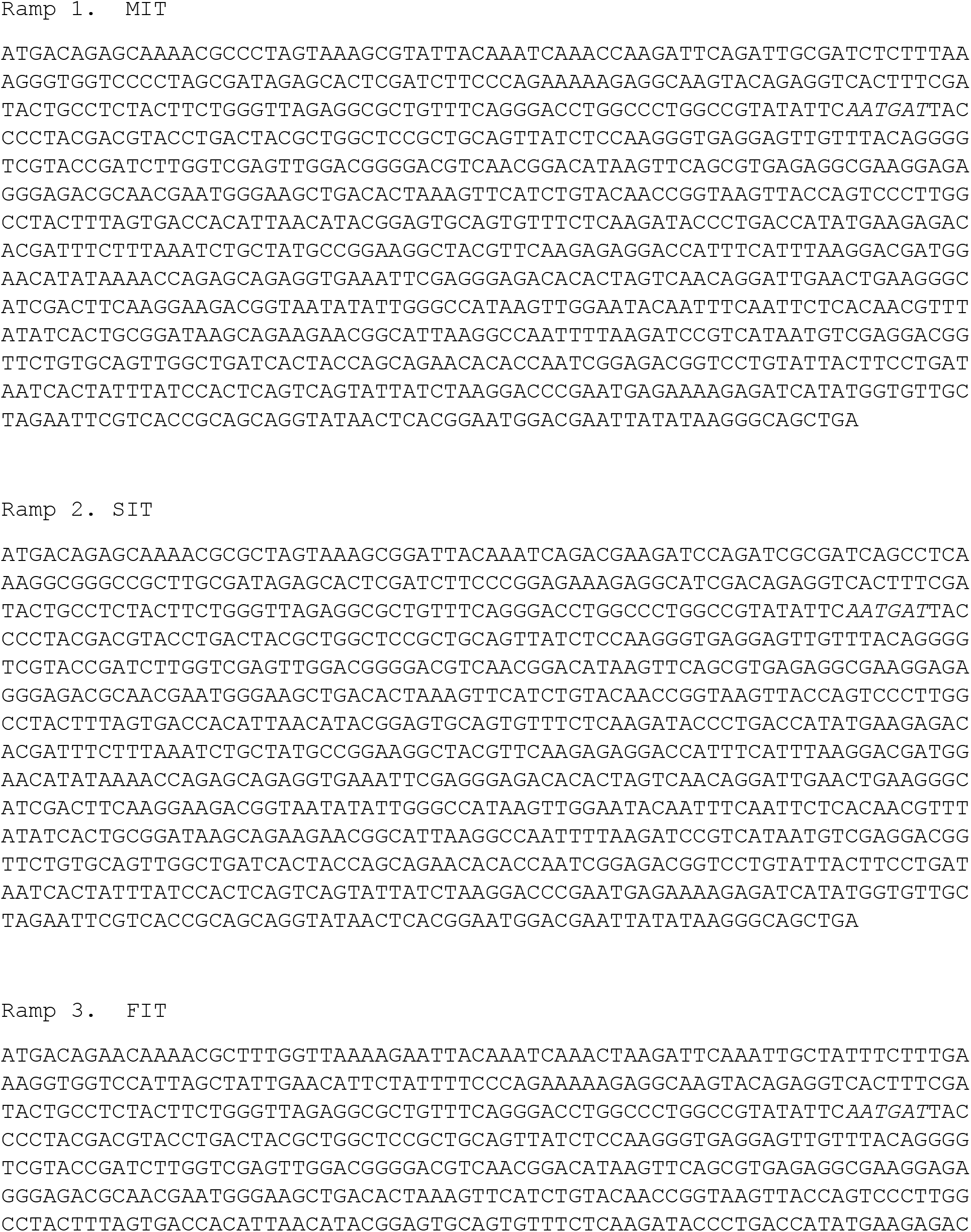
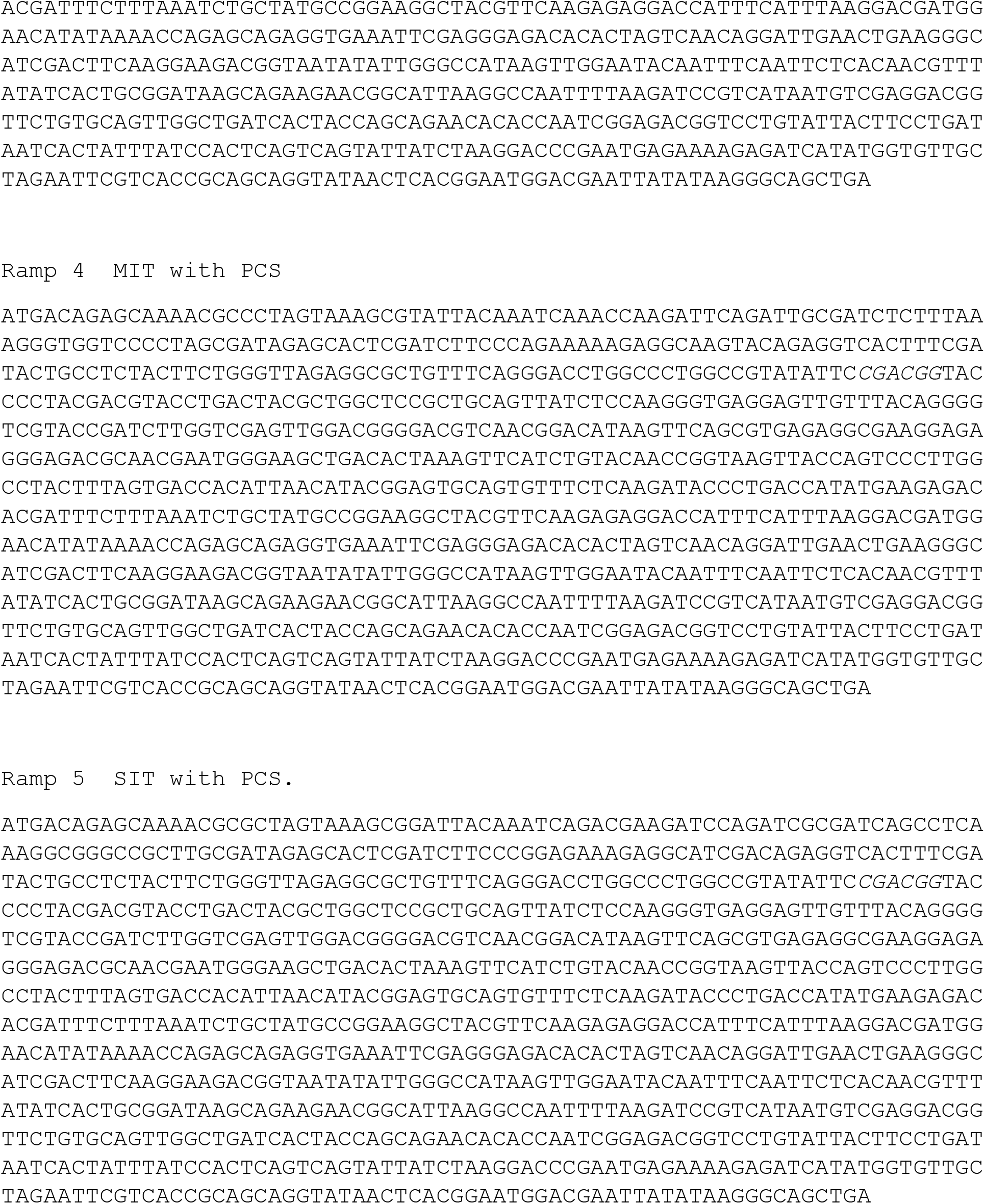
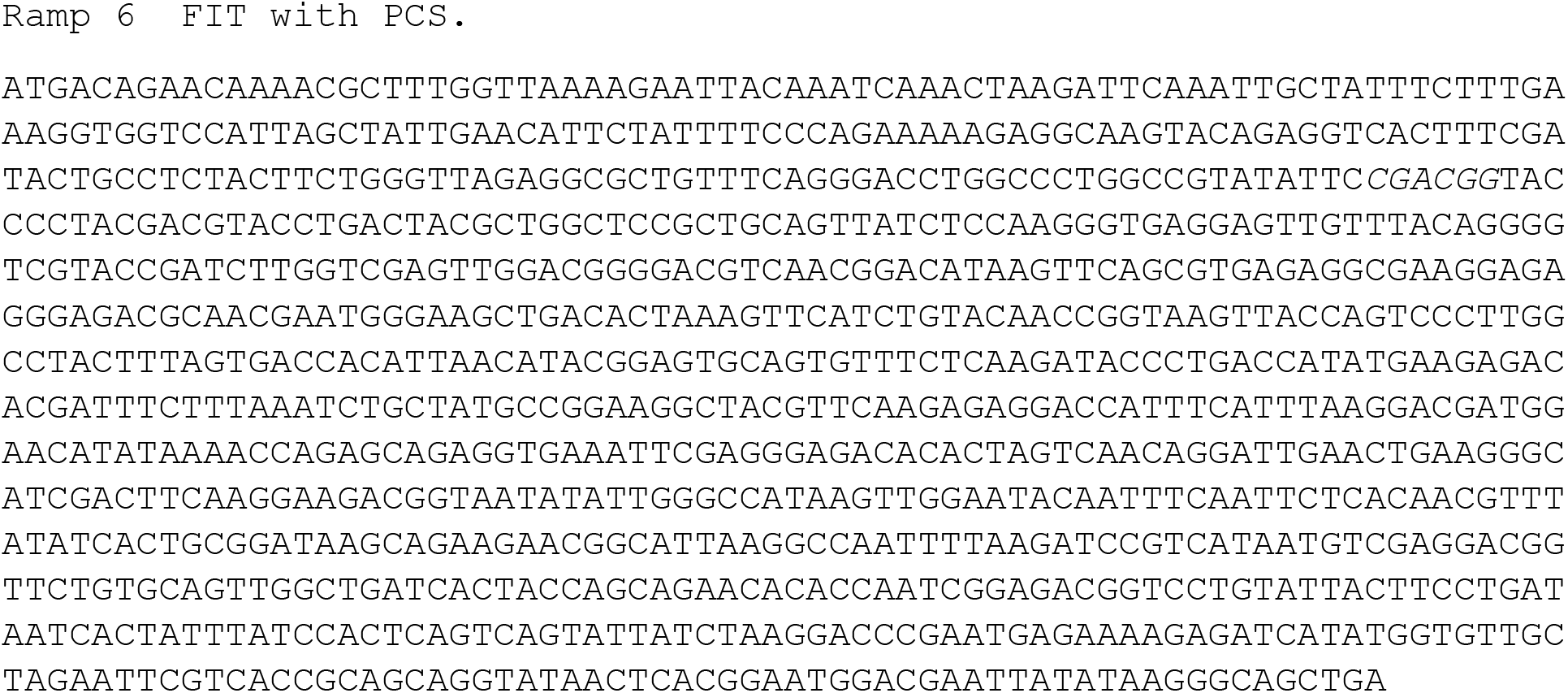
Ramp gene sequences.

