## Appendix 1: R code for analysis for "No Ramp Needed: Spandrels, Statistics, and a Slippery Slope"

---

Title: "No Ramp Needed: Spandrels, Statistics, and a Slippery Slope"

Authors: "Richard Sejour, Janet Leatherwood, Alisa Yurovsky, Bruce Futcher"

#Contact:

#files can be accessed at https://drive.google.com/drive/folders/1NIBFwiFD6gcPQ6E5FxRxXTV2iv9LNVSN?usp=sharing

output: pdf_document

---

#####Package citations.

```{r,echo=F,eval=F,include=F}

###All code, statistics, and figures were generated using The R Project for Statistical Computing (R Core Team, 2022). This code works best as an R Markdown file. If this is a text file, select all, copy everything in this file, and paste everything into a new R Markdown session. The following packages were used: rBLAST (version 0.99.2; Hahsler and Nagar, 2019), openxlsx (version 4.2.5.1; Schauberger and Walker, 2022), ggplot2 (version 3.3.6; Wickham, 2016), seqinr (version 4.2-16; Charif and Lobry, 2007), car (version 3.1-0; Fox and Weisberg, 2019), plyr (version 1.8.7; Wickham, 2011), readr (version 2.1.3; Wickham, Hester, and Bryan 2022), readxl (version 1.4.1; Wickham and Bryan, 2022), Rmisc (version 1.5.1; Hope, 2022), gtools (version 3.9.3; Bolker, Warnes, and Lumley 2022), scales (version 1.21; Wickham and Seidel 2022),stringr (version 1.5; Wickham 2022), and base (version 4.3; R Core Team, 2022).

# Bolker B, Warnes G, Lumley T (2022). _gtools: Various R Programming Tools_. https://CRAN.R-project.org/package=gtools. gtools package version 3.9.3.

# Charif D, Lobry J (2007). “SeqinR 1.0-2: a contributed package to the R project for statistical computing devoted to biological sequences retrieval and analysis.” In Bastolla U, Porto M, Roman H, Vendruscolo M (eds.), Structural approaches to sequence evolution: Molecules, networks, populations, series Biological and Medical Physics, Biomedical Engineering, 207-232. Springer Verlag, New York. ISBN : 978-3-540-35305-8. seqinr package version 4.2-16.

# Fox J, Weisberg S (2019). An R Companion to Applied Regression, Third edition. Sage, Thousand Oaks CA. https://socialsciences.mcmaster.ca/jfox/Books/Companion/. car package version 3.1-0.

# Hahsler M, Nagar A (2019). _rBLAST: R Interface for the Basic Local Alignment Search Tool_. https://github.com/mhahsler/rBLAST. rBLAST package version 0.99.2.

# Hope RM (2022). _Rmisc: Ryan Miscellaneous_. https://CRAN.R-project.org/package=Rmisc. Rmisc package version 1.5.1.

# R Core Team (2022). R: A language and environment for statistical computing. R Foundation for Statistical Computing, Vienna, Austria. URL https://www.R-project.org/. R version 4.2.1

# Schauberger P, Walker A (2022). _openxlsx: Read, Write and Edit xlsx Files_. https://CRAN.R-project.org/package=openxlsx. openxlsx package version 4.2.5.1.

# Wickham H (2011). “The Split-Apply-Combine Strategy for Data Analysis.” Journal of Statistical Software, 40(1), 1–29. https://www.jstatsoft.org/v40/i01/. plyr package version 1.8.7.

# Wickham H (2016). ggplot2: Elegant Graphics for Data Analysis. Springer-Verlag New York. ISBN 978-3-319-24277-4, https://ggplot2.tidyverse.org. ggplot2 package version 3.3.6.

# Wickham H (2022). stringr: Simple, Consistent Wrappers for Common String Operations. https://stringr.tidyverse.org, https://github.com/tidyverse/stringr. stringr package version 1.5.

# Wickham H, Bryan J (2022). _readxl: Read Excel Files_. https://CRAN.R-project.org/package=readxl. readxl package version 1.4.1.

# Wickham H, Hester J, Bryan J (2022). _readr: Read Rectangular Text Data_. https://CRAN.R-project.org/package=readr. readr package version 2.1.3.

# Wickham H, Seidel D (2022). scales: Scale Functions for Visualization. https://scales.r-lib.org, https://github.com/r-lib/scales. scales version 1.21.

```

###Installing the NCBI BLAST software and interfact for R.

```{r,echo=F,eval=F,include=F}

###This research used  NCBI BLAST software (version 2.13.0+). Download the latest version of the NCBI local BLAST tool from https://blast.ncbi.nlm.nih.gov/Blast.cgi?PAGE_TYPE=BlastDocs&DOC_TYPE=Download 

###The interface for the NCBI BLAST tool is controlled by the package rBLAST This is not found in the CRAN repository, so you have to download it using the code below. After you install the rBLAST package once, you never have to download it again unless there is an updated version. If the first code doesn't work, then use the one below it. This research uses version 0.99.2 of rBLAST.

remotes::install_github("mhahsler/rBLAST")

install.packages('rBLAST', repos = 'https://mhahsler.r-universe.dev')

###The default settings for the NCBI tool allocates an absurdly large amount of virtual memory which may exceed your computer specifications. As described in the biostars link below, for Windows OS you can set the environment variable BLASTDB_LMDB_MAP_SIZE to 10000000

#https://www.biostars.org/p/413294/

```

###Loading the packages and all functions. Run this chunk at the start of each session.

```{r, include=F}

###These are important packages that need to be loaded at the start of each session.

Packages<-c("rBLAST","openxlsx","ggplot2","seqinr","car","plyr","readr","readxl","Rmisc","gtools","scales","stringr")

lapply(Packages, library, character.only = TRUE)

###Most of the code requires an input directory containing the data to be analyzed, and an output directory where the analysis will be exported. R will give an error message if there are back-slashes \ in the directory path (which windows systems use in their directory path), so replace all back-slashes to forward slashes /. In some cases, the input directory will use the "outputdatabasedirectory" object because some data that has been analyzed will be subject to further analysis. The input directory can be changed to be an object in the environment.

inputdatabasedirectory<-"D:/Data Input/"

outputdatabasedirectory<-"D:/Data Output/"

###Below will dictate the size of the figures after you knit the file. If this is an RMarkdown file, then you can knit and export all of the figures and statistics without changing anything, as long as you keep all of the file names the same as in the examples.

knitr::opts_chunk$set(fig.height=7.5, fig.width=12,fig.align='center')

###Functions for calculating elapsed time. I created an experimental code that will calculate and report the total time that it takes to complete the code. Days, hours, minutes, and seconds are reported. Set your computer's internal clock to 24 hours. I tried to think of all of the conditions that may impact the elapsed time calculations (such as if the seconds at the start is lower than the seconds at the end), but I probably did not catch all the conditions. Leap years are also taken into account.

###Completion time output - start. "message.begin" which is an optional string of a message to be reported after the function is over.

timelapsebegin.function<-function(message.begin = "STARTED -"){
  
  timestartedtemp<-strftime(Sys.time())
  
  startyear<-unlist(strsplit(timestartedtemp,"-",fixed=T))[1]
  
  startmonth<-unlist(strsplit(timestartedtemp,"-",fixed=T))[2]
  
  startday<-unlist(strsplit(timestartedtemp,"-",fixed=T))[3]
  
  startday<-unlist(strsplit(startday," ",fixed=T))[1]
  
  starthour<-unlist(strsplit(timestartedtemp,":",fixed=T))[1]
  
  starthour<-unlist(strsplit(starthour," ",fixed=T))[2]
  
  startminutes<-unlist(strsplit(timestartedtemp,":",fixed=T))[2]
  
  startseconds<-unlist(strsplit(timestartedtemp,":",fixed=T))[3]
  
  months<-c("January","February","March","April","May","June","July","August","September","October","November","December")
  
  timestarted<-paste0(message.begin," ",months[as.numeric(startmonth)],", ",startday," ",startyear," - ",starthour,":",startminutes,":",startseconds)
  
  timebegin<-list(as.numeric(startyear),as.numeric(startmonth),as.numeric(startday),as.numeric(starthour),as.numeric(startminutes),as.numeric(startseconds),timestarted)
  
  return(timebegin)
}

###Completion time output - end. The arguments from timelapseend.function() are the output from the timelapsebegin.function() except for "message.fin" which is an optional string of a message to be reported after the function is over.

timelapseend.function<-function(startyear, startmonth, startday, starthour, startminutes, startseconds, timestarted, message.fin = "COMPLETED -"){
  
  timeendedtemp<-strftime(Sys.time())
  
  endfinyear<-unlist(strsplit(timeendedtemp,"-",fixed=T))[1]
  
  endfinmonth<-unlist(strsplit(timeendedtemp,"-",fixed=T))[2]
  
  endfinday<-unlist(strsplit(timeendedtemp,"-",fixed=T))[3]
  
  endfinday<-unlist(strsplit(endfinday," ",fixed=T))[1]
  
  endfinhour<-unlist(strsplit(timeendedtemp,":",fixed=T))[1]
  
  endfinhour<-unlist(strsplit(endfinhour," ",fixed=T))[2]
  
  endfinminutes<-unlist(strsplit(timeendedtemp,":",fixed=T))[2]
  
  endfinseconds<-unlist(strsplit(timeendedtemp,":",fixed=T))[3]
  
  months<-c("January","February","March","April","May","June","July","August","September","October","November","December")
  
  timeended<-paste0(months[as.numeric(endfinmonth)],", ",endfinday," ",endfinyear," - ",endfinhour,":",endfinminutes,":",endfinseconds)
    
  endfinseconds<-as.numeric(endfinseconds)
  
  endfinminutes<-as.numeric(endfinminutes)
  
  endfinhour<-as.numeric(endfinhour)
  
  endfinday<-as.numeric(endfinday)
  
  endfinmonth<-as.numeric(endfinmonth)
  
  endfinyear<-as.numeric(endfinyear)  
  
  if(startyear%%4!=0){
    
    calendar<-data.frame(months=c(rep(1,31),rep(2,28),rep(3,31),rep(4,30),rep(5,31),rep(6,30),rep(7,31),rep(8,31),rep(9,30),rep(10,31),rep(11,30),rep(12,31)),days=c(1:31,1:28,1:31,1:30,1:31,1:30,1:31,1:31,1:30,1:31,1:30,1:31),year=startyear)
  } else if(startyear%%4==0){
      
    calendar<-data.frame(months=c(rep(1,31),rep(2,29),rep(3,31),rep(4,30),rep(5,31),rep(6,30),rep(7,31),rep(8,31),rep(9,30),rep(10,31),rep(11,30),rep(12,31)),days=c(1:31,1:29,1:31,1:30,1:31,1:30,1:31,1:31,1:30,1:31,1:30,1:31),year=startyear)
  }
  
  yearrep<-1
  
  yeartemp<-startyear
  
  if(endfinyear-startyear>0){
    
    while(yearrep<(endfinyear-startyear)+1){
      
      yeartemp<-yeartemp+1
      
      if(yeartemp%%4!=0){
        
        calendartemp<-data.frame(months=c(rep(1,31),rep(2,28),rep(3,31),rep(4,30),rep(5,31),rep(6,30),rep(7,31),rep(8,31),rep(9,30),rep(10,31),rep(11,30),rep(12,31)),days=c(1:31,1:28,1:31,1:30,1:31,1:30,1:31,1:31,1:30,1:31,1:30,1:31),year=yeartemp)
      } else if(yeartemp%%4==0){
          
        calendartemp<-data.frame(months=c(rep(1,31),rep(2,29),rep(3,31),rep(4,30),rep(5,31),rep(6,30),rep(7,31),rep(8,31),rep(9,30),rep(10,31),rep(11,30),rep(12,31)),days=c(1:31,1:29,1:31,1:30,1:31,1:30,1:31,1:31,1:30,1:31,1:30,1:31),year=yeartemp)
      }
      
      calendar<-rbind(calendar,calendartemp)
      
      yearrep<-yearrep+1
    }
  }
  
  calendar$details<-""
  
  calendar$details[calendar$months==startmonth&calendar$days==startday&calendar$year==startyear]<-"start"
  
  calendar$details[calendar$months==endfinmonth&calendar$days==endfinday&calendar$year==endfinyear]<-"end"
  
  if(nrow(calendar[calendar$details=="start",])!=0 &nrow(calendar[calendar$details=="end",])!=0){
    
    calendar$details[which(calendar$details=="start"):which(calendar$details=="end")]<-"period"
  
    diffdays<-nrow(calendar[calendar$details=="period",])-1
  } else{
    
    diffdays<-1-1
  }
  
  if(startseconds>endfinseconds){
    
    elapseseconds<-60-(startseconds-endfinseconds)
  } else{
    
    elapseseconds<-endfinseconds-startseconds
  }
  
  elapsetotaltime<-((endfinhour*60)+(diffdays*24*60)+(endfinminutes))-((starthour*60)+(startminutes))
  
  elapsedays<-floor(elapsetotaltime/(24*60))
  
  elapsehours<-floor(((elapsetotaltime/(24*60))-elapsedays)*24)
  
  if(startseconds>endfinseconds){
    
    elapseminutes<-(round(((elapsetotaltime/(24*60))-elapsedays)*(60*24))-(elapsehours*60))-1
  
  } else{
      
    elapseminutes<-(round(((elapsetotaltime/(24*60))-elapsedays)*(60*24))-(elapsehours*60))
  }
  
  timefin<-paste0(message.fin," ",timeended," - ELAPSED TIME: ",   elapsedays," days ",elapsehours," hours ",elapseminutes," minutes ",elapseseconds," seconds")
  
  timefinoutput<-list(timestarted,timefin)
  
  return(timefinoutput)
}

####PART 1: INITIAL TRANSLATION SPEED CALCULATIONS

###Gene diagnostics to detect pseudogenes and dubious ORF from fasta file. "query.genes" is the list of genes as formatted from the read.fasta() function. "query.species" is an optional string for the species that the sequences originated from. "pseudogene.fasta.string" is an optional identifier in the fasta annotations that denote genes known to be pseudogenes. "dubiousORF.fasta.string" is the optional identifier in annotations that denote known dubious ORF. "fasta.updated" is an optional string for the last known date that the fasta file was uploaded to the queried website. "file.url" is the optional URL string to access the the fasta file. "notes" is an optional string for any purpose to accompany the output. "print.index" gives the option (True or False) to print the current numerical index in the fasta file that is being analyzed; for example, an index of 5 means that the fifth gene listed in the fasta file is currently being processed by the code. "message.start" is the message for the time elapse at the start of the function. "message.end" is the message for the time elapse after the function has completed.

checkORFfasta.function<-function(query.genes, query.species = "", pseudogene.fasta.string = "", dubiousORF.fasta.string = "", fasta.updated = "", file.url = "", notes = "", print.index = T, message.start = "STARTED -", message.end = "COMPLETED -"){
  
  ##Completion time output - start
  
  timebegin<-timelapsebegin.function(message.begin=message.start)
  
  print(timebegin[[7]])
  
  ##Code begins
  
  if(pseudogene.fasta.string!=""){
    
    query.ORFtemp2<-query.genes[grepl(pseudogene.fasta.string,getAnnot(query.genes),fixed = T)==T]
  } else{
    
    query.ORFtemp2<-list()
  }
  
  if(dubiousORF.fasta.string!=""){
    
    query.ORFtemp3<-query.genes[grepl(dubiousORF.fasta.string,getAnnot(query.genes),fixed = T)==T]
  } else{
    
    query.ORFtemp3<-list()
  }
  
  query.ORFtemp4<-query.genes[! query.genes %in% query.ORFtemp2]
  
  query.ORF<-query.ORFtemp4[! query.ORFtemp4 %in% query.ORFtemp3]
  
  ##Genuine ORF are made up of triplets of nucleotides that when divided by 3 are integers or whole numbers. This will check that all of the query ORF are divisible by 3, or else they are either truncated ORF sequences or are pseudogenes.
 
  xy<-1
  
  pseudogenes.check<-NULL
  
  while(xy<length(query.ORF)+1){
    
    nameslisttemp<-data.frame(Query.Species=query.species,Name=getName(query.ORF[xy]),Nucleotides.Represented=c2s(levels(factor(unlist(getSequence(query.ORF[xy]))))),Nucleotide_levels=length(levels(factor(unlist(getSequence(query.ORF[xy]))))),Nucleotides=length(unlist(getSequence(query.ORF[xy]))),Codons=((length(unlist(getSequence(query.ORF[xy]))))/3))
    
    if(nameslisttemp$Codons==round(nameslisttemp$Codons)){
      
      nameslisttemp$Integer<-"Yes"
    } else if(nameslisttemp$Codons!=round(nameslisttemp$Codons)){
      
      nameslisttemp$Integer<-"No"
    }
    
    nameslisttemp$Annotation=unlist(getAnnot(query.ORF[xy]))
    
    nameslisttemp$Dataset_updated=fasta.updated
    
    nameslisttemp$File=file.url
    
    nameslisttemp$Notes=notes
    
    pseudogenes.check=rbind(pseudogenes.check,nameslisttemp)
    
    if(print.index==T){
      
      print(xy)
    }
    
    xy=xy+1
  }
  
  query.ORFcds1<-query.ORF[pseudogenes.check$Name[pseudogenes.check$Integer=="No"|pseudogenes.check$Nucleotide_levels!=4]]
  
  query.ORFcdsonly<-query.ORF[!query.ORF %in% query.ORFcds1]
  
  if(nrow(pseudogenes.check[pseudogenes.check$Integer=="No",])==0&nrow(pseudogenes.check[pseudogenes.check$Integer=="Yes",])==length(query.ORF)){
    
    print("GENES ARE OKAY!!!")
  } else {
    
    base::warning("FASTA LIST HAS INCOMPLETE ORF OR THERE ARE PSEUDOGENES?? CHECK pseudogenes.check LIST TO SEE WHICH GENES HAVE NUCLEOTIDE LENGTHS THAT ARE NOT INTEGERS")
  }
  
  gene<-query.ORFcdsonly
  
  checkyeast.output<-list(query.ORFtemp2,query.ORFtemp3,pseudogenes.check,gene)
  
  names(checkyeast.output)[1] <- "annotated.pseudogenes"

  names(checkyeast.output)[2] <- "annotated.dubiousORF"
  
  names(checkyeast.output)[3] <- "pseudogenes.check"
  
  names(checkyeast.output)[4] <- "curated.genes"
  
  ##Completion time output - end
  
  timeended<-timelapseend.function(startyear=timebegin[[1]],startmonth=timebegin[[2]],startday=timebegin[[3]],starthour=timebegin[[4]],startminutes=timebegin[[5]],startseconds=timebegin[[6]],timestarted=timebegin[[7]],message.fin=message.end)
  
  print(timeended[[1]])
  
  print(timeended[[2]])
  
  return(checkyeast.output)
}

###Generating codon usage table. "query.genes" is the list of genes as formatted from the read.fasta() function. "query.species" is an optional string for the species that the sequences originated from."rrt.values" gives the option to assign a codon with its RRT value or codon-specific translation speed; this must be a data frame with 2 columns, and the column with the codons must have "Codons" as the column name. "fasta.updated" is an optional string for the last known date that the fasta file was uploaded to the queried website. "file.url" is the optional URL string in which the fasta can be accessed. "notes" is an optional string for any purpose to accompany the output. "save.file" gives the option (True or False) to save and export the results as an excel file directly into an assigned directory. If you choose to save the file using this code, then "outputfilename" is the string of the file name (default extension will be ".xlsx") for the output and "directorysave" should be a string for the directory to save the file. "print.index" gives the option (True or False) to print the current numerical index in the fasta file that is being analyzed; for example, an index of 5 means that the fifth gene listed in the fasta file is currently being processed by the code. "message.start" is the message for the time elapse at the start of the function. "message.end" is the message for the time elapse after the function has completed.

codonusagetable.function<-function(query.genes, query.species = "", rrt.values = 0, fasta.updated ="", file.url = "", notes = "", save.file = F, outputfilename = "codon usage table.xlsx", directorysave = outputdatabasedirectory, print.index = T, message.start = "STARTED -", message.end = "COMPLETED -"){
  
  ##Completion time output - start
  
  timebegin<-timelapsebegin.function(message.begin=message.start)
  
  print(timebegin[[7]])
  
  ##Code begins
  
  compiledcodonusageoutputframeone<-rep(list(NA),length(query.genes))
  
  generep<-1
  
  while (generep<(length(query.genes)+1)){
    
    if(print.index==T){
      
      print(generep)
    }
    
    dnasequence<-getSequence(query.genes[[generep]])
  
    firstposition<-1
    
    indexxxx<-1
    
    codonlistframeone<-rep(NA,(length(dnasequence))/3)
    
    while (firstposition<length(dnasequence)){
      codonlistframeone[indexxxx]<-toupper(c2s(dnasequence[firstposition:(firstposition+2)]))
      
      firstposition<-firstposition+3
      
      indexxxx<-indexxxx+1
    }
  
  compiledcodonusageoutputframeone[generep]<-list(codonlistframeone)
  
  names(compiledcodonusageoutputframeone)[generep] <- getName(query.genes[[generep]])
  
  generep<-generep+1
  }
  
  Globalcodons<-data.frame(table(unlist(compiledcodonusageoutputframeone)))
  
  names(Globalcodons)[1] <- "Codons"
  
  names(Globalcodons)[2] <- "Frame 1 (Coding) Observed Counts"
  
  Globalcodons$`Codon Proportion`<-Globalcodons$`Frame 1 (Coding) Observed Counts`/sum(Globalcodons$`Frame 1 (Coding) Observed Counts`)
  
  ##Translating codons into amino acid
  
  Globalcodons$`Amino Acid`<-"ZZZZZZ"
  
  codonx<-1
  
  while(codonx<(nrow(Globalcodons)+1)){
    
    Globalcodons$`Amino Acid`[codonx]<-translate(s2c(as.character(Globalcodons$Codons[codonx])))
    
    codonx<-codonx+1
  }
  
  ##Generating Order of synonymous codons
  
  Globalcodons$`Amino Acid`<-factor(Globalcodons$`Amino Acid`)
  
  Globalcodons$`AA proportion`<-"ZZZZZZZ"
  
  Globalcodons$`AA order`<-"ZZZZZZZ"
  
  compiled<-NULL
  
  generep<-1
  
  while(generep<length(levels(Globalcodons$`Amino Acid`))+1){
    
    aaco<-Globalcodons[Globalcodons$`Amino Acid` == levels(Globalcodons$`Amino Acid`)[generep],]
    
    aaco$`AA proportion`<-aaco$`Frame 1 (Coding) Observed Counts`/sum(aaco$`Frame 1 (Coding) Observed Counts`)
    
    aaco<-aaco[order(aaco$`AA proportion`),]
    
    aaco$`AA order`<-nrow(aaco):1
    aaco$`AA order`[1]<-"+"
    
    compiled<-rbind(compiled,aaco)
    
    generep<-generep+1
  }
  
  Globalcodons<-compiled
  
  Globalcodons$Symbol<-paste0(Globalcodons$Codons,":",Globalcodons$`AA order`,":",Globalcodons$`Amino Acid`)
  
  ##Adding RRT values, aka codon-specific translation speed
  
  if(length(rrt.values)>1){
    
    rrt_values<-rrt.values
    
    Globalcodons<-join(Globalcodons,rrt_values,by="Codons",type="full", match="all")
  }
  
  Globalcodons$Query.Species<-query.species
  
  Globalcodons$ORF<-length(compiledcodonusageoutputframeone)
  
  Globalcodons$Fasta.Updated<-fasta.updated
  
  Globalcodons$Fasta.URL<-file.url
  
  Globalcodons$Notes<-notes
  
  Globalcodons<-Globalcodons[order(Globalcodons$Codons),]
  
  if(save.file==T){
    
    wb<-createWorkbook()
    
    addWorksheet(wb, "Codon Usage")
    
    addWorksheet(wb, "Sampled ORF")
    
    writeData(wb,sheet="Codon Usage",x=Globalcodons)
    
    writeData(wb,sheet="Sampled ORF",x=data.frame(Name=names(compiledcodonusageoutputframeone)))
    
    saveWorkbook(wb, paste0(directorysave,outputfilename), overwrite = T)
  }
  
  ##Completion time output - end
  
  timeended<-timelapseend.function(startyear=timebegin[[1]],startmonth=timebegin[[2]],startday=timebegin[[3]],starthour=timebegin[[4]],startminutes=timebegin[[5]],startseconds=timebegin[[6]],timestarted=timebegin[[7]],message.fin=message.end)
  
  print(timeended[[1]])
  
  print(timeended[[2]])
  
  return(Globalcodons)
}

###Generating translation speed at the N-termini and C-termini. "query.genes" is the list of genes as formatted from the read.fasta() function. "query.species" is an optional string for the species that the sequences originated from. "rampzonelength.fiveprime" indicates the codon length of the window to measure translation speed at the N-termini. "rampzonelength.threeprime" indicates the codon length of the window to measure translation speed at the C-termini. "codons.translationspeed" is the data frame of codons with corresponding translation speeds, which must be formatted as a dataframe; all codons in the data frame must have a numerical codon-specific translation speed, the column with the codons must have "Codons" as the column name, and the column with the codon-specific translation speeds must have "RRT" as the column name. "save.file" gives the option (True or False) to save the translation speed output as an excel file with 3 tabs (default extension will be ".xlsx"). If you choose to save the file using this code, then "outputfilename" is the string of the file name for the output and "directorysave" should be a string for the directory to save the file. "print.index" gives the option (True or False) to print the current numerical index in the fasta file that is being analyzed; for example, an index of 5 means that the fifth gene listed in the fasta file is currently being processed by the code. "message.start" is the message for the time elapse at the start of the function. "message.end" is the message for the time elapse after the function has completed.

translationspeed.function<-function(query.genes, query.species = "", rampzonelength.fiveprime = 40, rampzonelength.threeprime = 40, codons.translationspeed, save.file = F, outputfilename = "N-termini and C-termini translation speed.xlsx", directorysave = outputdatabasedirectory, print.index = T, message.start = "STARTED -",message.end = "COMPLETED -"){

  ##Completion time output - start
  
  timebegin<-timelapsebegin.function(message.begin=message.start)
  
  print(timebegin[[7]])

  ##Code starts
  
  Globalcodons<-codons.translationspeed
  
  generep<-1

  sampledgenes<-data.frame(Name="ZZZZZ",Sampled="ZZZZZ",Nucleotides=99999,Query.Annotation="ZZZZZ",Number=1:length(gene))
  
  rampcomp<-data.frame(Name="ZZZZZ",AverageRRTFirst=99999,AverageRRTFirstnostartcodon=99999,AverageRRTentireminusfirst=99999,AverageRRTThreePrime=99999,AverageRRTentireminuslast=99999,AverageRRTentiregene=99999,AverageRRTentiregenenostartcodon=99999,Nucleotides=99999,Query.Annotation="ZZZZZ",Number=1:length(gene))
  
  while (generep<(length(gene)+1)){
    
    if(print.index==T){
      
      print(generep)
    }
    
    orfsample<-gene[generep]
    
    dnasequence<-unlist(getSequence(orfsample))
    
    genename<-getName(orfsample)
  
    if(length(dnasequence)>(3*rampzonelength.fiveprime)+3&length(dnasequence)>(3*rampzonelength.threeprime)+3){
        
      ##This starts index at start codon to the end of the gene omitting the last (stop) codon

      firstposition<-1
      
      indexxxx<-1
      
      entiregene<-rep(NA,(length(dnasequence)/3)-3)
      
      while (firstposition<length(dnasequence)-3){
        
        entiregene[indexxxx]<-Globalcodons$RRT[Globalcodons$Codons==toupper(c2s(dnasequence[firstposition:(firstposition+2)]))]
        
        firstposition<-firstposition+3
        
        indexxxx<-indexxxx+1
      }
      
      sampledgenes$Name[generep]<-genename
      
      sampledgenes$Sampled[generep]<-"Sampled"
      
      sampledgenes$Nucleotides[generep]<-length(dnasequence)
      
      sampledgenes$Query.Annotation[generep]<-unlist(getAnnot(orfsample))
 
      rampcomp$Name[generep]<-genename
      
      rampcomp$AverageRRTFirst[generep]<-mean(entiregene[1:rampzonelength.fiveprime])
      
      rampcomp$AverageRRTFirstnostartcodon[generep]<-mean(entiregene[2:rampzonelength.fiveprime])
      
      rampcomp$AverageRRTThreePrime[generep]<-mean(entiregene[(length(entiregene)-(rampzonelength.threeprime)+1):length(entiregene)])
      
      rampcomp$AverageRRTentireminusfirst[generep]<-mean(entiregene[(rampzonelength.fiveprime+1):length(entiregene)])
      
      rampcomp$AverageRRTentireminuslast[generep]<-mean(entiregene[2:(length(entiregene)-(rampzonelength.threeprime))])
      
      rampcomp$AverageRRTentiregene[generep]<-mean(entiregene)
      
      rampcomp$AverageRRTentiregenenostartcodon[generep]<-mean(entiregene[2:length(entiregene)])
      
      rampcomp$Nucleotides[generep]<-length(dnasequence)
      
      rampcomp$Query.Annotation[generep]<-unlist(getAnnot(orfsample))
  
    ##Rejected because ORF is too short
      
    } else if (length(dnasequence)<(3*rampzonelength.fiveprime)+3){
        
      sampledgenes$Name[generep]<-genename
        
      sampledgenes$Sampled[generep]<-"Too Short"
        
      sampledgenes$Nucleotides[generep]<-length(dnasequence)
        
      sampledgenes$Query.Annotation[generep]<-unlist(getAnnot(orfsample))
    } else if (length(dnasequence)==(3*rampzonelength.fiveprime)+3){
        
      sampledgenes$Name[generep]<-genename
        
      sampledgenes$Sampled[generep]<-"Too Short"
        
      sampledgenes$Nucleotides[generep]<-length(dnasequence)
        
      sampledgenes$Query.Annotation[generep]<-unlist(getAnnot(orfsample))
    } else if (length(dnasequence)<(3*rampzonelength.threeprime)+3){
        
      sampledgenes$Name[generep]<-genename
        
      sampledgenes$Sampled[generep]<-"Too Short"
        
      sampledgenes$Nucleotides[generep]<-length(dnasequence)
        
      sampledgenes$Query.Annotation[generep]<-unlist(getAnnot(orfsample))
    } else if (length(dnasequence)==(3*rampzonelength.threeprime)+3){
        
      sampledgenes$Name[generep]<-genename
        
      sampledgenes$Sampled[generep]<-"Too Short"
        
      sampledgenes$Nucleotides[generep]<-length(dnasequence)
        
      sampledgenes$Query.Annotation[generep]<-unlist(getAnnot(orfsample))
    }
  
    generep=generep+1
  }
  
  rampcomp$RatiovsRest=rampcomp$AverageRRTFirst/rampcomp$AverageRRTentireminusfirst
  
  rampcomp$RationostartcodonvsRest=rampcomp$AverageRRTFirstnostartcodon/rampcomp$AverageRRTentireminusfirst
  
  rampcomp$log2RationostartcodonvsRest<-log2(rampcomp$RationostartcodonvsRest)
  
  rampcomp$log2RatiothreeprimevsResttemp<-rampcomp$AverageRRTThreePrime/rampcomp$AverageRRTentireminuslast
  
  rampcomp$log2RatiothreeprimevsRest<-log2(rampcomp$log2RatiothreeprimevsResttemp)
  
  rampcomp$log2Rationostartcodonvsthreeprime<-log2(rampcomp$AverageRRTFirstnostartcodon/rampcomp$AverageRRTThreePrime)
  
  rampcomp$Query.Species<-query.species
  
  rampcomp<-rampcomp[,c("Name","Nucleotides","AverageRRTFirst","AverageRRTFirstnostartcodon","AverageRRTentireminusfirst","AverageRRTThreePrime","AverageRRTentireminuslast","AverageRRTentiregene","AverageRRTentiregenenostartcodon","RatiovsRest","RationostartcodonvsRest","log2RationostartcodonvsRest","log2RatiothreeprimevsRest","log2Rationostartcodonvsthreeprime","Query.Species","Query.Annotation")]
  
  names(rampcomp)[names(rampcomp) == "AverageRRTFirst"] <- paste0("AverageRRTFirst",rampzonelength.fiveprime)
  
  names(rampcomp)[names(rampcomp) == "AverageRRTFirstnostartcodon"] <- paste0("AverageRRTFirst",rampzonelength.fiveprime,"nostartcodon")
  
  names(rampcomp)[names(rampcomp) == "AverageRRTentireminusfirst"] <- paste0("AverageRRTentireminusfirst",rampzonelength.fiveprime)
  
  names(rampcomp)[names(rampcomp) == "AverageRRTThreePrime"] <- paste0("AverageRRTThreePrime",rampzonelength.threeprime)
  
  names(rampcomp)[names(rampcomp) == "AverageRRTentireminuslast"] <- paste0("AverageRRTentireminuslast",rampzonelength.threeprime)
  
  names(rampcomp)[names(rampcomp) == "RatiovsRest"] <- paste0("Ratio",rampzonelength.fiveprime,"vsRest")
  
  names(rampcomp)[names(rampcomp) == "RationostartcodonvsRest"] <- paste0("Ratio",rampzonelength.fiveprime,"nostartcodonvsRest")
  
  names(rampcomp)[names(rampcomp) == "log2RationostartcodonvsRest"] <- paste0("log2Ratio",rampzonelength.fiveprime,"nostartcodonvsRest")
  
  names(rampcomp)[names(rampcomp) == "log2RatiothreeprimevsRest"] <- paste0("log2Ratiothreeprime",rampzonelength.threeprime,"vsRest")
  
  names(rampcomp)[names(rampcomp) == "log2Rationostartcodonvsthreeprime"] <- paste0("log2Ratio",rampzonelength.fiveprime,"nostartcodonvsthreeprime",rampzonelength.threeprime)
  
  sampledgenes$Query.Species<-query.species

  sampledgenes<-sampledgenes[,c("Name","Nucleotides","Sampled","Query.Species","Query.Annotation")]
  
  omittedORF<-sampledgenes[sampledgenes$Sampled=="Too Short",]
  
  names(omittedORF)[names(omittedORF) == "Sampled"] <- "Rejected"
  
  rampcomp<-rampcomp[rampcomp$Name!="ZZZZZ",]
  
  rampcomp$Nucleotides<-as.numeric(rampcomp$Nucleotides)
  
  if(((nrow(rampcomp)+nrow(omittedORF))==length(gene))&((nrow(omittedORF)+nrow(sampledgenes[!sampledgenes$Sampled=="Too Short",]))==length(gene))){
    
    if(save.file==T){
      
      wb<-createWorkbook()
      
      addWorksheet(wb, paste0("First ",rampzonelength.fiveprime," codons"))
     
      addWorksheet(wb, paste0("Sampled ORF"))
      
      addWorksheet(wb, paste0("Omitted ORF"))
      
      writeData(wb,sheet=paste0("First ",rampzonelength.fiveprime," codons"),x=rampcomp)
      
      writeData(wb,sheet=paste0("Sampled ORF"),x=sampledgenes[sampledgenes$Sampled=="Sampled",])
      
      writeData(wb,sheet=paste0("Omitted ORF"),x=omittedORF)
      
      saveWorkbook(wb, paste0(directorysave,outputfilename), overwrite = T)
    }
      
    print("SUCCESS!!!")
  
  } else {
    
    base::warning("SOMETHING IS WRONG??")
  }
  
  output<-list(rampcomp,sampledgenes[sampledgenes$Sampled=="Sampled",],omittedORF)
  
  names(output)[1]<-paste0("First ",rampzonelength.fiveprime," codons")
  
  names(output)[2]<-"Sampled ORF"
  
  names(output)[3]<-"Omitted ORF"

  ##Completion time output - end
  
  timeended<-timelapseend.function(startyear=timebegin[[1]],startmonth=timebegin[[2]],startday=timebegin[[3]],starthour=timebegin[[4]],startminutes=timebegin[[5]],startseconds=timebegin[[6]],timestarted=timebegin[[7]],message.fin=message.end)
  
  print(timeended[[1]])
  
  print(timeended[[2]])

  return(output)
}

####PART 2: PROTEIN BLASTS AND CONSERVATION SCORES CALCULATIONS

###Function for making BLASTS

#blast_help(type = "blastp")
#To check if NCBI tool is correctly installed use Sys.which("blastp")
#To find out version installed use system("blastp -version")

###BLASTS of Full-Length Proteins

###The default settings of the NCBI BLAST will be used, except that the maximum number of alignments (1000000000) will be returned for every BLAST, otherwise the default is 500 alignments. The annotations for the subject hits are added after all BLASTS have been conducted. Don't worry if you get an error message that reads "Error in read.table(outfile, sep = ",", quote = "") : no lines available in input"; that just means that a query sequence had no BLAST hits, which will be recorded in the BLAST statistics. To find out the meaning of qstart, sstart, send, etc. view this link https://www.ncbi.nlm.nih.gov/books/NBK279684/  "queryproteinsstringset" is the query list of proteins which must be formatted using the readAAStringSet() function. "querydescriptives" must be a data frame containing the names, annotations, and other important features of the query sequences; the BLAST function refers to specific column names in the "querydescriptives" data frame, so refer to the example BLAST function to know exactly how to format the column names or else this code won't work. "make.database" gives the option (True or False) if you want to create the subject database in your home directory; each subject database is permanent and will not leave your home directory if you quit R session. "load.database" gives the option (True or False) to load all of the proteins in the local subject database as an object in the environment using the read.fasta() function; this takes a long time because the subject database will oftentimes have hundreds of thousands of sequences. "proteindatabase" is the string containing the directory path where the local subject database is located; by default this should be your home directory since the NCBI tool automatically creates the local database in your home directory. "query.species" is an optional string for the species that the query sequences originated from. "proteinspan" is an optional string to convey the region of the query sequence that will be BLASTed. "number.of.proteins" is a numerical value of how many proteins are queried by the BLAST function; this value must be the exact same as the length of proteins in "querydescriptives". "bitscore.threshold" is the minimum bitscore value from each sequence alignment to include in the BLAST results; the total number of sequence alignments a query has will be recorded regardless of bitscore value, but all BLAST descriptions (alignment regions, subject annotations, qstart, E value, etc) from alignments the same or higher than "bitscore.threshold" will be returned. "rawblastresultsname" should be the string of the name for the data frame containing the raw BLAST results. "sampledblastresultsname" is the string for the name of the data frame that contain statistics regarding each query protein BLAST results. "save.file" gives the option (True or False) to save the BLAST output as an R file with the extension ".RData". If you choose to save the file using this code, then "rdatablastname" is the string of the file name (requires extension ".RData") for the output and "directorysave" should be a string for the directory to save the file. "print.index" gives the option (True or False) to print the current numerical index in the fasta file that is being analyzed; for example, an index of 5 means that the fifth protein listed in the fasta file is currently being BLASTed. "save.csv" gives the option (True or False) to save and export all of the BLAST results as a file with the .csv extension. If you choose to export all of the BLAST results as csv, then two csv files will be saved; "blastfilename" is the string for the name of csv file with the raw BLAST results, and "blastreportname" is the string for the name of the csv file with the statistics regarding each query protein BLAST results. "message.start" is the message for the time elapse at the start of the function. "message.end" is the message for the time elapse after the function has completed.

proteinBLAST.function<-function(queryproteinsstringset, querydescriptives, make.database = T, load.database = T, proteindatabase,query.species = "", proteinspan = "Full Length", number.of.proteins, bitscore.threshold = 50, rawblastresultsname = "rawblastresults", sampledblastresultsname = "sampledblastresults" , save.file = F, directorysave = outputdatabasedirectory, rdatablastname = "Protein BLASTS List.RData", save.csv = F, blastfilename = "rawblastresults.csv", blastreportname = "sampledblastresults.csv", print.index = T, message.start = "STARTED -", message.end = "COMPLETED -"){
  
  ##Completion time output - start
  
  timebegin<-timelapsebegin.function(message.begin=message.start)
  
  print(timebegin[[7]])
  
  ##Making and loading subject database

  if (make.database==T){
    
    rBLAST::makeblastdb(proteindatabase,dbtype = "prot",args = "")
    
    tempmess<-timelapseend.function(startyear=timebegin[[1]],startmonth=timebegin[[2]],startday=timebegin[[3]],starthour=timebegin[[4]],startminutes=timebegin[[5]],startseconds=timebegin[[6]],timestarted=timebegin[[7]],message.fin="CREATED SUBJECT DATABASE - ")

    print(tempmess[[2]])
    
  } else if (make.database!=T){
    
    tempmess<-timelapseend.function(startyear=timebegin[[1]],startmonth=timebegin[[2]],startday=timebegin[[3]],starthour=timebegin[[4]],startminutes=timebegin[[5]],startseconds=timebegin[[6]],timestarted=timebegin[[7]],message.fin="DID NOT MAKE SUBJECT DATABASE - ")

    print(tempmess[[2]])
  }

  if (load.database==T){
    
    subjectdatabase<-seqinr::read.fasta(proteindatabase,seqtype="AA")
    
    tempmess<-timelapseend.function(startyear=timebegin[[1]],startmonth=timebegin[[2]],startday=timebegin[[3]],starthour=timebegin[[4]],startminutes=timebegin[[5]],startseconds=timebegin[[6]],timestarted=timebegin[[7]],message.fin="SUBJECT SEQUENCES ADDED - ")

    print(tempmess[[2]])
  } else if (load.database!=T){
    
    tempmess<-timelapseend.function(startyear=timebegin[[1]],startmonth=timebegin[[2]],startday=timebegin[[3]],starthour=timebegin[[4]],startminutes=timebegin[[5]],startseconds=timebegin[[6]],timestarted=timebegin[[7]],message.fin="DID NOT LOAD SUBJECT SEQUENCES - ")

    print(tempmess[[2]])
  }

  blast.table<-rBLAST::blast(db=proteindatabase,type = "blastp")
  
  tempmess<-timelapseend.function(startyear=timebegin[[1]],startmonth=timebegin[[2]],startday=timebegin[[3]],starthour=timebegin[[4]],startminutes=timebegin[[5]],startseconds=timebegin[[6]],timestarted=timebegin[[7]],message.fin="BLAST in process...")

  print(tempmess[[2]])
  
  ##Protein BLAST code begins
  
  rawblastresults=NULL
  
  sampledblastresults<-data.frame(Name="ZZZZZ",Query.Species=query.species,Total.Hits=9999999,Highest.Bitscore=9999999,Bitscore50.Hits=9999999,Amino.Acids=9999999,Query.Span=proteinspan,Date.of.BLAST="ZZZZZ",Time.of.BLAST="ZZZZZ",Annotation="ZZZZZ",Number=1:length(queryproteinsstringset))
  
  generep<-1
  
  while(generep <(number.of.proteins+1)){
  
    if(print.index==T){
      
      print(generep)
    }
    
    timestartedtemp<-strftime(Sys.time())
    
    startyear<-unlist(strsplit(timestartedtemp,"-",fixed=T))[1]
    
    startmonth<-unlist(strsplit(timestartedtemp,"-",fixed=T))[2]
    
    startday<-unlist(strsplit(timestartedtemp,"-",fixed=T))[3]
    
    startday<-unlist(strsplit(startday," ",fixed=T))[1]
    
    starthour<-unlist(strsplit(timestartedtemp,":",fixed=T))[1]
    
    starthour<-unlist(strsplit(starthour," ",fixed=T))[2]
    
    startminutes<-unlist(strsplit(timestartedtemp,":",fixed=T))[2]
    
    startseconds<-unlist(strsplit(timestartedtemp,":",fixed=T))[3]
    
    months<-c("January","February","March","April","May","June","July","August","September","October","November","December")
    
    blast.table2<-predict(blast.table, queryproteinsstringset[queryproteinsstringset@ranges@NAMES==querydescriptives$stringset.name[generep]],custom_format= 'qaccver qlen saccver slen pident length mismatch gapopen qstart qend sstart send evalue score bitscore qseq sseq',BLAST_args="-num_alignments 1000000000")
  
    if(nrow(blast.table2>0)){
      
      blast.table2$Query.Species<-query.species
      
      blast.table2$Query.Span<-proteinspan
      
      blast.table2$Date.of.BLAST<-paste0(months[as.numeric(startmonth)],", ",startday," ",startyear)
    
      blast.table2$Time.of.BLAST<-paste0(starthour,":",startminutes,":",startseconds)
      
      rawblastresults<-rbind(rawblastresults,blast.table2)
      
      sampledblastresults$Name[generep]<-querydescriptives$gene.name[generep]
      
      sampledblastresults$Total.Hits[generep]<-nrow(blast.table2)
      
      sampledblastresults$Highest.Bitscore[generep]<-blast.table2$bitscore[1]
      
      sampledblastresults$Bitscore50.Hits[generep]<-nrow(blast.table2[!blast.table2$bitscore<bitscore.threshold,])
      
      sampledblastresults$Amino.Acids[generep]<-querydescriptives$Amino.Acids[generep]
      
      sampledblastresults$Date.of.BLAST[generep]<-paste0(months[as.numeric(startmonth)],", ",startday," ",startyear)
      
      sampledblastresults$Time.of.BLAST[generep]<-paste0(starthour,":",startminutes,":",startseconds)
      
      sampledblastresults$Annotation[generep]<-querydescriptives$annotation[generep]
    } else{
      
      sampledblastresults$Name[generep]<-querydescriptives$gene.name[generep]
    
      sampledblastresults$Total.Hits[generep]<-nrow(blast.table2)
    
      sampledblastresults$Highest.Bitscore[generep]<-0
    
      sampledblastresults$Bitscore50.Hits[generep]<-nrow(blast.table2[!blast.table2$bitscore<bitscore.threshold,])
    
      sampledblastresults$Amino.Acids[generep]<-querydescriptives$Amino.Acids[generep]
    
      sampledblastresults$Date.of.BLAST[generep]<-paste0(months[as.numeric(startmonth)],", ",startday," ",startyear)
    
      sampledblastresults$Time.of.BLAST[generep]<-paste0(starthour,":",startminutes,":",startseconds)
    
      sampledblastresults$Annotation[generep]<-querydescriptives$annotation[generep]
    }
    
    generep<-generep+1
  }
  
  ##Completion time output - end
  
  timeended<-timelapseend.function(startyear=timebegin[[1]],startmonth=timebegin[[2]],startday=timebegin[[3]],starthour=timebegin[[4]],startminutes=timebegin[[5]],startseconds=timebegin[[6]],timestarted=timebegin[[7]],message.fin="FINISHED RAW BLASTS -")
  
  print(timeended[[2]])

  ##Adding annotations
  
  blastresults<-list(rawblastresults,sampledblastresults)

  names(blastresults)[1]<-rawblastresultsname
  
  names(blastresults)[2]<-sampledblastresultsname
  
  ##The NCBI BLAST program cannot parse annotations for the subject database, so the annotations will have to be added manually. The BLAST code can take a long time to complete, can be very resource intensive, and can cause crashes due to memory issues. For this reason the raw BLAST results will be saved before the annotations are added just in case, so if a crash does happen the BLAST output can be loaded and the annotations added after a fresh reboot.
  
  if(save.file==T){
        
    tempmess<-timelapseend.function(startyear=timebegin[[1]],startmonth=timebegin[[2]],startday=timebegin[[3]],starthour=timebegin[[4]],startminutes=timebegin[[5]],startseconds=timebegin[[6]],timestarted=timebegin[[7]],message.fin="Saving BLAST results without annotations...")

    print(tempmess[[2]])

    save(blastresults,file=paste0(directorysave,rdatablastname))
  }
  
  blastresults[[rawblastresultsname]]$Subject.Annotation<-"ZZZZZ"
  
  blastrep=1
  
  tempmess<-timelapseend.function(startyear=timebegin[[1]],startmonth=timebegin[[2]],startday=timebegin[[3]],starthour=timebegin[[4]],startminutes=timebegin[[5]],startseconds=timebegin[[6]],timestarted=timebegin[[7]],message.fin="Adding annotations in process...")

  print(tempmess[[2]])
  
  while(blastrep<nrow(blastresults[[rawblastresultsname]])+1){
    
    blastresults[[rawblastresultsname]]$Subject.Annotation[blastrep]<-unlist(getAnnot(subjectdatabase[blastresults[[rawblastresultsname]]$saccver[blastrep]]))
    
    blastrep=blastrep+1
  }
  
  ##This file rewrites the raw BLAST output and now contains contains the BLAST results with the annotations for each hit from the subject database.
  
  if(save.file==T){
        
    tempmess<-timelapseend.function(startyear=timebegin[[1]],startmonth=timebegin[[2]],startday=timebegin[[3]],starthour=timebegin[[4]],startminutes=timebegin[[5]],startseconds=timebegin[[6]],timestarted=timebegin[[7]],message.fin="Saving BLAST results with annotations...")

    print(tempmess[[2]])

    save(blastresults,file=paste0(directorysave,rdatablastname))
  }
  
  if(save.csv==T){

    write_csv(data.frame(blastresults[[rawblastresultsname]]),file = paste0(directorysave,blastfilename),col_names = T)
    
    write_csv(data.frame(blastresults[[sampledblastresultsname]]),file = paste0(directorysave,blastreportname),col_names = T)
  }
  
  ##Completion time output - end
  
  timeended<-timelapseend.function(startyear=timebegin[[1]],startmonth=timebegin[[2]],startday=timebegin[[3]],starthour=timebegin[[4]],startminutes=timebegin[[5]],startseconds=timebegin[[6]],timestarted=timebegin[[7]],message.fin=message.end)
  
  print(timeended[[1]])
  
  print(timeended[[2]])

  return(blastresults)
}

###Compiling all of the parts from BLASTS

###BLAST are computationally intensive and will require more RAM than my computer can handle to avoid crashes due to running out of memory. Due to memory and time constraints, the protein list will be split into 6 parts and then the blasts will be combined after each part has successfully completed. In order for this to work, all of the parts have to be saved with the following format: "pt" followed by the numerical order of the subset; so if the first subset of sequences are queried, then it should be saved as ".....pt1.RData". Also, each subset should be saved with the exact same file name except for the numerical subset value. "blastfiledirectory" should be a string for the directory containing the subsets of the BLAST results. "blastpartfilename" should be a string for the file name (requires extension ".RData") for each subset of the BLAST results that were saved; this won't work if the file names are different not counting the numerical subset. Since the BLAST function saves the BLAST output as a list with two data frames (raw BLAST results and the BLAST statistics), "sampledindex" is the list index containing the BLAST statistics and "rawblastindex" is the index containing the raw BLAST results; use the default for both indexes unless you manually changed the function. "querydescriptives" must be a data frame containing the names, annotations, and other important features of the query sequences; the BLAST function refers to specific column names in the "querydescriptives" data frame, so refer to the example BLAST function to know exactly how to format the column names or else this code won't work. "rawblastresultsname" should be a string for the name of the data frame containing the raw BLAST results. "sampledblastresultsname" should be a string for the name of the data frame that contain statistics regarding each query protein BLAST results. "save.file" gives the option (True or False) to save the BLAST output as an R file with the extension ".RData". If you choose to save the file using this code, then "rdatablastname" should be a string for the file name (requires extension ".RData") of the output and "directorysave" should be a string for the directory to save the file. "description" is an optional string that will be printed after the compilation has been completed. "message.start" is the message for the time elapse at the start of the function. "message.end" is the message for the time elapse after the function has completed.

compilationBLAST.function<-function(blastfiledirectory = outputdatabasedirectory, blastpartfilename = "Protein BLASTS List", sampledindex = 2, querydescriptives, rawblastindex = 1, rawblastresultsname = "rawblastresults", sampledblastresultsname = "sampledblastresults", save.file = F, directorysave = outputdatabasedirectory, rdatablastname = "Protein BLASTS List compilation",description = "", message.start = "STARTED -", message.end = "COMPLETED -"){
  
  ##Completion time output - start
  
  timebegin<-timelapsebegin.function(message.begin=message.start)
  
  print(timebegin[[7]])
  
  ##Compilation core code begins
  
  tempmess<-timelapseend.function(startyear=timebegin[[1]],startmonth=timebegin[[2]],startday=timebegin[[3]],starthour=timebegin[[4]],startminutes=timebegin[[5]],startseconds=timebegin[[6]],timestarted=timebegin[[7]],message.fin="Loading files...")

  print(tempmess[[2]])
  
  load(paste0(blastfiledirectory,blastpartfilename, " pt1.RData"))
  
  blastresults1<-blastresults
  
  load(paste0(blastfiledirectory,blastpartfilename, " pt2.RData"))
  
  blastresults2<-blastresults
  
  load(paste0(blastfiledirectory,blastpartfilename, " pt3.RData"))
  
  blastresults3<-blastresults
  
  load(paste0(blastfiledirectory,blastpartfilename, " pt4.RData"))
  
  blastresults4<-blastresults
  
  load(paste0(blastfiledirectory,blastpartfilename, " pt5.RData"))
  
  blastresults5<-blastresults
  
  load(paste0(blastfiledirectory,blastpartfilename, " pt6.RData"))
  
  blastresults6<-blastresults
  
  remove(blastresults)
  
  tempmess<-timelapseend.function(startyear=timebegin[[1]],startmonth=timebegin[[2]],startday=timebegin[[3]],starthour=timebegin[[4]],startminutes=timebegin[[5]],startseconds=timebegin[[6]],timestarted=timebegin[[7]],message.fin="Compiling files...")

  print(tempmess[[2]])
  
  comprawblastssampled<-rbind(blastresults1[[sampledindex]],blastresults2[[sampledindex]],blastresults3[[sampledindex]],blastresults4[[sampledindex]],blastresults5[[sampledindex]],blastresults6[[sampledindex]])
  
  compfullrawblasts<-rbind(blastresults1[[rawblastindex]],blastresults2[[rawblastindex]],blastresults3[[rawblastindex]],blastresults4[[rawblastindex]],blastresults5[[rawblastindex]],blastresults6[[rawblastindex]])
  
  remove(blastresults1,blastresults2,blastresults3,blastresults4,blastresults5,blastresults6)

  blastresultscomp<-list(compfullrawblasts,comprawblastssampled)
  
  names(blastresultscomp)[1]<-rawblastresultsname
  
  names(blastresultscomp)[2]<-sampledblastresultsname

  ##If the 6-part split of the query protein sequences is correct, then the concatenation of the 6 parts should have the exact same gene names and order as the original object that had all of the query names. If this is true, then the file will be saved. 
  
  querydescriptives<-querydescriptives[querydescriptives$gene.name%in%comprawblastssampled$Name,]
  
  if(identical(comprawblastssampled$Name,querydescriptives$gene.name)==T){
    
    print(paste0("SUCCESS!!! ",description," BLASTS COMPILATION ARE IN CORRECT ORDER!!!"))
  } else if(identical(comprawblastssampled$Name,querydescriptives$gene.name)!=T){
    
    base::warning(paste0("FAIL?? ",description," BLASTS COMPILATION ARE NOT IN ORDER??"))
  }
  
  if(save.file==T){
    
    tempmess<-timelapseend.function(startyear=timebegin[[1]],startmonth=timebegin[[2]],startday=timebegin[[3]],starthour=timebegin[[4]],startminutes=timebegin[[5]],startseconds=timebegin[[6]],timestarted=timebegin[[7]],message.fin="Saving...")

    print(tempmess[[2]])
    
    save(blastresultscomp,file=paste0(outputdatabasedirectory,rdatablastname))
  }
  
  ##Completion time output - end
  
  timeended<-timelapseend.function(startyear=timebegin[[1]],startmonth=timebegin[[2]],startday=timebegin[[3]],starthour=timebegin[[4]],startminutes=timebegin[[5]],startseconds=timebegin[[6]],timestarted=timebegin[[7]],message.fin=message.end)
  
  print(timeended[[1]])
  
  print(timeended[[2]])
  
  return(blastresultscomp)
}

###Adding species to BLAST results

###The default NCBI tool has no way of determining the species of the sequences from the subject database. This custom function will extract the species from the subject annotations accompanied with each sequence alignment. Also, for each query sequence, this function will further curate the BLAST results by selecting the alignment with the highest bitscore from each unique species from the subject database; this will eliminate duplicate BLAST hits that a query has for multiple proteins based on partial sequence similarities. "data" is the data frame containing the raw BLAST results (this is the compilation file if the query sequences were subset). "blastslist" should be the data frame containing the statistics from the BLAST results. "protein.database.analysis" gives the option (True or False) to determine unique species ONLY among the subject database NOT among the BLAST results. "query.species" is an optional string for the species that the query sequences originated from. "description" is an optional string that contains further details pertaining to the specific type of analysis being done. "save.file" gives the option (True or False) to save the BLAST output as an R file with the extension ".RData" as well as several csv files. If "protein.database.analysis" and "save.file" are both set to True, then "curatedblastsname" should be a string for the name of the csv file containing the parsed species from each entry of the subject database. "exportblastcsv" gives the option (True or False) to export the the BLAST results that are curated to only contain sequence alignments with the highest bitscore for each query BLAST; this is turned off by default because the file can be so large that it causes R to crash when trying to save. If "exportblastcsv" is set to True, then "uniquecuratedblastsname" should be a string for the name of the csv file that contains all BLASTS hits from unique organisms (regardless of bitscore) for each query. If "save.file" is set to True, then "countuniquesubjectname" should be a string for the name of the csv file detailing the number of BLAST hits that each unique species has among the sequence alignments; "countuniquequeryname" should be a string for the name of the csv file detailing the number of BLAST hits from unique species for each query gene. If "save.file" is set to True, then the curated BLAST results will be exported as a RData file consisting of a list of the following data.frames: "dataname" should be a string that will name the data frame consisting of all of the BLAST results initially input as "data" except that now, "dataname" has the parsed species for all of the BLAST alignments; "sampledblastresultsname" should be a string that will name the data frame containing the statistics of the BLAST results which is the same as what was input as "blastslist"; "uniqueorganismname" should be a string that will name the data frame that has the curated BLAST results with each query having a single alignment from each species with the highest bitscore. "print.index" gives the option (True or False) to print the current row of the particular data frame being analyzed. "rdatablastname" should be a string for the file name (requires extension ".RData") of the output and "directorysave" should be a string for the directory to save the file. "message.start" is the message for the time elapse at the start of the function. "message.end" is the message for the time elapse after the function has completed.

uniquespecies.function<-function(data, blastslist, protein.database.analysis = F, query.species = "", description = "", save.file = F, curatedblastsname = "RAW DATA - Every Entry in Subject Database.csv", exportblastcsv = F, uniquecuratedblastsname = "CURATED - Every Entry.csv", countuniquesubjectname = "Every UNIQUE Organism.csv", countuniquequeryname = "Unique Organisms Hits For Each Query.csv", dataname = "comp.rawblastresults", sampledblastresultsname = "comp.sampledblastresults", uniqueorganismname = "uniqueorganisms", print.index = T , rdatablastname = "Protein BLASTS List compilation.RData", directorysave = outputdatabasedirectory, message.start = "STARTED -", message.end = "COMPLETED -"){
  
  speciesoutput<-rep(NA,nrow(data))
  
  generep<-1
  
  ##Completion time output - start
  
  timebegin<-timelapsebegin.function(message.begin=message.start)
  
  print(timebegin[[7]])
  
  tempmess<-timelapseend.function(startyear=timebegin[[1]],startmonth=timebegin[[2]],startday=timebegin[[3]],starthour=timebegin[[4]],startminutes=timebegin[[5]],startseconds=timebegin[[6]],timestarted=timebegin[[7]],message.fin="Parsing species name for each BLAST hit...")
  
  print(tempmess[[2]])
  
  ##Unique species core code begins
  
  while(generep<(nrow(data)+1)){
    
    if(print.index==T){
      
      print(generep)
    }
  
    if(grepl("[[",data$Subject.Annotation[generep],fixed = T)==T&grepl("pir||",data$Subject.Annotation[generep],fixed = T)!=T&grepl("sp]",data$Subject.Annotation[generep],fixed = T)!=T){
      
      tempannotfirst<-unlist(strsplit(data$Subject.Annotation[generep],"[[",fixed=T))[2]
  
      tempannotsecond<-unlist(strsplit(tempannotfirst,"]",fixed=T))
      
      speciesoutput[generep]<-c2s(c(unlist(strsplit(tempannotsecond," ",fixed=T))[1]," ",unlist(strsplit(tempannotsecond," ",fixed=T))[3]))
    } else if(grepl("[[",data$Subject.Annotation[generep],fixed = T)!=T&grepl("pir||",data$Subject.Annotation[generep],fixed = T)!=T&grepl("sp]",data$Subject.Annotation[generep],fixed = T)!=T){
    
        if(length(unlist(strsplit(c2s(data$Subject.Annotation[generep]),"[",fixed=T)))<3){
        
          tempannotfirst<-unlist(strsplit(data$Subject.Annotation[generep],"[",fixed=T))[2]
        
          tempannotsecond<-unlist(strsplit(tempannotfirst,"]",fixed=T))[1]
  
        speciesoutput[generep]<-c2s(c(unlist(strsplit(tempannotsecond," ",fixed=T))[1]," ",unlist(strsplit(tempannotsecond," ",fixed=T))[2]))
        
          if(grepl(",",speciesoutput[generep],fixed = T)==T&grepl("=",speciesoutput[generep],fixed = T)!=T){
              
            speciesoutput[generep]<-unlist(strsplit(speciesoutput[generep],",",fixed=T))[1]
          } else if(grepl(",",speciesoutput[generep],fixed = T)==T&grepl("=",speciesoutput[generep],fixed = T)==T){
            
            speciesoutput[generep]<-unlist(strsplit(speciesoutput[generep],"=",fixed=T))[1]
          } else if(grepl(",",speciesoutput[generep],fixed = T)!=T){
            
            speciesoutput[generep]<-speciesoutput[generep]
          }
          
        } else if(length(unlist(strsplit(c2s(data$Subject.Annotation[generep]),"[",fixed=T)))>2){
    
          tempannotfirst<-unlist(strsplit(data$Subject.Annotation[generep],"[",fixed=T))
          
          tempannotsecond<-unlist(strsplit(tempannotfirst[length(tempannotfirst)],"]",fixed=T))
    
          speciesoutput[generep]<-c2s(c(unlist(strsplit(tempannotsecond," ",fixed=T))[1]," ",unlist(strsplit(tempannotsecond," ",fixed=T))[2]))
          
            if(grepl(",",speciesoutput[generep],fixed = T)==T&grepl("=",speciesoutput[generep],fixed = T)!=T){
                
              speciesoutput[generep]<-unlist(strsplit(speciesoutput[generep],",",fixed=T))[1]
            } else if(grepl(",",speciesoutput[generep],fixed = T)==T&grepl("=",speciesoutput[generep],fixed = T)==T){
              speciesoutput[generep]<-unlist(strsplit(speciesoutput[generep],"=",fixed=T))[1]
            } else if(grepl(",",speciesoutput[generep],fixed = T)!=T){
              
              speciesoutput[generep]<-speciesoutput[generep]
            } 
        }
      
    } else if (grepl("pir||",data$Subject.Annotation[generep],fixed = T)==T&grepl("sp]",data$Subject.Annotation[generep],fixed = T)!=T){
  
      tempannotfirst<-unlist(strsplit(data$Subject.Annotation[generep],"yeast ",fixed=T))[2]
      
      tempannotsecond<-unlist(strsplit(tempannotfirst,"(",fixed=T))[2]
      
      tempannothird<-unlist(strsplit(tempannotsecond,")",fixed=T))[1]
      
      speciesoutput[generep]<-c2s(c(unlist(strsplit(tempannothird," ",fixed=T))[1]," ",unlist(strsplit(tempannothird," ",fixed=T))[2]))
    } else if (grepl("sp]",data$Subject.Annotation[generep],fixed = T)==T){
      
      tempannotfirst<-unlist(strsplit(data$Subject.Annotation[generep],"[",fixed=T))
      
      tempannotsecond<-c2s(c(s2c(unlist(strsplit(tempannotfirst[length(tempannotfirst)],"]",fixed=T))),"."))
  
      speciesoutput[generep]<-c2s(c(unlist(strsplit(tempannotsecond," ",fixed=T))[1]," ",unlist(strsplit(tempannotsecond," ",fixed=T))[2]))
    }
    
    generep<-generep+1
  }
  
  data$Organism<-speciesoutput
  
  data1<-data
  
  data$Organism[grepl("NA NA",data$Organism,fixed = T)==T]<-"Candida albicans"
  
  ##Index 217440 belongs to Cyberlindnera jadinii https://www.ncbi.nlm.nih.gov/protein/CEP21445.1
  
  data$Organism[grepl("Cyber NA",data$Organism,fixed = T)==T]<-"Cyberlindnera jadinii"
  
  data$Organism[grepl("Starmerella cf.",data$Organism,fixed = T)==T]<-"Starmerella sorbosivorans"
  
  ##Hansenula is an obsolete genus and every Hansenula species has been renamed/reclassified.
  
  data$Organism[grepl("Hansenula polymorpha",data$Organism,fixed = T)==T]<-"Ogataea polymorpha"
  
  data$Organism[grepl("Pichia angusta",data$Organism,fixed = T)==T]<-"Ogataea polymorpha"
  
  data$Organism[grepl("Hansenula saturnus",data$Organism,fixed = T)==T]<-"Cyberlindnera saturnus"
  
  data$Organism[grepl("Williopsis saturnus",data$Organism,fixed = T)==T]<-"Cyberlindnera saturnus"
  
  ##Dekkera bruxellensis and Brettanomyces bruxellensis are the same fungi at different growth stages, so they will be treated as the same species.
  
  data$Organism[grepl("Dekkera bruxellensis",data$Organism,fixed = T)==T]<-"Brettanomyces bruxellensis"
  
  data<-data[!grepl("uncultured",data$Organism,fixed = T),]
  
  ##The Metschnikowia pulcherrima subclade includes yeast affiliated, yet distinct from existing Metschnikowia, but not enough taxonomic evidence has been provided to classify them as their own species. As such, all entries with Metschnikowia aff. will be omitted.
  
  data<-data[!grepl("aff.",data$Organism,fixed = T),]
  
  data<-data[!grepl("NA",data$Organism,fixed = T),]
  
  ##Some entries only have the genus but not the name of the species, such as [Candida sp.]. However, only unique species will be considered, so all entries that don't specify species will be omitted.
  
  data<-data[!grepl("sp.",data$Organism,fixed = T),]
  
  data<-data[!grepl("yeast",data$Organism,fixed = T),]
  
  data<-data[!grepl("synthetic construct",data$Organism,fixed = T),]
  
  if(protein.database.analysis!=T){
  
    data$qaccver<-factor(data$qaccver)
  
    finaldataset<-rep(list(NA),length(levels(data$qaccver)))
  
    tempmess<-timelapseend.function(startyear=timebegin[[1]],startmonth=timebegin[[2]],startday=timebegin[[3]],starthour=timebegin[[4]],startminutes=timebegin[[5]],startseconds=timebegin[[6]],timestarted=timebegin[[7]],message.fin="Parsing unique species for each query...")
    
    print(tempmess[[2]])
  
    generep<-1
  
    while(generep<(length(levels(data$qaccver))+1)){
      
      if(print.index==T){
        
        print(generep)
      }
      
      finaldatasettemp<-data[data$qaccver==levels(data$qaccver)[generep],]
      
      finaldatasettemp<-finaldatasettemp[order(finaldatasettemp$bitscore,decreasing = T),]
      
      finaldataset[generep]<-list(finaldatasettemp[!duplicated(finaldatasettemp$Organism),])
      
      names(finaldataset)[generep]<-levels(data$qaccver)[generep]
    
      generep<-generep+1
    }
  
  compunique<-rbind.fill(finaldataset)
  } else if (protein.database.analysis==T){
    
    compunique<-data
  }
  
  finalunique<-data[!duplicated(data$Organism),]
  
  finaluniquesubject<-finalunique
  
  finaluniquesubject$`Number of Entries`<-99999
  
  tempmess<-timelapseend.function(startyear=timebegin[[1]],startmonth=timebegin[[2]],startday=timebegin[[3]],starthour=timebegin[[4]],startminutes=timebegin[[5]],startseconds=timebegin[[6]],timestarted=timebegin[[7]],message.fin="Determining how many BLAST hits each query had in dataset...")
  
  print(tempmess[[2]])  
  
  generep<-1
  
  while(generep<nrow(finaluniquesubject)+1){
    
    if(print.index==T){
      
      print(generep)
    }
    
    finaluniquesubject$`Number of Entries`[generep]<-nrow(compunique[compunique$Organism==finaluniquesubject$Organism[generep],])
    
    generep<-generep+1
  }
  
  finaluniquesubject$Analysis<-description
  
  finaluniquesubject<-finaluniquesubject[,c("Organism","Number of Entries","Analysis")]

  tempmess<-timelapseend.function(startyear=timebegin[[1]],startmonth=timebegin[[2]],startday=timebegin[[3]],starthour=timebegin[[4]],startminutes=timebegin[[5]],startseconds=timebegin[[6]],timestarted=timebegin[[7]],message.fin="Determining how many BLAST hits there are for every unique species in dataset...")
  
  print(tempmess[[2]])  
  
  if(protein.database.analysis!=T){
    
    finaluniquequery<-data[!duplicated(data$qaccver),]
    
    finaluniquequery$`Number of Unique Organisms`<-99999
    
    finaluniquequery$qaccver<-as.factor(finaluniquequery$qaccver)
    
    generep<-1
    
    while(generep<nrow(finaluniquequery)+1){
      
      if(print.index==T){
      
        print(generep)
      }
      
      finaluniquequery$`Number of Unique Organisms`[generep]<-nrow(compunique[compunique$qaccver==finaluniquequery$qaccver[generep],])
      
      generep<-generep+1
    }
    
    finaluniquequery$Organism<-query.species
  
    finaluniquequery$Analysis<-description
  
    finaluniquequery<-finaluniquequery[,c("qaccver","Number of Unique Organisms","Analysis")]
    
    names(finaluniquequery)[names(finaluniquequery) == "qaccver"] <- "Query.Name"
    
    curatedBLASTSresults<-list(blastslist,data1,compunique,finaluniquequery,finaluniquesubject)
    
    names(curatedBLASTSresults)[1]<-sampledblastresultsname
    
    names(curatedBLASTSresults)[2]<-dataname
    
    names(curatedBLASTSresults)[3]<-uniqueorganismname
    
    names(curatedBLASTSresults)[4]<-"queryblasthits.uniqueorganisms"
    
    names(curatedBLASTSresults)[5]<-"count.uniqueorganisms"
    
    if(save.file==T){
      
      tempmess<-timelapsebegin.function(message.begin="Saving files...")

      tempmess<-timelapseend.function(startyear=timebegin[[1]],startmonth=timebegin[[2]],startday=timebegin[[3]],starthour=timebegin[[4]],startminutes=timebegin[[5]],startseconds=timebegin[[6]],timestarted=timebegin[[7]],message.fin="Saving files...")
  
      print(tempmess[[2]]) 
      
      ##finaluniquequery contains the number of BLAST hits from unique organisms for each query gene
      
      save(curatedBLASTSresults,file=paste0(directorysave,rdatablastname))
    
      write_csv(finaluniquequery,file = paste0(directorysave,countuniquequeryname),col_names = T)
    }
    
  } else if(protein.database.analysis==T&save.file==T){
      
    ##data1 contains the parsed organism species from each entry of the subject database. The file will be too long and big to open in excel, but it can be saved as a csv file and uploaded into R.
      
    write_csv(data1,file = paste0(directorysave,curatedblastsname),col_names = T)
  }

  if(save.file==T){
    
    ##finaluniquesubject reveals the number of BLAST hits that each unique species has.
    
    write_csv(finaluniquesubject,file = paste0(directorysave,countuniquesubjectname),col_names = T)
  }
  
  if(exportblastcsv==T){
          
    tempmess<-timelapseend.function(startyear=timebegin[[1]],startmonth=timebegin[[2]],startday=timebegin[[3]],starthour=timebegin[[4]],startminutes=timebegin[[5]],startseconds=timebegin[[6]],timestarted=timebegin[[7]],message.fin="Exporting all BLAST results...")

    print(tempmess[[2]]) 
      
    ##compunique contains all BLAST hits from unique organisms (regardless of bitscore) for each query. This is turned off by default because the file can be so large that it causes R to crash when trying to save.
    
    write_csv(compunique,file = paste0(directorysave,uniquecuratedblastsname),col_names = T)
  }
  
  ##Completion time output - end
  
  timeended<-timelapseend.function(startyear=timebegin[[1]],startmonth=timebegin[[2]],startday=timebegin[[3]],starthour=timebegin[[4]],startminutes=timebegin[[5]],startseconds=timebegin[[6]],timestarted=timebegin[[7]],message.fin=message.end)
  
  print(timeended[[1]])
  
  print(timeended[[2]])
  
  print(paste0("There are ",nrow(finaluniquesubject)," Unique Organisms in ",description))

  if(protein.database.analysis!=T){
        
    return(curatedBLASTSresults)
  }
}

###Protein conservation scores analysis and output

###Protein conservation scores are calculated using a weighted algorithm."curatedblastsdata" should be the data frame that has the curated BLAST results with each query having a single alignment from each species with the highest bitscore. "sampleddata" should be a data frame that has the statistics from the BLAST results. "querystart" should be a string ("Beginning", "Middle", or "End") identifying the region to start the protein conservation score calculations; default is "Beginning" which is the start of the query proteins. "homologystart" should be the region that the BLAST algorithm determines where the alignment begins in the query or subject sequences; the default is "qstart" which is a numerical value that indicates the start of the alignment within the query. "analysis" is an optional string that refers to the length of amino acids spanning the conservation score as well as the region (beginning, middle or end). "query.species" is an optional string for the species that the query sequences originated from. "querydescriptives" must be a data frame containing the names, annotations, and other important features of the query sequences; the BLAST function refers to specific column names in the "querydescriptives" data frame, so refer to the example BLAST function to know exactly how to format the column names or else this code won't work. "bitscore.threshold" is the minimum bitscore value from each sequence alignment that will be included in the protein score calculations. "rampzonelength" is the length of amino acids used as the basis for homology, and is the length for the range of values in the applied as a weight to the conservation scores; for example, a "rampzonelength" of 40 means that the code will count how many unique species have a qstart (or sstart, or qend, etc.) at every position from 1-40, with the first amino acid having the greatest weight of 40, and the last (fortieth) amino acid having the smallest weight of 1. "print.index" gives the option (True or False) to print the current numerical index that is being analyzed; for example, an index of 5 means that the fifth protein in the "querydescriptives" data frame is currently being processed by the code. "description" is an optional string that contains further details pertaining to the specific type of analysis being done. "message.start" is the message for the time elapse at the start of the function. "message.end" is the message for the time elapse after the function has completed.

weightedproportion.function<-function(curatedblastsdata, sampleddata, querystart = "Beginning", homologystart = "qstart", analysis = "", query.species = "", querydescriptives, bitscore.threshold = 50, rampzonelength = 40, print.index = T, description = "", message.start = "STARTED -", message.end = "COMPLETED -"){

  ##Completion time output - start
  
  timebegin<-timelapsebegin.function(message.begin=message.start)
  
  print(timebegin[[7]])
  
  ##Code for protein conservation scores begins
  
  curatedblasts<-curatedblastsdata[!curatedblastsdata$bitscore<bitscore.threshold,]
  
  names(curatedblasts)[names(curatedblasts) == homologystart] <- "homology"

  generep<-1
  
  geneconservation<-data.frame(Name="ZZZZZ",Common.Name="ZZZZZ",Amino.Acids=99999,Unique.Organisms=99999,Unique.Organisms.Bitscore.Threshold=99999,BLASTS.Rampzone.window.start=99999,BLASTS.Rampzone.window.finish=99999,BLASTS.Rampzone.match=99999,Lowest.match.Anywhere=99999,Highest.match.Anywhere=99999,Lowest.match.Rampzone=99999,Highest.match.Rampzone=99999,Sum.Weighted.Proportion=99999,Details="ZZZZZ",Annotation="ZZZZZ",Number=1:nrow(querydescriptives))
  
  while(generep<nrow(querydescriptives)+1){
    
    if(print.index==T){
      
      print(generep)
    }
    
    proteinname<-querydescriptives$gene.name[generep]
    
    tempblast<-curatedblasts[curatedblasts$qaccver==proteinname,]
    
    if (querystart=="Beginning"){
      
      tempblastsrampzone<-tempblast[!tempblast$homology>rampzonelength,]
      
      blastrep<-1
      
      blastrep.start<-1
    } else if (querystart=="Middle"){
      
      tempblastsrampzone<-tempblast[!tempblast$homology>(round(querydescriptives$Amino.Acids[querydescriptives$gene.name==proteinname]/2)+(rampzonelength/2)),]
      
      tempblastsrampzone<-tempblastsrampzone[!tempblastsrampzone$homology<((round(querydescriptives$Amino.Acids[querydescriptives$gene.name==proteinname]/2)-(rampzonelength/2))+1),]
      
      blastrep<-((round(querydescriptives$Amino.Acids[querydescriptives$gene.name==proteinname]/2)-(rampzonelength/2))+1)
      
      blastrep.start<-((round(querydescriptives$Amino.Acids[querydescriptives$gene.name==proteinname]/2)-(rampzonelength/2))+1)
    } else if (querystart=="End"){
      
      tempblastsrampzone<-tempblast[!tempblast$homology>(querydescriptives$Amino.Acids[querydescriptives$gene.name==proteinname]),]
      
      tempblastsrampzone<-tempblastsrampzone[!tempblastsrampzone$homology<(querydescriptives$Amino.Acids[querydescriptives$gene.name==proteinname])-(rampzonelength)+1,]
      
      blastrep<-(querydescriptives$Amino.Acids[querydescriptives$gene.name==proteinname])-(rampzonelength)+1
      
      blastrep.start<-(querydescriptives$Amino.Acids[querydescriptives$gene.name==proteinname])-(rampzonelength)+1
    }    

    comp.homologycount=NULL
    
    if(nrow(tempblastsrampzone)!=0){
        
      while(blastrep<max(tempblastsrampzone$homology)+1){
        
        homologycount<-data.frame(matchposition=blastrep,count=nrow(tempblastsrampzone[tempblastsrampzone$homology==blastrep,]))
        
        comp.homologycount=rbind(comp.homologycount,homologycount)
    
        blastrep=blastrep+1
      }
      
      comp.homologycount$proportion<-comp.homologycount$count/nrow(tempblast)
      
        if (querystart=="End"){
          
          comp.homologycount$weight<-(abs(nrow(comp.homologycount)-rampzonelength)+1):rampzonelength
        } else if (querystart!="End"){
          
          comp.homologycount$weight<-rampzonelength:(abs(nrow(comp.homologycount)-rampzonelength)+1)
        }

      comp.homologycount$weightXproportion<-comp.homologycount$proportion*comp.homologycount$weight
         
      geneconservation$Name[generep]<-proteinname
      
      geneconservation$Common.Name[generep]<-querydescriptives$common.name[querydescriptives$gene.name==proteinname]
      
      geneconservation$Amino.Acids[generep]<-querydescriptives$Amino.Acids[querydescriptives$gene.name==proteinname]
      
      geneconservation$Unique.Organisms[generep]<-nrow(curatedblastsdata[curatedblastsdata$qaccver==proteinname,])
      
      geneconservation$Sum.Weighted.Proportion[generep]<-sum(comp.homologycount$weightXproportion)
      
      geneconservation$Unique.Organisms.Bitscore.Threshold[generep]<-nrow(tempblast)
      
      geneconservation$BLASTS.Rampzone.window.start[generep]<-blastrep.start
      
      geneconservation$BLASTS.Rampzone.window.finish[generep]<-blastrep-1
      
      geneconservation$BLASTS.Rampzone.match[generep]<-nrow(tempblastsrampzone)
      
      geneconservation$Lowest.match.Anywhere[generep]<-min(tempblast$homology)
      
      geneconservation$Highest.match.Anywhere[generep]<-max(tempblast$homology)
      
      geneconservation$Lowest.match.Rampzone[generep]<-min(tempblastsrampzone$homology)
      
      geneconservation$Highest.match.Rampzone[generep]<-max(tempblastsrampzone$homology)
      
      geneconservation$Details[generep]<-"Ok"
      
      geneconservation$Annotation[generep]<-querydescriptives$annotation[querydescriptives$gene.name==proteinname]
    } else if((nrow(tempblastsrampzone)==0)&(nrow(tempblast)!=0)){

      geneconservation$Name[generep]<-proteinname
      
      geneconservation$Common.Name[generep]<-querydescriptives$common.name[querydescriptives$gene.name==proteinname]
      
      geneconservation$Amino.Acids[generep]<-querydescriptives$Amino.Acids[querydescriptives$gene.name==proteinname]
      
      geneconservation$Unique.Organisms[generep]<-nrow(curatedblastsdata[curatedblastsdata$qaccver==proteinname,])
      
      geneconservation$Sum.Weighted.Proportion[generep]<-sum(comp.homologycount$weightXproportion)
      
      geneconservation$Unique.Organisms.Bitscore.Threshold[generep]<-nrow(tempblast)
      
      geneconservation$BLASTS.Rampzone.window.start[generep]<-blastrep.start
      
      geneconservation$BLASTS.Rampzone.window.finish[generep]<-0
      
      geneconservation$BLASTS.Rampzone.match[generep]<-nrow(tempblastsrampzone)
      
      geneconservation$Lowest.match.Anywhere[generep]<-min(tempblast$homology)
      
      geneconservation$Highest.match.Anywhere[generep]<-max(tempblast$homology)
      
      geneconservation$Lowest.match.Rampzone[generep]<-0
      
      geneconservation$Highest.match.Rampzone[generep]<-0
      
      geneconservation$Details[generep]<-paste0("No BLASTs with match among ",analysis)
      
      geneconservation$Annotation[generep]<-querydescriptives$annotation[querydescriptives$gene.name==proteinname]
    } else if((nrow(tempblastsrampzone)==0)&(nrow(tempblast)==0)&(nrow(curatedblastsdata[curatedblastsdata$qaccver==proteinname,]))!=0){
          
      geneconservation$Name[generep]<-proteinname
      
      geneconservation$Common.Name[generep]<-querydescriptives$common.name[querydescriptives$gene.name==proteinname]
      
      geneconservation$Amino.Acids[generep]<-querydescriptives$Amino.Acids[querydescriptives$gene.name==proteinname]
      
      geneconservation$Unique.Organisms[generep]<-nrow(curatedblastsdata[curatedblastsdata$qaccver==proteinname,])
      
      geneconservation$Sum.Weighted.Proportion[generep]<-sum(comp.homologycount$weightXproportion)
      
      geneconservation$Unique.Organisms.Bitscore.Threshold[generep]<-nrow(tempblast)
      
      geneconservation$BLASTS.Rampzone.window.start[generep]<-blastrep.start
      
      geneconservation$BLASTS.Rampzone.window.finish[generep]<-0
      
      geneconservation$BLASTS.Rampzone.match[generep]<-nrow(tempblastsrampzone)
      
      geneconservation$Lowest.match.Anywhere[generep]<-0
      
      geneconservation$Highest.match.Anywhere[generep]<-0
      
      geneconservation$Lowest.match.Rampzone[generep]<-0
      
      geneconservation$Highest.match.Rampzone[generep]<-0
      
      geneconservation$Details[generep]<-paste0("All BLASTs have bitscore lower than ",bitscore.threshold)
      
      geneconservation$Annotation[generep]<-querydescriptives$annotation[querydescriptives$gene.name==proteinname]
    } else if((nrow(curatedblastsdata[curatedblastsdata$qaccver==proteinname,]))==0){
          
      geneconservation$Name[generep]<-proteinname
      
      geneconservation$Common.Name[generep]<-querydescriptives$common.name[querydescriptives$gene.name==proteinname]
      
      geneconservation$Amino.Acids[generep]<-querydescriptives$Amino.Acids[querydescriptives$gene.name==proteinname]
      
      geneconservation$Unique.Organisms[generep]<-0
      
      geneconservation$Sum.Weighted.Proportion[generep]<-0
      
      geneconservation$Unique.Organisms.Bitscore.Threshold[generep]<-0
      
      geneconservation$BLASTS.Rampzone.window.start[generep]<-0
      
      geneconservation$BLASTS.Rampzone.window.finish[generep]<-0
      
      geneconservation$BLASTS.Rampzone.match[generep]<-0
      
      geneconservation$Lowest.match.Anywhere[generep]<-0
      
      geneconservation$Highest.match.Anywhere[generep]<-0
      
      geneconservation$Lowest.match.Rampzone[generep]<-0
      
      geneconservation$Highest.match.Rampzone[generep]<-0
      
      geneconservation$Details[generep]<-"No BLAST Results"
      
      geneconservation$Annotation[generep]<-querydescriptives$annotation[querydescriptives$gene.name==proteinname]
    }
  
    generep<-generep+1 
  }
  
  geneconservation<-join(geneconservation,sampleddata[,c("Name","Total.Hits","Annotation")],by=c("Name","Annotation"),type="full", match="all")
  
  geneconservation$Query.Span<-description
  
  geneconservation$Query.Species<-query.species

  names(geneconservation)[names(geneconservation) == "Total.Hits"] <- "Total.BLASTS"
  
  geneconservation[is.na(geneconservation)] <- 0

  geneconservation$Bitscore.Threshold<-bitscore.threshold
  
  geneconservation$Analysis<-analysis
  
  geneconservation$Rampzone.Start<-querystart
  
  geneconservation<-geneconservation[,c("Name","Common.Name","Amino.Acids","Total.BLASTS","Unique.Organisms","Unique.Organisms.Bitscore.Threshold","BLASTS.Rampzone.window.start","BLASTS.Rampzone.window.finish","BLASTS.Rampzone.match","Lowest.match.Anywhere","Highest.match.Anywhere","Lowest.match.Rampzone","Highest.match.Rampzone","Sum.Weighted.Proportion","Rampzone.Start","Analysis","Bitscore.Threshold","Details","Query.Species","Query.Span","Annotation")]
  
  names(geneconservation)[names(geneconservation) == "Annotation"] <- "Query.Annotation"

  names(geneconservation)[names(geneconservation) == "BLASTS.Rampzone.match"] <- paste0("BLASTS.Rampzone.",homologystart)
  
  names(geneconservation)[names(geneconservation) == "Lowest.match.Anywhere"] <- paste0("Lowest.",homologystart,".Anywhere")
  
  names(geneconservation)[names(geneconservation) == "Highest.match.Anywhere"] <- paste0("Highest.",homologystart,".Anywhere")
  
  names(geneconservation)[names(geneconservation) == "Lowest.match.Rampzone"] <- paste0("Lowest.",homologystart,".Rampzone")
  
  names(geneconservation)[names(geneconservation) == "Highest.match.Rampzone"] <- paste0("Highest.",homologystart,".Rampzone")
  
  geneconservation<-geneconservation[order(geneconservation$Name,decreasing = F),]

  ##Completion time output - end
  
  timeended<-timelapseend.function(startyear=timebegin[[1]],startmonth=timebegin[[2]],startday=timebegin[[3]],starthour=timebegin[[4]],startminutes=timebegin[[5]],startseconds=timebegin[[6]],timestarted=timebegin[[7]],message.fin=message.end)
  
  print(timeended[[1]])
  
  print(timeended[[2]])
  
  return(geneconservation)
}

```

####PART 1: INITIAL TRANSLATION SPEED CALCULATIONS

###Generating codon usage table.

```{r,echo=F,eval=F}

###Gene diagnostics to detect pseudogenes and dubious ORF from fasta file
### The file contains all Saccharomyces cerevisiae protein-coding genes, and can be located using the url  http://sgd-archive.yeastgenome.org/sequence/S288C_reference/orf_dna/ This particular fasta file is orf_coding, which  is NOT supposed to have any ORF derived from dubious genes or pseudogenes. However, I discovered that in fact, this file has 12 pseudogenes that will mess up your analyses. To fix this, I am subsetting the original fasta file to exclude all genes annotated with pseudogenes (designated by the ", pseudogene," string), as well as dubious genes (designated by the ", Dubious ORF," string). The workable object, with 6022 genes, now has ORF that have been verified to produce protein products.

yeastgenes<-read.fasta(paste0(inputdatabasedirectory,"orf_coding_R64-3-1_20210421.fasta"))

###Gene diagnostics to detect pseudogenes and dubious ORF from fasta file. "query.genes" is the list of genes as formatted from the read.fasta() function. "query.species" is an optional string for the species that the sequences originated from. "pseudogene.fasta.string" is an optional identifier in the fasta annotations that denote genes known to be pseudogenes. "dubiousORF.fasta.string" is the optional identifier in annotations that denote known dubious ORF. "fasta.updated" is an optional string for the last known date that the fasta file was uploaded to the queried website. "file.url" is the optional URL string to access the the fasta file. "notes" is an optional string for any purpose to accompany the output. "print.index" gives the option (True or False) to print the current numerical index in the fasta file that is being analyzed; for example, an index of 5 means that the fifth gene listed in the fasta file is currently being processed by the code. "message.start" is the message for the time elapse at the start of the function. "message.end" is the message for the time elapse after the function has completed.

checkyeast.output<-checkORFfasta.function(query.genes=yeastgenes,query.species="Saccharomyces cerevisiae",pseudogene.fasta.string=", pseudogene,",dubiousORF.fasta.string=", Dubious ORF,",fasta.updated ="April 22, 2021",file.url = "http://sgd-archive.yeastgenome.org/sequence/S288C_reference/orf_dna/",notes="", print.index=T,message.start="Started S. cerevisiae genes diagnostics -",message.end="Finished S. cerevisiae genes diagnostics -")

yeastgenes.diagnostics<-checkyeast.output$pseudogenes.check

yeast.annotated.psuedogenes<-checkyeast.output$annotated.pseudogenes

yeast.annotated.dubiousORF<-checkyeast.output$annotated.dubiousORF

gene<-checkyeast.output$curated.genes

###Generating codon usage table

yeast.rrt.values <- read_excel(paste0(inputdatabasedirectory,"Supplemental Table 1, RRT values.xlsx"), 
    col_names = T)

###RRT is the average relative frequency of a codon that a ribosome occupies over 10 positions, and therefore each position is expected to have a frequency of 0.1 (see Figure 3 in the Gardin et al. paper). In the supplementary Table, these RRT values are specifically for position 6 which is believed to be the A-site in Saccharomyces cerevisiae. Therefore, all RRT values have to be multiplied by 10 to reflect the RRT that is specific to the ribosome occupancy at the A-site.

yeast.rrt.values$RRT<-(yeast.rrt.values$`RRT (Ribosome Residence Time)`)*10

names(yeast.rrt.values)[1] <- "Codons"

yeast.rrt.values<-yeast.rrt.values[,c("Codons","RRT")]

###Generating codon usage table. "query.genes" is the list of genes as formatted from the read.fasta() function. "query.species" is an optional string for the species that the sequences originated from."rrt.values" gives the option to assign a codon with its RRT value or codon-specific translation speed; this must be a data frame with 2 columns, and the column with the codons must have "Codons" as the column name. "fasta.updated" is an optional string for the last known date that the fasta file was uploaded to the queried website. "file.url" is the optional URL string in which the fasta can be accessed. "notes" is an optional string for any purpose to accompany the output. "save.file" gives the option (True or False) to save and export the results as an excel file directly into an assigned directory. If you choose to save the file using this code, then "outputfilename" is the string of the file name (default extension will be ".xlsx") for the output and "directorysave" should be a string for the directory to save the file. "print.index" gives the option (True or False) to print the current numerical index in the fasta file that is being analyzed; for example, an index of 5 means that the fifth gene listed in the fasta file is currently being processed by the code. "message.start" is the message for the time elapse at the start of the function. "message.end" is the message for the time elapse after the function has completed.

yeastcodonusage<-codonusagetable.function(query.genes=gene,query.species="Saccharomyces cerevisiae",rrt.values=yeast.rrt.values,fasta.updated ="April 22, 2021",file.url = "http://sgd-archive.yeastgenome.org/sequence/S288C_reference/orf_dna/",notes="",save.file = T,outputfilename="Saccharomyces cerevisiae codon usage table.xlsx",directorysave = outputdatabasedirectory,print.index=T,message.start="Started - S. cerevisiae codon usage -",message.end="Finished S. cerevisiae codon usage -")

```

###Generating translation speed at the N-termini and C-termini.

```{r,echo=F,eval=F}

yeastgenes<-read.fasta(paste0(inputdatabasedirectory,"orf_coding_R64-3-1_20210421.fasta"))

checkyeast.output<-checkORFfasta.function(query.genes=yeastgenes,query.species="Saccharomyces cerevisiae",pseudogene.fasta.string=", pseudogene,",dubiousORF.fasta.string=", Dubious ORF,",fasta.updated ="April 22, 2021",file.url = "http://sgd-archive.yeastgenome.org/sequence/S288C_reference/orf_dna/",notes="", print.index=T,message.start="Started S. cerevisiae genes diagnostics -",message.end="Finished S. cerevisiae genes diagnostics -")

gene<-checkyeast.output$curated.genes

yeastcodonusage<-data.frame(read_excel(paste0(outputdatabasedirectory,"Saccharomyces cerevisiae codon usage table.xlsx"),col_names = T),check.names = F,stringsAsFactors = F)

###Generating translation speed at the N-termini and C-termini. "query.genes" is the list of genes as formatted from the read.fasta() function. "query.species" is an optional string for the species that the sequences originated from. "rampzonelength.fiveprime" indicates the codon length of the window to measure translation speed at the N-termini. "rampzonelength.threeprime" indicates the codon length of the window to measure translation speed at the C-termini. "codons.translationspeed" is the data frame of codons with corresponding translation speeds, which must be formatted as a dataframe; all codons in the data frame must have a numerical codon-specific translation speed, the column with the codons must have "Codons" as the column name, and the column with the codon-specific translation speeds must have "RRT" as the column name. "save.file" gives the option (True or False) to save the translation speed output as an excel file with 3 tabs (default extension will be ".xlsx"). If you choose to save the file using this code, then "outputfilename" is the string of the file name for the output and "directorysave" should be a string for the directory to save the file. "print.index" gives the option (True or False) to print the current numerical index in the fasta file that is being analyzed; for example, an index of 5 means that the fifth gene listed in the fasta file is currently being processed by the code. "message.start" is the message for the time elapse at the start of the function. "message.end" is the message for the time elapse after the function has completed.

wtatg30<-translationspeed.function(query.genes=gene,query.species="Saccharomyces cerevisiae",rampzonelength.fiveprime = 30,rampzonelength.threeprime = 30,codons.translationspeed = yeastcodonusage, save.file = T,outputfilename ="Initial Translation Speed Table 30.xlsx",directorysave=outputdatabasedirectory,print.index = T,message.start="Started - wt translation speed table 30 -",message.end="Finished wt translation speed table 30 -")

wtatg40<-translationspeed.function(query.genes=gene,query.species="Saccharomyces cerevisiae",rampzonelength.fiveprime = 40,rampzonelength.threeprime = 40,codons.translationspeed = yeastcodonusage, save.file = T,outputfilename ="Initial Translation Speed Table 40.xlsx",directorysave=outputdatabasedirectory,print.index = F,message.start="Started - wt translation speed table 40 -",message.end="Finished wt translation speed table 40 -")

wtatg50<-translationspeed.function(query.genes=gene,query.species="Saccharomyces cerevisiae",rampzonelength.fiveprime = 50,rampzonelength.threeprime = 50,codons.translationspeed = yeastcodonusage, save.file = T,outputfilename ="Initial Translation Speed Table 50.xlsx",directorysave=outputdatabasedirectory,print.index = F,message.start="Started - wt translation speed table 50 -",message.end="Finished wt translation speed table 50 -")

wtatg60<-translationspeed.function(query.genes=gene,query.species="Saccharomyces cerevisiae",rampzonelength.fiveprime = 60,rampzonelength.threeprime = 60,codons.translationspeed = yeastcodonusage, save.file = T,outputfilename ="Initial Translation Speed Table 60.xlsx",directorysave=outputdatabasedirectory,print.index = F,message.start="Started - wt translation speed table 60 -",message.end="Finished wt translation speed table 60 -")

wtatg70<-translationspeed.function(query.genes=gene,query.species="Saccharomyces cerevisiae",rampzonelength.fiveprime = 70,rampzonelength.threeprime = 70,codons.translationspeed = yeastcodonusage, save.file = T,outputfilename ="Initial Translation Speed Table 70.xlsx",directorysave=outputdatabasedirectory,print.index = F,message.start="Started - wt translation speed table 70 -",message.end="Finished wt translation speed table 70 -")

wtatg80<-translationspeed.function(query.genes=gene,query.species="Saccharomyces cerevisiae",rampzonelength.fiveprime = 80,rampzonelength.threeprime = 80,codons.translationspeed = yeastcodonusage, save.file = T,outputfilename ="Initial Translation Speed Table 80.xlsx",directorysave=outputdatabasedirectory,print.index = F,message.start="Started - wt translation speed table 80 -",message.end="Finished wt translation speed table 80 -")

wtatg90<-translationspeed.function(query.genes=gene,query.species="Saccharomyces cerevisiae",rampzonelength.fiveprime = 90,rampzonelength.threeprime = 90,codons.translationspeed = yeastcodonusage, save.file = T,outputfilename ="Initial Translation Speed Table 90.xlsx",directorysave=outputdatabasedirectory,print.index = F,message.start="Started - wt translation speed table 90 -",message.end="Finished wt translation speed table 90 -")

wtatg100<-translationspeed.function(query.genes=gene,query.species="Saccharomyces cerevisiae",rampzonelength.fiveprime = 100,rampzonelength.threeprime = 100,codons.translationspeed = yeastcodonusage, save.file = T,outputfilename ="Initial Translation Speed Table 100.xlsx",directorysave=outputdatabasedirectory,print.index = F,message.start="Started - wt translation speed table 100 -",message.end="Finished wt translation speed table 100 -")

###Since ATG tends to be depleted at the start of genes, the initial translation speed is heavily influenced by ATG since it is one of the fastest codons. To account for this, all ATG will be "neutralized" with the average translation speed across all ORF at every position. In essence, the influence of ATG on translation speed in the first 40 codons vs rest is partialed out. 

atgneut.yeastcodonusage<-yeastcodonusage

atgneut.yeastcodonusage$cumRRT<-atgneut.yeastcodonusage$`Frame 1 (Coding) Observed Counts`*atgneut.yeastcodonusage$RRT

globalrrt<-sum(atgneut.yeastcodonusage$cumRRT)/sum(atgneut.yeastcodonusage$`Frame 1 (Coding) Observed Counts`)

atgneut.yeastcodonusage$RRT[atgneut.yeastcodonusage$Codons=="ATG"]<-globalrrt

atgneutralized40<-translationspeed.function(query.genes=gene,query.species="Saccharomyces cerevisiae",rampzonelength.fiveprime = 40,rampzonelength.threeprime=40,codons.translationspeed =atgneut.yeastcodonusage,save.file = T,outputfilename ="Initial Translation Speed Table ATG Neutralized 40.xlsx",directorysave=outputdatabasedirectory,print.index = F,message.start="Started atg neutralized translation speed table -",message.end="Finished atg neutralized translation speed table -")

```

####PART 2: PROTEIN BLASTS AND CONSERVATION SCORES CALCULATIONS

###Downloading, curating, and creating SUBJECT database.

```{r,echo=F,eval=F}

####Downloading protein sequences: Include Saccharomycotina (Taxonomy ID: 147537) and then I will remove all sequences from Saccharomyces (Taxonomy ID: 4930). To download, go to send file and then select fasta and then default order. 

# https://www.ncbi.nlm.nih.gov/Taxonomy/Browser/wwwtax.cgi?mode=Info&id=147537&lvl=3&lin=f&keep=1&srchmode=1&unlock

# https://www.uniprot.org/uniprotkb?dir=ascend&query=(taxonomy_id:147537)&sort=organism_name

###The ddbj, embl, genbank, refseq, and pir will be downloaded individually as fasta files from ncbi. Under the "Sources Database" section, click customize and only select the databases to be downloaded. Hover over the name of the database you want to download; you have to click the database (it will turn blue and there will be a check next to it). Make sure that only Fungi is selected for species. The site is buggy, so I always refresh the page in-between downloads. For uniprot, make sure that the taxonomy ID for Saccharomycotina is 147537.

###Combining Saccharomycotina fasta files.

sact1<-read.fasta(paste0(inputdatabasedirectory,"Saccharomycotina ddbj.fasta"),seqtype = "AA")

sact2<-read.fasta(paste0(inputdatabasedirectory,"Saccharomycotina embl.fasta"),seqtype = "AA")

sact3<-read.fasta(paste0(inputdatabasedirectory,"Saccharomycotina genbank.fasta"),seqtype = "AA")

sact4<-read.fasta(paste0(inputdatabasedirectory,"Saccharomycotina refseq.fasta"),seqtype = "AA")

sact5<-read.fasta(paste0(inputdatabasedirectory,"Saccharomycotina pir.fasta"),seqtype = "AA")

sact6<-read.fasta(paste0(inputdatabasedirectory,"Saccharomycotina uniprot.fasta"),seqtype = "AA")

####Since 6 databases were combined, I am checking to see if there are duplicate submissions in the combined Saccharomycotina dataset.

sactcombinefasta<-c(sact1,sact2,sact3,sact4,sact5,sact6)

sactcombinefastatest<-c(sact1,sactcombinefasta)

sexf<-getName(sactcombinefasta[duplicated(getName(sactcombinefasta))])

sexftest<-getName(sactcombinefastatest[duplicated(getName(sactcombinefastatest))])

if(length(sexf)==0&length(sexftest)==length(sact1)){
  
  print("SUCCESS THERE ARE NO DUPLICATE SUBMISSIONS!!!")
} else {
  
  base::stop("FAIL?? THERE ARE DUPLICATE SUBMISSIONS??")
}

###The ddbj, refseq, genbank, and esembl databases annotate genus and species in brackets []; the pir database annotates species in parentheses (); and the uniprot database annotates species with "OS=". Therefore, the most straight-forward way to get rid of all Saccharomyces species is by omitting all annotations that have "[Saccharomyces", or " (Saccharomyces", or "OS=Saccharomyces". However, there is added complexity, because a paper annotated Kazachstania barnettii and Geotrichum candidum sequences with the cerevisiae strain that the protein shares homology with, so it looks likes [Saccharomyces cerevisiae][Kazachstania barnettii] or [Saccharomyces cerevisiae][Geotrichum candidum]. https://www.ncbi.nlm.nih.gov/protein/CAD1785470.1 These entries are really for Kazachstania barnettii or Geotrichum candidum and contain no Saccharomyces relationship (besides protein homology). This code will identify and isolate all sequences annotated as Saccharomyces, but will keep entries that have "[Kazachstania barnettii]" or "[Geotrichum candidum]" in its annotation. 

rawcompileddatabase<-sactcombinefasta

generep<-1

saccharomycesspecies<-NULL

timebegin.subjectdatabase<-timelapsebegin.function(message.begin="Started compiling subject database -")

print(timebegin.subjectdatabase[[7]])

while (generep<(length(rawcompileddatabase)+1)){
  
  print(generep)
  
  if(grepl("[Saccharomyces",getAnnot(rawcompileddatabase[generep]),fixed = T)==T&grepl("[Kazachstania barnettii]",getAnnot(rawcompileddatabase[generep]),fixed = T)==F&grepl("[Geotrichum candidum]",getAnnot(rawcompileddatabase[generep]),fixed = T)==F){
    
    saccharomycesspecies<-c(saccharomycesspecies,generep)
  } else if(grepl("OS=Saccharomyces",getAnnot(rawcompileddatabase[generep]),fixed = T)==T){
    
    saccharomycesspecies<-c(saccharomycesspecies,generep)
  } else if(grepl(" (Saccharomyces",getAnnot(rawcompileddatabase[generep]),fixed = T)==T){
    
    saccharomycesspecies<-c(saccharomycesspecies,generep)
  }
  
  generep<-generep+1
}

timeended.subjectdatabase<-timelapseend.function(startyear=timebegin.subjectdatabase[[1]],startmonth=timebegin.subjectdatabase[[2]],startday=timebegin.subjectdatabase[[3]],starthour=timebegin.subjectdatabase[[4]],startminutes=timebegin.subjectdatabase[[5]],startseconds=timebegin.subjectdatabase[[6]],timestarted=timebegin.subjectdatabase[[7]],message.fin="Finished compiling subject database -  ")

print(timeended.subjectdatabase[[1]])

print(timeended.subjectdatabase[[2]])

remove(timeended.subjectdatabase)

print(paste0("Removing all Saccharomyces species from subject database..."))

rawcompileddatabasecurated<-rawcompileddatabase[-saccharomycesspecies]

saccharomycotina.list<-list(Saccharomycotina.full=rawcompileddatabase,Saccharomyces=saccharomycesspecies)

print(paste0("Saving..."))

save(saccharomycotina.list, file=paste0(outputdatabasedirectory,"rawcompiledsaccharomycotinadatabase.RData"))

remove(saccharomycotina.list)

save(rawcompileddatabasecurated, file=paste0(outputdatabasedirectory,"rawcompileddatabasecurated.RData"))

timeended.subjectdatabase<-timelapseend.function(startyear=timebegin.subjectdatabase[[1]],startmonth=timebegin.subjectdatabase[[2]],startday=timebegin.subjectdatabase[[3]],starthour=timebegin.subjectdatabase[[4]],startminutes=timebegin.subjectdatabase[[5]],startseconds=timebegin.subjectdatabase[[6]],timestarted=timebegin.subjectdatabase[[7]],message.fin="Finished saving full subject database -  ")

print(timeended.subjectdatabase[[1]])

print(timeended.subjectdatabase[[2]])

remove(timeended.subjectdatabase)

###This code will show all of the Saccharomyces species omitted from the Saccharomycotina database. This was used to identify if there were any sequences that had weird annotations that caused the code to remove non-Saccharomyces sequences. It seems like the omitted sequences were exclusively strains from Saccharomyces, whether haploid or diploid.

generep<-1

outputsaccharomycesspecies<-rep(NA,length(saccharomycesspecies))

annotoutputsaccharomycesspecies<-rep(NA,length(saccharomycesspecies))

while(generep<(length(saccharomycesspecies)+1)){
  
  print(generep)

  if(grepl(" OS=",getAnnot(rawcompileddatabase[saccharomycesspecies[generep]]),fixed = T)==T){
  yoyo<-c2s(getAnnot(rawcompileddatabase[saccharomycesspecies[generep]]))
  
  tempannotfirst<-unlist(strsplit(yoyo,"OS=Saccharomyces",fixed=T))[2]
  
  tempannotsecond<-s2c(unlist(strsplit(tempannotfirst," OX=",fixed=T))[1])

  outputsaccharomycesspecies[generep]<-c2s(c("Saccharomyces",c2s(tempannotsecond[-length(tempannotsecond)])))
  
  annotoutputsaccharomycesspecies[generep]<-unlist(getAnnot(rawcompileddatabase[saccharomycesspecies[generep]]))
  } else if(grepl("[Saccharomyces",getAnnot(rawcompileddatabase[saccharomycesspecies[generep]]),fixed = T)==T){
    
    yoyo<-c2s(getAnnot(rawcompileddatabase[saccharomycesspecies[generep]]))
    
    tempannotfirst<-unlist(strsplit(yoyo,"[Saccharomyces",fixed=T))[2]
    
    tempannotsecond<-s2c(tempannotfirst)
    
    outputsaccharomycesspecies[generep]<-c2s(c("Saccharomyces",c2s(tempannotsecond[-length(tempannotsecond)])))
    
    annotoutputsaccharomycesspecies[generep]<-unlist(getAnnot(rawcompileddatabase[saccharomycesspecies[generep]]))
  } else if(grepl("(Saccharomyces",getAnnot(rawcompileddatabase[saccharomycesspecies[generep]]),fixed = T)==T){
    
    yoyo<-c2s(getAnnot(rawcompileddatabase[saccharomycesspecies[generep]]))
    
    tempannotfirst<-unlist(strsplit(yoyo,"(Saccharomyces",fixed=T))[2]
    
    tempannotsecond<-s2c(tempannotfirst)
    
    outputsaccharomycesspecies[generep]<-c2s(c("Saccharomyces",c2s(tempannotsecond[-length(tempannotsecond)])))
    
    annotoutputsaccharomycesspecies[generep]<-unlist(getAnnot(rawcompileddatabase[saccharomycesspecies[generep]]))
  }
  
  generep<-generep+1
}

alloutputsaccharomycesspecies<-data.frame(Species.Omitted=outputsaccharomycesspecies,Full.Annotation=annotoutputsaccharomycesspecies)

nodupoutputsaccharomycesspecies<-alloutputsaccharomycesspecies[!duplicated(alloutputsaccharomycesspecies$Species.Omitted),]

nodupoutputsaccharomycesspecies$`Number of Entries`<-99999

generep<-1

while(generep<nrow(nodupoutputsaccharomycesspecies)+1){

  print(generep)
  
  nodupoutputsaccharomycesspecies$`Number of Entries`[generep]<-nrow(alloutputsaccharomycesspecies[alloutputsaccharomycesspecies$Species.Omitted==nodupoutputsaccharomycesspecies$Species.Omitted[generep],])
    
  generep<-generep+1
}

nodupoutputsaccharomycesspecies<-nodupoutputsaccharomycesspecies[,c("Species.Omitted","Number of Entries","Full.Annotation")]

wb<-createWorkbook()

addWorksheet(wb, "Every Entry")

addWorksheet(wb, "Unique Species")

writeData(wb,sheet="Every Entry",x=alloutputsaccharomycesspecies)

writeData(wb,sheet="Unique Species",x=nodupoutputsaccharomycesspecies)

saveWorkbook(wb, paste0(outputdatabasedirectory,"Omitted Saccharomyces Sequences.xlsx"), overwrite = T)

### The problem is that the uniprot sequences annotate species by "OS=" This inconsistency will affect how species are parsed later in the code since brackets or parentheses are needed to parse species. Also, the getAnnot function() returns a character with a >, and the write.fasta() function puts a > by default. Therefore, there will be two > after using write.fasta making the file unreadable. The > has to be removed from all annotations. Annotations will be changed to have all species in brackets, and the annotations which will be stored in a list. Then, a new fasta file will be made with all of the Saccharomycotina sequences (minus Saccharomyces) as the subject database.

templist<-rep(list(NA),length(rawcompileddatabasecurated))

generep<-1

while (generep<(length(rawcompileddatabasecurated)+1)){
  
  print(generep)
  
  if(grepl(" OS=",getAnnot(rawcompileddatabasecurated[generep]),fixed = T)==T){
    yoyo<-c2s(getAnnot(rawcompileddatabasecurated[generep]))
    
    ssksk<-s2c(yoyo)
    
    tempannotfirst<-unlist(strsplit(yoyo," OS=",fixed=T))[2]
    
    tempannotsecond<-unlist(strsplit(tempannotfirst," OX=",fixed=T))[1]
    
    remadeannotat<-c("[",s2c(tempannotsecond),"]")
    
    kikip<-c2s(c(ssksk[-1]," ",remadeannotat))
    
    templist[generep]<-list(kikip)
  } else if(grepl(" OS=",getAnnot(rawcompileddatabasecurated[generep]),fixed = T)!=T){
    
    yoyo<-c2s(getAnnot(rawcompileddatabasecurated[generep]))
    
    ssksk<-s2c(yoyo)
    
    kikip<-c2s(ssksk[-1])
    
    templist[generep]<-list(kikip)
  } else if(grepl("[Saccharomyces",getAnnot(rawcompileddatabasecurated[generep]),fixed = T)==T&grepl("[Kazachstania barnettii]",getAnnot(rawcompileddatabasecurated[generep]),fixed = T)==T){
    
    yoyo<-c2s(getAnnot(rawcompileddatabasecurated[generep]))
    
    tempannotfirst<-unlist(strsplit(yoyo,"[Saccharomyces",fixed=T))[1]
    
    kikip<-c2s(c(s2c(tempannotfirst)[c(-1)],"[Kazachstania barnettii]"))
    
    templist[generep]<-list(kikip)
  } else if(grepl("[Saccharomyces",getAnnot(rawcompileddatabasecurated[generep]),fixed = T)==T&grepl("[Geotrichum candidum]",getAnnot(rawcompileddatabasecurated[generep]),fixed = T)==T){
    
    yoyo<-c2s(getAnnot(rawcompileddatabasecurated[generep]))
    
    tempannotfirst<-unlist(strsplit(yoyo,"[Saccharomyces",fixed=T))[1]
    
    kikip<-c2s(c(s2c(tempannotfirst)[c(-1)],"[Geotrichum candidum]"))
    
    templist[generep]<-list(kikip)
  }
  
  generep<-generep+1
}

if((length(sactcombinefasta)-length(saccharomycesspecies))==length(rawcompileddatabasecurated)){
  
  print("SUCCESS!!! Saccharomycotina without Saccharomyces created!!!")
} else {
  
  base::stop("FAIL?? LENGTH OF SUBJECT DATABASE IS DIFFERENT THAN WHAT IS EXPECTED FOLLOWING THE REMOVAL OF SACCHAROMYCES??")
}

write.fasta(rawcompileddatabasecurated,file=paste0(outputdatabasedirectory,"Curated Saccharomycotina dataset with no Saccharomyces.fasta"),names=templist)

timeended.subjectdatabase<-timelapseend.function(startyear=timebegin.subjectdatabase[[1]],startmonth=timebegin.subjectdatabase[[2]],startday=timebegin.subjectdatabase[[3]],starthour=timebegin.subjectdatabase[[4]],startminutes=timebegin.subjectdatabase[[5]],startseconds=timebegin.subjectdatabase[[6]],timestarted=timebegin.subjectdatabase[[7]],message.fin="Finished compiling subject database without Saccharomyces -  ")

print(timeended.subjectdatabase[[1]])

print(timeended.subjectdatabase[[2]])

remove(sact1,sact2,sact3,sact4,sact5,sact6,sactcombinefasta,sactcombinefastatest,sexftest,saccharomycesspecies,rawcompileddatabase,rawcompileddatabasecurated,templist,alloutputsaccharomycesspecies,nodupoutputsaccharomycesspecies)

### The NCBI BLAST function won't work if there are spaces or - in your directory or file name. Also, the default location of the subject database is the default/home directory, and this cannot be changed. I changed the name of "Curated Saccharomycotina dataset with no Saccharomyces.fasta" to "Saccharomycotina_without_Saccharomyces.fasta" and put it my default directory (D:).

```

###Splitting and saving query proteins as fasta.

```{r,echo=F,eval=F}

###The query sequences are S. cerevisiae proteins.

#http://sgd-archive.yeastgenome.org/sequence/S288C_reference/orf_protein/

queryproteinfile<-paste0(inputdatabasedirectory,"orf_trans_R64-3-1_20210421.fasta")

###The BLAST function only recognizes sequences that are formatted as StringSet data.

fullqueryproteinsstringset<-readAAStringSet(queryproteinfile, format = "fasta")

###Since the BLASTS are long and there might be an interest in pseudogenes and dubious ORF, BLASTS will be done on all proteins, and then the data will be subset later.

fullqueryproteins.blast<-seqinr::read.fasta(queryproteinfile, seqtype = "AA")

fullquerydescriptives<-data.frame(gene.name="ZZZZZ",common.name="ZZZZZ",annotation="ZZZZZ",stringset.name="ZZZZZ",Amino.Acids=9999999,Blank=1:length(fullqueryproteins.blast))

generep<-1

while(generep<(nrow(fullquerydescriptives)+1)){
  fullquerydescriptives$gene.name[generep]<-getName(fullqueryproteins.blast[generep])
  
  fullquerydescriptives$common.name[generep]<-unlist(strsplit(unlist(getAnnot(fullqueryproteins.blast[generep]))," ",fixed=T))[2]
  
  fullquerydescriptives$annotation[generep]<-unlist(getAnnot(fullqueryproteins.blast[generep]))
  
  fullquerydescriptives$stringset.name[generep]<-c2s(s2c(c2s(getAnnot(fullqueryproteins.blast[generep])))[-1])
  
  fullquerydescriptives$Amino.Acids[generep]<-length(unlist(getSequence(fullqueryproteins.blast[generep])))-1
  
  generep<-generep+1
}

###Generating FASTA of proteins spanning from midpoint to stop codon

###In order to do BLAST beginning in the middle of proteins, the full-length query protein sequences will be split from the midpoint of the protein until the stop codon. These truncated sequences will be saved as a fasta file.

sequences.middlebottom<-rep(list(NA), length(fullqueryproteins.blast))

annotationlist.middlebottom<-rep(list(NA), length(fullqueryproteins.blast))

sampled.middlebottom<-data.frame(Name="ZZZZZ",FullProteinlength=9999999,MidProteinlength=9999999,Annotation="ZZZZZ",Order=1:length(fullqueryproteins.blast))

generep<-1

while (generep<(length(fullqueryproteins.blast)+1)) {

  wholesequence<-unlist(getSequence(fullqueryproteins.blast[generep]))
  
  middlebottom<-wholesequence[floor((length(wholesequence)-1)/2):length(wholesequence)]
  
  if(unlist(middlebottom[length(middlebottom)])!="*"&unlist(wholesequence[length(wholesequence)])=="*"){
    middlebottom<-c(middlebottom,"*")
  }
  
  sequences.middlebottom[generep]<-list(middlebottom)
  
  names(sequences.middlebottom)[generep]<-getName(fullqueryproteins.blast[generep])
  
  annotationlist.middlebottom[generep]<-c2s(s2c(unlist(getAnnot(fullqueryproteins.blast[generep])))[-1])
  
  names(annotationlist.middlebottom)[generep]<-getName(fullqueryproteins.blast[generep])

  sampled.middlebottom$Name[generep]<-getName(fullqueryproteins.blast[generep])
  
  sampled.middlebottom$FullProteinlength[generep]<-(length(wholesequence)-1)
  
  sampled.middlebottom$MidProteinlength[generep]<-(length(middlebottom)-1)
  
  sampled.middlebottom$Annotation[generep]<-unlist(getAnnot(fullqueryproteins.blast[generep]))
  
  generep<-generep+1
}

sampled.middlebottom$`Full/Mid`<-round(sampled.middlebottom$FullProteinlength/sampled.middlebottom$MidProteinlength,digits = 0.01)

###Generating FASTA of proteins ranging from start to midpoint of ORF

###Similarly half of the protein sequences will be prepared for the BLASTS, but this time it will span from the start codon to the midpoint of each protein. The midpoint-to-end sequences and start-to-midpoint sequences are cut in half in a precise way that when combined they have the exact sequences as the full wildtype sequence.

sequences.topmiddle<-rep(list(NA), length(fullqueryproteins.blast))

annotationlist.topmiddle<-rep(list(NA), length(fullqueryproteins.blast))

sampled.topmiddle<-data.frame(Name="ZZZZZ",FullProteinlength=9999999,MidProteinlength=9999999,Annotation="ZZZZZ",Order=1:length(fullqueryproteins.blast))

generep<-1

while(generep<(length(fullqueryproteins.blast)+1)){

  wholesequence<-unlist(getSequence(fullqueryproteins.blast[generep]))
  
  middletop<-wholesequence[1:((round(length(wholesequence)-1)/2)-1)]

  sequences.topmiddle[generep]<-list(middletop)
  
  names(sequences.topmiddle)[generep]<-getName(fullqueryproteins.blast[generep])
  
  annotationlist.topmiddle[generep]<-c2s(s2c(unlist(getAnnot(fullqueryproteins.blast[generep])))[-1])
  
  names(annotationlist.topmiddle)[generep]<-getName(fullqueryproteins.blast[generep])

  sampled.topmiddle$Name[generep]<-getName(fullqueryproteins.blast[generep])
  
  sampled.topmiddle$FullProteinlength[generep]<-(length(wholesequence)-1)
  
  sampled.topmiddle$MidProteinlength[generep]<-length(middletop)
  
  sampled.topmiddle$Annotation[generep]<-unlist(getAnnot(fullqueryproteins.blast[generep]))
  
  generep<-generep+1
}

sampled.topmiddle$`Full/Mid`<-round(sampled.topmiddle$FullProteinlength/sampled.topmiddle$MidProteinlength,digits = 0.01)

##This code verifies that all of the proteins are split exactly in the middle, and that the first amino acid of the second half immediately follows the last amino acid of the first half.

verify.splitgenes<-data.frame(Name="ZZZZZ",Amino.Acids=99999,Full="ZZZZZ",First.Half="ZZZZZ",Second.Half="ZZZZZ",Combined="ZZZZZ",FullvsCombined="ZZZZZ",Index=1:length(fullqueryproteins.blast))

generep<-1

while(generep<length(fullqueryproteins.blast)+1){
  
  genename<-names(sequences.topmiddle[generep])
  
  comp.sequences<-c(unlist(getSequence(sequences.topmiddle[genename])),unlist(getSequence(sequences.middlebottom[genename])))
  
  verify.splitgenes$Name[generep]<-genename
  
  verify.splitgenes$Amino.Acids[generep]<-length((unlist(getSequence(fullqueryproteins.blast[genename]))))-1
  
  verify.splitgenes$Full[generep]<-c2s(unlist(getSequence(fullqueryproteins.blast[genename])))
  
  verify.splitgenes$First.Half[generep]<-c2s(unlist(getSequence(sequences.topmiddle[genename])))
  
  verify.splitgenes$Second.Half[generep]<-c2s(unlist(getSequence(sequences.middlebottom[genename])))
  
  verify.splitgenes$Combined[generep]<-c2s(comp.sequences)
  
  verify.splitgenes$FullvsCombined[generep]<-if(identical(comp.sequences,unlist(getSequence(fullqueryproteins.blast[genename])))==T){
    "Same"
  } else{
    "Different"
  }
  
  generep<-generep+1
}

if(nrow(verify.splitgenes[verify.splitgenes$FullvsCombined=="Same",])==length(sequences.middlebottom)&nrow(verify.splitgenes[verify.splitgenes$FullvsCombined=="Same",])==length(sequences.topmiddle)){
    
  if(identical(names(sequences.topmiddle),names(annotationlist.topmiddle))==T&nrow(sampled.topmiddle[sampled.topmiddle$`Full/Mid`==2,])==nrow(sampled.topmiddle)){
    
    print("SUCCESS!!! FIRST HALF SEQUENCES IS HALF THE LENGTH OF FULL-LENGTH SEQUENCES!!!")
  } else if(identical(names(sequences.topmiddle),names(annotationlist.topmiddle))!=T){
    
    base::warning("FAIL?? FIRST HALF SEQUENCES AND ANNOTATIONS DO NOT MATCH??")
  } else if(nrow(sampled.topmiddle[sampled.topmiddle$`Full/Mid`==2,])!=nrow(sampled.topmiddle)){
    
    base::warning("FAIL?? FIRST HALF SEQUENCES NOT CUT IN HALF??")
  } else if(identical(names(sequences.topmiddle),names(annotationlist.topmiddle))!=T&nrow(sampled.topmiddle[sampled.topmiddle$`Full/Mid`==2,])!=nrow(sampled.topmiddle)){
    
    base::warning("COMPLETE FAIL?? NOTHING WORKED FOR FIRST HALF??")
  }
  
  if(identical(names(sequences.middlebottom),names(annotationlist.middlebottom))==T&nrow(sampled.middlebottom[sampled.middlebottom$`Full/Mid`==2,])==nrow(sampled.middlebottom)){
    
    print("SUCCESS!!! SECOND HALF SEQUENCES IS HALF THE LENGTH OF FULL-LENGTH SEQUENCES!!!")
  } else if(identical(names(sequences.middlebottom),names(annotationlist.middlebottom))!=T){
    
    base::warning("FAIL?? SECOND HALF SEQUENCES AND ANNOTATIONS DO NOT MATCH??")
  } else if(nrow(sampled.middlebottom[sampled.middlebottom$`Full/Mid`==2,])!=nrow(sampled.middlebottom)){
    
    base::warning("FAIL?? SECOND HALF SEQUENCES NOT CUT IN HALF??")
  } else if(identical(names(sequences.middlebottom),names(annotationlist.middlebottom))!=T&nrow(sampled.middlebottom[sampled.middlebottom$`Full/Mid`==2,])!=nrow(sampled.middlebottom)){
    
    base::warning("COMPLETE FAIL?? NOTHING WORKED FOR SECOND HALF??")
  }
  
  print("SUCCESS!!! FULL-LENGTH SEQUENCES ARE THE SAME AS THE COMBINED FIRST AND SECOND HALF SEQUENCES!!!")
} else{
  
  base::stop("FAIL?? FIRST HALF AND SECOND HALF ARE DIFFERENT FROM FULL-LENGTH SEQUENCE??")
}

write.fasta(sequences.topmiddle,file=paste0(outputdatabasedirectory,"All Saccharomyces cerevisiae ORF from start to midpoint.fasta"),names=annotationlist.topmiddle)

write.fasta(sequences.middlebottom,file=paste0(outputdatabasedirectory,"All Saccharomyces cerevisiae ORF from midpoint to stop.fasta"),names=annotationlist.middlebottom)

```

###BLASTS of Full-Length Proteins.

```{r,echo=F,eval=F}

###The BLASTS are computationally intensive and will require more RAM than my computer can handle to avoid crashes due to running out of memory. Due to memory and time constraints, the protein list will be split into 6 parts and then the blasts will be combined later. To further conserve RAM-space, it is best to remove all objects that are not necessary for proteinBLAST.function; this can be easily done by selecting grid view in the environment, selecting what is not needed, and then clearing those objects. For around 3000000 hits, it takes about 17 hours for the BLASTS to complete, and an additional 30 hours for the annotations to be added.

queryproteinfile<-paste0(inputdatabasedirectory,"orf_trans_R64-3-1_20210421.fasta")

###The BLAST function only recognizes sequences that are formatted as StringSet data

fullqueryproteinsstringset<-readAAStringSet(queryproteinfile, format = "fasta")

###Since the BLASTS are long and there might be an interest in pseudogenes and dubious ORF, BLASTS will be done on all proteins, and then the data will be subset later.

fullqueryproteins.blast<-seqinr::read.fasta(queryproteinfile, seqtype = "AA")

fullquerydescriptives<-data.frame(gene.name="ZZZZZ",common.name="ZZZZZ",annotation="ZZZZZ",stringset.name="ZZZZZ",Amino.Acids=9999999,Blank=1:length(fullqueryproteins.blast))

generep<-1

while(generep<(nrow(fullquerydescriptives)+1)){
  
  fullquerydescriptives$gene.name[generep]<-getName(fullqueryproteins.blast[generep])

  fullquerydescriptives$common.name[generep]<-unlist(strsplit(unlist(getAnnot(fullqueryproteins.blast[generep]))," ",fixed=T))[2]
  
  fullquerydescriptives$annotation[generep]<-unlist(getAnnot(fullqueryproteins.blast[generep]))
  
  fullquerydescriptives$stringset.name[generep]<-c2s(s2c(c2s(getAnnot(fullqueryproteins.blast[generep])))[-1])
  
  fullquerydescriptives$Amino.Acids[generep]<-length(unlist(getSequence(fullqueryproteins.blast[generep])))-1
  
  generep<-generep+1
}

gene.all<-fullqueryproteinsstringset

genename.all<-fullquerydescriptives

split1numba<-round(length(gene.all)/6)

split2numba<-split1numba+split1numba+1

split3numba<-split1numba+split2numba+1

split4numba<-split1numba+split3numba+1

split5numba<-split1numba+split4numba+1

split6numba<-length(gene.all)

gene.pt1<-gene.all[1:split1numba]

genename.pt1<-genename.all[1:split1numba,]
  
gene.pt2<-gene.all[(split1numba+1):split2numba]

genename.pt2<-genename.all[(split1numba+1):split2numba,]

gene.pt3<-gene.all[(split2numba+1):split3numba]

genename.pt3<-genename.all[(split2numba+1):split3numba,]

gene.pt4<-gene.all[(split3numba+1):split4numba]

genename.pt4<-genename.all[(split3numba+1):split4numba,]

gene.pt5<-gene.all[(split4numba+1):split5numba]

genename.pt5<-genename.all[(split4numba+1):split5numba,]

gene.pt6<-gene.all[(split5numba+1):split6numba]

genename.pt6<-genename.all[(split5numba+1):split6numba,]

###The default settings of the NCBI BLAST will be used, except that the maximum number of alignments (1000000000) will be returned for every BLAST, otherwise the default is 500 alignments. The annotations for the subject hits are added after all BLASTS have been conducted. Don't worry if you get an error message that reads "Error in read.table(outfile, sep = ",", quote = "") : no lines available in input"; that just means that a query sequence had no BLAST hits, which will be recorded in the BLAST statistics. To find out the meaning of qstart, sstart, send, etc. view this link https://www.ncbi.nlm.nih.gov/books/NBK279684/  "queryproteinsstringset" is the query list of proteins which must be formatted using the readAAStringSet() function. "querydescriptives" must be a data frame containing the names, annotations, and other important features of the query sequences; the BLAST function refers to specific column names in the "querydescriptives" data frame, so refer to the example BLAST function to know exactly how to format the column names or else this code won't work. "make.database" gives the option (True or False) if you want to create the subject database in your home directory; each subject database is permanent and will not leave your home directory if you quit R session. "load.database" gives the option (True or False) to load all of the proteins in the local subject database as an object in the environment using the read.fasta() function; this takes a long time because the subject database will oftentimes have hundreds of thousands of sequences. "proteindatabase" is the string containing the directory path where the local subject database is located; by default this should be your home directory since the NCBI tool automatically creates the local database in your home directory. "query.species" is an optional string for the species that the query sequences originated from. "proteinspan" is an optional string to convey the region of the query sequence that will be BLASTed. "number.of.proteins" is a numerical value of how many proteins are queried by the BLAST function; this value must be the exact same as the length of proteins in "querydescriptives". "bitscore.threshold" is the minimum bitscore value from each sequence alignment to include in the BLAST results; the total number of sequence alignments a query has will be recorded regardless of bitscore value, but all BLAST descriptions (alignment regions, subject annotations, qstart, E value, etc) from alignments the same or higher than "bitscore.threshold" will be returned. "rawblastresultsname" should be the string of the name for the data frame containing the raw BLAST results. "sampledblastresultsname" is the string for the name of the data frame that contain statistics regarding each query protein BLAST results. "save.file" gives the option (True or False) to save the BLAST output as an R file with the extension ".RData". If you choose to save the file using this code, then "rdatablastname" is the string of the file name (requires extension ".RData") for the output and "directorysave" should be a string for the directory to save the file. "print.index" gives the option (True or False) to print the current numerical index in the fasta file that is being analyzed; for example, an index of 5 means that the fifth protein listed in the fasta file is currently being BLASTed. "save.csv" gives the option (True or False) to save and export all of the BLAST results as a file with the .csv extension. If you choose to export all of the BLAST results as csv, then two csv files will be saved; "blastfilename" is the string for the name of csv file with the raw BLAST results, and "blastreportname" is the string for the name of the csv file with the statistics regarding each query protein BLAST results. "message.start" is the message for the time elapse at the start of the function. "message.end" is the message for the time elapse after the function has completed.

fulllengthproteindatabase<-"D:/Saccharomycotina_without_Saccharomyces.fasta"

fullblastoutput.pt1<-proteinBLAST.function(queryproteinsstringset = gene.pt1, querydescriptives = genename.pt1, make.database = T, load.database = T, proteindatabase = fulllengthproteindatabase, query.species = "Saccharomyces cerevisiae", proteinspan = "Full Length", number.of.proteins = length(gene.pt1), bitscore.threshold = 50, rawblastresultsname = "pt1.fullrawblastresults", sampledblastresultsname = "pt1.fullsampledblastresults", save.file = T, directorysave = outputdatabasedirectory, rdatablastname = "Full Protein BLASTS List pt1.RData", save.csv = F, print.index = T, message.start = "STARTED FULL LENGTH BLASTS pt1 -", message.end = "FINISHED FULL LENGTH BLASTS pt1 -")

###All BLASTS under every context will be run using the exact same subject database, so the "make.database" argument is set to False after the database has been initially created.

fullblastoutput.pt2<-proteinBLAST.function(queryproteinsstringset = gene.pt2, querydescriptives = genename.pt2, make.database = F, load.database = T, proteindatabase = fulllengthproteindatabase, query.species = "Saccharomyces cerevisiae", proteinspan = "Full Length", number.of.proteins = length(gene.pt2), bitscore.threshold = 50, rawblastresultsname = "pt2.fullrawblastresults", sampledblastresultsname = "pt2.fullsampledblastresults", save.file = T, directorysave = outputdatabasedirectory, rdatablastname = "Full Protein BLASTS List pt2.RData", save.csv = F, print.index = T, message.start = "STARTED FULL LENGTH BLASTS pt2 -", message.end = "FINISHED FULL LENGTH BLASTS pt2 -")

fullblastoutput.pt3<-proteinBLAST.function(queryproteinsstringset = gene.pt3, querydescriptives = genename.pt3, make.database = F, load.database = T, proteindatabase = fulllengthproteindatabase, query.species = "Saccharomyces cerevisiae", proteinspan = "Full Length", number.of.proteins = length(gene.pt3), bitscore.threshold = 50, rawblastresultsname = "pt3.fullrawblastresults", sampledblastresultsname = "pt3.fullsampledblastresults", save.file = T, directorysave = outputdatabasedirectory, rdatablastname = "Full Protein BLASTS List pt3.RData", save.csv = F, print.index = T, message.start = "STARTED FULL LENGTH BLASTS pt3 -", message.end = "FINISHED FULL LENGTH BLASTS pt3 -")

fullblastoutput.pt4<-proteinBLAST.function(queryproteinsstringset = gene.pt4, querydescriptives = genename.pt4, make.database = F, load.database = T, proteindatabase = fulllengthproteindatabase, query.species = "Saccharomyces cerevisiae", proteinspan = "Full Length", number.of.proteins = length(gene.pt4), bitscore.threshold = 50, rawblastresultsname = "pt4.fullrawblastresults", sampledblastresultsname = "pt4.fullsampledblastresults", save.file = T, directorysave = outputdatabasedirectory, rdatablastname = "Full Protein BLASTS List pt4.RData", save.csv = F, print.index = T, message.start = "STARTED FULL LENGTH BLASTS pt4 -", message.end = "FINISHED FULL LENGTH BLASTS pt4 -")

fullblastoutput.pt5<-proteinBLAST.function(queryproteinsstringset = gene.pt5, querydescriptives = genename.pt5, make.database = F, load.database = T, proteindatabase = fulllengthproteindatabase, query.species = "Saccharomyces cerevisiae", proteinspan = "Full Length", number.of.proteins = length(gene.pt5), bitscore.threshold = 50, rawblastresultsname = "pt5.fullrawblastresults", sampledblastresultsname = "pt5.fullsampledblastresults", save.file = T, directorysave = outputdatabasedirectory, rdatablastname = "Full Protein BLASTS List pt5.RData", save.csv = F, print.index = T, message.start = "STARTED FULL LENGTH BLASTS pt5 -", message.end = "FINISHED FULL LENGTH BLASTS pt5 -")

fullblastoutput.pt6<-proteinBLAST.function(queryproteinsstringset = gene.pt6, querydescriptives = genename.pt6, make.database = F, load.database = T, proteindatabase = fulllengthproteindatabase, query.species = "Saccharomyces cerevisiae", proteinspan = "Full Length", number.of.proteins = length(gene.pt6), bitscore.threshold = 50, rawblastresultsname = "pt6.fullrawblastresults", sampledblastresultsname = "pt6.fullsampledblastresults", save.file = T, directorysave = outputdatabasedirectory, rdatablastname = "Full Protein BLASTS List pt6.RData", save.csv = F, print.index = T, message.start = "STARTED FULL LENGTH BLASTS pt6 -", message.end = "FINISHED FULL LENGTH BLASTS pt6 -")

```

###BLASTS of Midpoint-to-Stop of Proteins.

```{r,echo=F,eval=F}

middlebottomqueryproteins.blast<-read.fasta(paste0(outputdatabasedirectory,"All Saccharomyces cerevisiae ORF from midpoint to stop.fasta"), seqtype="AA")

middlebottomqueryproteinsstringset<-readAAStringSet(paste0(outputdatabasedirectory,"All Saccharomyces cerevisiae ORF from midpoint to stop.fasta"), format = "fasta")

middlebottomquerydescriptives<-data.frame(gene.name="ZZZZZ",common.name="ZZZZZ",annotation="ZZZZZ",stringset.name="ZZZZZ",Amino.Acids=9999999,Blank=1:length(middlebottomqueryproteins.blast))

generep<-1

while(generep<(nrow(middlebottomquerydescriptives)+1)){
  
  middlebottomquerydescriptives$gene.name[generep]<-getName(middlebottomqueryproteins.blast[generep])
  
  middlebottomquerydescriptives$common.name[generep]<-unlist(strsplit(unlist(getAnnot(middlebottomqueryproteins.blast[generep]))," ",fixed=T))[2]
  
  middlebottomquerydescriptives$annotation[generep]<-unlist(getAnnot(middlebottomqueryproteins.blast[generep]))
  
  middlebottomquerydescriptives$stringset.name[generep]<-c2s(s2c(c2s(getAnnot(middlebottomqueryproteins.blast[generep])))[-1])
  
  middlebottomquerydescriptives$Amino.Acids[generep]<-length(unlist(getSequence(middlebottomqueryproteins.blast[generep])))-1
  
  generep<-generep+1
}

gene.all<-middlebottomqueryproteinsstringset

genename.all<-middlebottomquerydescriptives

split1numba<-round(length(gene.all)/6)

split2numba<-split1numba+split1numba+1

split3numba<-split1numba+split2numba+1

split4numba<-split1numba+split3numba+1

split5numba<-split1numba+split4numba+1

split6numba<-length(gene.all)

gene.pt1<-gene.all[1:split1numba]

genename.pt1<-genename.all[1:split1numba,]
  
gene.pt2<-gene.all[(split1numba+1):split2numba]

genename.pt2<-genename.all[(split1numba+1):split2numba,]

gene.pt3<-gene.all[(split2numba+1):split3numba]

genename.pt3<-genename.all[(split2numba+1):split3numba,]

gene.pt4<-gene.all[(split3numba+1):split4numba]

genename.pt4<-genename.all[(split3numba+1):split4numba,]

gene.pt5<-gene.all[(split4numba+1):split5numba]

genename.pt5<-genename.all[(split4numba+1):split5numba,]

gene.pt6<-gene.all[(split5numba+1):split6numba]

genename.pt6<-genename.all[(split5numba+1):split6numba,]

###All BLASTS under every context will be run using the exact same subject database, so the "make.database" argument is set to False after the database has been initially created.

fulllengthproteindatabase<-"D:/Saccharomycotina_without_Saccharomyces.fasta"

middlebottomblastoutput.pt1<-proteinBLAST.function(queryproteinsstringset = gene.pt1, querydescriptives = genename.pt1, make.database = F, load.database = T, proteindatabase = fulllengthproteindatabase, query.species = "Saccharomyces cerevisiae", proteinspan = "midpoint to stop", number.of.proteins = length(gene.pt1), bitscore.threshold = 50, rawblastresultsname = "pt1.middlebottomrawblastresults", sampledblastresultsname = "pt1.middlebottomsampledblastresults", save.file = T, directorysave = outputdatabasedirectory, rdatablastname = "Midpoint-to-Stop Protein BLASTS List pt1.RData", save.csv = F, print.index = T, message.start ="STARTED MIDDLE BOTTOM BLASTS pt1 -", message.end = "FINISHED MIDDLE BOTTOM BLASTS pt1 -")

middlebottomblastoutput.pt2<-proteinBLAST.function(queryproteinsstringset = gene.pt2, querydescriptives = genename.pt2, make.database = F, load.database = T, proteindatabase = fulllengthproteindatabase, query.species = "Saccharomyces cerevisiae", proteinspan = "midpoint to stop", number.of.proteins = length(gene.pt2), bitscore.threshold = 50, rawblastresultsname = "pt2.middlebottomrawblastresults", sampledblastresultsname = "pt2.middlebottomsampledblastresults", save.file = T, directorysave = outputdatabasedirectory, rdatablastname = "Midpoint-to-Stop Protein BLASTS List pt2.RData", save.csv = F, print.index = T, message.start ="STARTED MIDDLE BOTTOM BLASTS pt2 -", message.end = "FINISHED MIDDLE BOTTOM BLASTS pt2 -")

middlebottomblastoutput.pt3<-proteinBLAST.function(queryproteinsstringset = gene.pt3, querydescriptives = genename.pt3, make.database = F, load.database = T, proteindatabase = fulllengthproteindatabase, query.species = "Saccharomyces cerevisiae", proteinspan = "midpoint to stop", number.of.proteins = length(gene.pt3), bitscore.threshold = 50, rawblastresultsname = "pt3.middlebottomrawblastresults", sampledblastresultsname = "pt3.middlebottomsampledblastresults", save.file = T, directorysave = outputdatabasedirectory, rdatablastname = "Midpoint-to-Stop Protein BLASTS List pt3.RData", save.csv = F, print.index = T, message.start ="STARTED MIDDLE BOTTOM BLASTS pt3 -", message.end = "FINISHED MIDDLE BOTTOM BLASTS pt3 -")

middlebottomblastoutput.pt4<-proteinBLAST.function(queryproteinsstringset = gene.pt4, querydescriptives = genename.pt4, make.database = F, load.database = T, proteindatabase = fulllengthproteindatabase, query.species= "Saccharomyces cerevisiae",proteinspan ="midpoint to stop", number.of.proteins = length(gene.pt4), bitscore.threshold = 50, rawblastresultsname = "pt4.middlebottomrawblastresults", sampledblastresultsname = "pt4.middlebottomsampledblastresults", save.file = T, directorysave = outputdatabasedirectory, rdatablastname = "Midpoint-to-Stop Protein BLASTS List pt4.RData", save.csv = F, print.index = T, message.start ="STARTED MIDDLE BOTTOM BLASTS pt4 -", message.end = "FINISHED MIDDLE BOTTOM BLASTS pt4 -")

middlebottomblastoutput.pt5<-proteinBLAST.function(queryproteinsstringset = gene.pt5, querydescriptives = genename.pt5, make.database = F, load.database = T, proteindatabase = fulllengthproteindatabase, query.species = "Saccharomyces cerevisiae", proteinspan = "midpoint to stop", number.of.proteins = length(gene.pt5), bitscore.threshold = 50, rawblastresultsname = "pt5.middlebottomrawblastresults", sampledblastresultsname = "pt5.middlebottomsampledblastresults", save.file = T, directorysave = outputdatabasedirectory, rdatablastname = "Midpoint-to-Stop Protein BLASTS List pt5.RData", save.csv = F, print.index = T, message.start ="STARTED MIDDLE BOTTOM BLASTS pt5 -", message.end = "FINISHED MIDDLE BOTTOM BLASTS pt5 -")

middlebottomblastoutput.pt6<-proteinBLAST.function(queryproteinsstringset = gene.pt6, querydescriptives = genename.pt6, make.database = F, load.database = T, proteindatabase = fulllengthproteindatabase, query.species = "Saccharomyces cerevisiae", proteinspan = "midpoint to stop", number.of.proteins = length(gene.pt6), bitscore.threshold = 50, rawblastresultsname = "pt6.middlebottomrawblastresults", sampledblastresultsname = "pt6.middlebottomsampledblastresults", save.file = T, directorysave = outputdatabasedirectory, rdatablastname = "Midpoint-to-Stop Protein BLASTS List pt6.RData", save.csv = F, print.index = T, message.start ="STARTED MIDDLE BOTTOM BLASTS pt6 -", message.end = "FINISHED MIDDLE BOTTOM BLASTS pt6 -")

```

###BLASTS of Start to Midpoint of Proteins.

```{r,echo=F,eval=F}

topmiddlequeryproteins.blast<-read.fasta(paste0(outputdatabasedirectory,"All Saccharomyces cerevisiae ORF from start to midpoint.fasta"), seqtype="AA")

topmiddlequeryproteinsstringset<-readAAStringSet(paste0(outputdatabasedirectory,"All Saccharomyces cerevisiae ORF from start to midpoint.fasta"), format = "fasta")

topmiddlequerydescriptives<-data.frame(gene.name="ZZZZZ",common.name="",annotation="ZZZZZ",stringset.name="ZZZZZ",Amino.Acids=9999999,Blank=1:length(topmiddlequeryproteins.blast))

generep<-1

while(generep<(nrow(topmiddlequerydescriptives)+1)){
  
  topmiddlequerydescriptives$gene.name[generep]<-getName(topmiddlequeryproteins.blast[generep])
  
  topmiddlequerydescriptives$common.name[generep]<-unlist(strsplit(unlist(getAnnot(topmiddlequeryproteins.blast[generep]))," ",fixed=T))[2]
  
  topmiddlequerydescriptives$annotation[generep]<-unlist(getAnnot(topmiddlequeryproteins.blast[generep]))
  
  topmiddlequerydescriptives$stringset.name[generep]<-c2s(s2c(c2s(getAnnot(topmiddlequeryproteins.blast[generep])))[-1])
  
  topmiddlequerydescriptives$Amino.Acids[generep]<-length(unlist(getSequence(topmiddlequeryproteins.blast[generep])))
  
  generep<-generep+1
}

gene.all<-topmiddlequeryproteinsstringset

genename.all<-topmiddlequerydescriptives

split1numba<-round(length(gene.all)/6)

split2numba<-split1numba+split1numba+1

split3numba<-split1numba+split2numba+1

split4numba<-split1numba+split3numba+1

split5numba<-split1numba+split4numba+1

split6numba<-length(gene.all)

gene.pt1<-gene.all[1:split1numba]

genename.pt1<-genename.all[1:split1numba,]
  
gene.pt2<-gene.all[(split1numba+1):split2numba]

genename.pt2<-genename.all[(split1numba+1):split2numba,]

gene.pt3<-gene.all[(split2numba+1):split3numba]

genename.pt3<-genename.all[(split2numba+1):split3numba,]

gene.pt4<-gene.all[(split3numba+1):split4numba]

genename.pt4<-genename.all[(split3numba+1):split4numba,]

gene.pt5<-gene.all[(split4numba+1):split5numba]

genename.pt5<-genename.all[(split4numba+1):split5numba,]

gene.pt6<-gene.all[(split5numba+1):split6numba]

genename.pt6<-genename.all[(split5numba+1):split6numba,]

###All BLASTS under every context will be run using the exact same subject database, so the "make.database" argument is set to False after the database has been initially created.

fulllengthproteindatabase<-"D:/Saccharomycotina_without_Saccharomyces.fasta"

topmiddleblastoutput.pt1<-proteinBLAST.function(queryproteinsstringset = gene.pt1, querydescriptives = genename.pt1, make.database = F, load.database = T, proteindatabase = fulllengthproteindatabase, query.species = "Saccharomyces cerevisiae", proteinspan = "start to midpoint", number.of.proteins = length(gene.pt1), bitscore.threshold = 50, rawblastresultsname = "pt1.topmiddlerawblastresults", sampledblastresultsname = "pt1.topmiddlesampledblastresults", save.file = T, directorysave = outputdatabasedirectory, rdatablastname = "Start-to-Midpoint Protein BLASTS List pt1.RData",save.csv = F, print.index = T, message.start ="FINISHED TOP MIDDLE BOTTOM BLASTS pt1 -",message.end = "FINISHED TOP MIDDLE BOTTOM BLASTS pt1 -")

topmiddleblastoutput.pt2<-proteinBLAST.function(queryproteinsstringset = gene.pt2, querydescriptives = genename.pt2, make.database = F, load.database = T, proteindatabase = fulllengthproteindatabase, query.species = "Saccharomyces cerevisiae", proteinspan = "start to midpoint", number.of.proteins = length(gene.pt2), bitscore.threshold = 50, rawblastresultsname = "pt2.topmiddlerawblastresults", sampledblastresultsname = "pt2.topmiddlesampledblastresults", save.file = T, directorysave = outputdatabasedirectory, rdatablastname = "Start-to-Midpoint Protein BLASTS List pt2.RData", save.csv = F, print.index = T, message.start ="FINISHED TOP MIDDLE BOTTOM BLASTS pt2 -", message.end="FINISHED TOP MIDDLE BOTTOM BLASTS pt2 -")

topmiddleblastoutput.pt3<-proteinBLAST.function(queryproteinsstringset = gene.pt3, querydescriptives = genename.pt3, make.database =F , load.database = T, proteindatabase = fulllengthproteindatabase, query.species = "Saccharomyces cerevisiae", proteinspan = "start to midpoint", number.of.proteins = length(gene.pt3), bitscore.threshold = 50, rawblastresultsname = "pt3.topmiddlerawblastresults", sampledblastresultsname = "pt3.topmiddlesampledblastresults", save.file = T, directorysave = outputdatabasedirectory, rdatablastname = "Start-to-Midpoint Protein BLASTS List pt3.RData", save.csv = F, print.index = T, message.start ="FINISHED TOP MIDDLE BOTTOM BLASTS pt3 -", message.end = "FINISHED TOP MIDDLE BOTTOM BLASTS pt3 -")

topmiddleblastoutput.pt4<-proteinBLAST.function(queryproteinsstringset = gene.pt4, querydescriptives = genename.pt4, make.database = F, load.database = T, proteindatabase = fulllengthproteindatabase, query.species = "Saccharomyces cerevisiae", proteinspan = "start to midpoint", number.of.proteins = length(gene.pt4), bitscore.threshold = 50, rawblastresultsname = "pt4.topmiddlerawblastresults", sampledblastresultsname = "pt4.topmiddlesampledblastresults", save.file = T, directorysave = outputdatabasedirectory, rdatablastname = "Start-to-Midpoint Protein BLASTS List pt4.RData", save.csv = F, print.index = T, message.start ="FINISHED TOP MIDDLE BOTTOM BLASTS pt4 -", message.end = "FINISHED TOP MIDDLE BOTTOM BLASTS pt4 -")

topmiddleblastoutput.pt5<-proteinBLAST.function(queryproteinsstringset = gene.pt5, querydescriptives = genename.pt5, make.database = F, load.database = T, proteindatabase = fulllengthproteindatabase, query.species = "Saccharomyces cerevisiae", proteinspan = "start to midpoint", number.of.proteins = length(gene.pt5), bitscore.threshold = 50, rawblastresultsname = "pt5.topmiddlerawblastresults", sampledblastresultsname = "pt5.topmiddlesampledblastresults", save.file = T, directorysave = outputdatabasedirectory, rdatablastname = "Start-to-Midpoint Protein BLASTS List pt5.RData", save.csv = F, print.index = T, message.start ="FINISHED TOP MIDDLE BOTTOM BLASTS pt5 -", message.end = "FINISHED TOP MIDDLE BOTTOM BLASTS pt5 -")

topmiddleblastoutput.pt6<-proteinBLAST.function(queryproteinsstringset = gene.pt6, querydescriptives = genename.pt6, make.database = F, load.database = T, proteindatabase = fulllengthproteindatabase, query.species = "Saccharomyces cerevisiae", proteinspan = "start to midpoint", number.of.proteins = length(gene.pt6), bitscore.threshold = 50, rawblastresultsname = "pt6.topmiddlerawblastresults", sampledblastresultsname = "pt6.topmiddlesampledblastresults", save.file = T, directorysave = outputdatabasedirectory, rdatablastname = "Start-to-Midpoint Protein BLASTS List pt6.RData", save.csv = F, print.index = T, message.start ="FINISHED TOP MIDDLE BOTTOM BLASTS pt6 -", message.end = "FINISHED TOP MIDDLE BOTTOM BLASTS pt6 -")

```

###Compiling all of the parts from BLASTS.

```{r,echo=F,eval=F}

###Compiling all of the parts from the Full Length BLAST

queryproteinfile<-paste0(inputdatabasedirectory,"orf_trans_R64-3-1_20210421.fasta")

fullqueryproteins.blast<-seqinr::read.fasta(queryproteinfile, seqtype = "AA")

fullquerydescriptives<-data.frame(gene.name="ZZZZZ",common.name="ZZZZZ",annotation="ZZZZZ",stringset.name="ZZZZZ",Amino.Acids=9999999,Blank=1:length(fullqueryproteins.blast))

generep<-1

while(generep<(nrow(fullquerydescriptives)+1)){
  
  fullquerydescriptives$gene.name[generep]<-getName(fullqueryproteins.blast[generep])
  
  fullquerydescriptives$common.name[generep]<-unlist(strsplit(unlist(getAnnot(fullqueryproteins.blast[generep]))," ",fixed=T))[2]
  
  fullquerydescriptives$annotation[generep]<-unlist(getAnnot(fullqueryproteins.blast[generep]))
  
  fullquerydescriptives$stringset.name[generep]<-c2s(s2c(c2s(getAnnot(fullqueryproteins.blast[generep])))[-1])
  
  fullquerydescriptives$Amino.Acids[generep]<-length(unlist(getSequence(fullqueryproteins.blast[generep])))-1
  
  generep<-generep+1
}

###BLAST are computationally intensive and will require more RAM than my computer can handle to avoid crashes due to running out of memory. Due to memory and time constraints, the protein list will be split into 6 parts and then the blasts will be combined after each part has successfully completed. In order for this to work, all of the parts have to be saved with the following format: "pt" followed by the numerical order of the subset; so if the first subset of sequences are queried, then it should be saved as ".....pt1.RData". Also, each subset should be saved with the exact same file name except for the numerical subset value. "blastfiledirectory" should be a string for the directory containing the subsets of the BLAST results. "blastpartfilename" should be a string for the file name (requires extension ".RData") for each subset of the BLAST results that were saved; this won't work if the file names are different not counting the numerical subset. Since the BLAST function saves the BLAST output as a list with two data frames (raw BLAST results and the BLAST statistics), "sampledindex" is the list index containing the BLAST statistics and "rawblastindex" is the index containing the raw BLAST results; use the default for both indexes unless you manually changed the function. "querydescriptives" must be a data frame containing the names, annotations, and other important features of the query sequences; the BLAST function refers to specific column names in the "querydescriptives" data frame, so refer to the example BLAST function to know exactly how to format the column names or else this code won't work. "rawblastresultsname" should be a string for the name of the data frame containing the raw BLAST results. "sampledblastresultsname" should be a string for the name of the data frame that contain statistics regarding each query protein BLAST results. "save.file" gives the option (True or False) to save the BLAST output as an R file with the extension ".RData". If you choose to save the file using this code, then "rdatablastname" should be a string for the file name (requires extension ".RData") of the output and "directorysave" should be a string for the directory to save the file. "description" is an optional string that will be printed after the compilation has been completed. "message.start" is the message for the time elapse at the start of the function. "message.end" is the message for the time elapse after the function has completed.

fullblastresultscomp<-compilationBLAST.function(blastfiledirectory = outputdatabasedirectory, blastpartfilename = "Full Protein BLASTS List", sampledindex = 2, querydescriptives = fullquerydescriptives, rawblastindex = 1, rawblastresultsname = "comp.fullrawblastresults",  sampledblastresultsname = "comp.fullsampledblastresults", save.file = T, directorysave = outputdatabasedirectory, rdatablastname = "Full Protein BLASTS List compilation.RData", description = "FULL LENGTH", message.start = "Started full compilation-", message.end = "Finished full compilation -")

###Compiling all of the parts from the Midpoint-to-Stop BLASTS

remove(fullblastresultscomp)

middlebottomqueryproteins.blast<-read.fasta(paste0(outputdatabasedirectory,"All Saccharomyces cerevisiae ORF from midpoint to stop.fasta"), seqtype="AA")

middlebottomquerydescriptives<-data.frame(gene.name="ZZZZZ",common.name="ZZZZZ",annotation="ZZZZZ",stringset.name="ZZZZZ",Amino.Acids=9999999,Blank=1:length(middlebottomqueryproteins.blast))

generep<-1

while(generep<(nrow(middlebottomquerydescriptives)+1)){
  middlebottomquerydescriptives$gene.name[generep]<-getName(middlebottomqueryproteins.blast[generep])
  
  middlebottomquerydescriptives$common.name[generep]<-unlist(strsplit(unlist(getAnnot(middlebottomqueryproteins.blast[generep]))," ",fixed=T))[2]
  
  middlebottomquerydescriptives$annotation[generep]<-unlist(getAnnot(middlebottomqueryproteins.blast[generep]))
  
  middlebottomquerydescriptives$stringset.name[generep]<-c2s(s2c(c2s(getAnnot(middlebottomqueryproteins.blast[generep])))[-1])
  
  middlebottomquerydescriptives$Amino.Acids[generep]<-length(unlist(getSequence(middlebottomqueryproteins.blast[generep])))-1
  
  generep<-generep+1
}

middlebottomblastresultscomp<-compilationBLAST.function(blastfiledirectory = outputdatabasedirectory, blastpartfilename = "Midpoint-to-Stop Protein BLASTS List", sampledindex = 2,  querydescriptives = middlebottomquerydescriptives, rawblastindex = 1, rawblastresultsname = "comp.middlebottomrawblastresults",  sampledblastresultsname = "comp.middlebottomsampledblastresults", save.file = T, directorysave = outputdatabasedirectory, rdatablastname = "Midpoint-to-Stop Protein BLASTS List compilation.RData",  description = "MIDDLE BOTTOM", message.start = "Started Midpoint-to-Stop compilation-", message.end = "Finished Midpoint-to-Stop compilation -")

###Compiling all of the parts from the Start-to-Midpoint BLASTS

remove(middlebottomblastresultscomp)

topmiddlequeryproteins.blast<-read.fasta(paste0(outputdatabasedirectory,"All Saccharomyces cerevisiae ORF from start to midpoint.fasta"), seqtype="AA")

topmiddlequerydescriptives<-data.frame(gene.name="ZZZZZ",common.name="ZZZZZ",annotation="ZZZZZ",stringset.name="ZZZZZ",Amino.Acids=9999999,Blank=1:length(topmiddlequeryproteins.blast))

generep<-1

while(generep<(nrow(topmiddlequerydescriptives)+1)){
  topmiddlequerydescriptives$gene.name[generep]<-getName(topmiddlequeryproteins.blast[generep])
  
  topmiddlequerydescriptives$common.name[generep]<-unlist(strsplit(unlist(getAnnot(topmiddlequeryproteins.blast[generep]))," ",fixed=T))[2]
  
  topmiddlequerydescriptives$annotation[generep]<-unlist(getAnnot(topmiddlequeryproteins.blast[generep]))
  
  topmiddlequerydescriptives$stringset.name[generep]<-c2s(s2c(c2s(getAnnot(topmiddlequeryproteins.blast[generep])))[-1])
  
  topmiddlequerydescriptives$Amino.Acids[generep]<-length(unlist(getSequence(topmiddlequeryproteins.blast[generep])))
  
  generep<-generep+1
}

topmiddleblastresultscomp<-compilationBLAST.function(blastfiledirectory = outputdatabasedirectory, blastpartfilename = "Start-to-Midpoint Protein BLASTS List", sampledindex = 2,  querydescriptives = topmiddlequerydescriptives, rawblastindex = 1, rawblastresultsname = "comp.topmiddlerawblastresults",  sampledblastresultsname = "comp.topmiddlesampledblastresults", save.file = T, directorysave = outputdatabasedirectory, rdatablastname = "Start-to-Midpoint Protein BLASTS List compilation.RData",  description = "TOP MIDDLE", message.start = "Started Start-to-Midpoint compilation-", message.end = "Finished Start-to-Midpoint compilation -")

remove(topmiddleblastresultscomp)

```

###Adding species to BLAST results.

```{r,echo=F,eval=F}

###The default NCBI tool has no way of determining the species of the sequences from the subject database. This custom function will extract the species from the subject annotations accompanied with each sequence alignment. Also, for each query sequence, this function will further curate the BLAST results by selecting the alignment with the highest bitscore from each unique species from the subject database; this will eliminate duplicate BLAST hits that a query has for multiple proteins based on partial sequence similarities. "data" is the data frame containing the raw BLAST results (this is the compilation file if the query sequences were subset). "blastslist" should be the data frame containing the statistics from the BLAST results. "protein.database.analysis" gives the option (True or False) to determine unique species ONLY among the subject database NOT among the BLAST results. "query.species" is an optional string for the species that the query sequences originated from. "description" is an optional string that contains further details pertaining to the specific type of analysis being done. "save.file" gives the option (True or False) to save the BLAST output as an R file with the extension ".RData" as well as several csv files. If "protein.database.analysis" and "save.file" are both set to True, then "curatedblastsname" should be a string for the name of the csv file containing the parsed species from each entry of the subject database. "exportblastcsv" gives the option (True or False) to export the the BLAST results that are curated to only contain sequence alignments with the highest bitscore for each query BLAST; this is turned off by default because the file can be so large that it causes R to crash when trying to save. If "exportblastcsv" is set to True, then "uniquecuratedblastsname" should be a string for the name of the csv file that contains all BLASTS hits from unique organisms (regardless of bitscore) for each query. If "save.file" is set to True, then "countuniquesubjectname" should be a string for the name of the csv file detailing the number of BLAST hits that each unique species has among the sequence alignments; "countuniquequeryname" should be a string for the name of the csv file detailing the number of BLAST hits from unique species for each query gene. If "save.file" is set to True, then the curated BLAST results will be exported as a RData file consisting of a list of the following data.frames: "dataname" should be a string that will name the data frame consisting of all of the BLAST results initially input as "data" except that now, "dataname" has the parsed species for all of the BLAST alignments; "sampledblastresultsname" should be a string that will name the data frame containing the statistics of the BLAST results which is the same as what was input as "blastslist"; "uniqueorganismname" should be a string that will name the data frame that has the curated BLAST results with each query having a single alignment from each species with the highest bitscore. "print.index" gives the option (True or False) to print the current row of the particular data frame being analyzed. "rdatablastname" should be a string for the file name (requires extension ".RData") of the output and "directorysave" should be a string for the directory to save the file. "message.start" is the message for the time elapse at the start of the function. "message.end" is the message for the time elapse after the function has completed.

fulllengthproteindatabase<-"D:/Saccharomycotina_without_Saccharomyces.fasta"

subjectdatabase<-seqinr::read.fasta(fulllengthproteindatabase,seqtype="AA")

proteindatabaseannotations<-data.frame(Subject.Annotation=unlist(getAnnot(subjectdatabase)))

remove(subjectdatabase)

blastsuniqueorganisms.subjectdatabase<-uniquespecies.function(data = proteindatabaseannotations, protein.database.analysis = T, query.species = "Saccharomyces cerevisiae", description = "Entire Subject Protein Database", save.file = T, curatedblastsname = "RAW DATA - Every Entry in Saccharomycotina without Saccharomyces protein database.csv", exportblastcsv = T, uniquecuratedblastsname = "CURATED - Every Entry in Saccharomycotina without Saccharomyces protein database.csv", countuniquesubjectname = "Every UNIQUE Organism in Saccharomycotina without Saccharomyces protein database.csv", print.index = T, directorysave = outputdatabasedirectory, message.start = "STARTED SUBJECT DATABASE UNIQUE ORGANSISMS -", message.end = "FINISHED SUBJECT DATABASE UNIQUE ORGANSISMS -")

remove(proteindatabaseannotations)

remove(blastsuniqueorganisms.subjectdatabase)

###Curating BLASTS for Full-Length Proteins

load(paste0(outputdatabasedirectory,"Full Protein BLASTS List compilation.RData"))

blastsuniqueorganisms.full<-uniquespecies.function(data = blastresultscomp$comp.fullrawblastresults, blastslist = blastresultscomp$comp.fullsampledblastresults, protein.database.analysis = F, query.species = "Saccharomyces cerevisiae", description = "Full Length BLASTS", save.file = T, exportblastcsv = T, uniquecuratedblastsname = "CURATED - Every Entry in Full Protein BLASTS.csv", countuniquesubjectname = "Every UNIQUE Organism in Full Protein BLASTS.csv", countuniquequeryname = "Unique Organisms Hits For Each Query in Full Protein BLASTS.csv", dataname = "comp.fullrawblastresults", sampledblastresultsname = "comp.fullsampledblastresults", uniqueorganismname = "comp.fulluniqueorganisms", rdatablastname="Full Protein BLASTS List curated compilation.RData", print.index = F, directorysave = outputdatabasedirectory, message.start = "STARTED FULL LENGTH BLASTS UNIQUE ORGANSISMS -", message.end = "FINISHED FULL LENGTH BLASTS UNIQUE ORGANSISMS -")

remove(blastresultscomp)

remove(blastsuniqueorganisms.full)

###Curating BLASTS for Start-to-Midpoint Proteins

load(paste0(outputdatabasedirectory,"Start-to-Midpoint Protein BLASTS List compilation.RData"))

blastsuniqueorganisms.topmiddle<-uniquespecies.function(data = blastresultscomp$comp.topmiddlerawblastresults, blastslist = blastresultscomp$comp.topmiddlesampledblastresults, protein.database.analysis = F, query.species = "Saccharomyces cerevisiae", description = "Start to Midpoint BLASTS", save.file = T, exportblastcsv = T, uniquecuratedblastsname = "CURATED - Every Entry in Start-to-Midpoint Protein BLASTS.csv", countuniquesubjectname = "Every UNIQUE Organism in Start-to-Midpoint Protein BLASTS.csv", countuniquequeryname = "Unique Organisms Hits For Each Query in Start-to-Midpoint Protein BLASTS.csv", dataname = "comp.topmiddlerawblastresults", sampledblastresultsname = "comp.topmiddlesampledblastresults", uniqueorganismname = "comp.topmiddleuniqueorganisms", rdatablastname = "Start-to-Midpoint Protein BLASTS List curated compilation.RData", print.index = F, directorysave = outputdatabasedirectory, message.start = "STARTED TOP MIDDLE BLASTS UNIQUE ORGANSISMS -", message.end = "FINISHED TOP MIDDLE BLASTS UNIQUE ORGANSISMS -")

remove(blastresultscomp)

remove(blastsuniqueorganisms.topmiddle)

###Curating BLASTS for Midpoint-to-End Proteins

load(paste0(outputdatabasedirectory,"Midpoint-to-Stop Protein BLASTS List compilation.RData"))

blastsuniqueorganisms.middlebottom<-uniquespecies.function(data = blastresultscomp$comp.middlebottomrawblastresults, blastslist = blastresultscomp$comp.middlebottomsampledblastresults, protein.database.analysis = F, query.species = "Saccharomyces cerevisiae", description = "Midpoint to Stop BLASTS", save.file = T, exportblastcsv = T, uniquecuratedblastsname = "CURATED - Every Entry in Midpoint-to-Stop Protein BLASTS.csv", countuniquesubjectname = "Every UNIQUE Organism in Midpoint-to-Stop Protein BLASTS.csv", countuniquequeryname = "Unique Organisms Hits For Each Query in Midpoint-to-Stop Protein BLASTS.csv", dataname = "comp.middlebottomrawblastresults", sampledblastresultsname = "comp.middlebottomsampledblastresults", uniqueorganismname = "comp.middlebottomuniqueorganisms", rdatablastname = "Midpoint-to-Stop Protein BLASTS List curated compilation.RData", print.index = F, directorysave = outputdatabasedirectory, message.start = "STARTED MIDDLE BOTTOM BLASTS UNIQUE ORGANSISMS -", message.end = "FINISHED MIDDLE BOTTOM BLASTS UNIQUE ORGANSISMS -")

remove(blastresultscomp)

remove(blastsuniqueorganisms.middlebottom)

```

###Protein conservation scores analysis and output.

```{r,echo=F,eval=F}

###Protein conservation scores are calculated using a weighted algorithm."curatedblastsdata" should be the data frame that has the curated BLAST results with each query having a single alignment from each species with the highest bitscore. "sampleddata" should be a data frame that has the statistics from the BLAST results. "querystart" should be a string ("Beginning", "Middle", or "End") identifying the region to start the protein conservation score calculations; default is "Beginning" which is the start of the query proteins. "homologystart" should be the region that the BLAST algorithm determines where the alignment begins in the query or subject sequences; the default is "qstart" which is a numerical value that indicates the start of the alignment within the query. "analysis" is an optional string that refers to the length of amino acids spanning the conservation score as well as the region (beginning, middle or end). "query.species" is an optional string for the species that the query sequences originated from. "querydescriptives" must be a data frame containing the names, annotations, and other important features of the query sequences; the BLAST function refers to specific column names in the "querydescriptives" data frame, so refer to the example BLAST function to know exactly how to format the column names or else this code won't work. "bitscore.threshold" is the minimum bitscore value from each sequence alignment that will be included in the protein score calculations. "rampzonelength" is the length of amino acids used as the basis for homology, and is the length for the range of values in the applied as a weight to the conservation scores; for example, a "rampzonelength" of 40 means that the code will count how many unique species have a qstart (or sstart, or qend, etc.) at every position from 1-40, with the first amino acid having the greatest weight of 40, and the last (fortieth) amino acid having the smallest weight of 1. "print.index" gives the option (True or False) to print the current numerical index that is being analyzed; for example, an index of 5 means that the fifth protein in the "querydescriptives" data frame is currently being processed by the code. "description" is an optional string that contains further details pertaining to the specific type of analysis being done. "message.start" is the message for the time elapse at the start of the function. "message.end" is the message for the time elapse after the function has completed.

###Protein conservation scores for Full-length BLASTS

load(paste0(outputdatabasedirectory,"Full Protein BLASTS List curated compilation.RData"))

queryproteinfile<-paste0(inputdatabasedirectory,"orf_trans_R64-3-1_20210421.fasta")

fullqueryproteins.blast<-seqinr::read.fasta(queryproteinfile, seqtype = "AA")

fullquerydescriptives<-data.frame(gene.name="ZZZZZ",common.name="ZZZZZ",annotation="ZZZZZ",stringset.name="ZZZZZ",Amino.Acids=9999999,Blank=1:length(fullqueryproteins.blast))

generep<-1

while(generep<(nrow(fullquerydescriptives)+1)){
  
  fullquerydescriptives$gene.name[generep]<-getName(fullqueryproteins.blast[generep])
  
  fullquerydescriptives$common.name[generep]<-unlist(strsplit(unlist(getAnnot(fullqueryproteins.blast[generep]))," ",fixed=T))[2]
  
  fullquerydescriptives$annotation[generep]<-unlist(getAnnot(fullqueryproteins.blast[generep]))
  
  fullquerydescriptives$stringset.name[generep]<-c2s(s2c(c2s(getAnnot(fullqueryproteins.blast[generep])))[-1])
  
  fullquerydescriptives$Amino.Acids[generep]<-length(unlist(getSequence(fullqueryproteins.blast[generep])))-1
  
  generep<-generep+1
}

conservationscores.fullproteins.qstart.beginning<-weightedproportion.function(curatedblastsdata = curatedBLASTSresults$comp.fulluniqueorganisms, sampleddata = curatedBLASTSresults$comp.fullsampledblastresults, querystart = "Beginning", homologystart = "qstart", analysis = "First 40 Amino Acids", query.species = "Saccharomyces cerevisiae", querydescriptives = fullquerydescriptives, bitscore.threshold = 50, rampzonelength = 40, description = "Qstart Beginning Full Length Proteins", print.index = T, message.start = "Started qstart beginning full -", message.end = "qstart BEGINNING FULL PROTEIN CONSERVATION SCORES FINISHED -")

conservationscores.fullproteins.sstart.beginning<-weightedproportion.function(curatedblastsdata = curatedBLASTSresults$comp.fulluniqueorganisms, sampleddata = curatedBLASTSresults$comp.fullsampledblastresults, querystart = "Beginning", homologystart = "sstart", analysis = "First 40 Amino Acids", query.species = "Saccharomyces cerevisiae", querydescriptives = fullquerydescriptives, bitscore.threshold = 50, rampzonelength = 40, description = "Sstart Beginning Full Length Proteins", print.index = F, message.start = "Started sstart beginning full", message.end = "sstart BEGINNING FULL PROTEIN CONSERVATION SCORES FINISHED -")

conservationscores.fullproteins.qstart.middle<-weightedproportion.function(curatedblastsdata = curatedBLASTSresults$comp.fulluniqueorganisms, sampleddata = curatedBLASTSresults$comp.fullsampledblastresults, querystart = "Middle", homologystart = "qstart", analysis = "Middle 40 Amino Acids", query.species = "Saccharomyces cerevisiae", querydescriptives = fullquerydescriptives, bitscore.threshold = 50, rampzonelength = 40, description = "Qstart Middle Full Length Proteins", print.index = F, message.start = "Started qstart middle full -", message.end = "qstart MIDDLE FULL PROTEIN CONSERVATION SCORES FINISHED -")

conservationscores.fullproteins.sstart.middle<-weightedproportion.function(curatedblastsdata = curatedBLASTSresults$comp.fulluniqueorganisms, sampleddata = curatedBLASTSresults$comp.fullsampledblastresults, querystart = "Middle", homologystart = "sstart", analysis = "Middle 40 Amino Acids", query.species = "Saccharomyces cerevisiae", querydescriptives = fullquerydescriptives, bitscore.threshold = 50, rampzonelength = 40, description = "Sstart Middle Full Length Proteins", print.index = F, message.start = "Started sstart middle full -", message.end = "sstart MIDDLE FULL PROTEIN CONSERVATION SCORES FINISHED -")

conservationscores.fullproteins.qend.end<-weightedproportion.function(curatedblastsdata = curatedBLASTSresults$comp.fulluniqueorganisms, sampleddata = curatedBLASTSresults$comp.fullsampledblastresults, querystart = "End", homologystart = "qend", analysis = "Last 40 Amino Acids", query.species = "Saccharomyces cerevisiae", querydescriptives = fullquerydescriptives, bitscore.threshold = 50, rampzonelength = 40, description = "Qend End Full Length Proteins", print.index = F, message.start = "Started qend beginning full -", message.end = "qend END FULL PROTEIN CONSERVATION SCORES FINISHED -")

conservationscores.fullproteins.send.end<-weightedproportion.function(curatedblastsdata = curatedBLASTSresults$comp.fulluniqueorganisms, sampleddata = curatedBLASTSresults$comp.fullsampledblastresults, querystart = "End", homologystart = "send", analysis = "Last 40 Amino Acids", query.species = "Saccharomyces cerevisiae", querydescriptives = fullquerydescriptives, bitscore.threshold = 50, rampzonelength = 40, description = "Send End Full Length Proteins", print.index = F, message.start = "Started send beginning full -", message.end = "send END FULL PROTEIN CONSERVATION SCORES FINISHED -")

###Protein conservation scores for Midpoint-to-Stop Protein BLASTS

remove(curatedBLASTSresults)

load(paste0(outputdatabasedirectory,"Midpoint-to-Stop Protein BLASTS List curated compilation.RData"))

middlebottomqueryproteins.blast<-read.fasta(paste0(outputdatabasedirectory,"All Saccharomyces cerevisiae ORF from midpoint to stop.fasta"), seqtype="AA")

middlebottomquerydescriptives<-data.frame(gene.name="ZZZZZ",common.name="ZZZZZ",annotation="ZZZZZ",stringset.name="ZZZZZ",Amino.Acids=9999999,Blank=1:length(middlebottomqueryproteins.blast))

generep<-1

while(generep<(nrow(middlebottomquerydescriptives)+1)){
  
  middlebottomquerydescriptives$gene.name[generep]<-getName(middlebottomqueryproteins.blast[generep])
  
  middlebottomquerydescriptives$common.name[generep]<-unlist(strsplit(unlist(getAnnot(middlebottomqueryproteins.blast[generep]))," ",fixed=T))[2]
  
  middlebottomquerydescriptives$annotation[generep]<-unlist(getAnnot(middlebottomqueryproteins.blast[generep]))
  
  middlebottomquerydescriptives$stringset.name[generep]<-c2s(s2c(c2s(getAnnot(middlebottomqueryproteins.blast[generep])))[-1])
  
  middlebottomquerydescriptives$Amino.Acids[generep]<-length(unlist(getSequence(middlebottomqueryproteins.blast[generep])))-1
  
  generep<-generep+1
}

conservationscores.middlebottomproteins.qstart.beginning<-weightedproportion.function(curatedblastsdata = curatedBLASTSresults$comp.middlebottomuniqueorganisms, sampleddata = curatedBLASTSresults$comp.middlebottomsampledblastresults, querystart = "Beginning", homologystart = "qstart", analysis = "First 40 Amino Acids", query.species = "Saccharomyces cerevisiae", querydescriptives = middlebottomquerydescriptives, bitscore.threshold = 50, rampzonelength = 40, description = "Qstart Beginning Midpoint to Stop Proteins", print.index = F, message.start = "Started qstart beginning middlebottom -", message.end = "qstart BEGINNING middlebottom PROTEIN CONSERVATION SCORES FINISHED -")

conservationscores.middlebottomproteins.qend.end<-weightedproportion.function(curatedblastsdata = curatedBLASTSresults$comp.middlebottomuniqueorganisms, sampleddata = curatedBLASTSresults$comp.middlebottomsampledblastresults, querystart = "End", homologystart = "qend", analysis = "Last 40 Amino Acids", query.species = "Saccharomyces cerevisiae", querydescriptives= middlebottomquerydescriptives, bitscore.threshold = 50, rampzonelength = 40, description = "Qend End Midpoint to Stop Proteins", print.index = F, message.start = "Started qend beginning middlebottom -", message.end = "qend END middlebottom PROTEIN CONSERVATION SCORES FINISHED -")

conservationscores.middlebottomproteins.sstart.beginning<-weightedproportion.function(curatedblastsdata = curatedBLASTSresults$comp.middlebottomuniqueorganisms, sampleddata = curatedBLASTSresults$comp.middlebottomsampledblastresults, querystart = "Beginning", homologystart = "sstart", analysis = "First 40 Amino Acids", query.species = "Saccharomyces cerevisiae", querydescriptives = middlebottomquerydescriptives, bitscore.threshold = 50, rampzonelength = 40, description = "Sstart Beginning Midpoint to Stop Proteins", print.index = F, message.start = "Started sstart beginning middlebottom -", message.end = "sstart BEGINNING middlebottom PROTEIN CONSERVATION SCORES FINISHED -")

conservationscores.middlebottomproteins.send.end<-weightedproportion.function(curatedblastsdata = curatedBLASTSresults$comp.middlebottomuniqueorganisms, sampleddata = curatedBLASTSresults$comp.middlebottomsampledblastresults, querystart = "End", homologystart = "send", analysis = "Last 40 Amino Acids", query.species = "Saccharomyces cerevisiae", querydescriptives = middlebottomquerydescriptives, bitscore.threshold = 50, rampzonelength = 40, description = "Send End Midpoint to Stop Proteins", print.index = F, message.start = "Started send beginning middlebottom -", message.end = "send END middlebottom PROTEIN CONSERVATION SCORES FINISHED -")

###Protein conservation scores for Start-to-Midpoint Protein BLASTS

remove(curatedBLASTSresults)

load(paste0(outputdatabasedirectory,"Start-to-Midpoint Protein BLASTS List curated compilation.RData"))

topmiddlequeryproteins.blast<-read.fasta(paste0(outputdatabasedirectory,"All Saccharomyces cerevisiae ORF from start to midpoint.fasta"), seqtype="AA")

topmiddlequerydescriptives<-data.frame(gene.name="ZZZZZ",common.name="ZZZZZ",annotation="ZZZZZ",stringset.name="ZZZZZ",Amino.Acids=9999999,Blank=1:length(topmiddlequeryproteins.blast))

generep<-1

while(generep<(nrow(topmiddlequerydescriptives)+1)){
  
  topmiddlequerydescriptives$gene.name[generep]<-getName(topmiddlequeryproteins.blast[generep])
  
  topmiddlequerydescriptives$common.name[generep]<-unlist(strsplit(unlist(getAnnot(topmiddlequeryproteins.blast[generep]))," ",fixed=T))[2]
  
  topmiddlequerydescriptives$annotation[generep]<-unlist(getAnnot(topmiddlequeryproteins.blast[generep]))
  
  topmiddlequerydescriptives$stringset.name[generep]<-c2s(s2c(c2s(getAnnot(topmiddlequeryproteins.blast[generep])))[-1])
  
  topmiddlequerydescriptives$Amino.Acids[generep]<-length(unlist(getSequence(topmiddlequeryproteins.blast[generep])))
  
  generep<-generep+1
}

conservationscores.topmiddleproteins.qstart.beginning<-weightedproportion.function(curatedblastsdata = curatedBLASTSresults$comp.topmiddleuniqueorganisms, sampleddata = curatedBLASTSresults$comp.topmiddlesampledblastresults, querystart = "Beginning", homologystart = "qstart", analysis = "First 40 Amino Acids", query.species = "Saccharomyces cerevisiae", querydescriptives = topmiddlequerydescriptives, bitscore.threshold = 50, rampzonelength = 40, description = "Qstart Beginning Start to Midpoint Proteins", print.index = F, message.start = "Started qstart beginning topmiddle -", message.end = "qstart BEGINNING topmiddle PROTEIN CONSERVATION SCORES FINISHED -")

conservationscores.topmiddleproteins.qend.end<-weightedproportion.function(curatedblastsdata = curatedBLASTSresults$comp.topmiddleuniqueorganisms, sampleddata = curatedBLASTSresults$comp.topmiddlesampledblastresults, querystart = "End", homologystart = "qend", analysis = "Last 40 Amino Acids", query.species = "Saccharomyces cerevisiae", querydescriptives = topmiddlequerydescriptives, bitscore.threshold = 50, rampzonelength = 40, description = "Qend End Start to Midpoint Proteins", print.index = F, message.start = "Started qend beginning topmiddle -", message.end = "qend MIDDLE topmiddle PROTEIN CONSERVATION SCORES FINISHED -")

conservationscores.topmiddleproteins.sstart.beginning<-weightedproportion.function(curatedblastsdata = curatedBLASTSresults$comp.topmiddleuniqueorganisms, sampleddata = curatedBLASTSresults$comp.topmiddlesampledblastresults, querystart = "Beginning", homologystart = "sstart", analysis = "First 40 Amino Acids", query.species = "Saccharomyces cerevisiae", querydescriptives = topmiddlequerydescriptives, bitscore.threshold = 50, rampzonelength = 40, description = "Sstart Beginning Start to Midpoint Proteins", print.index = F, message.start = "Started sstart beginning topmiddle -", message.end = "sstart BEGINNING topmiddle PROTEIN CONSERVATION SCORES FINISHED -")

conservationscores.topmiddleproteins.send.end<-weightedproportion.function(curatedblastsdata = curatedBLASTSresults$comp.topmiddleuniqueorganisms, sampleddata = curatedBLASTSresults$comp.topmiddlesampledblastresults, querystart = "End", homologystart = "send", analysis = "Last 40 Amino Acids", query.species = "Saccharomyces cerevisiae", querydescriptives = topmiddlequerydescriptives, bitscore.threshold = 50, rampzonelength = 40, description = "Send End Start to Midpoint Proteins", print.index = F, message.start = "Started send beginning topmiddle -", message.end = "send MIDDLE topmiddle PROTEIN CONSERVATION SCORES FINISHED -")

remove(curatedBLASTSresults)

###Exporting protein conservation scores as excel file.

wb<-createWorkbook()

addWorksheet(wb, "Full qstart beginning")

addWorksheet(wb, "Full sstart beginning")

addWorksheet(wb, "Full qstart middle")

addWorksheet(wb, "Full sstart middle")

addWorksheet(wb, "Full qend end")

addWorksheet(wb, "Full send end")

addWorksheet(wb, "Start-Mid qstart beginning")

addWorksheet(wb, "Start-Mid sstart beginning")

addWorksheet(wb, "Start-Mid qend end")

addWorksheet(wb, "Start-Mid send end")

addWorksheet(wb, "Mid-Stop qstart beginning")

addWorksheet(wb, "Mid-Stop sstart beginning")

addWorksheet(wb, "Mid-Stop qend end")

addWorksheet(wb, "Mid-Stop send end")

writeData(wb,sheet="Full qstart beginning",x=conservationscores.fullproteins.qstart.beginning)

writeData(wb,sheet="Full sstart beginning",x=conservationscores.fullproteins.sstart.beginning)

writeData(wb,sheet="Full qstart middle",x=conservationscores.fullproteins.qstart.middle)

writeData(wb,sheet="Full sstart middle",x=conservationscores.fullproteins.sstart.middle)

writeData(wb,sheet="Full qend end",x=conservationscores.fullproteins.qend.end)

writeData(wb,sheet="Full send end",x=conservationscores.fullproteins.send.end)

writeData(wb,sheet="Start-Mid qstart beginning",x=conservationscores.topmiddleproteins.qstart.beginning)

writeData(wb,sheet="Start-Mid sstart beginning",x=conservationscores.topmiddleproteins.sstart.beginning)

writeData(wb,sheet="Start-Mid qend end",x=conservationscores.topmiddleproteins.qend.end)

writeData(wb,sheet="Start-Mid send end",x=conservationscores.topmiddleproteins.send.end)

writeData(wb,sheet="Mid-Stop qstart beginning",x=conservationscores.middlebottomproteins.qstart.beginning)

writeData(wb,sheet="Mid-Stop sstart beginning",x=conservationscores.middlebottomproteins.sstart.beginning)

writeData(wb,sheet="Mid-Stop qend end",x=conservationscores.middlebottomproteins.qend.end)

writeData(wb,sheet="Mid-Stop send end",x=conservationscores.middlebottomproteins.send.end)

saveWorkbook(wb, paste0(outputdatabasedirectory,"Protein Conservation Scores from Bitscore 50 BLASTS.xlsx"), overwrite = T)

```

####PART 3: FIGURES AND STATISTICS

###Loading files.

```{r,echo=F}

###Below will dictate the size of the figures after you knit the file. If this is an RMarkdown file, then you can knit and export all of the figures and statistics without changing anything, as long as you keep all of the file names the same as in the examples.

knitr::opts_chunk$set(fig.height=7.5, fig.width=12,fig.align='center')

###Begin time elapse for generating all the figures.

timebegin<-timelapsebegin.function(message.begin="STARTED FIGURES -")

print(timebegin[[7]])

yeastcodonusage<-data.frame(read_excel(paste0(outputdatabasedirectory,"Saccharomyces cerevisiae codon usage table.xlsx"),col_names = T),check.names = F,stringsAsFactors = F)

yeastcodonusage$`Amino Acid`<-as.factor(yeastcodonusage$`Amino Acid`)

###Short nucleotides (300 nt or shorter) will be omitted because they will be translated to proteins that are shorter than 100 amino acids which when split into half for the BLASTS may lead to misleading homology.

initialtranslationspeed<-data.frame(read_excel(paste0(outputdatabasedirectory,"Initial Translation Speed Table 40.xlsx"),col_names = T),check.names = F,stringsAsFactors = F)

initialtranslationspeed<-initialtranslationspeed[initialtranslationspeed$Nucleotides>300,]

initialtranslationspeed$inverserrt.beginningvsrest<-log2((1/(initialtranslationspeed$AverageRRTFirst40nostartcodon/initialtranslationspeed$AverageRRTentireminusfirst40)))

initialtranslationspeed$inverserrt.endvsrest<-log2((1/(initialtranslationspeed$AverageRRTThreePrime40/initialtranslationspeed$AverageRRTentireminuslast40)))

###These files are when ATG is "neutralized" by replacing ATG's natural RRT with the average RRT across all protein-coding region.

initialtranslationspeedneut<-data.frame(read_excel(paste0(outputdatabasedirectory,"Initial Translation Speed Table ATG Neutralized 40.xlsx"),col_names = T),check.names = F,stringsAsFactors = F)

initialtranslationspeedneut<-initialtranslationspeedneut[initialtranslationspeedneut$Nucleotides>300,]

initialtranslationspeedneut$inverserrt.beginningvsrest<-log2((1/(initialtranslationspeedneut$AverageRRTFirst40nostartcodon/initialtranslationspeedneut$AverageRRTentireminusfirst40)))

initialtranslationspeedneut$inverserrt.endvsrest<-log2((1/(initialtranslationspeedneut$AverageRRTThreePrime40/initialtranslationspeedneut$AverageRRTentireminuslast40)))

```

###Translation speed statistics at the N-termini and C-termini.

```{r,echo=F}

###First 30 codons

print(paste0("###First 30 codons"))

initialtranslationspeed30<-data.frame(read_excel(paste0(outputdatabasedirectory,"Initial Translation Speed Table 30.xlsx"),col_names = T),check.names = F,stringsAsFactors = F)

initialtranslationspeed30<-initialtranslationspeed30[initialtranslationspeed30$Nucleotides>300,]

print(paste0("mean(log2(first30RRT/restRRT))= ",signif(mean(initialtranslationspeed30$log2Ratio30nostartcodonvsRest),digits = 5)))

first30vsrestpvaluewt<-wilcox.test(initialtranslationspeed30$AverageRRTFirst30nostartcodon,initialtranslationspeed30$AverageRRTentireminusfirst30,alternative="two.sided",paired=F)[["p.value"]]

print(paste0("5' translation speed of first 30 codons vs. rest p= ",signif(first30vsrestpvaluewt,digits = 5)))

###First 40 codons

print(paste0("###First 40 codons"))

initialtranslationspeed40<-data.frame(read_excel(paste0(outputdatabasedirectory,"Initial Translation Speed Table 40.xlsx"),col_names = T),check.names = F,stringsAsFactors = F)

initialtranslationspeed40<-initialtranslationspeed40[initialtranslationspeed40$Nucleotides>300,]

print(paste0("mean(log2(first40RRT/restRRT))= ",signif(mean(initialtranslationspeed40$log2Ratio40nostartcodonvsRest),digits = 5)))

first40vsrestpvaluewt<-wilcox.test(initialtranslationspeed40$AverageRRTFirst40nostartcodon,initialtranslationspeed40$AverageRRTentireminusfirst40,alternative="two.sided",paired=F)[["p.value"]]

print(paste0("5' translation speed of first 40 codons vs. rest p= ",signif(first40vsrestpvaluewt,digits = 5)))

###First 50 codons

print(paste0("###First 50 codons"))

initialtranslationspeed50<-data.frame(read_excel(paste0(outputdatabasedirectory,"Initial Translation Speed Table 50.xlsx"),col_names = T),check.names = F,stringsAsFactors = F)

initialtranslationspeed50<-initialtranslationspeed50[initialtranslationspeed50$Nucleotides>300,]

print(paste0("mean(log2(first50RRT/restRRT))= ",signif(mean(initialtranslationspeed50$log2Ratio50nostartcodonvsRest),digits = 5)))

first50vsrestpvaluewt<-wilcox.test(initialtranslationspeed50$AverageRRTFirst50nostartcodon,initialtranslationspeed50$AverageRRTentireminusfirst50,alternative="two.sided",paired=F)[["p.value"]]

print(paste0("5' translation speed of first 50 codons vs. rest p= ",signif(first50vsrestpvaluewt,digits = 5)))

###First 60 codons

print(paste0("###First 60 codons"))

initialtranslationspeed60<-data.frame(read_excel(paste0(outputdatabasedirectory,"Initial Translation Speed Table 60.xlsx"),col_names = T),check.names = F,stringsAsFactors = F)

initialtranslationspeed60<-initialtranslationspeed60[initialtranslationspeed60$Nucleotides>300,]

print(paste0("mean(log2(first60RRT/restRRT))= ",signif(mean(initialtranslationspeed60$log2Ratio60nostartcodonvsRest),digits = 5)))

first60vsrestpvaluewt<-wilcox.test(initialtranslationspeed60$AverageRRTFirst60nostartcodon,initialtranslationspeed60$AverageRRTentireminusfirst60,alternative="two.sided",paired=F)[["p.value"]]

print(paste0("5' translation speed of first 60 codons vs. rest p= ",signif(first60vsrestpvaluewt,digits = 5)))

###First 70 codons

print(paste0("###First 70 codons"))

initialtranslationspeed70<-data.frame(read_excel(paste0(outputdatabasedirectory,"Initial Translation Speed Table 70.xlsx"),col_names = T),check.names = F,stringsAsFactors = F)

initialtranslationspeed70<-initialtranslationspeed70[initialtranslationspeed70$Nucleotides>300,]

print(paste0("mean(log2(first70RRT/restRRT))= ",signif(mean(initialtranslationspeed70$log2Ratio70nostartcodonvsRest),digits = 5)))

first70vsrestpvaluewt<-wilcox.test(initialtranslationspeed70$AverageRRTFirst70nostartcodon,initialtranslationspeed70$AverageRRTentireminusfirst70,alternative="two.sided",paired=F)[["p.value"]]

print(paste0("5' translation speed of first 70 codons vs. rest p= ",signif(first70vsrestpvaluewt,digits = 5)))

###First 80 codons

print(paste0("###First 80 codons"))

initialtranslationspeed80<-data.frame(read_excel(paste0(outputdatabasedirectory,"Initial Translation Speed Table 80.xlsx"),col_names = T),check.names = F,stringsAsFactors = F)

initialtranslationspeed80<-initialtranslationspeed80[initialtranslationspeed80$Nucleotides>300,]

print(paste0("mean(log2(first80RRT/restRRT))= ",signif(mean(initialtranslationspeed80$log2Ratio80nostartcodonvsRest),digits = 5)))

first80vsrestpvaluewt<-wilcox.test(initialtranslationspeed80$AverageRRTFirst80nostartcodon,initialtranslationspeed80$AverageRRTentireminusfirst80,alternative="two.sided",paired=F)[["p.value"]]

print(paste0("5' translation speed of first 80 codons vs. rest p= ",signif(first80vsrestpvaluewt,digits = 5)))

###First 90 codons

print(paste0("###First 90 codons"))

initialtranslationspeed90<-data.frame(read_excel(paste0(outputdatabasedirectory,"Initial Translation Speed Table 90.xlsx"),col_names = T),check.names = F,stringsAsFactors = F)

initialtranslationspeed90<-initialtranslationspeed90[initialtranslationspeed90$Nucleotides>300,]

print(paste0("mean(log2(first90RRT/restRRT))= ",signif(mean(initialtranslationspeed90$log2Ratio90nostartcodonvsRest),digits = 5)))

first90vsrestpvaluewt<-wilcox.test(initialtranslationspeed90$AverageRRTFirst90nostartcodon,initialtranslationspeed90$AverageRRTentireminusfirst90,alternative="two.sided",paired=F)[["p.value"]]

print(paste0("5' translation speed of first 90 codons vs. rest p= ",signif(first90vsrestpvaluewt,digits = 5)))

###First 100 codons

print(paste0("###First 100 codons"))

initialtranslationspeed100<-data.frame(read_excel(paste0(outputdatabasedirectory,"Initial Translation Speed Table 100.xlsx"),col_names = T),check.names = F,stringsAsFactors = F)

initialtranslationspeed100<-initialtranslationspeed100[initialtranslationspeed100$Nucleotides>300,]

print(paste0("mean(log2(first100RRT/restRRT))= ",signif(mean(initialtranslationspeed100$log2Ratio100nostartcodonvsRest),digits = 5)))

first100vsrestpvaluewt<-wilcox.test(initialtranslationspeed100$AverageRRTFirst100nostartcodon,initialtranslationspeed100$AverageRRTentireminusfirst100,alternative="two.sided",paired=F)[["p.value"]]

print(paste0("5' translation speed of first 100 codons vs. rest p= ",signif(first100vsrestpvaluewt,digits = 5)))

```

###Figure 1: Calculation of translation speed confirms slow initial translation (SIT).

```{r,echo=F}

###Figure 1: Calculation of translation speed confirms slow initial translation (SIT).

initialtranslationspeed<-data.frame(read_excel(paste0(outputdatabasedirectory,"Initial Translation Speed Table 40.xlsx"),col_names = T),check.names = F,stringsAsFactors = F)

initialtranslationspeed<-initialtranslationspeed[initialtranslationspeed$Nucleotides>300,]

initialtranslationspeed$inverserrt.beginningvsrest<-log2((1/(initialtranslationspeed$AverageRRTFirst40nostartcodon/initialtranslationspeed$AverageRRTentireminusfirst40)))

initialtranslationspeed$inverserrt.endvsrest<-log2((1/(initialtranslationspeed$AverageRRTThreePrime40/initialtranslationspeed$AverageRRTentireminuslast40)))

genetemp<-read.fasta(paste0(inputdatabasedirectory,"orf_coding_R64-3-1_20210421.fasta"))

gene<-genetemp[initialtranslationspeed$Name]

fiveprimecodonlist<-rep(list(NULL),(max(initialtranslationspeed$Nucleotides)/3)-1)

fiveprimerrtlist<-rep(list(NULL),(max(initialtranslationspeed$Nucleotides)/3)-1)

stopinframe<-NULL

ATGatstart<-NULL

ATGnotatstart<-NULL

generep<-1

while (generep<(length(gene)+1)){
  
  dnasequence<-getSequence(gene[[generep]])

  firstposition<-1
  
  indexxxx<-1

  while(firstposition<length(dnasequence)-3){
  
    fiveprimecodonlist[[indexxxx]]<-append(fiveprimecodonlist[[indexxxx]],toupper(c2s(dnasequence[firstposition:(firstposition+2)])))
    
    fiveprimerrtlist[[indexxxx]]<-append(fiveprimerrtlist[[indexxxx]],yeastcodonusage$RRT[yeastcodonusage$Codons==toupper(c2s(dnasequence[firstposition:(firstposition+2)]))])
      
    if(firstposition<121&toupper(c2s(dnasequence[firstposition:(firstposition+2)]))=="TAG"){
      
      stopinframe<-c(stopinframe,getName(gene[generep]))
    } else if(firstposition<121&toupper(c2s(dnasequence[firstposition:(firstposition+2)]))=="TAA"){
      
      stopinframe<-c(stopinframe,getName(gene[generep]))
    } else if(firstposition<121&toupper(c2s(dnasequence[firstposition:(firstposition+2)]))=="TGA"){
      
      stopinframe<-c(stopinframe,getName(gene[generep]))
    }
    
    if(firstposition==1&toupper(c2s(dnasequence[firstposition:(firstposition+2)]))=="ATG"){
    
    ATGatstart<-c(ATGatstart,getName(gene[generep]))
    } else if(firstposition==1&toupper(c2s(dnasequence[firstposition:(firstposition+2)]))!="ATG"){
    
    ATGnotatstart<-c(ATGnotatstart,getName(gene[generep]))
    }
    
    firstposition<-firstposition+3
    
    indexxxx<-indexxxx+1
  }

  generep<-generep+1
}

###There are 17 genes with in-frame stop codons within the first 40 codons. These are mitochondrial genes that are polycistronic.

genestopcodons40<-gene[stopinframe]

compiledcodonusageoutputframeone<-rep(list(NA),length(genestopcodons40))

generep<-1

while (generep<(length(genestopcodons40)+1)){
  
  dnasequence<-getSequence(genestopcodons40[[generep]])

  firstposition<-1
  
  indexxxx<-1
  
  fiveprimecodonlistframeone<-rep(NA,40)
  
  while (firstposition<121){
  
    fiveprimecodonlistframeone[indexxxx]<-toupper(c2s(dnasequence[firstposition:(firstposition+2)]))
  
    firstposition<-firstposition+3
  
    indexxxx<-indexxxx+1
  }

  compiledcodonusageoutputframeone[generep]<-list(fiveprimecodonlistframeone)
  
  names(compiledcodonusageoutputframeone)[generep] <- getName(genestopcodons40[[generep]])
  
  generep<-generep+1
}

averagerrteveryposition<-data.frame(codonposition=99999,count=99999,meanRRT=99999,index=1:length(fiveprimerrtlist))

generep<-1

while(generep<(length(fiveprimerrtlist)+1)){
  
  averagerrteveryposition$codonposition[generep]<-generep
  
  averagerrteveryposition$count[generep]<-length(unlist(fiveprimerrtlist[generep]))
  
  averagerrteveryposition$meanRRT[generep]<-mean(unlist(fiveprimerrtlist[generep]))
  
  generep<-generep+1
}

##The inverse of the average RRT will be plot to match Tuller's figure.

averagerrteveryposition$inversemeanrrt<-1/averagerrteveryposition$meanRRT

yeastcodonusage$cumRRT<-yeastcodonusage$`Frame 1 (Coding) Observed Counts`*yeastcodonusage$RRT

globalrrt<-sum(yeastcodonusage$cumRRT)/sum(yeastcodonusage$`Frame 1 (Coding) Observed Counts`)

###In order to match Tuller's figures, For each codon position the inverse of the average RRT will be plotted for the first 200 codons minus the start codon since every yeast gene has ATG as the first codon.

lineplot<-function(data){
  
  ggplot(data=data, aes(x=codonposition, y=inversemeanrrt))+
  
  geom_line()+
  
  geom_hline(yintercept = 1/globalrrt,linetype="dotted")+
  
  geom_vline(xintercept = 40,linetype="dotted")+
  
  labs(title="",x="Distance from Start Codon",y="1/RRT")+
    
  theme(panel.grid.major = element_line(size = 0.5, colour = 'black'),
        
        panel.grid.major.x = element_line(size = 0.5, colour = NA),
        
        panel.grid.major.y = element_line(size = 0.5, colour = NA),
        
        panel.grid.minor = element_line(colour = NA),
      
        plot.title = element_text(hjust = 0.5,size = 15),
      
        axis.title.x = element_text(size=37),
      
        axis.text.x = element_text(family="sans", vjust=0.9,hjust=0.5,size=30,angle=0),
      
        axis.text.y = element_text(size=30),
      
        axis.title.y = element_text(size=37),
      
        panel.background = element_rect(fill = 'white', colour = 'black'))+
    
  scale_y_continuous(breaks = scales::pretty_breaks(n = 8),limits = c(0.94,0.990))
}

data2<-averagerrteveryposition[2:200,]

lineplot(data = data2)

###Figure 1 statistics

print(paste0("###Figure 1 statistics"))

corr<-cor.test(averagerrteveryposition$inversemeanrrt[2:200], averagerrteveryposition$codonposition[2:200],method = "spearman")

corr

paste0("mean 5' first 40 RRT= ",signif(mean(initialtranslationspeed$AverageRRTFirst40nostartcodon),digits = 5))

paste0("mean 5' 41:end RRT= ",signif(mean(initialtranslationspeed$AverageRRTentireminusfirst40),digits = 5))

paste0("mean(log2(first40RRT/restRRT))= ",signif(mean(initialtranslationspeed$log2Ratio40nostartcodonvsRest),digits = 5))

first40vsrestpvaluewt<-wilcox.test(initialtranslationspeed$AverageRRTFirst40nostartcodon,initialtranslationspeed$AverageRRTentireminusfirst40,alternative="two.sided",paired=F)[["p.value"]]

paste0("5' translation speed of first 40 codons vs. rest p= ",signif(first40vsrestpvaluewt,digits = 5))

paste0("ATG neutralized mean 5' first 40 RRT== ",signif(mean(initialtranslationspeedneut$AverageRRTFirst40nostartcodon),digits = 5))

paste0("ATG neutralized mean 5' 41:end RRT= ",signif(mean(initialtranslationspeedneut$AverageRRTentireminusfirst40),digits = 5))

paste0("ATG neutralized mean(log2(first40RRT/restRRT))= ",signif(mean(initialtranslationspeedneut$log2Ratio40nostartcodonvsRest),digits = 5))

paste0("equation is  y= ",signif(lm(data2$inversemeanrrt~data2$codonposition)[["coefficients"]][[1]],digits=4)," + ", signif(lm(data2$inversemeanrrt~data2$codonposition)[["coefficients"]][[2]],digits=4),"x")

```

###Figure 2:  Codon usage in the Slow Initial Translation (SIT) region.

```{r,echo=F}

###Figure 2A: Codon usage in the Slow Initial Translation (SIT) region.

allorfcodonsfirst40<-unlist(fiveprimecodonlist[2:40])

allorfcodonsrest<-unlist(fiveprimecodonlist[41:length(fiveprimecodonlist)])

first40codons<-data.frame(table(allorfcodonsfirst40))

names(first40codons)[1]<-"Codons"

names(first40codons)[names(first40codons) == "Freq"] <- "Frequency"

first40codons<-join(first40codons,yeastcodonusage[,c("Codons","Amino Acid","RRT")],by="Codons",type="full", match="all")

restcodons<-data.frame(table(allorfcodonsrest))

names(restcodons)[1]<-"Codons"

names(restcodons)[names(restcodons) == "Freq"] <- "Frequency"

restcodons<-join(restcodons,yeastcodonusage[,c("Codons","Amino Acid","RRT")],by="Codons",type="full", match="all")

first40codons$first40Proportion<-first40codons$Frequency/sum(first40codons$Frequency,na.rm = T)

restcodons$restProportion<-restcodons$Frequency/sum(restcodons$Frequency)

first40vsrest<-join(first40codons,restcodons,by=c("Codons","Amino Acid","RRT"),type="full", match="all")

first40vsrest$beginningvsrestmeancodonfoldchange<-first40vsrest$first40Proportion/first40vsrest$restProportion

globalvssit<-join(yeastcodonusage,first40vsrest,by=c("Codons","Amino Acid","RRT"),match = "all",type = "full")

globalvssit$Codons<-factor(globalvssit$Codons)

globalvssit2<-globalvssit[order(globalvssit$`Codon Proportion`),]

globalvssit3<-globalvssit2[order(globalvssit2$`Amino Acid`),]

globalvssit3<-droplevels(globalvssit3[!globalvssit3$`Amino Acid`=="*",])

globalvssit3$newsymbol<-"ZZZZZ"

generep<-1

while(generep<nrow(globalvssit3)+1){
  
  globalvssit3$newsymbol[generep]<-paste0(globalvssit3$`Amino Acid`[generep],"-", globalvssit3$Codons[generep])
  
  generep<-generep+1
}

colorlist<-c(rep(c("black","gray50"),10))

aminocolors<-data.frame(AA=levels(globalvssit3$`Amino Acid`),colors=colorlist)

names(aminocolors)[names(aminocolors) == "AA"] <- "Amino Acid"

newdata<-join(globalvssit3,aminocolors,type="full",match = "all",by="Amino Acid")

newdata<-newdata[!newdata$`Amino Acid`=="*",]

colorlistfin<-newdata$colors

barcodonplot<-function(data,codonname){
  
  ggplot(data=data)+
    
  geom_col(data = data,aes(x = factor(newsymbol,levels =newsymbol ),y = beginningvsrestmeancodonfoldchange,fill=factor(newsymbol,levels =newsymbol )),width = .75, position = "dodge")+
    
  geom_hline(yintercept = 1,linetype="dashed")+
    
  labs(title="",x="Codons",y="Usage (2:40/Rest)")+
    
  theme(panel.grid.major = element_line(size = 0.5, colour = 'black'),
        
        panel.grid.major.x = element_line(size = 0.5, colour = NA),
        
        panel.grid.major.y = element_line(size = 0.5, colour = NA),
        
        panel.grid.minor = element_line(colour = NA),
        
        plot.title = element_text(hjust = 0.5,size = 13),
        
        axis.title.x = element_text(size=37),
        
        axis.text.x = element_text(family="sans", vjust=0.3,hjust=1,size=13,angle=90),
        
        axis.text.y = element_text(size=30),
        
        axis.title.y = element_text(size=30),
        
        panel.background = element_rect(fill = 'white', colour = 'black'),
        
        legend.position="none")+
    
  scale_y_continuous(expand=c(0,0),breaks = scales::pretty_breaks(n = 10), limits=c(0,(max(data$beginningvsrestmeancodonfoldchange)*1.2)))+
    
  scale_fill_manual(values=colorlistfin)
}

testname=paste("First 40 Codons no start codon")

barcodonplot(data=newdata,codonname = testname)

###Figure 2B: Absolute usage of each leucine codon in the SIT.

newdata<-first40codons[first40codons$`Amino Acid`=="L",]

names(newdata)[names(newdata) == "first40Proportion"] <- "Codon Proportion"

newdata$zone<-"first40"

newdata2<-yeastcodonusage[,c("Codons","Frame 1 (Coding) Observed Counts","Codon Proportion","Amino Acid","RRT")]

newdata2<-newdata2[newdata2$`Amino Acid`=="L",]

names(newdata2)[names(newdata2) == "Frame 1 (Coding) Observed Counts"] <- "Frequency"

newdata2$zone<-"global"

newdata3<-rbind(newdata,newdata2)

newdata3$Codons=ordered(newdata3$Codons, levels = c("TTA","TTG","CTA","CTT","CTG","CTC"))

newdata3$zone=ordered(newdata3$zone, levels = c("global","first40"))

barcodonplot<-function(data,codonname){
  ggplot(data=data)+
    
  geom_col(data = data,aes(x = Codons,y = `Codon Proportion`,fill=zone),width = .75, position = "dodge")+
    
  labs(title="",x="Leucine Codons",y="Usage")+
    
  theme(panel.grid.major = element_line(size = 0.5, colour = 'black'),
        
        panel.grid.major.x = element_line(size = 0.5, colour = NA),
        
        panel.grid.major.y = element_line(size = 0.5, colour = NA),
        
        panel.grid.minor = element_line(colour = NA),
        
        plot.title = element_text(hjust = 0.5,size = 15),
        
        axis.title.x = element_text(size=40),
        
        axis.text.x = element_text(family="sans", vjust=0.3,hjust=0.5,size=27,angle=90),
        
        axis.text.y = element_text(size=35),
        
        axis.title.y = element_text(size=45),
        
        legend.title = element_text(color = "black", size = 28),
        
        legend.text = element_text(size=28),
        
        panel.background = element_rect(fill = 'white', colour = 'black'),
        
        legend.key=element_rect(fill="white"))+
    
  scale_y_continuous(expand=c(0,0),breaks = c(0.01,0.02,0.03,0.04), limits=c(0,0.048))+
    
  scale_fill_manual(name="",breaks=c("global", "first40"),labels=c("Global", "First 40"),values = c("black", "gray50"))
}

barcodonplot(data=newdata3,codonname = testname)

```

###Figure 3: The N-termini of proteins can vary in evolution. Figure 3 was generated using the online NCBI BLAST tool. https://blast.ncbi.nlm.nih.gov/Blast.cgi?PROGRAM=blastp&PAGE_TYPE=BlastSearch&LINK_LOC=blasthome

###Figure 4: Conservation of S. cerevisiae proteins over the N-terminal, Middle, and C-terminal 40 amino acids.

```{r,echo=F}

###All proteins that have full lengths shorter than 100 amino acids (303 nucleotides) will be omitted. The reason is that short amino acids are likely to have partial homology with other proteins even if they are not genuine orthologues. Since BLASTs were done on proteins that were split into the top and bottom halves, half-lengthed proteins that are shorter than 50 nucleotides will be omitted from analyses. Also, We only considered conservation scores from proteins that had homology with at least 40 unique species in Saccharomycotina, and therefore all proteins that have fewer than 40 highly-conserved orthlogues will be omitted from all analyses. 

###Using protein conservation scores from first half of proteins.

topmiddlebeginning<-data.frame(read_excel(paste0(outputdatabasedirectory,"Protein Conservation Scores from Bitscore 50 BLASTS.xlsx"),sheet = "Start-Mid qstart beginning",col_names = T),check.names = F,stringsAsFactors = F)

topmiddlebeginning<-topmiddlebeginning[order(topmiddlebeginning$Name),]

topmiddlebeginning<-topmiddlebeginning[!topmiddlebeginning$Unique.Organisms.Bitscore.Threshold<40,]

topmiddlebeginning<-topmiddlebeginning[!topmiddlebeginning$Amino.Acids<50,]

topmiddlebeginning$topmiddle.protein.sumweightedscore<-topmiddlebeginning$Sum.Weighted.Proportion

topmiddlebeginning<-topmiddlebeginning[topmiddlebeginning$Total.BLASTS>0,]

topmiddlebeginning<-topmiddlebeginning[topmiddlebeginning$Details!="ALL BLASTS have bitscore lower than 50",]

topmiddlebeginning<-topmiddlebeginning[topmiddlebeginning$Details!="No BLASTS Results",]

load(paste0(outputdatabasedirectory,"Start-to-Midpoint Protein BLASTS List curated compilation.RData"))

highhomologysequences.all<-curatedBLASTSresults$comp.topmiddleuniqueorganisms[!curatedBLASTSresults$comp.topmiddleuniqueorganisms$bitscore<50,]

highhomologysequences.topmiddle<-highhomologysequences.all[highhomologysequences.all$qaccver%in%topmiddlebeginning$Name,]

print(paste0("BLASTs from the first half of proteins have ",nrow(highhomologysequences.topmiddle[!duplicated(highhomologysequences.topmiddle$Organism),]), " unique species with a bitscore of at least 50"))

print(paste0("BLAST of the first half of proteins have ",nrow(highhomologysequences.topmiddle)," sequences with a bitscore of at least 50"))

remove(curatedBLASTSresults,highhomologysequences.all)

###Using protein conservation scores from second half of proteins.

middlebottombeginning<-data.frame(read_excel(paste0(outputdatabasedirectory,"Protein Conservation Scores from Bitscore 50 BLASTS.xlsx"),sheet = "Mid-Stop qstart beginning",col_names = T),check.names = F,stringsAsFactors = F)

middlebottombeginning<-middlebottombeginning[order(middlebottombeginning$Name),]

middlebottombeginning<-middlebottombeginning[!middlebottombeginning$Unique.Organisms.Bitscore.Threshold<40,]

middlebottombeginning<-middlebottombeginning[!middlebottombeginning$Amino.Acids<50,]

middlebottombeginning$middlebottom.protein.sumweightedscore<-middlebottombeginning$Sum.Weighted.Proportion

middlebottombeginning<-middlebottombeginning[middlebottombeginning$Total.BLASTS>0,]

middlebottombeginning<-middlebottombeginning[middlebottombeginning$Details!="ALL BLASTS have bitscore lower than 50",]

middlebottombeginning<-middlebottombeginning[middlebottombeginning$Details!="No BLASTS Results",]

load(paste0(outputdatabasedirectory,"Midpoint-to-Stop Protein BLASTS List curated compilation.RData"))

highhomologysequences.all<-curatedBLASTSresults$comp.middlebottomuniqueorganisms[!curatedBLASTSresults$comp.middlebottomuniqueorganisms$bitscore<50,]

highhomologysequences.middlebottom<-highhomologysequences.all[highhomologysequences.all$qaccver%in%middlebottombeginning$Name,]

print(paste0("BLASTs from the second half of proteins have ",nrow(highhomologysequences.middlebottom[!duplicated(highhomologysequences.middlebottom$Organism),]), " unique species with a bitscore of at least 50"))


print(paste0("BLAST of the second half of proteins have ",nrow(highhomologysequences.middlebottom)," sequences with a bitscore of at least 50"))

remove(curatedBLASTSresults,highhomologysequences.all)

###Using protein conservation scores from first full-length proteins, but only at the end.

endconservation<-data.frame(read_excel(paste0(outputdatabasedirectory,"Protein Conservation Scores from Bitscore 50 BLASTS.xlsx"),sheet = "Full qend end",col_names = T),check.names = F,stringsAsFactors = F)

endconservation<-endconservation[order(endconservation$Name),]

endconservation<-endconservation[!endconservation$Unique.Organisms.Bitscore.Threshold<40,]

endconservation<-endconservation[!endconservation$Amino.Acids<50,]

endconservation$endconservation.protein.sumweightedscore<-endconservation$Sum.Weighted.Proportion

endconservation<-endconservation[endconservation$Total.BLASTS>0,]

endconservation<-endconservation[endconservation$Details!="ALL BLASTS have bitscore lower than 50",]

endconservation<-endconservation[endconservation$Details!="No BLASTS Results",]

load(paste0(outputdatabasedirectory,"Full Protein BLASTS List curated compilation.RData"))

highhomologysequences.all<-curatedBLASTSresults$comp.fulluniqueorganisms[!curatedBLASTSresults$comp.fulluniqueorganisms$bitscore<50,]

highhomologysequences.full<-highhomologysequences.all[highhomologysequences.all$qaccver%in%endconservation$Name,]

print(paste0("BLASTs from full-length proteins have ",nrow(highhomologysequences.full[!duplicated(highhomologysequences.full$Organism),])," unique species with a bitscore of at least 50"))

print(paste0("BLAST of full-length proteins have ",nrow(highhomologysequences.full)," sequences with a bitscore of at least 50"))

remove(curatedBLASTSresults,highhomologysequences.all)

###Compiling all of the protein conservation scores from different regions of BLAST.

compconservation<-list(topmiddlebeginning[,c("Name","Query.Annotation","topmiddle.protein.sumweightedscore")],middlebottombeginning[,c("Name","Query.Annotation","middlebottom.protein.sumweightedscore")],endconservation[,c("Name","Amino.Acids","Query.Annotation","endconservation.protein.sumweightedscore")])

topbottomend<-join_all(compconservation,by=c("Name","Query.Annotation"),type="full", match="all")

topbottomend<-topbottomend[topbottomend$Name%in%initialtranslationspeed$Name,]

topbottomend.nona<-topbottomend[complete.cases(topbottomend), ]

uniquetop<-topmiddlebeginning[topmiddlebeginning$Name%in%topbottomend.nona$Name,]

uniquebot<-middlebottombeginning[middlebottombeginning$Name%in%topbottomend.nona$Name,]

temp<-topbottomend.nona[,c("Name","Query.Annotation","topmiddle.protein.sumweightedscore")]

temp$zone<-"topmiddle"

names(temp)[names(temp) == "topmiddle.protein.sumweightedscore"] <- "value"

temp2<-topbottomend.nona[,c("Name","Query.Annotation","middlebottom.protein.sumweightedscore")]

temp2$zone<-"middlebottom"

names(temp2)[names(temp2) == "middlebottom.protein.sumweightedscore"] <- "value"

temp3<-topbottomend.nona[,c("Name","Query.Annotation","endconservation.protein.sumweightedscore")]

temp3$zone<-"end"

names(temp3)[names(temp3) == "endconservation.protein.sumweightedscore"] <- "value"

temp4<-rbind.fill(list(temp,temp2,temp3))

###Figure 4A: First 40 protein conservation scores from BLAST on first half of proteins.

lineplot<-function(data){
  
  ggplot(data=data, aes(x=value, fill=zone, alpha=zone))+
    
  geom_histogram(colour="black",binwidth=.5, position="identity")+
    
  labs(title = "",x="Protein Conservation Scores",y="Frequency")+
    
  theme(panel.grid.major = element_line(size = 0.5, colour = 'black'),
        
        panel.grid.major.x = element_line(size = 0.5, colour = NA),
        
        panel.grid.major.y = element_line(size = 0.5, colour = NA),
        
        panel.grid.minor = element_line(colour = NA),
        
        plot.title = element_text(hjust = 0.5,size = 15),
        
        axis.title.x = element_text(size=35),
        
        axis.text.x = element_text(family="sans", vjust=0.3,hjust=0.5,size=30,angle=0),
        
        axis.text.y = element_text(size=35),
        
        axis.title.y = element_text(size=35),

        panel.background = element_rect(fill = 'white', colour = 'black'),
        
        legend.position="none")+
    
  annotate(geom="text", x=20, y=800, size=12,label="First 40", color="black")+

  scale_y_continuous(breaks = scales::pretty_breaks(n = 10),limits=c(0,1300),expand=c(0,0))+
  
  scale_fill_manual(values = c("#000000"))+
  
  scale_alpha_manual(values=c(0.4)) 
}

lineplot(data=temp4[temp4$zone=="topmiddle",])

###Figure 4B: Middle 40 protein conservation scores from BLAST on second half of proteins.

lineplot<-function(data){ggplot(data=data, aes(x=value, fill=zone, alpha=zone))+
    
  geom_histogram(colour="black",binwidth=.5, position="identity")+
    
  labs(title = "",x="Protein Conservation Scores",y="Frequency")+
    
  theme(panel.grid.major = element_line(size = 0.5, colour = 'black'),
        
        panel.grid.major.x = element_line(size = 0.5, colour = NA),
        
        panel.grid.major.y = element_line(size = 0.5, colour = NA),
        
        panel.grid.minor = element_line(colour = NA),
        
        plot.title = element_text(hjust = 0.5,size = 15),
        
        axis.title.x = element_text(size=35),
        
        axis.text.x = element_text(family="sans", vjust=0.3,hjust=0.5,size=30,angle=0),
        
        axis.text.y = element_text(size=35),
        
        axis.title.y = element_text(size=35),

        panel.background = element_rect(fill = 'white', colour = 'black'),
        
        legend.position="none")+
  
  annotate(geom="text", x=20, y=800, size=12,label="Middle 40", color="black")+

  scale_y_continuous(breaks = scales::pretty_breaks(n = 10),limits=c(0,1300),expand=c(0,0))+
  
  scale_fill_manual(values = c("green"))+
  
  scale_alpha_manual(values=c(0.5)) 
}

lineplot(data=temp4[temp4$zone=="middlebottom",])

###Figure 4C: Last 40 protein conservation scores from BLAST on full-length proteins.

lineplot<-function(data){ggplot(data=data, aes(x=value, fill=zone, alpha=zone))+
    
  geom_histogram(colour="black",binwidth=.5, position="identity")+
    
  labs(title = "",x="Protein Conservation Scores",y="Frequency")+
    
  theme(panel.grid.major = element_line(size = 0.5, colour = 'black'),
        
        panel.grid.major.x = element_line(size = 0.5, colour = NA),
        
        panel.grid.major.y = element_line(size = 0.5, colour = NA),
        
        panel.grid.minor = element_line(colour = NA),
        
        plot.title = element_text(hjust = 0.5,size = 15),
        
        axis.title.x = element_text(size=35),
        
        axis.text.x = element_text(family="sans", vjust=0.3,hjust=0.5,size=30,angle=0),
        
        axis.text.y = element_text(size=35),
        
        axis.title.y = element_text(size=35),

        panel.background = element_rect(fill = 'white', colour = 'black'),
        
        legend.position="none")+
    
  annotate(geom="text", x=20, y=800, size=12,label="Last 40", color="black")+

  scale_y_continuous(breaks = scales::pretty_breaks(n = 10),limits=c(0,1300),expand=c(0,0))+
  
  scale_fill_manual(values = c("red"))+
  
  scale_alpha_manual(values=c(0.3)) 
}

lineplot(data=temp4[temp4$zone=="end",])

###Figure 4 statistics

print(paste0("###Figure 4 statistics"))

print(paste("mean of first 40 protein conservation scores = ",signif(mean(temp4$value[temp4$zone=="topmiddle"]),digits = 5)))

print(paste("mean of middle 40 protein conservation scores =",signif(mean(temp4$value[temp4$zone=="middlebottom"]),digits = 5)))

print(paste("mean of last 40 protein conservation scores =",signif(mean(temp4$value[temp4$zone=="end"]),digits = 5)))

print(paste0("first 40 amino acids vs middle 40 amino acids conservation ks.test p= ",signif(ks.test(temp4$value[temp4$zone=="topmiddle"],temp4$value[temp4$zone=="middlebottom"],alternative="two.sided",simulate.p.value = T, B = 5000)[["p.value"]],digits = 5)))

print(paste0("first 40 amino acids vs last 40 amino acids conservation ks.test p= ",signif(ks.test(temp4$value[temp4$zone=="topmiddle"],temp4$value[temp4$zone=="end"],alternative="two.sided",simulate.p.value = T, B = 5000)[["p.value"]],digits = 5)))

print(paste0("middle 40 amino acids vs last 40 amino acids conservation ks.test p= ",signif(ks.test(temp4$value[temp4$zone=="middlebottom"],temp4$value[temp4$zone=="end"],alternative="two.sided",simulate.p.value = T, B = 5000)[["p.value"]],digits = 5)))

```

###Figure 5: Slow Initial Translation is correlated with poor N-terminal conservation.

```{r,echo=F}

###Figure 5A: N-termini relative initial translation speed as response variable and protein conservation as explanatory variable.

initialtranslationspeed<-data.frame(read_excel(paste0(outputdatabasedirectory,"Initial Translation Speed Table 40.xlsx"),col_names = T),check.names = F,stringsAsFactors = F)

initialtranslationspeed<-initialtranslationspeed[initialtranslationspeed$Nucleotides>300,]

initialtranslationspeed$inverserrt.beginningvsrest<-log2((1/(initialtranslationspeed$AverageRRTFirst40nostartcodon/initialtranslationspeed$AverageRRTentireminusfirst40)))

initialtranslationspeed$inverserrt.endvsrest<-log2((1/(initialtranslationspeed$AverageRRTThreePrime40/initialtranslationspeed$AverageRRTentireminuslast40)))

rampvsprotein<-join(topbottomend.nona,initialtranslationspeed,by=c("Name","Query.Annotation"),type="full", match="all")

rampvsprotein<-rampvsprotein[complete.cases(rampvsprotein), ]

size<-(round(nrow(rampvsprotein)/3))

rampvsprotein<-rampvsprotein[order(rampvsprotein$topmiddle.protein.sumweightedscore,decreasing = T),]

top33percent<-rampvsprotein[1:(size+1),]

midbegin<-(round(nrow(rampvsprotein)/3))

middle33percent<-rampvsprotein[(midbegin+2):(midbegin+(size)),]

bottom33percent<-rampvsprotein[(midbegin+(size+1)):nrow(rampvsprotein),]

check1<-top33percent[top33percent$Name%in%middle33percent$Name,]

check2<-top33percent[top33percent$Name%in%bottom33percent$Name,]

check3<-middle33percent[middle33percent$Name%in%bottom33percent$Name,]

if(nrow(check1)==0&nrow(check2)==0&nrow(check3)==0){
  
  print("SUCCESS!!! NO DUPLICATES!!!!")
} else {
  
  base::stop(print("ERROR?? THERE ARE DUPLICATES??"))
}

if(nrow(top33percent)+nrow(middle33percent)+nrow(bottom33percent)==nrow(rampvsprotein)){
  
  print("SUCCESS!!! SPLIT INTO 3 PARTS")
} else {
    
  base::stop(print("ERROR?? SOME ARE MISSING??"))
}

rampvsprotein$size="ZZZZZ"

rampvsprotein$size[rampvsprotein$Name %in% top33percent$Name]<-"Top 33%"

rampvsprotein$size[rampvsprotein$Name %in% bottom33percent$Name]<-"Bottom 33%"

rampvsprotein$size[rampvsprotein$Name %in% middle33percent$Name]<-"Middle 33%"

rampvsprotein$size=ordered(rampvsprotein$size, levels = c("Bottom 33%","Middle 33%","Top 33%","Blank"))

tgrq<-summarySE(rampvsprotein,measurevar = "inverserrt.beginningvsrest",groupvars =c("size") )

means=tgrq

means$size=ordered(means$size, levels = c("Bottom 33%","Middle 33%","Top 33%"))

means$lower=means$inverserrt.beginningvsrest-means$ci

means$upper=means$inverserrt.beginningvsrest+means$ci

means=means[1:3,]

lineplot<-function(data){
 
  ggplot(data=data)+

  geom_col(data = data,aes(x = size,y = inverserrt.beginningvsrest),width = .75, position = "dodge")+
    
  geom_errorbar(data=means,aes(x = size,ymin=lower, ymax=upper,y=inverserrt.beginningvsrest),width=.4, color="black", alpha=1)+
    
  labs(title="", x="N-terminal Protein Conservation Score",y="5' log2(Relative Initial Translation Speed)")+
    
  theme(panel.grid.major = element_line(size = 0.5, colour = 'black'),
        
        panel.grid.major.x = element_line(size = 0.5, colour = NA),
        
        panel.grid.major.y = element_line(size = 0.5, colour = NA),
        
        panel.grid.minor = element_line(colour = NA),
        
        plot.title = element_text(hjust = 0.5,size = 15),
        
        axis.title.x = element_text(size=30),
        
        axis.text.x = element_text(family="sans", vjust=0.9,hjust=0.5,size=35,angle=0),
        
        axis.text.y = element_text(size=28),
        
        axis.title.y = element_text(size=23),
        
        panel.background = element_rect(fill = 'white', colour = 'black'))+
    
  scale_y_continuous(breaks = scales::pretty_breaks(n = 10),limits = c(-0.05,0),expand=c(0,0))
}

lineplot(data=means)

topvsbottomp<-wilcox.test(top33percent$inverserrt.beginningvsrest,bottom33percent$inverserrt.beginningvsrest,alternative="two.sided",paired=F)[["p.value"]]

print(paste0("topvsbottom p= ",signif(topvsbottomp,digits = 5)))

topvsmidp<-wilcox.test(top33percent$inverserrt.beginningvsrest,middle33percent$inverserrt.beginningvsrest,alternative="two.sided",paired=F)[["p.value"]]

print(paste0("topvsmid p= ",signif(topvsmidp,digits = 5)))

bottomvsmidp<-wilcox.test(bottom33percent$inverserrt.beginningvsrest,middle33percent$inverserrt.beginningvsrest,alternative="two.sided",paired=F)[["p.value"]]

print(paste0("bottomvsmid p= ",signif(bottomvsmidp,digits = 5)))

p.adjust.methods<-c("holm", "hochberg", "hommel", "bonferroni", "BH", "BY","fdr", "none")

nterminalbottopmid<-sapply(p.adjust.methods, function(meth) p.adjust(c(topvsbottomp,topvsmidp,bottomvsmidp), meth,n=3))

nterminalbottopmid

###Figure 5B: N-termini protein conservation as response variable and relative initial translation speed as explanatory variable.

size<-(round(nrow(rampvsprotein)/3))

rampvsprotein<-rampvsprotein[order(rampvsprotein$log2Ratio40nostartcodonvsRest,decreasing = T),]

top33percent<-rampvsprotein[1:(size+1),]

midbegin<-(round(nrow(rampvsprotein)/3))

middle33percent<-rampvsprotein[(midbegin+2):(midbegin+(size)),]

bottom33percent<-rampvsprotein[(midbegin+(size+1)):nrow(rampvsprotein),]

check1<-top33percent[top33percent$Name%in%middle33percent$Name,]

check2<-top33percent[top33percent$Name%in%bottom33percent$Name,]

check3<-middle33percent[middle33percent$Name%in%bottom33percent$Name,]

if(nrow(check1)==0&nrow(check2)==0&nrow(check3)==0){
  
  print("SUCCESS!!! NO DUPLICATES!!!!")
} else {
    
  base::stop(print("ERROR?? THERE ARE DUPLICATES??"))
}

if(nrow(top33percent)+nrow(middle33percent)+nrow(bottom33percent)==nrow(rampvsprotein)){
  
  print("SUCCESS!!! SPLIT INTO 3 PARTS")
} else {
    
  base::stop(print("ERROR?? SOME ARE MISSING??"))
}

rampvsprotein<-rampvsprotein[,!names(rampvsprotein)%in%c("size")]

rampvsprotein$size="ZZZZZ"

rampvsprotein$size[rampvsprotein$Name %in% top33percent$Name]<-"SIT"

rampvsprotein$size[rampvsprotein$Name %in% bottom33percent$Name]<-"FIT"

rampvsprotein$size[rampvsprotein$Name %in% middle33percent$Name]<-"MIT"

rampvsprotein$size=ordered(rampvsprotein$size, levels = c("SIT","MIT","FIT"))

tgrq<-summarySE(rampvsprotein,measurevar = "topmiddle.protein.sumweightedscore",groupvars =c("size") )

means=tgrq

means$size=ordered(means$size, levels = c("SIT","MIT","FIT"))

means$lower=means$topmiddle.protein.sumweightedscore-means$ci

means$upper=means$topmiddle.protein.sumweightedscore+means$ci

means=means[1:3,]

lineplot<-function(data){
  
  ggplot(data=data)+

  geom_col(data = data,aes(x = size,y = topmiddle.protein.sumweightedscore),width = .75, position = "dodge")+

  geom_errorbar(data=means,aes(x = size,ymin=lower, ymax=upper,y=topmiddle.protein.sumweightedscore),width=.4, color="black", alpha=1)+

  labs(title="", x="5' log2(Relative Initial Translation Speed)",y="N-terminal Protein Conservation Score")+
    
  theme(panel.grid.major = element_line(size = 0.5, colour = 'black'),
        
        panel.grid.major.x = element_line(size = 0.5, colour = NA),
        
        panel.grid.major.y = element_line(size = 0.5, colour = NA),
        
        panel.grid.minor = element_line(colour = NA),
        
        plot.title = element_text(hjust = 0.5,size = 15),
        
        axis.title.x = element_text(size=30),
        
        axis.text.x = element_text(family="sans", vjust=0.9,hjust=0.5,size=35,angle=0),
        
        axis.text.y = element_text(size=28),
        
        axis.title.y = element_text(size=23),
        
        panel.background = element_rect(fill = 'white', colour = 'black'))+
  
  scale_y_continuous(breaks = scales::pretty_breaks(n = 10),limits = c(0,41),expand=c(0,0))
}

lineplot(data=means)

sitvsfitp<-wilcox.test(top33percent$topmiddle.protein.sumweightedscore,bottom33percent$topmiddle.protein.sumweightedscore,alternative="two.sided",paired=F)[["p.value"]]

print(paste0("sitvsfit p= ",signif(sitvsfitp,digits = 5)))

sitvsmitp<-wilcox.test(top33percent$topmiddle.protein.sumweightedscore,middle33percent$topmiddle.protein.sumweightedscore,alternative="two.sided",paired=F)[["p.value"]]

print(paste0("sitvsmit p= ",signif(sitvsmitp,digits = 5)))

fitvsmitp<-wilcox.test(bottom33percent$topmiddle.protein.sumweightedscore,middle33percent$topmiddle.protein.sumweightedscore,alternative="two.sided",paired=F)[["p.value"]]

print(paste0("fitvsmit p= ",signif(fitvsmitp,digits = 5)))

p.adjust.methods<-c("holm", "hochberg", "hommel", "bonferroni", "BH", "BY","fdr", "none")

nterminalsitmitfit <- sapply(p.adjust.methods, function(meth) p.adjust(c(sitvsfitp,sitvsmitp,fitvsmitp), meth,n=3))

nterminalsitmitfit

```

###Figure 6: mRNA Expression and ribosome density analyses.

```{r,echo=F,message=F}

###Figure 6A: gene expression vs 5' relative initial translation speed.

###downloaded yeast mRNA transcript dataset from Quantification of the yeast transcriptome by single-molecule sequencing    

#https://www.nature.com/articles/nbt.1551#Sec17

#Supplementary Information: Supplementary Table 1

expressionlist<-read_excel(paste0(inputdatabasedirectory,"41587_2009_BFnbt1551_MOESM7_ESM.xls"), 
    col_names = F)

expressionfilenew<-expressionlist[-c(1:3),-c(1,17,18)]

colnames(expressionfilenew)<-expressionfilenew[1,]

expressionfilenew<-expressionfilenew[-c(1),]

expressionfilenew$Avg<-as.numeric(expressionfilenew$Avg)

expresscoding<-expressionfilenew[expressionfilenew$ORF%in%rampvsprotein$Name,]

expresscoding<-expresscoding[order(expresscoding$Avg,decreasing = T),]

expresscoding<-expresscoding[!expresscoding$Avg<10,]

names(expresscoding)[names(expresscoding) == "ORF"] <- "Name"

rampvsprotein$beginningvsmiddle.log2.conservedratios<-log2(rampvsprotein$topmiddle.protein.sumweightedscore/rampvsprotein$middlebottom.protein.sumweightedscore)

rampvsprotein$endvsmiddle.log2.conservedratios<-log2(rampvsprotein$endconservation.protein.sumweightedscore/rampvsprotein$middlebottom.protein.sumweightedscore)

expresscodingramp<-join(rampvsprotein,expresscoding,by="Name",type="full", match="all")

expresscodingramp<-expresscodingramp[complete.cases(expresscodingramp$Avg), ]

size<-(round(nrow(expresscodingramp)/3))

expresscodingramp<-expresscodingramp[order(expresscodingramp$Avg,decreasing = T),]

expresscodingramp<-expresscodingramp[,!names(expresscodingramp)%in%c("size")]

top33percent<-expresscodingramp[1:(size+1),]

midbegin<-(round(nrow(expresscodingramp)/3))

middle33percent<-expresscodingramp[(midbegin+2):(midbegin+(size)),]

bottom33percent<-expresscodingramp[(midbegin+(size+1)):nrow(expresscodingramp),]

check1<-top33percent[top33percent$Name%in%middle33percent$Name,]

check2<-top33percent[top33percent$Name%in%bottom33percent$Name,]

check3<-middle33percent[middle33percent$Name%in%bottom33percent$Name,]

if(nrow(check1)==0&nrow(check2)==0&nrow(check3)==0){
 
   print("SUCCESS!!! NO DUPLICATES!!!!")
} else {
    
  base::stop(print("ERROR?? THERE ARE DUPLICATES??"))
}

if(nrow(top33percent)+nrow(middle33percent)+nrow(bottom33percent)==nrow(expresscodingramp)){
  
  print("SUCCESS!!! SPLIT INTO 3 PARTS")
} else {
    
  base::stop(print("ERROR?? SOME ARE MISSING??"))
}

expresscodingramp$size="ZZZZZ"

expresscodingramp$size[expresscodingramp$Name %in% top33percent$Name]<-"Top 33%"

expresscodingramp$size[expresscodingramp$Name %in% bottom33percent$Name]<-"Bottom 33%"

expresscodingramp$size[expresscodingramp$Name %in% middle33percent$Name]<-"Middle 33%"

expresscodingramp$size=ordered(expresscodingramp$size, levels = c("Bottom 33%","Middle 33%","Top 33%","Blank"))

tgrq<-summarySE(expresscodingramp,measurevar = "inverserrt.beginningvsrest",groupvars =c("size") )

means=tgrq

means$size=ordered(means$size, levels = c("Bottom 33%","Middle 33%","Top 33%"))

means$lower=means$inverserrt.beginningvsrest-means$ci

means$upper=means$inverserrt.beginningvsrest+means$ci

means=means[1:3,]

lineplot<-function(data){
  
  ggplot(data=data)+

  geom_col(data = data,aes(x = size,y = inverserrt.beginningvsrest),width = .75, position = "dodge")+
    
  geom_errorbar(data=means,aes(x = size,ymin=lower, ymax=upper,y=inverserrt.beginningvsrest),width=.4, color="black", alpha=1)+ 
    
  labs(title="", x="mRNA Transcript Levels",y="5' log2(Relative Initial Translation Speed)")+
    
  theme(panel.grid.major = element_line(size = 0.5, colour = 'black'),
        
        panel.grid.major.x = element_line(size = 0.5, colour = NA),
        
        panel.grid.major.y = element_line(size = 0.5, colour = NA),
        
        panel.grid.minor = element_line(colour = NA),
        
        plot.title = element_text(hjust = 0.5,size = 15),
        
        axis.title.x = element_text(size=30),
        
        axis.text.x = element_text(family="sans", vjust=0.9,hjust=0.5,size=35,angle=0),
        
        axis.text.y = element_text(size=28),
        
        axis.title.y = element_text(size=23),
        
        panel.background = element_rect(fill = 'white', colour = 'black'))+
    
  scale_y_continuous(breaks = scales::pretty_breaks(n = 10),limits = c(-0.05,0),expand=c(0,0))
}

lineplot(data=means)

topvsbottomp<-wilcox.test(top33percent$inverserrt.beginningvsrest,bottom33percent$inverserrt.beginningvsrest,alternative="two.sided",paired=F)[["p.value"]]

print(paste0("topvsbottom p= ",signif(topvsbottomp,digits = 5)))

topvsmidp<-wilcox.test(top33percent$inverserrt.beginningvsrest,middle33percent$inverserrt.beginningvsrest,alternative="two.sided",paired=F)[["p.value"]]

print(paste0("topvsmid p= ",signif(topvsmidp,digits = 5)))

bottomvsmidp<-wilcox.test(bottom33percent$inverserrt.beginningvsrest,middle33percent$inverserrt.beginningvsrest,alternative="two.sided",paired=F)[["p.value"]]

print(paste0("bottomvsmid p= ",signif(bottomvsmidp,digits = 5)))

p.adjust.methods<-c("holm", "hochberg", "hommel", "bonferroni", "BH", "BY","fdr", "none")

mrnatopmidbottomspeed <- sapply(p.adjust.methods, function(meth) p.adjust(c(topvsbottomp,topvsmidp,bottomvsmidp), meth,n=3))

mrnatopmidbottomspeed

paste0("mean log2(first40RRT/restRRT) of proteins with top 33% mRNA expression= ",signif(mean(top33percent$log2Ratio40nostartcodonvsRest),digits = 5))

paste0("mean log2(first40RRT/restRRT) of proteins with bottom 33% mRNA expression= ",signif(mean(bottom33percent$log2Ratio40nostartcodonvsRest),digits = 5))

paste0("mean log2(first40RRT/restRRT) of proteins with middle 33% mRNA expression= ",signif(mean(middle33percent$log2Ratio40nostartcodonvsRest),digits = 5))

###Figure 6B: gene expression vs N-termini protein conservation.

size<-(round(nrow(expresscodingramp)/3))

expresscodingramp<-expresscodingramp[order(expresscodingramp$Avg,decreasing = T),]

expresscodingramp<-expresscodingramp[,!names(expresscodingramp)%in%c("size")]

top33percent<-expresscodingramp[1:(size+1),]

midbegin<-(round(nrow(expresscodingramp)/3))

middle33percent<-expresscodingramp[(midbegin+2):(midbegin+(size)),]

bottom33percent<-expresscodingramp[(midbegin+(size+1)):nrow(expresscodingramp),]

check1<-top33percent[top33percent$Name%in%middle33percent$Name,]

check2<-top33percent[top33percent$Name%in%bottom33percent$Name,]

check3<-middle33percent[middle33percent$Name%in%bottom33percent$Name,]

if(nrow(check1)==0&nrow(check2)==0&nrow(check3)==0){
  
  print("SUCCESS!!! NO DUPLICATES!!!!")
} else {
    
  base::stop(print("ERROR?? THERE ARE DUPLICATES??"))
}

if(nrow(top33percent)+nrow(middle33percent)+nrow(bottom33percent)==nrow(expresscodingramp)){
  
  print("SUCCESS!!! SPLIT INTO 3 PARTS")
} else {
    
  base::stop(print("ERROR?? SOME ARE MISSING??"))
}

expresscodingramp$size="ZZZZZ"

expresscodingramp$size[expresscodingramp$Name %in% top33percent$Name]<-"Top 33%"

expresscodingramp$size[expresscodingramp$Name %in% bottom33percent$Name]<-"Bottom 33%"

expresscodingramp$size[expresscodingramp$Name %in% middle33percent$Name]<-"Middle 33%"

expresscodingramp$size=ordered(expresscodingramp$size, levels = c("Bottom 33%","Middle 33%","Top 33%","Blank"))

tgrq<-summarySE(expresscodingramp,measurevar = "topmiddle.protein.sumweightedscore",groupvars =c("size") )

means=tgrq

means$size=ordered(means$size, levels = c("Bottom 33%","Middle 33%","Top 33%"))

means$lower=means$topmiddle.protein.sumweightedscore-means$ci

means$upper=means$topmiddle.protein.sumweightedscore+means$ci

means=means[1:3,]

lineplot<-function(data){
  
  ggplot(data=data)+
    
  geom_col(data = data,aes(x = size,y = topmiddle.protein.sumweightedscore),width = .75, position = "dodge")+
    
  geom_errorbar(data=means,aes(x = size,ymin=lower, ymax=upper,y=topmiddle.protein.sumweightedscore),width=.4, color="black", alpha=1)+ 
    
  labs(title="", x="mRNA Transcript Levels",y="N-terminal Protein Conservation Score")+
    
  theme(panel.grid.major = element_line(size = 0.5, colour = 'black'),
        
        panel.grid.major.x = element_line(size = 0.5, colour = NA),
        
        panel.grid.major.y = element_line(size = 0.5, colour = NA),
        
        panel.grid.minor = element_line(colour = NA),
        
        plot.title = element_text(hjust = 0.5,size = 15),
        
        axis.title.x = element_text(size=30),
        
        axis.text.x = element_text(family="sans", vjust=0.9,hjust=0.5,size=35,angle=0),
        
        axis.text.y = element_text(size=28),
        
        axis.title.y = element_text(size=23),
        
        panel.background = element_rect(fill = 'white', colour = 'black'))+
    
  scale_y_continuous(breaks = scales::pretty_breaks(n = 10),limits = c(0,41),expand=c(0,0))
}

lineplot(data=means)

topvsbottomp<-wilcox.test(top33percent$topmiddle.protein.sumweightedscore,bottom33percent$topmiddle.protein.sumweightedscore,alternative="two.sided",paired=F)[["p.value"]]

print(paste0("topvsbottom p= ",signif(topvsbottomp,digits = 5)))

topvsmidp<-wilcox.test(top33percent$topmiddle.protein.sumweightedscore,middle33percent$topmiddle.protein.sumweightedscore,alternative="two.sided",paired=F)[["p.value"]]

print(paste0("topvsmid p= ",signif(topvsmidp,digits = 5)))

bottomvsmidp<-wilcox.test(bottom33percent$topmiddle.protein.sumweightedscore,middle33percent$topmiddle.protein.sumweightedscore,alternative="two.sided",paired=F)[["p.value"]]

print(paste0("bottomvsmid p= ",signif(bottomvsmidp,digits = 5)))

p.adjust.methods<-c("holm", "hochberg", "hommel", "bonferroni", "BH", "BY","fdr", "none")

mrnatopmidbottomprotein <- sapply(p.adjust.methods, function(meth) p.adjust(c(topvsbottomp,topvsmidp,bottomvsmidp), meth,n=3))

mrnatopmidbottomprotein

paste0("mean protein conservation score of N-termini with top 33% mRNA expression= ",signif(mean(top33percent$topmiddle.protein.sumweightedscore),digits = 5))

paste0("mean protein conservation score of N-termini with bottom 33% mRNA expression= ",signif(mean(bottom33percent$topmiddle.protein.sumweightedscore),digits = 5))

paste0("mean protein conservation score of N-termini with middle 33% mRNA expression= ",signif(mean(middle33percent$topmiddle.protein.sumweightedscore)
,digits = 5))

###Figure 6C: Ribosome density vs 5' relative initial translation speed.

###Downloaded yeast ribosome density dataset from Genome-wide analysis of mRNA translation profiles in Saccharomyces cerevisiae

#https://www.pnas.org/doi/full/10.1073/pnas.0635171100#supplementary-materials  

#Supporting Information - 5171_Table3.xls

densitylist<-read_excel(paste0(inputdatabasedirectory,"5171_table3.xls"),col_names = F)

densitylistnew<-densitylist[-c(1:11),]

colnames(densitylistnew)<-densitylistnew[1,]

densitylistnew<-densitylistnew[-c(1),]

densitylistnew<-densitylistnew[!is.na(densitylistnew$Density),]

densitylistnew$Density<-as.numeric(densitylistnew$Density)

names(densitylistnew)[names(densitylistnew) == "YORF"] <- "Name"

densitylistnew<-densitylistnew[densitylistnew$Name%in%rampvsprotein$Name,]

expresscodingdensity<-join(rampvsprotein,densitylistnew,by="Name",type="full", match="all")

expresscodingdensity<-expresscodingdensity[complete.cases(expresscodingdensity$Density), ]

size<-(round(nrow(expresscodingdensity)/3))

expresscodingdensity<-expresscodingdensity[order(expresscodingdensity$Density,decreasing = T),]

expresscodingdensity<-expresscodingdensity[,!names(expresscodingdensity)%in%c("size")]

top33percent<-expresscodingdensity[1:(size+1),]

midbegin<-(round(nrow(expresscodingdensity)/3))

middle33percent<-expresscodingdensity[(midbegin+2):(midbegin+(size)),]

bottom33percent<-expresscodingdensity[(midbegin+(size+1)):nrow(expresscodingdensity),]

check1<-top33percent[top33percent$Name%in%middle33percent$Name,]

check2<-top33percent[top33percent$Name%in%bottom33percent$Name,]

check3<-middle33percent[middle33percent$Name%in%bottom33percent$Name,]

if(nrow(check1)==0&nrow(check2)==0&nrow(check3)==0){
  
  print("SUCCESS!!! NO DUPLICATES!!!!")
} else {
    
  base::stop(print("ERROR?? THERE ARE DUPLICATES??"))
}

if(nrow(top33percent)+nrow(middle33percent)+nrow(bottom33percent)==nrow(expresscodingdensity)){
  
  print("SUCCESS!!! SPLIT INTO 3 PARTS")
} else {
    
  base::stop(print("ERROR?? SOME ARE MISSING??"))
}

expresscodingdensity$size="ZZZZZ"

expresscodingdensity$size[expresscodingdensity$Name %in% top33percent$Name]<-"Top 33%"

expresscodingdensity$size[expresscodingdensity$Name %in% bottom33percent$Name]<-"Bottom 33%"

expresscodingdensity$size[expresscodingdensity$Name %in% middle33percent$Name]<-"Middle 33%"

expresscodingdensity$size=ordered(expresscodingdensity$size, levels = c("Bottom 33%","Middle 33%","Top 33%","Blank"))

tgrq<-summarySE(expresscodingdensity,measurevar = "inverserrt.beginningvsrest",groupvars =c("size") )

means=tgrq

means$size=ordered(means$size, levels = c("Bottom 33%","Middle 33%","Top 33%"))

means$lower=means$inverserrt.beginningvsrest-means$ci

means$upper=means$inverserrt.beginningvsrest+means$ci

means=means[1:3,]

lineplot<-function(data){
  
  ggplot(data=data)+
    
  geom_col(data = data,aes(x = size,y = inverserrt.beginningvsrest),width = .75, position = "dodge")+
    
  geom_errorbar(data=means,aes(x = size,ymin=lower, ymax=upper,y=inverserrt.beginningvsrest),width=.4, color="black", alpha=1)+ 
    
  labs(title="", x="Ribosome Density",y="5' log2(Relative Initial Translation Speed)")+
    
  theme(panel.grid.major = element_line(size = 0.5, colour = 'black'),
        
        panel.grid.major.x = element_line(size = 0.5, colour = NA),
        
        panel.grid.major.y = element_line(size = 0.5, colour = NA),
        
        panel.grid.minor = element_line(colour = NA),
        
        plot.title = element_text(hjust = 0.5,size = 15),
        
        axis.title.x = element_text(size=30),
        
        axis.text.x = element_text(family="sans", vjust=0.9,hjust=0.5,size=35,angle=0),
        
        axis.text.y = element_text(size=28),
        
        axis.title.y = element_text(size=23),
        
        panel.background = element_rect(fill = 'white', colour = 'black'))+
    
  scale_y_continuous(breaks = scales::pretty_breaks(n = 10),limits = c(-0.05,0),expand=c(0,0))
}

lineplot(data=means)

topvsbottomp<-wilcox.test(top33percent$inverserrt.beginningvsrest,bottom33percent$inverserrt.beginningvsrest,alternative="two.sided",paired=F)[["p.value"]]

print(paste0("topvsbottom p= ",signif(topvsbottomp,digits = 5)))

topvsmidp<-wilcox.test(top33percent$inverserrt.beginningvsrest,middle33percent$inverserrt.beginningvsrest,alternative="two.sided",paired=F)[["p.value"]]

print(paste0("topvsmid p= ",signif(topvsmidp,digits = 5)))

bottomvsmidp<-wilcox.test(bottom33percent$inverserrt.beginningvsrest,middle33percent$inverserrt.beginningvsrest,alternative="two.sided",paired=F)[["p.value"]]

print(paste0("bottomvsmid p= ",signif(bottomvsmidp,digits = 5)))

p.adjust.methods<-c("holm", "hochberg", "hommel", "bonferroni", "BH", "BY","fdr", "none")

ribosometopmidbottomspeed<- sapply(p.adjust.methods, function(meth) p.adjust(c(topvsbottomp,topvsmidp,bottomvsmidp), meth,n=3))

ribosometopmidbottomspeed

paste0("mean log2(first40RRT/restRRT) of proteins with top 33% ribosome density= ",signif(mean(top33percent$log2Ratio40nostartcodonvsRest),digits = 5))

paste0("mean log2(first40RRT/restRRT) of proteins with bottom 33% ribosome density= ",signif(mean(bottom33percent$log2Ratio40nostartcodonvsRest),digits = 5))

paste0("mean log2(first40RRT/restRRT) of proteins with middle 33% ribosome density= ",signif(mean(middle33percent$log2Ratio40nostartcodonvsRest),digits = 5))

##Figure 6D: Ribosome density vs N-termini protein conservation.

size<-(round(nrow(expresscodingdensity)/3))

expresscodingdensity<-expresscodingdensity[order(expresscodingdensity$Density,decreasing = T),]

expresscodingdensity<-expresscodingdensity[,!names(expresscodingdensity)%in%c("size")]

top33percent<-expresscodingdensity[1:(size+1),]

midbegin<-(round(nrow(expresscodingdensity)/3))

middle33percent<-expresscodingdensity[(midbegin+2):(midbegin+(size)),]

bottom33percent<-expresscodingdensity[(midbegin+(size+1)):nrow(expresscodingdensity),]

check1<-top33percent[top33percent$Name%in%middle33percent$Name,]

check2<-top33percent[top33percent$Name%in%bottom33percent$Name,]

check3<-middle33percent[middle33percent$Name%in%bottom33percent$Name,]

if(nrow(check1)==0&nrow(check2)==0&nrow(check3)==0){
  
  print("SUCCESS!!! NO DUPLICATES!!!!")
} else {
    
  base::stop(print("ERROR?? THERE ARE DUPLICATES??"))
}

if(nrow(top33percent)+nrow(middle33percent)+nrow(bottom33percent)==nrow(expresscodingdensity)){
  
  print("SUCCESS!!! SPLIT INTO 3 PARTS")
} else {
    
  base::stop(print("ERROR?? SOME ARE MISSING??"))
}
expresscodingdensity$size="ZZZZZ"

expresscodingdensity$size[expresscodingdensity$Name %in% top33percent$Name]<-"Top 33%"

expresscodingdensity$size[expresscodingdensity$Name %in% bottom33percent$Name]<-"Bottom 33%"

expresscodingdensity$size[expresscodingdensity$Name %in% middle33percent$Name]<-"Middle 33%"

expresscodingdensity$size=ordered(expresscodingdensity$size, levels = c("Bottom 33%","Middle 33%","Top 33%","Blank"))

tgrq<-summarySE(expresscodingdensity,measurevar = "topmiddle.protein.sumweightedscore",groupvars =c("size") )

means=tgrq

means$size=ordered(means$size, levels = c("Bottom 33%","Middle 33%","Top 33%"))

means$lower=means$topmiddle.protein.sumweightedscore-means$ci

means$upper=means$topmiddle.protein.sumweightedscore+means$ci

means=means[1:3,]

lineplot<-function(data){
  
  ggplot(data=data)+
    
  geom_col(data = data,aes(x = size,y = topmiddle.protein.sumweightedscore),width = .75, position = "dodge")+
    
  geom_errorbar(data=means,aes(x = size,ymin=lower, ymax=upper,y=topmiddle.protein.sumweightedscore),width=.4, color="black", alpha=1)+ 
    
  labs(title="", x="Ribosome Density",y="N-terminal Protein Conservation Score")+
    
  theme(panel.grid.major = element_line(size = 0.5, colour = 'black'),
        
        panel.grid.major.x = element_line(size = 0.5, colour = NA),
        
        panel.grid.major.y = element_line(size = 0.5, colour = NA),
        
        panel.grid.minor = element_line(colour = NA),
        
        plot.title = element_text(hjust = 0.5,size = 15),
        
        axis.title.x = element_text(size=30),
        
        axis.text.x = element_text(family="sans", vjust=0.9,hjust=0.5,size=35,angle=0),
        
        axis.text.y = element_text(size=28),
        
        axis.title.y = element_text(size=23),
        
        panel.background = element_rect(fill = 'white', colour = 'black'))+
    
  scale_y_continuous(breaks = scales::pretty_breaks(n = 10),limits = c(0,41),expand=c(0,0))
}

lineplot(data=means)

topvsbottomp<-wilcox.test(top33percent$topmiddle.protein.sumweightedscore,bottom33percent$topmiddle.protein.sumweightedscore,alternative="two.sided",paired=F)[["p.value"]]

print(paste0("topvsbottom p= ",signif(topvsbottomp,digits = 5)))

topvsmidp<-wilcox.test(top33percent$topmiddle.protein.sumweightedscore,middle33percent$topmiddle.protein.sumweightedscore,alternative="two.sided",paired=F)[["p.value"]]

print(paste0("topvsmid p= ",signif(topvsmidp,digits = 5)))

bottomvsmidp<-wilcox.test(bottom33percent$topmiddle.protein.sumweightedscore,middle33percent$topmiddle.protein.sumweightedscore,alternative="two.sided",paired=F)[["p.value"]]

print(paste0("bottomvsmid p= ",signif(bottomvsmidp,digits = 5)))

p.adjust.methods<-c("holm", "hochberg", "hommel", "bonferroni", "BH", "BY","fdr", "none")

ribosometopmidbottomprotein <- sapply(p.adjust.methods, function(meth) p.adjust(c(topvsbottomp,topvsmidp,bottomvsmidp), meth,n=3))

ribosometopmidbottomprotein

paste0("mean protein conservation score of N-termini with top 33% ribosome density= ",signif(mean(top33percent$topmiddle.protein.sumweightedscore),digits = 5))

paste0("mean protein conservation score of N-termini with bottom 33% ribosome density= ",signif(mean(bottom33percent$topmiddle.protein.sumweightedscore),digits = 5))

paste0("mean protein conservation score of N-termini with middle 33% ribosome density= ",signif(mean(middle33percent$topmiddle.protein.sumweightedscore),digits = 5))

```

###Figure 7: Slow Initial Translation inhibits gene expression.

```{r,echo=F,message=F,fig.height=7.5, fig.width=16,fig.align='center'}

###Figure 7: yeast were transformed with a bidirectional fluorescent reporter gene that has GFP with the first 40 codons recoded to have a SIT, MIT, FIT with and without a putative collision site.

mitvssit <- read_csv(paste0(inputdatabasedirectory,"MIT vs SIT/Statistics/021121_Ramp2_001-Batch_Analysis_11022021133313.csv"), 
    col_names = T)

fitsamples <- read_csv(paste0(inputdatabasedirectory,"FIT vs MIT vs SIT/Statistics/2182021_Ramp3-Batch_Analysis_18022021145033.csv"), 
    col_names = T)

yeastcodonusage<-data.frame(read_excel(paste0(outputdatabasedirectory,"Saccharomyces cerevisiae codon usage table.xlsx"),col_names = T),check.names = F,stringsAsFactors = F)

gene<-read.fasta(paste0(inputdatabasedirectory,"ramp-gfp constructs.fasta"))

generep<-1

syntheticgenes<-data.frame(Name="ZZZZZ",AverageRRTFirst41=99999,AverageRRTFirst41nostartcodon=99999,AverageRRTentireminusfirst41=99999,AverageRRTentiregene=99999,AverageRRTentiregenenostartcodon=99999,Nucleotides=99999,DNA.Sequence="ZZZZZ",Protein.Sequence="ZZZZZ",Number=1:length(gene))

rampcodonlist<-rep(list(NULL),length(gene))

ramprrtlist<-rep(list(NULL),length(gene))

codonusage<-rep(list(NULL),length(gene))

aausage<-rep(list(NULL),length(gene))

while (generep<(length(gene)+1)){
  
  dnasequence<-getSequence(gene[[generep]])
  
  genenametemp<-s2c(unlist(getAnnot(gene[generep])))
  
  genename<-c2s(genenametemp[-1])
  
  firstposition<-1
  
  indexxxx<-1
  
  syntheticgenescodonlist<-rep(NA,(length(dnasequence)/3))
  
  syntheticgenesrrtlist<-rep(NA,(length(dnasequence)/3))

  while(firstposition<length(dnasequence)){
    
    syntheticgenescodonlist[indexxxx]<-toupper(c2s(dnasequence[firstposition:(firstposition+2)]))
    
    syntheticgenesrrtlist[indexxxx]<-yeastcodonusage$RRT[yeastcodonusage$Codons==toupper(c2s(dnasequence[firstposition:(firstposition+2)]))]
    
    firstposition<-firstposition+3
    
    indexxxx<-indexxxx+1
  }
  
  data.frame(table(syntheticgenescodonlist))
  
  codonusagetemp<-data.frame(table(syntheticgenescodonlist))
  
  aausagetemp<-data.frame(table(translate(dnasequence)))
  
  names(codonusagetemp)[1] <- "Codons"
  
  names(codonusagetemp)[2] <- "Frame 1 (Coding) Observed Counts"
  
  codonusage[generep]<-list(codonusagetemp)
  
  names(codonusage)[generep] <- genename
  
  aausage[generep]<-list(aausagetemp)
  
  names(aausage)[generep] <- genename

  syntheticgenes$Name[generep]<-genename
  
  syntheticgenes$AverageRRTFirst41[generep]<-mean(syntheticgenesrrtlist[1:41])
  
  syntheticgenes$AverageRRTFirst41nostartcodon[generep]<-mean(syntheticgenesrrtlist[2:41])
  
  syntheticgenes$AverageRRTentireminusfirst41[generep]<-mean(syntheticgenesrrtlist[(41+1):length(syntheticgenesrrtlist)])
  
  syntheticgenes$AverageRRTentiregene[generep]<-mean(syntheticgenesrrtlist)
  
  syntheticgenes$AverageRRTentiregenenostartcodon[generep]<-mean(syntheticgenesrrtlist[2:length(syntheticgenesrrtlist)])
  
  syntheticgenes$Nucleotides[generep]<-length(dnasequence)
  
  syntheticgenes$DNA.Sequence[generep]<-c2s(toupper(dnasequence))
  
  syntheticgenes$Protein.Sequence[generep]<-c2s(translate(dnasequence))

  rampcodonlist[generep]<-list(syntheticgenescodonlist)
  
  names(rampcodonlist)[generep] <- genename
  
  ramprrtlist[generep]<-list(syntheticgenesrrtlist)
  
  names(ramprrtlist)[generep] <- genename

  generep<-generep+1
}

###Ramp 1 is MIT, Ramp 2 is SIT, Ramp 3 is FIT, Ramp 4 is MIT:PCS, Ramp 5 is SIT:PCS, and Ramp 6 is FIT:PCS

print(paste0("Ramp 1 is MIT, Ramp 2 is SIT, Ramp 3 is FIT"))

print(paste0("Ramp 4 is MIT:PCS, Ramp 5 is SIT:PCS, Ramp 6 is FIT:PCS"))

syntheticgenes$Ratio41nostartcodonvsRest=syntheticgenes$AverageRRTFirst41nostartcodon/syntheticgenes$AverageRRTentireminusfirst

syntheticgenes$log2Ratio41nostartcodonvsRest<-log2(syntheticgenes$Ratio41nostartcodonvsRest)

mitsitdat<-mitvssit[,c("Record Date","All Events FITC-A Mean","All Events PE-Texas Red-A Mean")]

mitsitdat$Name<-"ZZZZZ"

mitsitdat$Name[c(1,2,(nrow(mitsitdat)-1):nrow(mitsitdat))]<-"control"

mitsitdat$Name[c(3:(3+21))]<-"Ramp1. MIT"

mitsitdat$Name[c(25:(25+21))]<-"Ramp2. SIT"

mitsitdat$Name[c(47:(47+21))]<-"Ramp4. MIT with PCS"

mitsitdat$Name[c(69:(69+21))]<-"Ramp5. SIT with PCS"

mitsitdat$Name<-factor(mitsitdat$Name)

fitonlydat<-fitsamples[,c("Record Date","All Events FITC-A Mean","All Events PE-Texas Red-A Mean")]

fitonlydat$Name<-"ZZZZZ"

fitonlydat$Name[c(1,2,(nrow(fitonlydat)-1):nrow(fitonlydat))]<-"control"

fitonlydat$Name[c(3:(3+24))]<-"Ramp3. FIT"

fitonlydat$Name[c(28:(28+24))]<-"Ramp6. FIT with PCS"

fitonlydat$Name[c(53:(53+2))]<-"Ramp1. MIT"

fitonlydat$Name[c(56:(56+2))]<-"Ramp2. SIT"

fitonlydat$Name[c(59:(59+2))]<-"Ramp4. MIT with PCS"

fitonlydat$Name[c(62:(62+2))]<-"Ramp5. SIT with PCS"

fitonlydat$Name<-factor(fitonlydat$Name)

yeastfluorescence<-rbind(mitsitdat,fitonlydat)

yeastfluorescence$Name=ordered(yeastfluorescence$Name, levels = c("Ramp1. MIT","Ramp2. SIT","Ramp3. FIT","Ramp4. MIT with PCS","Ramp5. SIT with PCS","Ramp6. FIT with PCS","control"))

yeastfluorescence<-yeastfluorescence[order(yeastfluorescence$Name),]

generep<-1

while(generep<length(levels(yeastfluorescence$Name))+1){
  
  print(paste0(nrow(yeastfluorescence[yeastfluorescence$Name==levels(yeastfluorescence$Name)[generep],])," ",levels(yeastfluorescence$Name)[generep]," samples"))
  
  generep<-generep+1
}

yeastfluorescence$Translation.speed.size<-"ZZZZZ"

yeastfluorescence$Translation.speed.size[yeastfluorescence$Name=="Ramp1. MIT"]<-"MIT"

yeastfluorescence$Translation.speed.size[yeastfluorescence$Name=="Ramp2. SIT"]<-"SIT"

yeastfluorescence$Translation.speed.size[yeastfluorescence$Name=="Ramp3. FIT"]<-"FIT"

yeastfluorescence$Translation.speed.size[yeastfluorescence$Name=="Ramp4. MIT with PCS"]<-"MIT:PCS"

yeastfluorescence$Translation.speed.size[yeastfluorescence$Name=="Ramp5. SIT with PCS"]<-"SIT:PCS"

yeastfluorescence$Translation.speed.size[yeastfluorescence$Name=="Ramp6. FIT with PCS"]<-"FIT:PCS"

newsyntheticgenes<-syntheticgenes

newsyntheticgenes$inverseRRT<-log2(1/(newsyntheticgenes$AverageRRTFirst41nostartcodon/newsyntheticgenes$AverageRRTentireminusfirst41))

newsyntheticgenes$Translation.speed.size[newsyntheticgenes$Name=="Ramp1. MIT"]<-"MIT"

newsyntheticgenes$Translation.speed.size[newsyntheticgenes$Name=="Ramp2. SIT"]<-"SIT"

newsyntheticgenes$Translation.speed.size[newsyntheticgenes$Name=="Ramp3. FIT"]<-"FIT"

newsyntheticgenes$Translation.speed.size[newsyntheticgenes$Name=="Ramp4. MIT with PCS"]<-"MIT:PCS"

newsyntheticgenes$Translation.speed.size[newsyntheticgenes$Name=="Ramp5. SIT with PCS"]<-"SIT:PCS"

newsyntheticgenes$Translation.speed.size[newsyntheticgenes$Name=="Ramp6. FIT with PCS"]<-"FIT:PCS"

newsyntheticgenes<-newsyntheticgenes[,c("Name","Translation.speed.size","Nucleotides","AverageRRTFirst41","AverageRRTFirst41nostartcodon","AverageRRTentiregene","AverageRRTentiregenenostartcodon","Ratio41nostartcodonvsRest","log2Ratio41nostartcodonvsRest","inverseRRT","DNA.Sequence","Protein.Sequence")]

wb<-createWorkbook()

addWorksheet(wb, "Output")

writeData(wb,sheet="Output",x=newsyntheticgenes[,c("Name","Translation.speed.size","Nucleotides","AverageRRTFirst41","AverageRRTFirst41nostartcodon","AverageRRTentiregene","AverageRRTentiregenenostartcodon","Ratio41nostartcodonvsRest","log2Ratio41nostartcodonvsRest","DNA.Sequence","Protein.Sequence")])

saveWorkbook(wb, paste0(outputdatabasedirectory,"Synthetic GFP constructs statistics.xlsx"), overwrite = T)

###The MIT, FIT, and SIT all have the same amino acid sequence, but differ from the MIT:PCS, SIT:PCS, and SIT:PCS because the PCS encode for 2 different amino acids.

(sapply(list(newsyntheticgenes$Protein.Sequence[newsyntheticgenes$Translation.speed.size=="SIT"], newsyntheticgenes$Protein.Sequence[newsyntheticgenes$Translation.speed.size=="FIT"], newsyntheticgenes$Protein.Sequence[newsyntheticgenes$Translation.speed.size=="MIT:PCS"], newsyntheticgenes$Protein.Sequence[newsyntheticgenes$Translation.speed.size=="SIT:PCS"],newsyntheticgenes$Protein.Sequence[newsyntheticgenes$Translation.speed.size=="FIT:PCS"]), FUN = identical, newsyntheticgenes$Protein.Sequence[newsyntheticgenes$Translation.speed.size=="MIT"]))

(sapply(list(newsyntheticgenes$Protein.Sequence[newsyntheticgenes$Translation.speed.size=="MIT"], newsyntheticgenes$Protein.Sequence[newsyntheticgenes$Translation.speed.size=="SIT"], newsyntheticgenes$Protein.Sequence[newsyntheticgenes$Translation.speed.size=="FIT"], newsyntheticgenes$Protein.Sequence[newsyntheticgenes$Translation.speed.size=="SIT:PCS"],newsyntheticgenes$Protein.Sequence[newsyntheticgenes$Translation.speed.size=="FIT:PCS"]), FUN = identical, newsyntheticgenes$Protein.Sequence[newsyntheticgenes$Translation.speed.size=="MIT:PCS"]))

###Checks on codon usage and protein sequence.

(sapply(list(aausage$Ramp2,aausage$Ramp3,aausage$ramp4,aausage$Ramp5,aausage$Ramp6), FUN = identical, aausage$Ramp1))

(sapply(list(aausage$Ramp1,aausage$Ramp2,aausage$ramp3,aausage$Ramp5,aausage$Ramp6), FUN = identical, aausage$Ramp4))

(sapply(list(codonusage$Ramp2,codonusage$Ramp3,codonusage$ramp4,codonusage$Ramp5,codonusage$Ramp6), FUN = identical, codonusage$Ramp1))

(sapply(list(codonusage$Ramp1,codonusage$Ramp2,codonusage$ramp3,codonusage$Ramp5,codonusage$Ramp6), FUN = identical, codonusage$Ramp4))

yeastfluorescence<-join(yeastfluorescence,newsyntheticgenes[,c("Name","log2Ratio41nostartcodonvsRest","inverseRRT","Translation.speed.size")],by="Translation.speed.size")

yeastfluorescence$greenredratio<-yeastfluorescence$`All Events FITC-A Mean`/yeastfluorescence$`All Events PE-Texas Red-A Mean`

yeastfluorescence$log2ratio<-log2(yeastfluorescence$greenredratio)

yeastfluorescence$Translation.speed.size<-factor(yeastfluorescence$Translation.speed.size)

yeastfluorescence<-yeastfluorescence[yeastfluorescence$Translation.speed.size!="ZZZZZ",]

tgrq<-summarySE(yeastfluorescence,measurevar = "greenredratio",groupvars =c("Translation.speed.size") )

means=tgrq

means$Translation.speed.size=ordered(means$Translation.speed.size, levels = c("SIT","MIT","FIT","SIT:PCS","MIT:PCS","FIT:PCS"))

means$lower=means$greenredratio-means$ci

means$upper=means$greenredratio+means$ci

lineplot<-function(data){
  
  ggplot(data=data)+

  geom_col(data = data,aes(x = Translation.speed.size,y = greenredratio),width = .75, position = "dodge")+
    
  labs(title = "",x="",y="GFP/RFP")+
    
  geom_hline(yintercept = 1,linetype="dashed")+
    
  geom_errorbar(data=means,aes(x = Translation.speed.size,ymin=lower, ymax=upper,y=greenredratio),width=.4, color="black", alpha=1)+
    
  theme(panel.grid.major = element_line(size = 0.5, colour = 'black'),
        
        panel.grid.major.x = element_line(size = 0.5, colour = NA),
        
        panel.grid.major.y = element_line(size = 0.5, colour = NA),
        
        panel.grid.minor = element_line(colour = NA),
        
        plot.title = element_text(hjust = 0.5,size = 15),
        
        axis.title.x = element_text(size=60),
        
        axis.text.x = element_text(family="sans", vjust=0.3,hjust=0.5,size=36,angle=0),
        
        axis.text.y = element_text(size=30),
        
        axis.title.y = element_text(size=50),
        
        panel.background = element_rect(fill = 'white', colour = 'black'))+
    
  scale_y_continuous(breaks = scales::pretty_breaks(n = 6),limits=c(0,3),expand=c(0,0))+
  
  scale_x_discrete("",labels= c("SIT","MIT","FIT","SIT:PCS","MIT:PCS","FIT:PCS"))
}

lineplot(data=means)

###Statistics

print(paste0("SITvsmit p= ",signif(wilcox.test(yeastfluorescence$greenredratio[yeastfluorescence$Translation.speed.size=="SIT"],yeastfluorescence$greenredratio[yeastfluorescence$Translation.speed.size=="MIT"],alternative="two.sided",paired=F)[["p.value"]],digits = 5)))

print(paste0("SITvsFIT p= ",signif(wilcox.test(yeastfluorescence$greenredratio[yeastfluorescence$Translation.speed.size=="SIT"],yeastfluorescence$greenredratio[yeastfluorescence$Translation.speed.size=="FIT"],alternative="two.sided",paired=F)[["p.value"]],digits = 5)))

print(paste0("MITvsFIT p= ",signif(wilcox.test(yeastfluorescence$greenredratio[yeastfluorescence$Translation.speed.size=="MIT"],yeastfluorescence$greenredratio[yeastfluorescence$Translation.speed.size=="FIT"],alternative="two.sided",paired=F)[["p.value"]],digits = 5)))

paste0("mean GFP/RFP for SIT= ",signif(mean(yeastfluorescence$greenredratio[yeastfluorescence$Translation.speed.size=="SIT"]),digits = 5))

paste0("mean GFP/RFP for MIT= ",signif(mean(yeastfluorescence$greenredratio[yeastfluorescence$Translation.speed.size=="MIT"]),digits = 5))

paste0("mean GFP/RFP for FIT= ",signif(mean(yeastfluorescence$greenredratio[yeastfluorescence$Translation.speed.size=="FIT"]),digits = 5))

print(paste0("SIT:PCSvsMIT:PCS p= ",signif(wilcox.test(yeastfluorescence$greenredratio[yeastfluorescence$Translation.speed.size=="SIT:PCS"],yeastfluorescence$greenredratio[yeastfluorescence$Translation.speed.size=="MIT:PCS"],alternative="two.sided",paired=F)[["p.value"]],digits = 5)))

print(paste0("SIT:PCSvsFIT:PCS p= ",signif(wilcox.test(yeastfluorescence$greenredratio[yeastfluorescence$Translation.speed.size=="SIT:PCS"],yeastfluorescence$greenredratio[yeastfluorescence$Translation.speed.size=="FIT:PCS"],alternative="two.sided",paired=F)[["p.value"]],digits = 5)))

print(paste0("MIT:PCSvsFIT:PCS p= ",signif(wilcox.test(yeastfluorescence$greenredratio[yeastfluorescence$Translation.speed.size=="MIT:PCS"],yeastfluorescence$greenredratio[yeastfluorescence$Translation.speed.size=="FIT:PCS"],alternative="two.sided",paired=F)[["p.value"]],digits = 5)))

paste0("mean GFP/RFP for SIT:PCS= ",signif(mean(yeastfluorescence$greenredratio[yeastfluorescence$Translation.speed.size=="SIT:PCS"]),digits = 5))

paste0("mean GFP/RFP for MIT:PCS= ",signif(mean(yeastfluorescence$greenredratio[yeastfluorescence$Translation.speed.size=="MIT:PCS"]),digits = 5))

paste0("mean GFP/RFP for FIT:PCS= ",signif(mean(yeastfluorescence$greenredratio[yeastfluorescence$Translation.speed.size=="FIT:PCS"]),digits = 5))

```

###Figure S1: Translation speed at 3’ ends.

```{r,echo=F}

###Figure S1: Translation speed at C-termini.

initialtranslationspeed<-data.frame(read_excel(paste0(outputdatabasedirectory,"Initial Translation Speed Table 40.xlsx"),col_names = T),check.names = F,stringsAsFactors = F)

initialtranslationspeed<-initialtranslationspeed[initialtranslationspeed$Nucleotides>300,]

initialtranslationspeed$inverserrt.beginningvsrest<-log2((1/(initialtranslationspeed$AverageRRTFirst40nostartcodon/initialtranslationspeed$AverageRRTentireminusfirst40)))

initialtranslationspeed$inverserrt.endvsrest<-log2((1/(initialtranslationspeed$AverageRRTThreePrime40/initialtranslationspeed$AverageRRTentireminuslast40)))

genetemp<-read.fasta(paste0(inputdatabasedirectory,"orf_coding_R64-3-1_20210421.fasta"))

gene<-genetemp[initialtranslationspeed$Name]

rampzonelength<-40

threeprimecodonlist<-rep(list(NULL),(max(initialtranslationspeed$Nucleotides)/3)-2)

threeprimerrtlist<-rep(list(NULL),(max(initialtranslationspeed$Nucleotides)/3)-2)

generep<-1

while (generep<(length(gene)+1)){
  
  dnasequence<-getSequence(gene[[generep]])

  firstposition<-length(dnasequence)-5
  
  indexxxx<-1
  
  while (firstposition>3){
    
    threeprimecodonlist[[indexxxx]]<-append(threeprimecodonlist[[indexxxx]],toupper(c2s(dnasequence[firstposition:(firstposition+2)])))
    
    threeprimerrtlist[[indexxxx]]<-append(threeprimerrtlist[[indexxxx]],yeastcodonusage$RRT[yeastcodonusage$Codons==toupper(c2s(dnasequence[firstposition:(firstposition+2)]))])
    
    firstposition<-firstposition-3
    
    indexxxx<-indexxxx+1
  }

  generep<-generep+1
}

averagerrteveryposition3prime<-data.frame(codonposition=99999,count=99999,meanRRT=99999,index=length(threeprimerrtlist):1)

generep<-1

while(generep<(length(threeprimerrtlist)+1)){
  
  averagerrteveryposition3prime$codonposition[generep]<-generep
  
  averagerrteveryposition3prime$count[generep]<-length(unlist(threeprimerrtlist[generep]))
  
  averagerrteveryposition3prime$meanRRT[generep]<-mean(unlist(threeprimerrtlist[generep]))
  
  generep<-generep+1
}

averagerrteveryposition3prime$inversemeanrrt<-1/averagerrteveryposition3prime$meanRRT

yeastcodonusage$cumRRT<-yeastcodonusage$`Frame 1 (Coding) Observed Counts`*yeastcodonusage$RRT

globalrrt<-sum(yeastcodonusage$cumRRT)/sum(yeastcodonusage$`Frame 1 (Coding) Observed Counts`)

lineplot<-function(data){
  
  ggplot(data=data, aes(x=codonposition, y=inversemeanrrt))+
    
  geom_line()+
    
  geom_hline(yintercept = 1/globalrrt,linetype="dotted")+
    
  geom_vline(xintercept = rampzonelength,linetype="dotted")+
    
  labs(title="",x="Distance from Stop Codon",y="1/RRT")+
    
  theme(panel.grid.major = element_line(size = 0.5, colour = 'black'),
        
        panel.grid.major.x = element_line(size = 0.5, colour = NA),
        
        panel.grid.major.y = element_line(size = 0.5, colour = NA),
        
        panel.grid.minor = element_line(colour = NA),
        
        plot.title = element_text(hjust = 0.5,size = 15),
        
        axis.title.x = element_text(size=37),
        
        axis.text.x = element_text(family="sans", vjust=0.9,hjust=0.5,size=30,angle=0),
        
        axis.text.y = element_text(size=30),
        
        axis.title.y = element_text(size=37),
        
        panel.background = element_rect(fill = 'white', colour = 'black'))+
        
  scale_y_continuous(breaks = scales::pretty_breaks(n = 8),limits = c(0.94,0.990))+
  
  scale_x_continuous(trans = "reverse")
}

data2<-averagerrteveryposition3prime[1:200,]

lineplot(data = data2)

###Figure S1 statistics

print(paste0("###Figure S1 statistics"))

corr<-cor.test(averagerrteveryposition3prime$inversemeanrrt[1:200], averagerrteveryposition3prime$codonposition[1:200],method = "spearman")

corr

paste0("equation is  y= ",signif(lm(data2$inversemeanrrt~data2$codonposition)[["coefficients"]][[1]],digits=4)," + ", signif(lm(data2$inversemeanrrt~data2$codonposition)[["coefficients"]][[2]],digits=4),"x")

print(paste0("3' last 40 translation speed vs body p= ",signif(wilcox.test(initialtranslationspeed$AverageRRTThreePrime40,initialtranslationspeed$AverageRRTentireminuslast40,alternative="two.sided",paired=F)[["p.value"]],digits = 5)))

paste0("mean 3' translation speed of last 40 codons= ",signif(mean(initialtranslationspeed$AverageRRTThreePrime40),digits = 5))

paste0("mean 3' translation speed of body= ",signif(mean(initialtranslationspeed$AverageRRTentireminuslast40),digits = 5))

paste0("3' mean(log2(last40 codons/body))= ",signif(mean(initialtranslationspeed$log2Ratiothreeprime40vsRest),digits = 5))

###Statistics for the last 100 codons versus the rest of genes.

print(paste0("###Statistics for the last 100 codons versus the rest of genes."))

print(paste0("3' last 100 translation speed vs body p= ",signif(wilcox.test(initialtranslationspeed100$AverageRRTThreePrime100,initialtranslationspeed100$AverageRRTentireminuslast100,alternative="two.sided",paired=F)[["p.value"]],digits = 5)))

paste0("mean 3' translation speed of last 100 codons= ",signif(mean(initialtranslationspeed100$AverageRRTThreePrime100),digits = 5))

paste0("mean 3' translation speed of body= ",signif(mean(initialtranslationspeed100$AverageRRTentireminuslast100),digits = 5))

paste0("3' mean(log2(last 100 codons/body))= ",signif(mean(initialtranslationspeed100$log2Ratiothreeprime100),digits = 5))

```

###Figure S2: Distribution of translation speeds at 5’ and 3’ ends.

```{r,echo=F}

###Figure S2A: N-termini relative initial translation speed distribution.

initialtranslationspeed<-data.frame(read_excel(paste0(outputdatabasedirectory,"Initial Translation Speed Table 40.xlsx"),col_names = T),check.names = F,stringsAsFactors = F)

initialtranslationspeed<-initialtranslationspeed[initialtranslationspeed$Nucleotides>300,]

initialtranslationspeed$inverserrt.beginningvsrest<-log2((1/(initialtranslationspeed$AverageRRTFirst40nostartcodon/initialtranslationspeed$AverageRRTentireminusfirst40)))

initialtranslationspeed$inverserrt.endvsrest<-log2((1/(initialtranslationspeed$AverageRRTThreePrime40/initialtranslationspeed$AverageRRTentireminuslast40)))

lineplot<-function(data){
  
  ggplot(data=data, aes(x=inverserrt.beginningvsrest))+
    
  geom_histogram(colour="black",bins = 40)+
    
  labs(title = "",x="5' log2(Relative Initial Translation Speed)",y="Frequency")+
    
  theme(panel.grid.major = element_line(size = 0.5, colour = 'black'),
        
        panel.grid.major.x = element_line(size = 0.5, colour = NA),
        
        panel.grid.major.y = element_line(size = 0.5, colour = NA),
        
        panel.grid.minor = element_line(colour = NA),
        
        plot.title = element_text(hjust = 0.5,size = 15),
        
        axis.title.x = element_text(size=30),
        
        axis.text.x = element_text(family="sans", vjust=0.5,hjust=0.9,size=20,angle=90),
        
        axis.text.y = element_text(size=25),
        
        axis.title.y = element_text(size=35),
        
        panel.background = element_rect(fill = 'white', colour = 'black'))+
    
  scale_y_continuous(breaks = scales::pretty_breaks(n = 10),limits=c(0,800),expand=c(0,0))+
  
  scale_x_continuous(breaks = scales::pretty_breaks(n = 15),limits=c(-0.43,0.43))
}

lineplot(data=initialtranslationspeed)

print(paste("less than 0=",signif(((nrow(initialtranslationspeed[initialtranslationspeed$inverserrt.beginningvsrest<0,]))/length(initialtranslationspeed$inverserrt.beginningvsrest))*100,digits=5),"%  ","   mean=", signif(mean(initialtranslationspeed$inverserrt.beginningvsrest),digits = 5), "   range=",signif(min(initialtranslationspeed$inverserrt.beginningvsrest),digits = 5)," to ",signif(max(initialtranslationspeed$inverserrt.beginningvsrest),digits = 5)))

###Figure S2B: C-termini relative initial translation speed distribution.

lineplot<-function(data){
  
  ggplot(data=data, aes(x=inverserrt.endvsrest))+
    
  geom_histogram(colour="black",bins = 40)+
    
  labs(title = "",x="3' log2(Relative Terminal Translation Speed)",y="Frequency")+
    
  theme(panel.grid.major = element_line(size = 0.5, colour = 'black'),
        
        panel.grid.major.x = element_line(size = 0.5, colour = NA),
        
        panel.grid.major.y = element_line(size = 0.5, colour = NA),
        
        panel.grid.minor = element_line(colour = NA),
        
        plot.title = element_text(hjust = 0.5,size = 15),
        
        axis.title.x = element_text(size=30),
        
        axis.text.x = element_text(family="sans", vjust=0.5,hjust=0.9,size=20,angle=90),
        
        axis.text.y = element_text(size=25),
        
        axis.title.y = element_text(size=35),
        
        panel.background = element_rect(fill = 'white', colour = 'black'))+
    
  scale_y_continuous(breaks = scales::pretty_breaks(n = 10),limits=c(0,800),expand=c(0,0))+
  
  scale_x_continuous(breaks = scales::pretty_breaks(n = 15),limits=c(-0.43,0.43))
}

lineplot(data=initialtranslationspeed)

print(paste("less than 0=",signif(((nrow(initialtranslationspeed[initialtranslationspeed$inverserrt.endvsrest<0,]))/length(initialtranslationspeed$inverserrt.endvsrest))*100,digits=5),"%  ","   mean=", signif(mean(initialtranslationspeed$inverserrt.endvsrest),digits = 5), "   range=",signif(min(initialtranslationspeed$inverserrt.endvsrest),digits = 5)," to ",signif(max(initialtranslationspeed$inverserrt.endvsrest),digits = 5)))

```

###Figure S3: Codon speed and codon usage are correlated.

```{r,echo=F}

allorfcodonsfirst40<-unlist(fiveprimecodonlist[2:40])

allorfcodonsrest<-unlist(fiveprimecodonlist[41:length(fiveprimecodonlist)])

first40codons<-data.frame(table(allorfcodonsfirst40))

names(first40codons)[1]<-"Codons"

names(first40codons)[names(first40codons) == "Freq"] <- "Frequency"

first40codons<-join(first40codons,yeastcodonusage[,c("Codons","Amino Acid","RRT")],by="Codons",type="full", match="all")

restcodons<-data.frame(table(allorfcodonsrest))

names(restcodons)[1]<-"Codons"

names(restcodons)[names(restcodons) == "Freq"] <- "Frequency"

restcodons<-join(restcodons,yeastcodonusage[,c("Codons","Amino Acid","RRT")],by="Codons",type="full", match="all")

first40codons$first40Proportion<-first40codons$Frequency/sum(first40codons$Frequency,na.rm = T)

restcodons$restProportion<-restcodons$Frequency/sum(restcodons$Frequency)

### Figure S3A: correlation between codon usage and RRT across all ORF averaged across all positions. I took the inverse of RRT so the more right you go, the faster translation speed.

lineplot<-function(data){
  
  ggplot(data=data, aes(x=RRT, y=`Codon Proportion`))+
    
  geom_point(size=2.5)+
    
  geom_smooth(aes(color = NULL),method = "lm",se=T)+
    
  labs(title="", x=paste("1/RRT"),y="Global Codon Usage")+
    
  theme(panel.grid.major = element_line(size = 0.5, colour = 'black'),
        
        panel.grid.major.x = element_line(size = 0.5, colour = NA),
        
        panel.grid.major.y = element_line(size = 0.5, colour = NA),
        
        panel.grid.minor = element_line(colour = NA),
        
        plot.title = element_text(hjust = 0.5,size = 15),
        
        axis.title.x = element_text(size=40),
        
        axis.text.x = element_text(family="sans", vjust=0.9,hjust=0.5,size=37,angle=0),
        
        axis.text.y = element_text(size=37),
        
        axis.title.y = element_text(size=40),
        
        panel.background = element_rect(fill = 'white', colour = 'black'))+
    
  scale_y_continuous(expand=c(0,0),breaks = scales::pretty_breaks(n = 5), limits=c(0.0001,0.048))
}

data2=yeastcodonusage[!yeastcodonusage$`Amino Acid`=="*",]

data2$RRT<-1/data2$RRT

lineplot(data = data2)

corr<-cor.test(data2$RRT, data2$`Codon Proportion`,method = "spearman")

corr

paste0("Inverse RRT vs Codon Usage equation is  y= ",signif(lm(data2$`Codon Proportion`~data2$RRT)[["coefficients"]][[1]],digits=4)," + ", signif(lm(data2$`Codon Proportion`~data2$RRT)[["coefficients"]][[2]],digits=4),"x")

###Figure S3B: Correlation between first 40 codons at the beginning of genes/rest and average proportion across all ORF. 

first40vsrest<-join(first40codons,restcodons,by=c("Codons","Amino Acid","RRT"),type="full", match="all")

first40vsrest$beginningvsrestmeancodonfoldchange<-first40vsrest$first40Proportion/first40vsrest$restProportion

globalvssit<-join(yeastcodonusage,first40vsrest,by=c("Codons","Amino Acid","RRT"),match = "all",type = "full")

lineplot<-function(data){
  
  ggplot(data=data, aes(x=`Codon Proportion`, y=beginningvsrestmeancodonfoldchange))+
    
  geom_point(size=2.5)+
    
  geom_smooth(aes(color = NULL),method = "lm",se=T)+
    
  geom_hline(yintercept = 1,linetype="dotted")+
    
  labs(title="",x=paste("Global Usage"),y="Usage (2:40/Rest)")+
    
  theme(panel.grid.major = element_line(size = 0.5, colour = 'black'),
        
        panel.grid.major.x = element_line(size = 0.5, colour = NA),
        
        panel.grid.major.y = element_line(size = 0.5, colour = NA),
        
        panel.grid.minor = element_line(colour = NA),
        
        plot.title = element_text(hjust = 0.5,size = 15),
        
        axis.title.x = element_text(size=40),
        
        axis.text.x = element_text(family="sans", vjust=0.9,hjust=0.5,size=40,angle=0),
        
        axis.text.y = element_text(size=40),
        
        axis.title.y = element_text(size=40),
        
        panel.background = element_rect(fill = 'white', colour = 'black'))
}

data2=droplevels(globalvssit[!globalvssit$`Amino Acid`=="*",])

lineplot(data = data2)

corr<-cor.test(data2$`Codon Proportion`, data2$beginningvsrestmeancodonfoldchange,method = "spearman")

corr

paste0("Global Usage vs. Usage (2:40/Rest) equation is  y= ",signif(lm(data2$beginningvsrestmeancodonfoldchange~data2$`Codon Proportion`)[["coefficients"]][[1]],digits=4)," + ", signif(lm(data2$beginningvsrestmeancodonfoldchange~data2$`Codon Proportion`)[["coefficients"]][[2]],digits=4),"x")

###Figure S3C: Correlation between Sit/Rest and RRT.

lineplot<-function(data){ggplot(data=data, aes(x=RRT, y=beginningvsrestmeancodonfoldchange))+
    
  geom_point(size=2.5)+
    
  geom_smooth(aes(color = NULL),method = "lm",se=T)+
    
  geom_hline(yintercept = 1,linetype="dotted")+
    
  labs(title="",x=paste("1/RRT"),y="Usage (2:40/Rest)")+
    
  theme(panel.grid.major = element_line(size = 0.5, colour = 'black'),
        
        panel.grid.major.x = element_line(size = 0.5, colour = NA),
        
        panel.grid.major.y = element_line(size = 0.5, colour = NA),
        
        panel.grid.minor = element_line(colour = NA),
        
        plot.title = element_text(hjust = 0.5,size = 15),
        
        axis.title.x = element_text(size=40),
        
        axis.text.x = element_text(family="sans", vjust=0.9,hjust=0.5,size=40,angle=0),
        
        axis.text.y = element_text(size=40),
        
        axis.title.y = element_text(size=40),
        
        panel.background = element_rect(fill = 'white', colour = 'black'))
}

data2=first40vsrest[!first40vsrest$`Amino Acid`=="*",]

data2$RRT<-1/data2$RRT

lineplot(data = data2)

corr<-cor.test(data2$RRT, data2$beginningvsrestmeancodonfoldchange,method = "spearman")

corr

paste0("1/RRT vs. Usage (2:40/Rest) equation is  y= ",signif(lm(data2$beginningvsrestmeancodonfoldchange~data2$RRT)[["coefficients"]][[1]],digits=4)," + ", signif(lm(data2$beginningvsrestmeancodonfoldchange~data2$RRT)[["coefficients"]][[2]],digits=4),"x")

```

###Figure S4: Comparison of conservation scores at the N- and C-termini. 

```{r,echo=F}

###Figure S4: First 40 protein conservation scores vs last 40 protein conservation scores.

temp<-topbottomend.nona[,c("Name","Query.Annotation","topmiddle.protein.sumweightedscore")]

temp$zone<-"topmiddle"

names(temp)[names(temp) == "topmiddle.protein.sumweightedscore"] <- "value"

temp3<-topbottomend.nona[,c("Name","Query.Annotation","endconservation.protein.sumweightedscore")]

temp3$zone<-"end"

names(temp3)[names(temp3) == "endconservation.protein.sumweightedscore"] <- "value"

temp4<-rbind(temp,temp3)

lineplot<-function(data){
  
  ggplot(data=data, aes(x=value, fill=zone, alpha=zone))+
    
  geom_histogram(colour="black",binwidth=.5, position="identity")+
    
  labs(title = "",x="Protein Conservation Scores",y="Frequency")+
    
  theme(panel.grid.major = element_line(size = 0.5, colour = 'black'),
        
        panel.grid.major.x = element_line(size = 0.5, colour = NA),
        
        panel.grid.major.y = element_line(size = 0.5, colour = NA),
        
        panel.grid.minor = element_line(colour = NA),
        
        plot.title = element_text(hjust = 0.5,size = 15),
        
        axis.title.x = element_text(size=35),
        
        axis.text.x = element_text(family="sans", vjust=0.3,hjust=0.5,size=30,angle=0),
        
        axis.text.y = element_text(size=35),
        
        axis.title.y = element_text(size=35),
        
        legend.title= element_blank(),
        
        legend.text = element_text(size=30),
        
        panel.background = element_rect(fill = 'white', colour = 'black'),
        
        legend.key=element_rect(fill="white"),
        
        legend.position = c(0.5, 0.55),
        
        legend.direction = "horizontal",
        
        legend.spacing.x  = unit(0.01, 'cm'),
        
        legend.spacing.y = unit(0.5, 'cm'))+
    
        scale_alpha_manual(values=c(0.3,0.4))+
    
  guides(alpha = "none",fill = guide_legend(byrow = TRUE,override.aes = list(linetype=0,size=12,alpha = c(0.4,0.3))))+
  
  scale_y_continuous(breaks = scales::pretty_breaks(n = 10),limits=c(0,1300),expand=c(0,0))+
  
  scale_fill_manual(name="", breaks=c("topmiddle","end"), labels=c("First 40        ","Last 40"), values = c("#000000", "red"))
}

lineplot(data=temp4)

###First 40 vs Middle 40 vs Last 40

temp<-topbottomend.nona[,c("Name","Query.Annotation","topmiddle.protein.sumweightedscore")]

temp$zone<-"topmiddle"

names(temp)[names(temp) == "topmiddle.protein.sumweightedscore"] <- "value"

temp2<-topbottomend.nona[,c("Name","Query.Annotation","middlebottom.protein.sumweightedscore")]

temp2$zone<-"middlebottom"

names(temp2)[names(temp2) == "middlebottom.protein.sumweightedscore"] <- "value"

temp3<-topbottomend.nona[,c("Name","Query.Annotation","endconservation.protein.sumweightedscore")]

temp3$zone<-"end"

names(temp3)[names(temp3) == "endconservation.protein.sumweightedscore"] <- "value"

temp4<-rbind.fill(list(temp,temp2,temp3))

lineplot<-function(data){
  
  ggplot(data=data, aes(x=value, fill=zone,alpha=zone))+
    
  geom_histogram(colour="black",binwidth=.5, position="identity")+
    
  labs(title = "",x="Protein Conservation Scores",y="Frequency")+
    
  theme(panel.grid.major = element_line(size = 0.5, colour = 'black'),
        
        panel.grid.major.x = element_line(size = 0.5, colour = NA),
        
        panel.grid.major.y = element_line(size = 0.5, colour = NA),
        
        panel.grid.minor = element_line(colour = NA),
        
        plot.title = element_text(hjust = 0.5,size = 15),
        
        axis.title.x = element_text(size=35),
        
        axis.text.x = element_text(family="sans", vjust=0.3,hjust=0.5,size=30,angle=0),
        
        axis.text.y = element_text(size=35),
        
        axis.title.y = element_text(size=35),
        
        legend.title = element_text(color = "black", size = 60),
        
        legend.text = element_text(size=30),
        
        panel.background = element_rect(fill = 'white', colour = 'black'),
        
        legend.key=element_rect(fill="white"),
        
        legend.position = c(0.5, 0.55),
        
        legend.direction = "vertical",
        
        ###to increase the spacing between the legend symbol and text, change legend.spacing.x  = unit(0.01, 'cm')
        legend.spacing.x  = unit(0.01, 'cm'),
        
        ###to increase the spacing between the top and bottom legend items change legend.spacing.y = unit(0.5, 'cm'))
        
        legend.spacing.y = unit(0.5, 'cm'))+
  
  scale_alpha_manual(values=c(0.3,0.5,0.4))+
    
  guides(alpha = "none",fill = guide_legend(override.aes = list(linetype=0,size=12,alpha = c(0.4,0.5,0.3))))+
    
  scale_y_continuous(breaks = scales::pretty_breaks(n = 10),limits=c(0,1300),expand=c(0,0))+
    
  scale_fill_manual(name="", breaks=c("topmiddle", "middlebottom","end"), labels=c("First 40", "Middle 40","Last 40"), values = c("#000000", "green", "red"))
}

lineplot(data=temp4)

###Figure S4 statistics

print(paste0("###Figure S4 statistics"))

print(paste("mean of first 40 protein conservation scores = ",signif(mean(temp4$value[temp4$zone=="topmiddle"]),digits = 5)))

print(paste("mean of middle 40 protein conservation scores =",signif(mean(temp4$value[temp4$zone=="middlebottom"]),digits = 5)))

print(paste("mean of last 40 protein conservation scores =",signif(mean(temp4$value[temp4$zone=="end"]),digits = 5)))

print(paste0("first 40 amino acids vs middle 40 amino acids conservation ks.test p= ",signif(ks.test(temp4$value[temp4$zone=="topmiddle"],temp4$value[temp4$zone=="middlebottom"],alternative="two.sided",simulate.p.value = T, B = 5000)[["p.value"]],digits = 5)))

print(paste0("first 40 amino acids vs last 40 amino acids conservation ks.test p= ",signif(ks.test(temp4$value[temp4$zone=="topmiddle"],temp4$value[temp4$zone=="end"],alternative="two.sided",simulate.p.value = T, B = 5000)[["p.value"]],digits = 5)))

print(paste0("middle 40 amino acids vs last 40 amino acids conservation ks.test p= ",signif(ks.test(temp4$value[temp4$zone=="middlebottom"],temp4$value[temp4$zone=="end"],alternative="two.sided",simulate.p.value = T, B = 5000)[["p.value"]],digits = 5)))

```

###Figure S5: Slow 3’ Translation is correlated with poor C-terminal conservation.

```{r,echo=F}

###Figure S5A: C-termini relative initial translation speed as response variable and protein conservation as explanatory variable.

initialtranslationspeed<-data.frame(read_excel(paste0(outputdatabasedirectory,"Initial Translation Speed Table 40.xlsx"),col_names = T),check.names = F,stringsAsFactors = F)

initialtranslationspeed<-initialtranslationspeed[initialtranslationspeed$Nucleotides>300,]

initialtranslationspeed$inverserrt.beginningvsrest<-log2((1/(initialtranslationspeed$AverageRRTFirst40nostartcodon/initialtranslationspeed$AverageRRTentireminusfirst40)))

initialtranslationspeed$inverserrt.endvsrest<-log2((1/(initialtranslationspeed$AverageRRTThreePrime40/initialtranslationspeed$AverageRRTentireminuslast40)))

rampvsprotein<-join(topbottomend.nona,initialtranslationspeed,by=c("Name","Query.Annotation"),type="full", match="all")

rampvsprotein<-rampvsprotein[complete.cases(rampvsprotein), ]

size<-(round(nrow(rampvsprotein)/3))

rampvsprotein<-rampvsprotein[order(rampvsprotein$endconservation.protein.sumweightedscore,decreasing = T),]

top33percent<-rampvsprotein[1:(size+1),]

midbegin<-(round(nrow(rampvsprotein)/3))

middle33percent<-rampvsprotein[(midbegin+2):(midbegin+(size)),]

bottom33percent<-rampvsprotein[(midbegin+(size+1)):nrow(rampvsprotein),]

check1<-top33percent[top33percent$Name%in%middle33percent$Name,]

check2<-top33percent[top33percent$Name%in%bottom33percent$Name,]

check3<-middle33percent[middle33percent$Name%in%bottom33percent$Name,]

if(nrow(check1)==0&nrow(check2)==0&nrow(check3)==0){
  
  print("SUCCESS!!! NO DUPLICATES!!!!")
} else {
    
  base::stop(print("ERROR?? THERE ARE DUPLICATES??"))
}

if(nrow(top33percent)+nrow(middle33percent)+nrow(bottom33percent)==nrow(rampvsprotein)){
  
  print("SUCCESS!!! SPLIT INTO 3 PARTS")
} else {
    
  base::stop(print("ERROR?? SOME ARE MISSING??"))
}

rampvsprotein<-rampvsprotein[,!names(rampvsprotein)%in%c("size")]

rampvsprotein$size="ZZZZZ"

rampvsprotein$size[rampvsprotein$Name %in% top33percent$Name]<-"Top 33%"

rampvsprotein$size[rampvsprotein$Name %in% bottom33percent$Name]<-"Bottom 33%"

rampvsprotein$size[rampvsprotein$Name %in% middle33percent$Name]<-"Middle 33%"

rampvsprotein$size=ordered(rampvsprotein$size, levels = c("Bottom 33%","Middle 33%","Top 33%","Blank"))

tgrq<-summarySE(rampvsprotein,measurevar = "inverserrt.endvsrest",groupvars =c("size") )

means=tgrq

means$size=ordered(means$size, levels = c("Bottom 33%","Middle 33%","Top 33%"))

means$lower=means$inverserrt.endvsrest-means$ci

means$upper=means$inverserrt.endvsrest+means$ci

means=means[1:3,]

lineplot<-function(data){
  
  ggplot(data=data) +

  geom_col(data = data,aes(x = size,y = inverserrt.endvsrest),width = .75, position = "dodge")+
    
  geom_errorbar(data=means,aes(x = size,ymin=lower, ymax=upper,y=inverserrt.endvsrest),width=.4, color="black", alpha=1)+ 
    
  labs(title="", x="C-terminal Protein Conservation Score",y="3' log2(Relative Terminal Translation Speed)")+
    
  theme(panel.grid.major = element_line(size = 0.5, colour = 'black'),
        
        panel.grid.major.x = element_line(size = 0.5, colour = NA),
        
        panel.grid.major.y = element_line(size = 0.5, colour = NA),
        
        panel.grid.minor = element_line(colour = NA),
        
        plot.title = element_text(hjust = 0.5,size = 15),
        
        axis.title.x = element_text(size=30),
        
        axis.text.x = element_text(family="sans", vjust=0.9,hjust=0.5,size=35,angle=0),
        
        axis.text.y = element_text(size=28),
        
        axis.title.y = element_text(size=23),
        
        panel.background = element_rect(fill = 'white', colour = 'black'))+
    
  scale_y_continuous(breaks = scales::pretty_breaks(n = 10))
}

lineplot(data=means)

###Figure S5A statistics

print(paste0("###Figure S5A statistics"))

topvsbottomp<-wilcox.test(top33percent$inverserrt.endvsrest,bottom33percent$inverserrt.endvsrest,alternative="two.sided",paired=F)[["p.value"]]

print(paste0("topvsbottom p= ",signif(topvsbottomp,digits = 5)))

topvsmidp<-wilcox.test(top33percent$inverserrt.endvsrest,middle33percent$inverserrt.endvsrest,alternative="two.sided",paired=F)[["p.value"]]

print(paste0("topvsmid p= ",signif(topvsmidp,digits = 5)))

bottomvsmidp<-wilcox.test(bottom33percent$inverserrt.endvsrest,middle33percent$inverserrt.endvsrest,alternative="two.sided",paired=F)[["p.value"]]

print(paste0("bottomvsmid p= ",signif(bottomvsmidp,digits = 5)))

p.adjust.methods<-c("holm", "hochberg", "hommel", "bonferroni", "BH", "BY","fdr", "none")

cterminalbottopmid <- sapply(p.adjust.methods, function(meth) p.adjust(c(topvsbottomp,topvsmidp,bottomvsmidp), meth,n=3))

cterminalbottopmid

###Figure S5B: C-termini protein conservation as response variable and relative initial translation speed as explanatory variable.

size<-(round(nrow(rampvsprotein)/3))

rampvsprotein<-rampvsprotein[order(rampvsprotein$log2Ratiothreeprime40vsRest,decreasing = T),]

top33percent<-rampvsprotein[1:(size+1),]

midbegin<-(round(nrow(rampvsprotein)/3))

middle33percent<-rampvsprotein[(midbegin+2):(midbegin+(size)),]

bottom33percent<-rampvsprotein[(midbegin+(size+1)):nrow(rampvsprotein),]

check1<-top33percent[top33percent$Name%in%middle33percent$Name,]

check2<-top33percent[top33percent$Name%in%bottom33percent$Name,]

check3<-middle33percent[middle33percent$Name%in%bottom33percent$Name,]

if(nrow(check1)==0&nrow(check2)==0&nrow(check3)==0){
  
  print("SUCCESS!!! NO DUPLICATES!!!!")
} else {
    
  base::stop(print("ERROR?? THERE ARE DUPLICATES??"))
}

if(nrow(top33percent)+nrow(middle33percent)+nrow(bottom33percent)==nrow(rampvsprotein)){
  
  print("SUCCESS!!! SPLIT INTO 3 PARTS")
} else {
   
  base::stop(print("ERROR?? SOME ARE MISSING??"))
}

rampvsprotein<-rampvsprotein[,!names(rampvsprotein)%in%c("size")]

rampvsprotein$size="ZZZZZ"

rampvsprotein$size[rampvsprotein$Name %in% top33percent$Name]<-"STT"

rampvsprotein$size[rampvsprotein$Name %in% bottom33percent$Name]<-"FTT"

rampvsprotein$size[rampvsprotein$Name %in% middle33percent$Name]<-"MTT"

rampvsprotein$size=ordered(rampvsprotein$size, levels = c("STT","MTT","FTT"))

tgrq<-summarySE(rampvsprotein,measurevar = "endconservation.protein.sumweightedscore",groupvars =c("size") )

means=tgrq

means$size=ordered(means$size, levels = c("STT","MTT","FTT"))

means$lower=means$endconservation.protein.sumweightedscore-means$ci

means$upper=means$endconservation.protein.sumweightedscore+means$ci

means=means[1:3,]

lineplot<-function(data){
  
  ggplot(data=data) +

  geom_col(data = data,aes(x = size,y = endconservation.protein.sumweightedscore),width = .75, position = "dodge")+
    
  geom_errorbar(data=means,aes(x = size,ymin=lower, ymax=upper,y=endconservation.protein.sumweightedscore),width=.4, color="black", alpha=1)+
  labs(title="", x="3' log2(Relative Terminal Translation Speed)",y="C-terminal Protein Conservation Score")+
    
  theme(panel.grid.major = element_line(size = 0.5, colour = 'black'),
        
        panel.grid.major.x = element_line(size = 0.5, colour = NA),
        
        panel.grid.major.y = element_line(size = 0.5, colour = NA),
        
        panel.grid.minor = element_line(colour = NA),
        
        plot.title = element_text(hjust = 0.5,size = 15),
        
        axis.title.x = element_text(size=30),
        
        axis.text.x = element_text(family="sans", vjust=0.9,hjust=0.5,size=35,angle=0),
        
        axis.text.y = element_text(size=28),
        
        axis.title.y = element_text(size=23),
        
        panel.background = element_rect(fill = 'white', colour = 'black'))+
    
  scale_y_continuous(breaks = scales::pretty_breaks(n = 10),limits = c(0,41),expand=c(0,0))
}

lineplot(data=means)

###Figure S5B statistics

print(paste0("###Figure S5B statistics"))

sttvfttp<-wilcox.test(top33percent$endconservation.protein.sumweightedscore,bottom33percent$endconservation.protein.sumweightedscore,alternative="two.sided",paired=F)[["p.value"]]

print(paste0("sttvsftt p= ",signif(sttvfttp,digits = 5)))

sttvsmttp<-wilcox.test(top33percent$endconservation.protein.sumweightedscore,middle33percent$endconservation.protein.sumweightedscore,alternative="two.sided",paired=F)[["p.value"]]

print(paste0("sttvsmtt p= ",signif(sttvsmttp,digits = 5)))

fttvsmtt<-wilcox.test(bottom33percent$endconservation.protein.sumweightedscore,middle33percent$endconservation.protein.sumweightedscore,alternative="two.sided",paired=F)[["p.value"]]

print(paste0("fttvsmtt p= ",signif(fttvsmtt,digits = 5)))

p.adjust.methods<-c("holm", "hochberg", "hommel", "bonferroni", "BH", "BY","fdr", "none")

cterminalsttfttmtt <- sapply(p.adjust.methods, function(meth) p.adjust(c(sttvfttp,sttvsmttp,fttvsmtt), meth,n=3))

cterminalsttfttmtt

```

###End time elapse for generating all the figures.

```{r,echo=F}

timeended<-timelapseend.function(startyear=timebegin[[1]],startmonth=timebegin[[2]],startday=timebegin[[3]],starthour=timebegin[[4]],startminutes=timebegin[[5]],startseconds=timebegin[[6]],timestarted=timebegin[[7]],message.fin="COMPLETED FIGURES -")

print(timeended[[1]])

print(timeended[[2]])
  
```
